# Readers move their eyes mindlessly using midbrain visuo-motor principles

**DOI:** 10.1101/2021.10.21.465242

**Authors:** Françoise Vitu, Hossein Adeli, Gregory J. Zelinsky

## Abstract

Saccadic eye movements rapidly shift our gaze over 100,000 times daily, enabling countless tasks ranging from driving to reading. Long regarded as a window to the mind^1^ and human information processing^2^, they are thought to be cortically/cognitively controlled movements aimed at objects/words of interest^3–10^. Saccades however involve a complex cerebral network^11–13^ wherein the contribution of phylogenetically older sensory-motor pathways^14–15^ remains unclear. Here we show using a neuro-computational approach^16^ that mindless visuo-motor computations, akin to reflexive orienting responses^17^ in neonates^18–19^ and vertebrates with little neocortex^15,20^, guide humans’ eye movements in a quintessentially cognitive task, reading. These computations occur in the superior colliculus, an ancestral midbrain structure^15^, that integrates retinal and (sub)cortical afferent signals^13^ over retinotopically organized, and size-invariant, neuronal populations^21^. Simply considering retinal and primary-visual-cortex afferents, which convey the distribution of luminance contrast over sentences (visual-saliency map^22^), we find that collicular population-averaging principles capture readers’ prototypical word-based oculomotor behavior^2^, leaving essentially rereading behavior unexplained. These principles reveal that inter-word spacing is unnecessary^23–24^, explaining metadata across languages and writing systems using only print size as a predictor^25–26^. Our findings demonstrate that saccades, rather than being a window into cognitive/linguistic processes, primarily reflect rudimentary visuo-motor mechanisms in the midbrain that survived brain-evolution pressure^27^.

## Introduction

Saccades are a central component of vision in vertebrates with non-homogeneous retina, enabling high-resolution (foveal) sampling of the environment during ensuing eye fixations^20^. In humans, these eye movements provide the visual details necessary for performing complex cognitive tasks. An assumption, fueled by over a century of research in fields ranging from visual search to reading^2^, is that human oculomotor behavior is predominantly under top-down cognitive control^3–10^. The capacity of superior primates^27^ to shift gaze purposely towards desired target locations arises primarily from frontal and parietal brain areas^11–12^. However, the superior colliculus (SC), a phylogenetically older midbrain structure^15^, remains a key brain hub^13^ that relays most neo-corticofugal fibers to the brainstem premotor circuits^28^, but also integrates retinal and primary-visual-cortex afferents^13^ as in lower vertebrates^14^. The role of ancestral visuo-tectal tracts, besides driving reflexive saccades towards peripheral onsets^17–19^, remains unknown. Here we use an SC model^16^ to investigate the extent these faster pathways^13^ determine where humans move their eyes in a natural task, reading.

During reading, inter-saccadic intervals are particularly brief (averaging 225 ms^2^). This, together with visual-acuity limitations and letter crowding^29^, constrains information extraction from the periphery and creates conditions favoring default bottom-up eye-movement control. Such constraints have been largely ignored, despite evidence suggesting that peripheral word-identification processes are neither fast enough^30^ nor necessary^31–33^ to fully account for readers’ oculomotor behavior. Most theories/models presume that saccades are guided optimally by information acquired/processed during fixations (Extended Data Table 1). In predominant word-based models, they are programmed towards the center of target word(-object)s, as determined by ongoing word-identification processes^4,8^ and/or educated guesses/strategies combined with (coarse) peripheral preview^7,9^. These models however are explanatory and established on the questionable premise that readers’ preferential eye-fixation patterns in words/text are deliberate^34^. Moreover, they must assume substantial oculomotor errors, notably saccadic-range error (SRE; a bias to shift gaze a constant distance forward)^35^ to address unexplained behavioral variability, notwithstanding evidence against such bias^36^.

MASC, our Model of Attention in the Superior Colliculus^16^, predicts oculomotor behavior by spatially integrating luminance-contrast signals, as conveyed by retino(-geniculo-striate)-tectal tracts (Fig. 1a; see Methods). In the SC, spatial coding is distributed over populations of neurons (point images) with large and overlapping receptive/movement fields^37^. Moreover, overrepresentation of space closer to the fovea is offset by increasing response-field size with eccentricity, resulting in an invariant point-image size across the visual-field representation^21,38^. This implies that input signals are averaged over constant-size populations, which in turn causes saccades to be biased toward the fovea-weighted spatial centroid of peripheral configurations^38–40^. MASC implements these SC-averaging principles over visual- and motor-point images, and uses winner-take-all to determine each new fixation location, with an inhibitory spatial tag inserted after each to simulate inhibition of saccade return (ISR)^41^ – MASC’s one fit parameter compared to the many used by top-down models.

**Figure 1.**
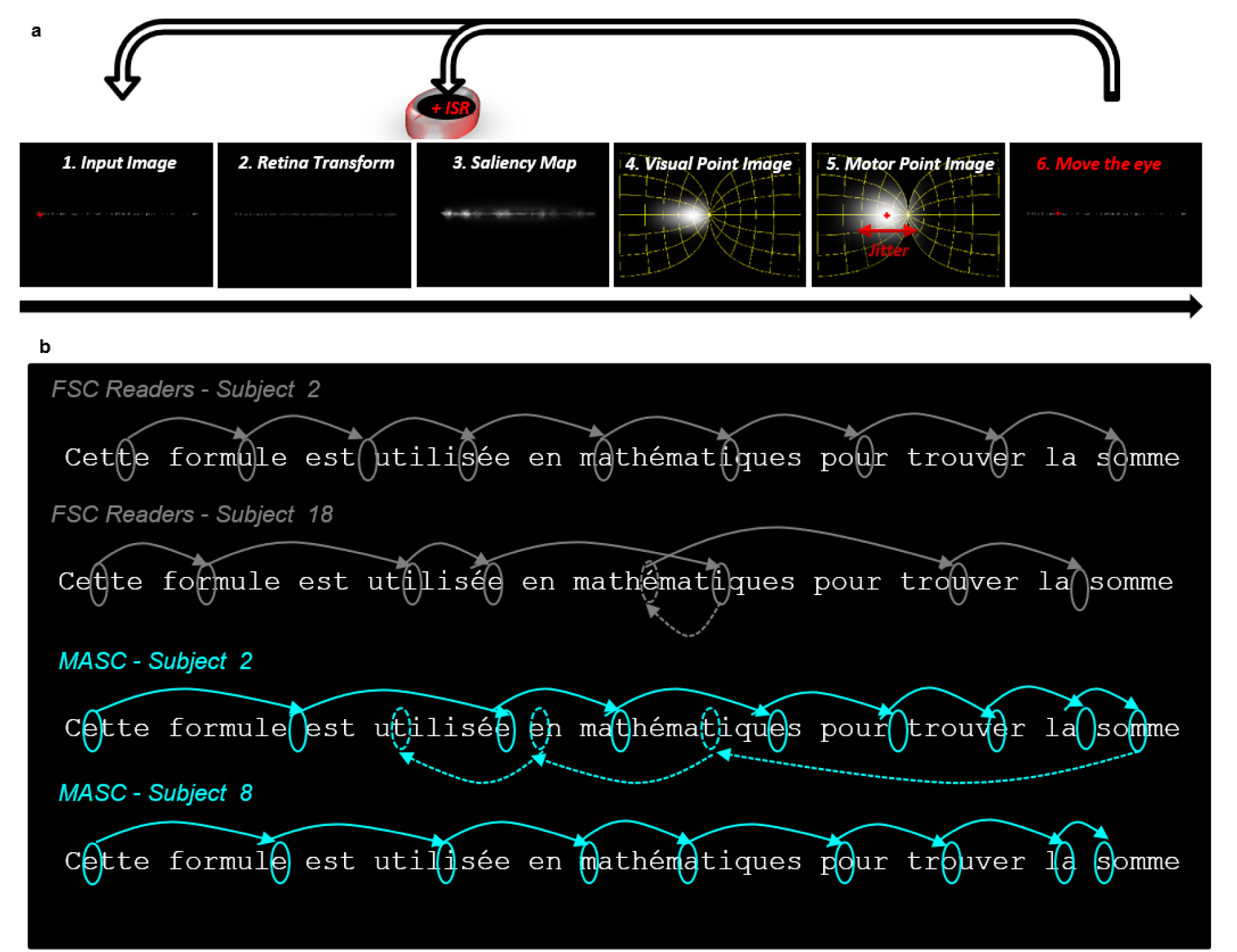
MASC’s main processing stages and its resulting reading-like behavior. **a,** On each fixation, the input image of the sentence (Panel 1) was blurred proportional to retinal eccentricity (Retina Transform –RT; Panel 2). A saliency map^22^ (distribution of oriented luminance contrast) was computed for the image (Panel 3), and then projected into SC space (Panel 4), taking into account the SC magnification factor^21^. Two cascaded averaging operations were made over translation-invariant visual- and (corresponding) motor-point images in SC space (Panels 4-5). A winner-take-all process identified the maximum population activity, and after jitter over the winning population (the horizontal red arrow in Panel 5) the location of the new fixation in visual space (Panel 6) was determined using inverse, efferent, mapping^21^. A reading scanpath was generated by repeating this process (upper horizontal arrows) but inserting after each saccade an inhibitory spatial tag (Inhibition of Saccade Return^41^; ISR) in the visual-saliency map at the fixated location (red circle above Panel 3). For further details see Methods. **b,** Example eye-movement patterns from FSC readers (Subjects 2 and 18; in grey) and MASC (Subjects 2 and 8; in cyan) over a randomly chosen sentence from the FSC corpus^31^.

## Results

### Readers’ oculomotor behavior is essentially mindless

Despite MASC being dumb, illiterate, and largely deterministic, its generated eye-movement behavior over sentences from the French-Sentence Corpus (FSC)^31^ was nearly indistinguishable from the behavior of humans reading the sentences for comprehension (Fig. 1b, Supplementary Videos 1-4). MASC mainly moved from left to right, though sometimes making regressive saccades^2^ (Fig. 2a). Compared to readers, MASC made more, and larger, regressions, but its forward saccades were just 1-letter (0.25°) shorter on average and as variable (Supplementary Tables 1-4).

**Figure 2.**
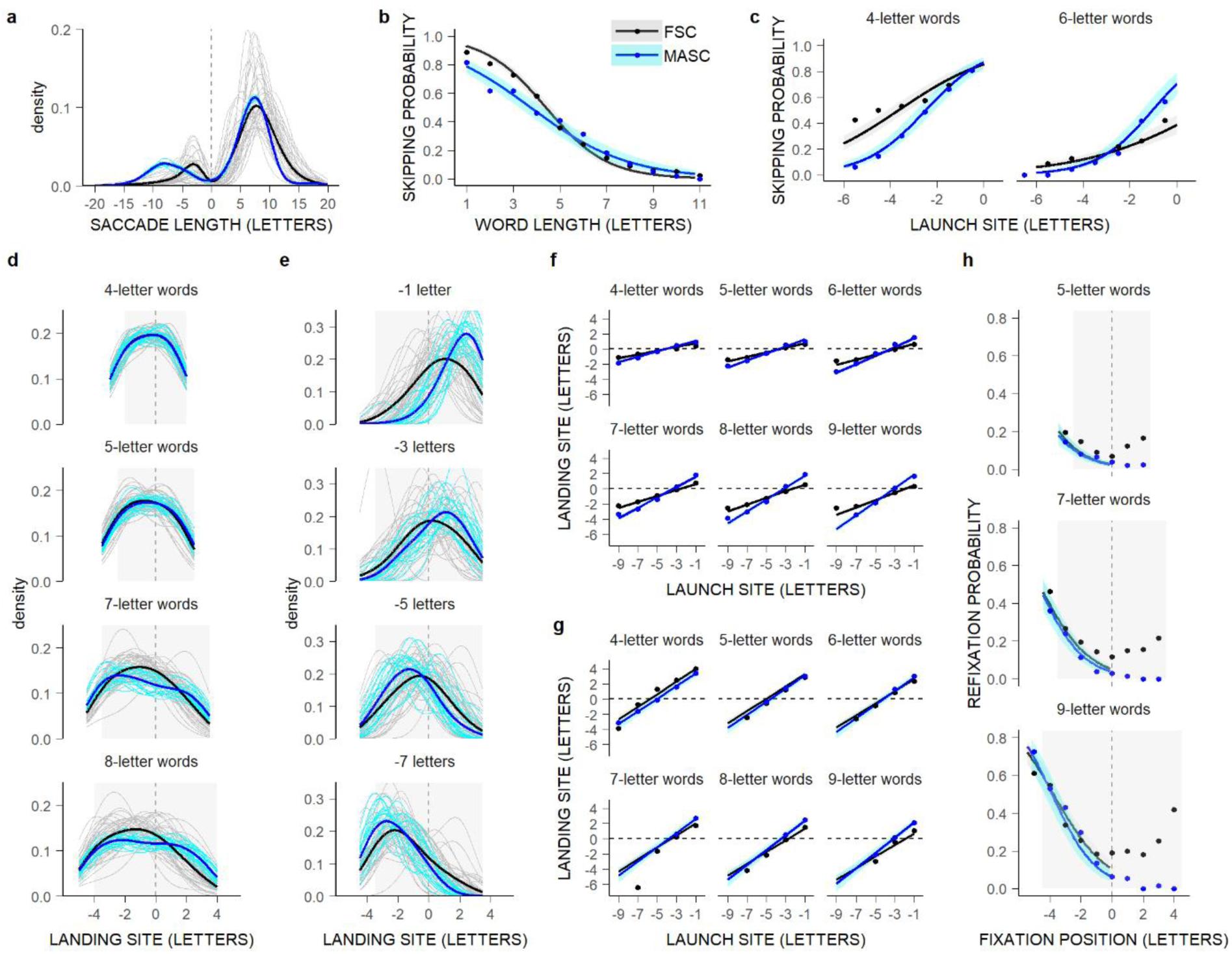
Illiterate visuo-motor principles in the SC account for prototypical word-based eye-movement behavior during reading. **a-h,** Comparison of the oculomotor behavior of MASC (blue/cyan) and FSC readers (black/grey), after matching both data sets for numbers of fixations (see Methods, Supplementary Methods 1). **a,** Probability density functions of saccade lengths (in letters) across and by subjects (thick and thin lines); positive/negative lengths: progressive/regressive saccades. **b,c,** Mean word-skipping probability (dots) as a function of word length (in letters; **b**), and for 4- and 6-letter words as a function of saccades’ launch-site distance to the space in front of the words (in letters; **c**, left and right panels), and partial effects (lines), with 0.95 confidence intervals (bands), computed from Generalized Linear-Mixed-effects Models (GLMMs; Supplementary Tables 5-6). **d,e** Probability density functions of within-word landing positions (in letters relative to the centers of words, represented by the vertical grey lines) across and by subjects for 4-,5-,7-, and 8-letter words (**d**), and for 7-letter words, separately for four launch-site distances (-1,-3,-5,-7 letters; **e**), representing PVL and launch-site effects respectively; grey-filled rectangle areas represent the words’ horizontal extent. **f-g,** Mean landing positions relative to the centers of 4- to 9-letter words (estimated by Gaussian-Mixture-Models (GMMs) fitted to individual landing-site distributions) as a function of launch-site distance, and partial effects, with 0.95 confidence intervals, computed from LMMs (Supplementary Tables 9-12); in **f**, within-word landing positions; in **g**, all landing positions, including also the endpoints of saccades falling short or landing beyond the words’ end^26,31^. **h,** Mean probability of refixating 4- to 9-letter words as a function of initial fixation location (in letters relative to the words’ centers), and partial effects, with 0.95 confidence intervals, computed from GLMMs for initial fixations in the first halves of words (Supplementary Table 13), representing the OVP effect. The small differences between MASC and FSC readers (**b-c**, **f-h**) are explained in Extended Data Fig. 1-3.

MASC also reproduced five prototypical forward eye-movement patterns taken as evidence for top-down guidance. Two relate to *word-skipping behavior,* readers’ tendency to more frequently skip words that are shorter^42^ and nearer to the saccades’ launch site^32^. Top-down models explain these patterns by arguing that shorter and less-eccentric words are (known to be) easier to process peripherally, making them less likely to be selected as the next saccade target^4–5,7–9^. MASC uses neither word-related knowledge nor top-down selection mechanisms, yet it predicted a reduction in skipping rate with both increasing word length and launch-site distance (Fig. 2b-c, Supplementary Tables 5-6). MASC skipped shorter and more eccentric words slightly less than FSC readers, but it skipped words as much as readers skipped rare words in their language^31^ and its behavior resembled humans viewing meaningless text^33^ (Extended Data Fig. 1). MASC therefore suggests that word-identification processes only mildly modulate word-skipping rate^31–32^.

Two other benchmark phenomena characterize the distributions of initial landing positions in words. The *Preferred-Viewing-Location (PVL) effect* refers to readers’ bias to fixate near the center of words, although closer to the words’ beginning as word length increases^43^. The *launch-site effect* refers to saccades landing further into words as they originate closer to the words’ beginning^35^. In word-based models, both phenomena reflect a word-center saccade-targeting strategy^7^ combined with SRE^4,8–9^. In other visual-span models, they result from eye-movement guidance toward the location minimizing uncertainty about the word being processed^5^. MASC does not compute word uncertainty, and it uses neither word-based saccade-targeting mechanisms nor SRE, yet it generated both PVL and launch-site effects. Its Gaussian-shaped landing-position distributions peaked near the center of 4-letter words, and shifted towards the beginning of longer words, as in FSC readers (Fig. 2d); only its landing positions in longer words showed more variability (Supplementary Tables 7-8). Moreover, as MASC’s saccades originated closer to the words’ beginning, they landed closer to the words’ end, or even beyond (Fig. 2e), yielding a linear relationship between launch-site distance and mean landing site. Slopes for MASC and FSC readers matched almost perfectly (Fig. 2f-g, Supplementary Tables 9-12). The tiny remaining differences, comparable in size to word-frequency effects, likely reflect language-related modulations of saccade amplitude^31^ (Extended Data Fig. 2).

Lastly, there is the *Optimal-Viewing-Position (OVP) phenomenon*, that corresponds to the increased likelihood of immediately refixating a word, particularly a long word, when the initial fixation location deviates from the word’s center^7^. Top-down models attribute this effect to word identification being (expected to be) less efficient when the eyes deviate from the words’ centers^4–5,7–9^. MASC is illiterate and generated no (regressive) refixations from the words’ end, yet still made more refixations when landing closer to the beginning of (longer) words, reproducing nearly perfectly the left wing of typically U-shaped OVP curves (Fig. 2h, Supplementary Table 13), like readers viewing meaningless text^33^ (Extended Data Fig. 3). This suggests that word-identification processes only partly contribute to the Refixation-OVP effect, accounting mainly for regressive (refixation) saccades. MASC indeed failed to generate regressions in additional benchmark conditions (Extended Data Fig. 4). It nevertheless captured regressions’ PVL effect (Supplementary Tables 14-15), thus indicating that these are programmed following the same visuo-motor principles as forward saccades.

### Mindless reading behavior reflects visual-saliency averaging in SC space

Our proposal that eye-movement guidance is essentially mindless is not completely new. However, researchers advancing this view assumed, unlike us, that either readers’ saccades are preprogrammed to move a constant distance forward regardless of encountered material^24,34,44^ (Extended Data Table 1), or that eye movements are guided by visual saliency alone^22^. Neither of these assumptions can predict reading behavior (Fig. 3). The constant-saccade-length model, besides not generating regressive saccades, lacks the variability in amplitude needed to predict most word-based phenomena. Even visual-saliency models using the same luminance-contrast distributions as MASC failed. They generated atypical distributions of saccade lengths, having landing positions biased towards word boundaries, skipping behavior inconsistent with word length, and a too-weak/strong (left-)OVP effect, all regardless of retinal transformation (RT; reduction in visual resolution with retinal eccentricity) or Gaussian averaging over the saliency map.

**Figure 3.**
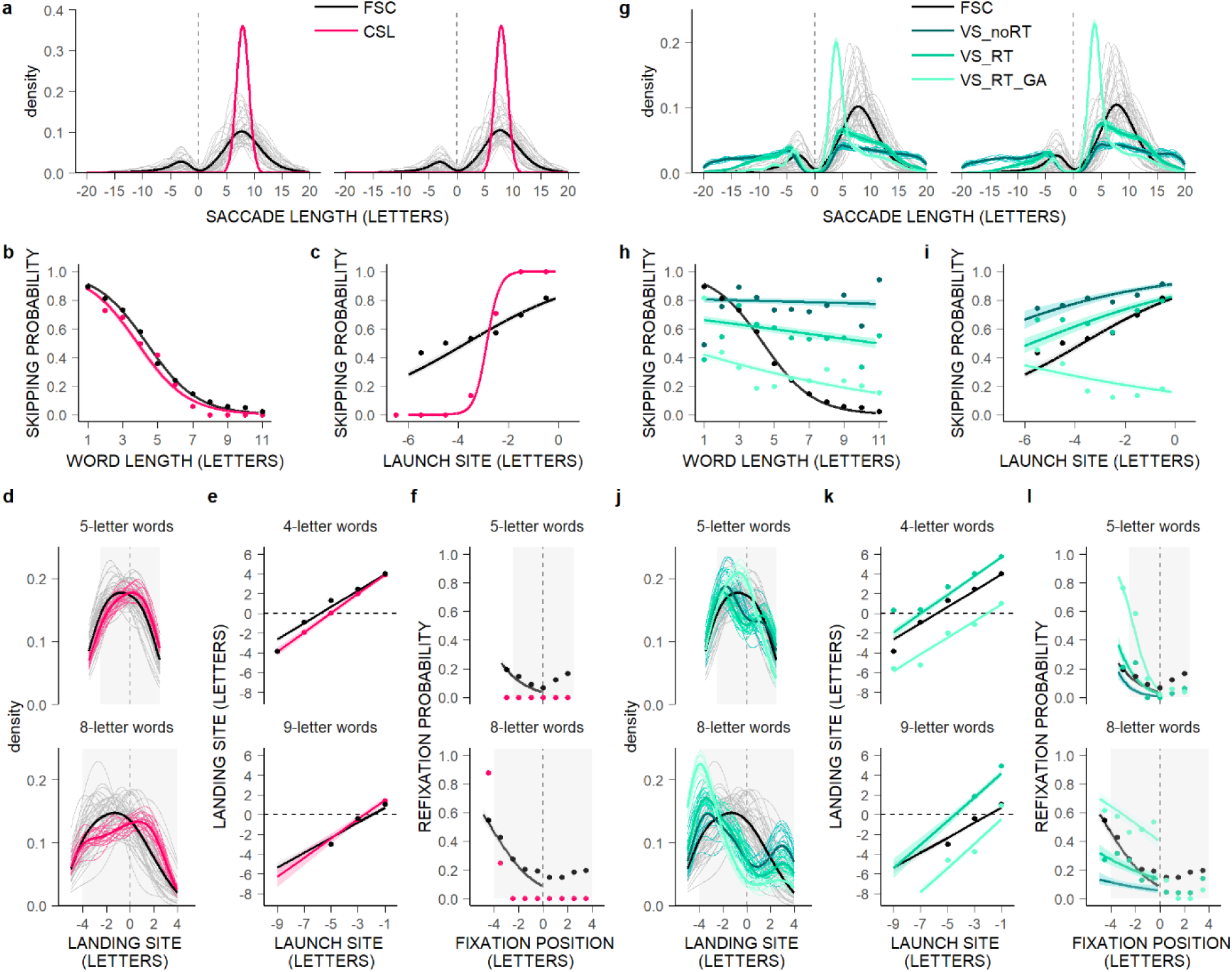
Constant-Saccade-Length and Visual-Saliency models fail to predict readers’ oculomotor behavior. **a-l,** Oculomotor behavior during the first pass over sentences for the Constant-Saccade-Length model (CSL in red; **a-f**), and for Visual-Saliency (VS) models (**g-l**) with or without RT (VS_RT and VS_noRT; medium and dark green) and Gaussian Averaging (VS_RT_GA; light green), compared to FSC readers (black) **–**see Methods, Supplementary Methods 2. **a,g,** Probability density functions of saccade lengths (in letters) across and by subjects (thick and thin lines); left panels: for comparison, for data sets matched for numbers of fixations (Fig. 2). **b-c,h-i,** Mean probability of word skipping (dots) as a function of word length (in letters; **b,h**), and for 4-letter words as a function of launch-site distance to the space in front of the words (in letters; **c,i**), and partial effects (lines), with 0.95 confidence intervals (bands), computed from GLMMs (Supplementary Tables 20-21). **d,j,** Probability density functions of within-word landing positions (in letters relative to the centers of words; vertical grey lines) across and by subjects for 5- and 8-letter words (the grey-filled rectangle areas). **e,k,** GMM-estimated mean of all landing positions, in letters relative to the centers of 4- and 9-letter words, as a function of launch site distance, and partial effects, with 0.95 confidence intervals, computed from LMMs (Supplementary Tables 26-27); VS_noRT was excluded because of a low *n* when data were split by word length and launch site. **f,l,** Mean within-word refixation probability as a function of initial fixation location (in letters relative to the centers of words) for 5- and 8-letter words, and partial effects, with 0.95 confidence intervals, computed from GLMMs but only for the left wing of OVP curves and excluding CSL which made zero refixations in 4- to 6-letter words (Supplementary Table 28).

These models crucially lacked cascaded averaging of luminance-contrast signals over translation-invariant visual- and motor-point images in SC space^16,21^ (Supplementary Tables 16-28), principles enabling prediction of readers’ stereotyped oculomotor behavior (although visual- or motor-only averaging already yielded reading-like behavior; Extended Data Fig. 5). MASC’s other processing stages (ISR, RT, population jitter; Fig. 1) contributed, but much less than spatial saliency averaging (Extended Data Fig. 6-7).

### Universals of reading behavior: Print size matters, but not inter-word spacing

A challenge when modeling reading behavior is to identify principles that generalize across many existing font types, print sizes, and text formats, as well as the world’s countless languages and writing systems. Existing models circumvented this challenge by taking letters as input, essentially agreeing that saccades are programmed in character coordinates regardless of print properties^2,7,44^ and that inter-word spacing, which enables fast text segmentation into (saccade-target) word(-object)s, is all that matters^4–5,7–9^ (Extended Data Table 1). However, these assumptions, specific to spaced Western-alphabetic languages, are controversial^23–24,26^ and imply that Eastern, alphabetic (Thai) and ideographic (Chinese/Japanese), scripts, that lack inter-word spacing, are read using less efficient word segmentation^45^ and/or different saccade-targeting strategies^45–46^, notwithstanding the universality of (most) word-based eye-movement patterns (Extended Data Fig. 8). MASC points a direction out of this impasse by suggesting that word segmentation is unnecessary, and inter-word spacing superfluous, for eye-movement guidance, and that most important is the spatial extent of the stimulus pattern(s), notably print size.

The assumption that *inter-word spacing* is crucial for eye-movement guidance rests on findings showing that readers make shorter forward saccades, and fixate slightly closer to the words’ beginning, when spaces in normally-spaced texts/sentences are removed or filled^2^. These behavioral changes are commonly attributed to increased difficulty in online peripheral-word segmentation/processing and saccade targeting. However, space-filling(/removal) is prone to confounds^23^ and effects at best speak to online foveal-word processing difficulty, being negligible when the one space following the fixated word is preserved regardless of peripheral linguistic content (Extended Data Fig. 9a-b). MASC lacks (foveal) word-identification processes, and therefore was largely unaffected by removing or filling inter-word spaces in FSC sentences (Fig 4a-f, Supplementary Tables 29-35). It also replicated the greater impact of space-removal compared to space-filling manipulations, showing this is due to space withdrawal making text narrower and consequently favoring shorter saccades^23^. MASC thus captures the minor role that inter-word spacing plays in online eye-movement guidance.

**Figure 4.**
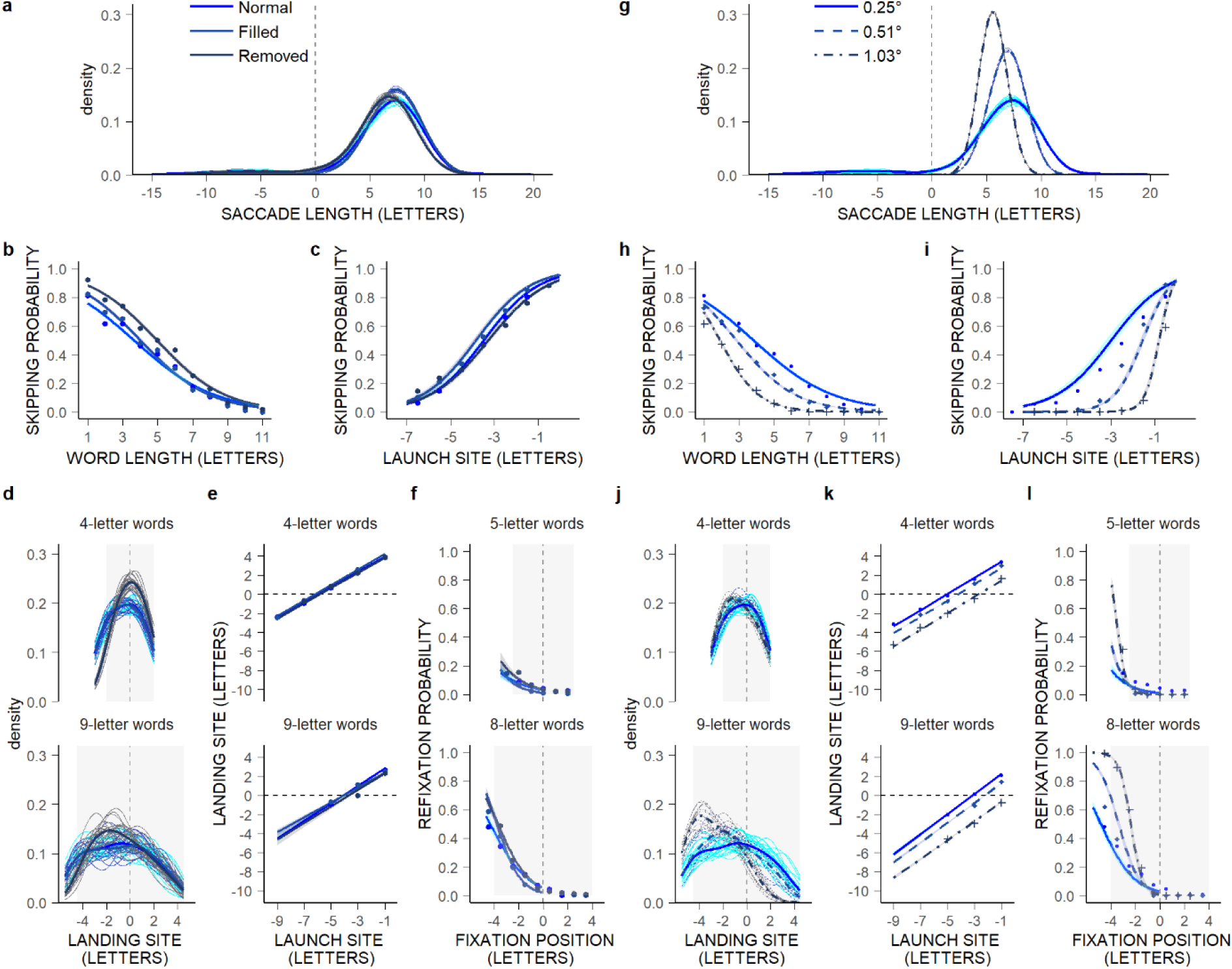
Illiterate visuo-motor principles in the SC reveal that the critical visual factor for eye-movement guidance during reading is character size, not inter-word spacing. Comparison of MASC’s oculomotor behavior over FSC sentences between three inter-word spacing conditions (**a-f)**, i.e., normal (original condition; blue), spaces removed (dark blue) and spaces filled (medium-dark blue), and three screen-width angles corresponding to three angular character sizes (**g-l)**, i.e., 0.25° (as in the original study), 0.51°, and 1.03° (solid, dashed, and dotted lines) – see Methods, Supplementary Methods 3. **a,g,** Probability density functions of saccade lengths (in letters) across and by subject (thick and thin lines). **b-c,h-i,** Mean probability of word skipping (dots) as a function of word length (in letters; **b,h**), and for 4-letter words as a function of launch-site distance (in letters relative to (**c**) the beginning of words and (**i**) the space in front of the words), and partial effects (lines), with 0.95 confidence intervals (bands), computed from GLMMs (Supplementary Tables 31-32, 37A,38). **d,j,** Probability density functions of within-word landing positions (in letters relative to the centers of words; vertical grey lines) across and by subjects for 4- and 9-letter words (grey-filled rectangle areas). **e,k,** GMM-estimated mean of all landing positions, in letters relative to the centers of 4- and 9-letter words, plotted as a function of launch-site distance, and partial effects, with 0.95 confidence intervals, computed from LMMs (Supplementary Tables 34, 40). **f,l**, Mean within-word refixation probability as a function of initial landing positions (in letters relative to the centers of words) for 5- and 8-letter words, and partial effects, with 0.95 confidence intervals, computed from GLMMs but only for the left wing of OVP curves (Supplementary Tables 35, 41).

*Character-print size*, unlike inter-word spacing, is thought to be unimportant for eye-movement behavior, due largely to a few influential studies reporting non-significant variations in the numbers of characters traversed with viewing distance (or angular print size)^2,7,44^. Yet, several studies reported significant effects of font size/type on the character count per saccade, and in all studies/languages, saccades’ angular extent increased with angular print size (Extended Data Fig. 9c-d). MASC, without any re-parametrization, predicted this relationship when tested on FSC sentences at three viewing distances. It also replicated changes in word-based behavior with increasing print size^25–26^: less word skipping (with increasing word length and eccentricity), more refixations (due to stronger OVP effect), and landing positions (much) closer to the beginning of (longer) words (Fig 4g-l, Supplementary Tables 36-41). MASC thus captures the major role played by character size, while revealing that readers’ saccades, rather than being aimed at specific (within-word) letter locations, are programmed to traverse angular distances regardless of letter/word units.

Inter-language comparisons indicate that, although *Chinese/Japanese readers* exhibit most word-based phenomena, they skip fewer words, refixate more words, and fixate preferentially the words’ first character(s) (Extended Data Fig. 8). Researchers attribute these patterns to a lack of inter-word spacing45, but MASC’s strikingly similar behavior over spaced French sentences when angular print size was multiplied by four suggests a simpler alternative–that these patterns result from Asian-language studies using character sizes two-to-four times greater and characters being the metric unit. Replotting word-skipping rate by the words’ angular extent erases differences between spaced-language studies using similar-sized fonts and shows *more* skipping for Chinese readers presented with larger characters, as predicted by MASC (Extended Data Fig. 9e). Relatedly, plotting landing-position distributions for comparable angular-sized words using angular-defined bins eliminates inter-language differences, revealing only a *weaker* PVL effect in large-printed (Chinese) words, consistent with MASC’s predictions (Extended Data Fig. 9f-g). MASC therefore evidences universal visuo-motor principles that generalize across spaced and unspaced languages, while raising crucial methodological issues.

## Discussion

Ancestral visuo-tectal tracts are classically regarded as purely reflexive pathways^17–18,47^. Their contribution to humans’ eye-movement behavior has been largely ignored under the common view that top-down cognitive control prevails^3–10^. Here we put these pathways back on center stage by generating reading-like oculomotor behavior over sentences using an SC model deprived of neocortical afferents^16^.

Readers move their eyes essentially forward, and in a stereotyped manner relative to word boundaries^2^, which existing models explain as top-down guidance to perceptually/lexically relevant locations combined with oculomotor errors/biases^4–5,7–9^. Our model captured these word-based phenomena, leaving mainly regressions-related behavior unexplained. This demonstrates that eye-movement behavior during reading is essentially mindless and only mildly modulated by cognitive/linguistic processes^31–32,40^. Relatedly, our study explains why readers’ oculomotor behavior over meaningless texts remains largely unperturbed^33^, which top-down models cannot explain without additional assumptions^4,8^. Top-down models spawned the belief that word(-object) segmentation is crucial for eye-movement guidance^4,7–9^ and consequently that Asian unspaced scripts are read more laboriously and/or differently compared to spaced texts^45–46^. We showed that the lack of inter-word spacing is not a limiting factor for oculomotor control^23–24^ –Chinese/Japanese readers behave differently simply because they were tested using much-larger print sizes than spaced-language readers. Character size matters^25–26^ but reading models taking letters as input ignore it. Our model replicated such evidence. It indicates that saccades during reading are programmed in visual-space coordinates using universal visuo-motor principles.

The principles we isolated involve luminance-contrast extraction in retina and V1, but the crucial step is visual-saliency averaging over constant-size visual- and motor-point images in SC space^21,38^. This hallmark SC visuo-motor transformation, estimated from macaque data, is what enabled our model, unlike visual-saliency models^22^ deprived of retino/cortico-tectal projections, to reproduce readers’ oculomotor behavior. This is also why our model outperformed scene-viewing models in a previous study^16^. Here we predicted fundamental word(/-object)-based eye-movement properties, that generalize to non-reading tasks^33,48^ and are already present in first-grade readers^49^. This suggests, in line with early maturation of visuo-tectal tracts/computations^19,47^, that visual-saliency averaging in SC space is an inborn principle determining where, by default, primates move their eyes regardless of task. Slower cognitive/attentional control^13,40^, essentially via descending projections to the SC^28^, intervenes secondarily by modulating this default neuronal-activity pattern^39–50^, all depending on peripheral-processing speed and fixation duration and hence stimulus and task.

Phylogenetic brain reorganization afforded humans with superior attention- and oculomotor-control systems^27,47^, but this did not lessen the role of ancestral visuo-motor pathways^14–15,17–18^. We established a baseline of midbrain eye-movement control during reading against which (universal) cognitive/linguistic processes/influences can now be properly studied. This baseline should inform reading education policy and provide a biomarker of visuo-motor deficits in clinical applications (low-vision, dyslexia, etc.). Future research will extend our approach to other tasks and species, to further understand the complex interplay between bottom-up and top-down oculomotor control.

## Methods

### MASC implementation

MASC is a neuro-computational model that takes pixels as input (i.e., image-based). Here it was nearly identical to MASC applied to the free viewing of natural scenes^16^. MASC predicted each new fixation location over sentences by going through the following sequence of processing stages (Fig. 1a): (1) *RT*, the blurring of the input image to simulate the gradual reduction in visual resolution with increasing retinal eccentricity (svistoolbox-1.0.5 Space Variant Imaging System; http://svi.cps.utexas.edu/software.shtml)^51^; (2) *Computation of a priority (*here a *visual-saliency) map* based on extraction of feature (hue, luminance and orientation) contrast at different spatial scales (GBVS toolbox; http://www.vision.caltech.edu/~harel/share/gbvs)^52^; (3) *Projection of this saliency map into SC space*, i.e., a two-dimensional array of retinotopically arranged and equally-spaced visually-responsive neurons with large receptive fields (as in superficial and intermediate SC layers^53–55^), which, due to the non-homogeneity of afferent projections, produces an overrepresentation of space closer to the fovea^21^; (4) *Cascaded averaging*^56^ of resulting activity over translation-invariant neuronal populations (or point images^57–60^), first in the visual map and then in a spatially-registered motor map^61^, implemented here by the projection of averaged visual activity onto a topographic layer of equally-spaced neurons having large movement fields (as in the intermediate and deeper SC layers^55,62–63^); (5) *Winner-take-all process* to identify the most active motor population; (6) *Location jitter*^64^ applied to the winning population (the only step not in MASC’s free-viewing version); (7) *Conversion back to visual space*, using an inverse efferent mapping^21^ to determine the next fixation location; and (8) *ISR*^41^, here defined as inhibition injected into the saliency map to prevent returning to image locations that were already fixated.

As further detailed in our original model paper^16^, projection of the visual-saliency map into SC space was done using an anisotropic logarithmic afferent-mapping function, as estimated in the monkey^21^. The diameter and sigma of the Gaussian window used for computation of visual- and motor-point images were fixed and estimated directly from monkey electrophysiological data^65^. Population-location jitter was rotation-symmetrical and had a sigma and diameter corresponding to ∼13% of motor-point images’ sigma and diameter, as previously estimated based on saccade-endpoint scatter in humans^64^. Given the SC magnification factor, this meant that larger saccades were more variable in size, as shown in saccade-targeting tasks^38,64^ and as also reported during reading^35^. Both the width and sigma of the ISR window were adjusted, by testing a range of diameter (1.07°-2.12°) and sigma (0.22°-0.90°) values (Extended Data Fig. 6g-l). The parameter pairs yielding the most reasonable fit of the observed distribution of saccade lengths were first selected. Then, the one yielding the best fit of word-skipping behavior, and PVL, launch-site, and refixation-OVP effects, was retained. The selected ISR window, used for all the simulations, had a diameter of 1.82° and a sigma of 0.37°, corresponding to 7.28 and 1.48 letters subtending 0.25° each.

### MASC dissection and model comparison

To determine the contribution of each processing step in MASC’s behavior, we first implemented six amputated versions of the model, each containing all of MASC’s processing steps, except for: (1) RT (MASC_noRT), (2) averaging over motor-point images (MASC_VISUAL), (3) averaging over visual-point images (MASC_MOTOR), (4) averaging over both visual- and motor-point images, in which case MASC turned into a pure Visual-Saliency (VS) model with RT (VS_RT), (5) averaging over both visual- and motor-point images and RT (VS_noRT), or (6) jitter over the winning population (MASC_noJITTER). Additionally, to estimate the contribution of cascaded averaging over translation-invariant visual- and motor-point images in SC space, we implemented four additional VS_RT models that applied Gaussian averaging (GA) directly to the saliency map (VS_RT_GA1-4) using windows of variable diameter and sigma (0.31°, 0.15°; 0.62°, 0.30°; 1.22°, 0.60°; 2.42°, 1.20°). Since the first three models gave results that were either similar to VS_RT or somewhere in between VS_RT and VS_RT_GA4, only the simulation results for VS_RT_GA4 are reported; this is referred to as VS_RT_GA.

Finally, we estimated the contribution of long-lasting ISR in MASC, using two additional model variants, one with ISR applied only to the current fixation (MASC_ISR_C) and another with ISR applied to both the current and the immediately prior fixations (MASC_ISR_1PC). Moreover, we implemented a Constant-Saccade-Length (CSL) model, one making exclusively forward saccades of nearly constant amplitude (1.75°, or 7 letters, the mean length of MASC’s forward saccades –Supplementary Table 4, with Gaussian noise of diameter 0.21° and sigma 0.105°). This allowed us to assess whether visual input is at all necessary to predict readers’ eye-movement behavior, while providing a definitive test of the previously proposed saccade-preprogramming hypothesis^24,44,66^.

### The French Sentence Corpus (FSC)

The FSC, created to investigate the influence of visual and linguistic variables on eye movements during reading, comprised a total of 316 pairs of one-line sentences read silently by 40 French-native adults whose eye movements were recorded with a Dual-Purkinje-Image Eye-Tracker (Ward Technical Consulting)^31^. The two sentences of a pair differed by a single word (the second word), that was either semantically related or unrelated to a following test word of variable frequency and length. The total set of 632 sentences was split into two lists of 316 sentences, each containing only one exemplar of a sentence pair, and an equal number of predictable and unpredictable sentences. Each participant saw only one list, and hence only one exemplar of each sentence pair, but all sentences were seen across all participants (Latin-Square Design). Note that, as in a main series of analyses of the original FSC study, all words in the sentences (that corresponded to our selection criteria – see Data Selection and Analyses), and not only the test words, were considered for analysis; this increased the number of observations per cell, and hence statistical power, without changing observed eye-movement patterns^31^. The properties of sentences and words are detailed in the original paper^31^.

Sentences, saved as bitmaps, were displayed one at a time on a gamma-corrected 21” CRT monitor, at a screen resolution of 1280×960 pixels. Each sentence appeared on the vertical midline of the screen, with its second character aligned with a previously displayed fixation bar in the left part of the screen. Each character space subtended 0.25 degrees of visual angle at a distance of 118 cm from the participants’ eyes. Each sentence remained on screen until the participant pressed a button, thus allowing sentence rereading at will. Comprehension was enforced by semantic-content questions presented randomly after 20% of the sentences (96% correct responses on average).

All participants in the FSC study gave their written informed consent prior to their participation in the experiment, that was conducted in accordance with the ethical standards laid down in the Declaration of Helsinki. This research was approved by the committee responsible for overseeing research conducted in human subjects at Aix-Marseille University (Comité d’éthique de l’université d’Aix-Marseille; Pierre-Jean Weiller, President).

### Model simulations

Both lists of 316 sentences from the FSC were input ten times to all models in our comparison set, except MASC_noJITTER (where multiple inputs were unnecessary), thus yielding a total of 20 runs per model. For each sentence, a given model generated saccades to bitmap locations until: (1) the buildup of ISR emptied activity on the visual-saliency map (for MASC and VS models), (2) there were less than about seven characters to the right of fixation (for CSL), or (3) a maximum of 20 fixations was reached. This 20-fixation termination criterion was determined empirically based on the number of fixations per sentence in FSC readers (mean: 11.63; 4.07-26.82), which was distributed normally when the few occurrences with more than 20 fixations (7.1% on average), typically associated with eye blinks and/or false tracks, were excluded. It was an upper bound ensuring that simulated and observed data sets could be matched for numbers of fixations or at least first-pass behavior over sentences (see Data selection and analysis). Accordingly, most models generated on average more fixations per sentence than FSC readers (MASC_IOR_1PC, MASC_IOR_C: 20; MASC, MASC_noRT, MASC_noJITTER: 19.99; MASC_MOTOR: 18.18; VS_noRT: 17.00; VS_RT: 16.03; MASC_VISUAL: 15.76). VS_RT_GA and CSL still made fewer fixations on average (10.47 and 6.81 respectively).

For the main set of simulations, the screen width angle was set to 20°, such that each character subtended about 0.25 degree of visual angle, as for FSC readers. However, to explore the role of character size, two additional width angles were tested (40° and 80°), so that each character subtended about 0.51° and 1.03° respectively. Additionally, to determine MASC’s predicted effect of inter-word spacing, FSC bitmaps were regenerated after removing or filling with x’s inter-word spaces in the corresponding sentences.

### Data selection and analysis

Simulated and observed oculomotor behavior were compared across the two lists of sentences from the FSC corpus. MASC was first opposed to FSC readers only to keep the comparison simple and directly test whether MASC predicted readers’ oculomotor behavior (Supplementary Methods 1). Then, comparison models were opposed to MASC and FSC readers to identify MASC’s critical processing steps (Supplementary Methods 2). Because the numbers of fixations per sentence differed between MASC and FSC readers and between MASC and other data sets, we implemented two different data-matching procedures respectively. The first procedure, for comparison between MASC and FSC readers, matched data sets for numbers of fixations. For a given sentence and model run, the number of fixations considered for analysis was determined by randomly sampling from the distribution of the numbers of fixations per sentence for FSC readers in the corresponding sentence pair (but excluding marginal trials with more than 20 fixations –see Model Simulations). The second procedure, used for model comparison, matched data sets for behavior by selecting the fixations made during the first pass over a sentence (i.e., all fixations from the start of reading a sentence until a regression or button press following the first eye pass on the rightmost fixated word). Compared to random sampling, this procedure more greatly reduces the number of fixations considered for analysis, but it allows comparison of the oculomotor behavior over a sentence before re-reading, regardless of how many fixations were necessary to achieve this behavior; it also allows fairer comparison with CSL, which made fewer fixations and never generated regressions.

For both comparison sets, exclusion criteria from the original FSC study^31^ were applied to the data. Specifically, fixations were excluded if they were (1) preceded or followed by an eye blink or other signal irregularity (which biases estimation of the fixation location; for FSC data only), (2) more than 1° above or below the screen midline where the sentence was displayed (and possibly unrelated to sentence reading), (3) preceded by a fixation more than 1° above/below the midline, (4) the last fixation on the line (biased by subsequent button press), or (5) preceded by a fixation that was the first fixation on the line (biased by fixation behavior on the prior fixation stimulus).

In saccade-length analyses, we measured the horizontal amplitude and direction of the saccade immediately preceding a selected fixation. Saccades launched from either the first or the last word in the sentence were excluded from analysis so as not to bias estimations of regression rate and forward/regressive saccade length; a few saccades greater than +/-20 letters were also identified and excluded. In word-based analyses, we measured the location of the selected fixation relative to the boundaries of a given critical word, either: (1) the word immediately to the right of the word from which the prior saccade was launched in both word- skipping and overall landing-position analyses (thus measuring whether the fixation was beyond the word’s end, and where it was located relative to the word’s center, respectively), (2) the fixated word in within-word landing-position analyses, or (3) the word the prior saccade was launched from in refixation-probability analyses (thus measuring whether the fixation remained on the word). Instances when the critical word was either the first or the last word in the sentence, or a word preceded or followed by punctuation, were excluded to avoid screen- border and beginning/end-sentence effects as well as underestimation of visual word length. Furthermore, to restrict our analyses to first-pass behavior on words (as classically done), cases were rejected when the critical word was previously fixated and the critical fixation in word-skipping and landing-position analyses, or the fixation prior to the critical fixation in refixation analyses, was neither a fixation preceded by a forward saccade nor the very-first fixation on a word. The number of cases that remained after these selections varied depending on the analysis and is reported in the Supplementary Tables’ legends.

### Gaussian-mixture modeling of saccade-length and landing-position distributions

Saccade-length and landing-position distributions were first visualized by plotting for each data set, individual and condition, corresponding probability density functions, with fixed 1-letter (0.25°) bandwidth and Gaussian kernel. Since several of the distributions had several modes, GMMs were first fitted to the data, using the *mclust* package (Version 5.2)^67^ in R (Version R-3.1.3)^68^. These provided an estimate of the number of mixture components in each of the distributions, as well as an estimate of the mean, variance, and proportion of cases (“*k*”) for each detected mode.

GMMs searched for 1 to 4 and 1 to 3 mixture components (“*G*” parameter) in saccade-length and landing-position distributions respectively; these numbers of components were motivated by the shape of the most irregular distributions, those generated by VS models. To optimize GMM fitting, we used a prior having three parameters: mean, scale, and shrinkage^67^. For a given data set, individual and condition, the mean and scale parameters were fixed. The mean parameter corresponded to the default-prior mean, that is the mean of saccade lengths or landing positions for this data set, individual and condition, unless this prevented an optimal fit; For saccade-length distributions, the mean parameter was set to an extreme negative value (-25 letters) to capture the often very-small mode associated with regressive saccades. The scale parameter, which was defined separately for each tested *G* value, corresponded to the default-prior scale, meaning the variance in saccade lengths, or landing positions, in the data set, individual and condition, divided by squared *G*. The shrinkage parameter for the prior on the mean was tuned over a range of shrinkage values (from 0.01, the default prior-shrinkage value, to 18).

Selection of the optimal model for a given data set, individual and condition was done in three steps. First, models were excluded for having more than one mixture component if: (1) the difference between the estimated means of adjacent modes was less than (or equal to) a given threshold (4.5 and 3.2 letters in saccade-length and landing-position analyses respectively), (2) there was no mixture component with a negative mean (only in saccade-length analyses and for the data sets containing regressive saccades, thus not CSL), or (3) the *k*-value of one of the mixture components was not greater than 0.3 (only in landing-position analyses). These empirically determined selection criteria reflect a compromise to capture the bi(tri)modality in the VS-models’ distributions and to reproduce two well-established findings from the literature, also present in FSC readers: (1) that saccade-length distributions are typically bimodal (with both a negative and a positive peak associated respectively with regressive and progressive saccades)^2^, and (2) that landing-position distributions are typically unimodal^35,43^. Moreover, *k* was set to a value greater than 0.3 in landing-position analyses to ensure that a given mixture component contained a reasonable minimal number of observations; these analyses, particularly when data were split by word length and launch site, relied on a much lower *n*. Conversely, having no *k*-threshold in saccade-length analyses meant that the often very small proportion of regressive saccades would be modeled. Second, the model with the maximal BIC (Bayesian Information Criterion^69^) value across the tested range of shrinkage-parameter values was selected separately for each G-value. Third, the model that was retained had by default two components in saccade-length analyses and a single component in landing-position analyses, unless there was strong evidence that a more complex (or a simpler) model better fitted the data, meaning that the BIC value was greater than that associated with the default model and the difference in BIC values was greater than 6^70^.

The distributions were then compared between data sets (and conditions), using parameter estimates from the corresponding optimal GMM models (Supplementary Methods 1-2, Supplementary Tables 1-4,7-12,14-19,22-27). First, to assess whether MASC and other comparison models reproduced readers’ typically unimodal and bimodal distributions of landing positions and saccade lengths respectively, two indexes were compared: (1) the proportion of 1-3 and 1-4 mixture components respectively, and (2) the proportion of cases belonging to the largest mixture component (i.e., the value of the highest *k* estimate; 1 in unimodal distributions) or, for saccade-length distributions, the ratio between the highest *k* estimate and the sum of *k* estimates separately for negative and positive modes. These first comparisons yet remained descriptive due to these indexes showing floor or ceiling effects in several data sets. However, they were completed, for saccade-length distributions, by statistical comparisons of the regression rate, estimated by summing all *k* estimates associated with a negative mode. Moreover, to determine whether the main part of the distributions was aligned between data sets and conditions, and showed comparable spread, both the mean and the standard deviation (SD) of the largest mixture component (that with the highest *k* value) were submitted to statistical tests, but separately for regressive and progressive saccades in saccade-length analyses.

### (Generalized)#Linear (Mixed Effect) modelling

To statistically compare the behavior of FSC readers and MASC, and to determine whether MASC outperformed the other models in our comparison set, (G)LMMs were fit to the data using the *(g)lmer* functions in the *lme4* package (Version 1.1-7)^71^ in R (Version R-3.1.3)^68^. GLMMs are logistic models that fit the probability distribution of binary data. Here, they were used to estimate word-skipping and within-word refixation rates. LMMs were fit to the GMM-estimated mean and SD of the largest mixture component in landing-position analyses. However, for GMM-estimated mean and SD of saccade lengths, as well as the proportion of regressive saccades, (G)LMs were fit to the data, because there was only one observation per subject.

(G)LMMs were implemented after checking the linearity of the relationship between each dependent variable and its predictors, leading us to remove extreme predictor values that were associated with a low *n* or yielded floor/ceiling effects. The default random structure included a random intercept by subject (and by sentence pair and word in word-skipping and refixation analyses) and random effects of all explanatory variables (except Data Set) by subject. If the model did not converge, simpler random structures were tested until convergence was attained: first, the random intercept by word, and then by sentence pair, was removed, and then random effects by subjects were progressively removed, but each time after testing the model with and without the correlation between random effects. The fixed structure included Data Set as a categorical predictor and other explanatory variables, as well as all interactions. When possible, explanatory variables were entered as continuous variables centered on their mean.

(G)L(M)M estimates are presented in Supplementary Tables 3-6,8,10,12-13,15,17-21,23,25,27-28, with the models’ random and fixed structures in the tables’ legend; fixed effects are described and commented in Supplementary Methods 1-2. The exact number of degrees of freedom for the t-values of fixed effects in LMMs remains undetermined. However, given the large number of observations, subjects, and items entering our analyses, t-distributions converged to a normal distribution. Therefore, we considered as significant, the effects whose *t*-value was greater than 2, which corresponds to a significance level of 5% in two-tailed tests^72^. Partial effects were computed (for visual representation) from the (G)LMMs’ fixed effects using the *ggpredict* function from the *ggeffects* package (Version 0.8.0) in R (Version R-3.5.3).

### Statistical analysis of inter-word-spacing and character-size effects

Additional analyses were conducted to estimate MASC’s simulated behavior over FSC sentences as a function of inter-word spacing and print size (see Model simulations), using the same procedure as in the main analyses. However, since FSC readers were tested only in the normal spacing condition and for characters subtending 0.25°, data set was not entered as a predictor. MASC’s estimated effects are shown in Supplementary Tables 29-41 and described in Supplementary Methods 3. They were compared to previously reported effects of inter-word spacing and print size (Extended Data Fig. 9a-d) as well as data from spaced- and unspaced-language studies using different font sizes (Extended Data Fig. 8, 9e-g).

## Data availability

All stimulus materials and data used in the present study are available through the Zenodo repository: https://doi.org/10.5281/zenodo.5338616.

## Code availability

The model code and the R-scripts that were used for data analysis and figure generation are available through the Zenodo repository: https://doi.org/10.5281/zenodo.5338616.

## Acknowledgments

F.V.: The behavioral component of this work was carried out within the Labex BLRI (ANR-11-LABX-0036) and the Institut Convergence ILCB (ANR-16-CONV-0002), which benefited from support from the French government, managed by the French National Agency for Research (ANR) and the Excellence Initiative of Aix-Marseille University (A*MIDEX). http://www.agence-nationale-recherche.fr/. It also received support from the French government under the Programme “Investissements d’Avenir”, Initiative d’Excellence d’Aix-Marseille Université via A*Midex funding (AMX-19-IET-004), and ANR (ANR-17-EURE-0029). FV would like to thank Eric Castet for his statistical advice. G.J.Z.: The computational component of this work benefitted from support from the National Science Foundation (Awards 1718014, 1734260, and 1763981) and the National Institutes of Health (Award R21HD093912).

## Author contributions

F.V. and G.J.Z. conceptualized the project in 2012; F.V., H.A. and G.J.Z. designed the research; H.A. and G.J.Z. implemented the model and ran the simulations; F.V. analyzed the simulation and subject data and did the literature review; F.V., H.A. and G.J.Z. wrote the paper.

---

**Extended Data Table 1.**
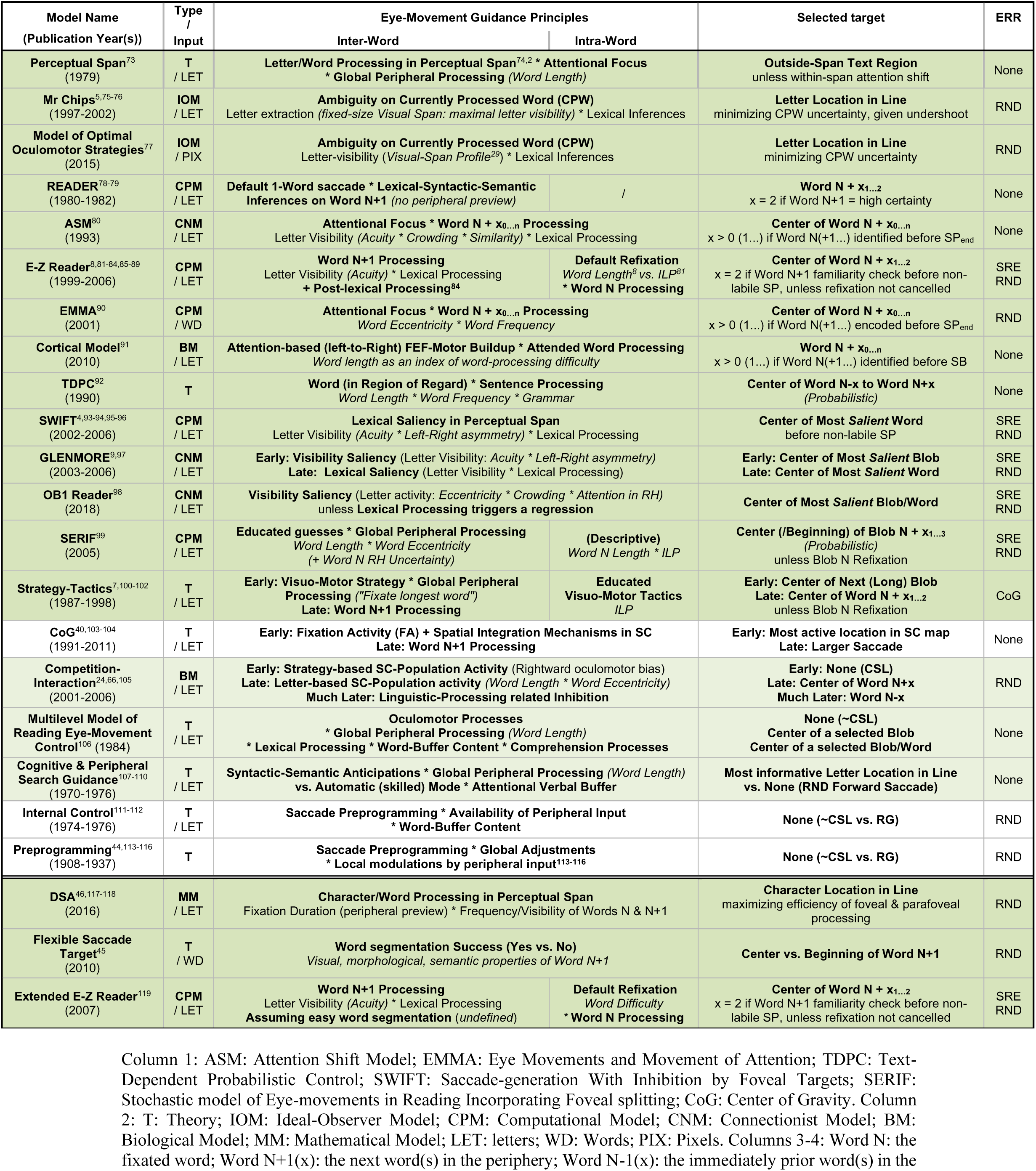
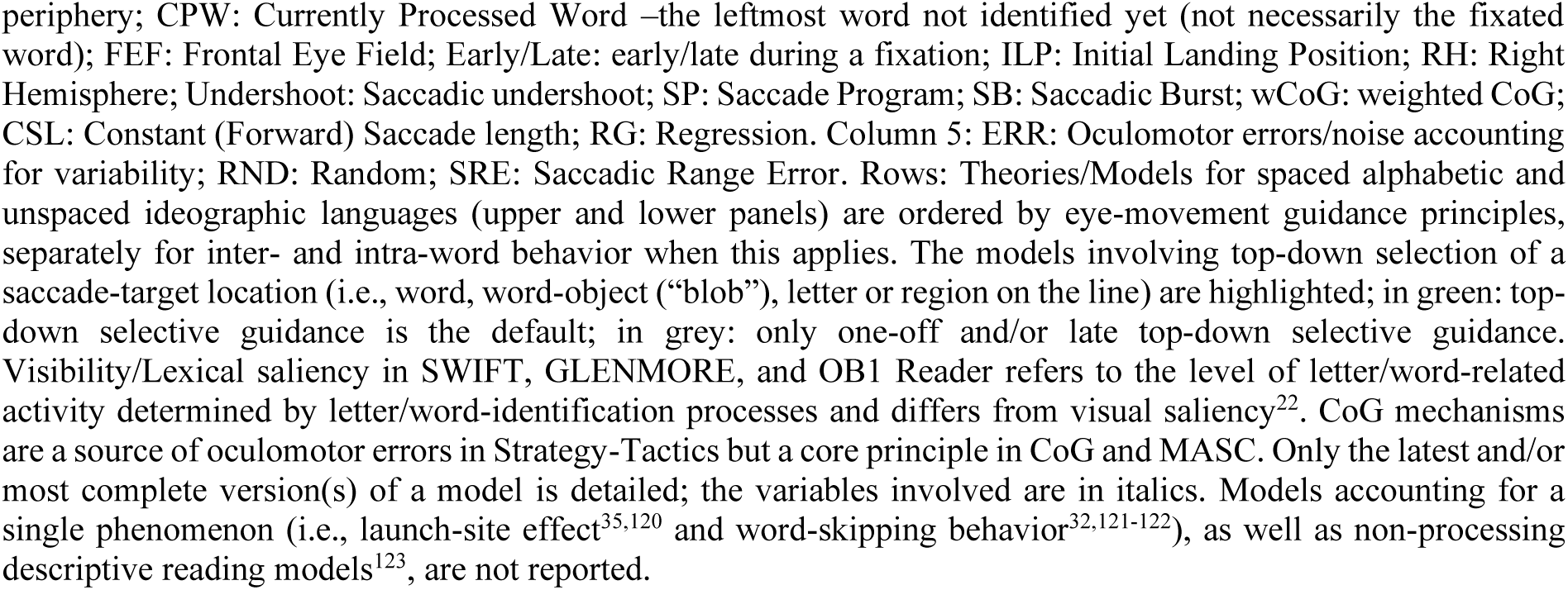
Models of eye-movement control during the reading of spaced and unspaced languages

| Model Name<br>(Publication Year(s)) | Type<br>/<br>Input | Eye-Movement Guidance Principles |  | Selected target | ERR |
| --- | --- | --- | --- | --- | --- |
|  |  | Inter-Word | Intra-Word |  |  |
| **Perceptual Span**<sup>73</sup><br>(1979) | **T**<br>/<br>LET | **Letter/Word Processing in Perceptual Span**<sup>74,2</sup> * **Attentional Focus**<br>* **Global Peripheral Processing** (*Word Length*) |  | **Outside-Span Text Region**<br>unless within-span attention shift | None |
| **Mr Chips**<sup>5,75-76</sup><br>(1997-2002) | **IOM**<br>/<br>LET | **Ambiguity on Currently Processed Word (CPW)**<br>Letter extraction (*fixed-size Visual Span: maximal letter visibility*) * Lexical Inferences |  | **Letter Location in Line**<br>minimizing CPW uncertainty, given undershoot | RND |
| **Model of Optimal Oculomotor Strategies**<sup>77</sup><br>(2015) | **IOM**<br>/<br>PIX | **Ambiguity on Currently Processed Word (CPW)**<br>Letter-visibility (*Visual-Span Profile*<sup>29</sup>) * Lexical Inferences |  | **Letter Location in Line**<br>minimizing CPW uncertainty | RND |
| **READER**<sup>78-79</sup><br>(1980-1982) | **CPM**<br>/<br>LET | **Default 1-Word saccade \* Lexical-Syntactic-Semantic Inferences on Word N+1** (*no peripheral preview*) | / | **Word N + x<sub>1...2</sub>**<br>x = 2 if Word N+1 = high certainty | None |
| **ASM**<sup>80</sup><br>(1993) | **CNM**<br>/<br>LET | **Attentional Focus \* Word N + x<sub>0...n</sub> Processing**<br>Letter Visibility (*Acuity \* Crowding \* Similarity*) * Lexical Processing |  | **Center of Word N + x<sub>0...n</sub>**<br>x > 0 (1...) if Word N(+1...) identified before SP<sub>end</sub> | None |
| **E-Z Reader**<sup>8,81-84,85-89</sup><br>(1999-2006) | **CPM**<br>/<br>LET | **Word N+1 Processing**<br>Letter Visibility (*Acuity*) * Lexical Processing<br>+ **Post-lexical Processing**<sup>84</sup> | **Default Refixation**<br>*Word Length*<sup>8</sup> vs. *ILP*<sup>81</sup><br>* **Word N Processing** | **Center of Word N + x<sub>1...2</sub>**<br>x = 2 if Word N+1 familiarity check before non-labile SP, unless refixation not cancelled | SRE<br>RND |
| **EMMA**<sup>90</sup><br>(2001) | **CPM**<br>/<br>WD | **Attentional Focus \* Word N + x<sub>0...n</sub> Processing**<br>*Word Eccentricity \* Word Frequency* |  | **Center of Word N + x<sub>0...n</sub>**<br>x > 0 (1...) if Word N(+1...) encoded before SP<sub>end</sub> | RND |
| **Cortical Model**<sup>91</sup><br>(2010) | **BM**<br>/<br>LET | **Attention-based (left-to-Right) FEF-Motor Buildup \* Attended Word Processing**<br>*Word length as an index of word-processing difficulty* |  | **Word N + x<sub>0...n</sub>**<br>x > 0 (1...) if Word N(+1...) identified before SB | None |
| **TDPC**<sup>92</sup><br>(1990) | **T** | **Word (in Region of Regard) \* Sentence Processing**<br>*Word Length \* Word Frequency \* Grammar* |  | **Center of Word N-x to Word N+x**<br>(*Probabilistic*) | None |
| **SWIFT**<sup>4,93-94,95-96</sup><br>(2002-2006) | **CPM**<br>/<br>LET | **Lexical Saliency in Perceptual Span**<br>Letter Visibility (*Acuity \* Left-Right asymmetry*) * Lexical Processing |  | **Center of Most Salient Word**<br>before non-labile SP | SRE<br>RND |
| **GLENMORE**<sup>9,97</sup><br>(2003-2006) | **CNM**<br>/<br>LET | **Early: Visibility Saliency** (Letter Visibility: *Acuity \* Left-Right asymmetry*)<br>**Late: Lexical Saliency** (Letter Visibility * Lexical Processing) |  | **Early: Center of Most Salient Blob**<br>**Late: Center of Most Salient Word** | SRE<br>RND |
| **OB1 Reader**<sup>98</sup><br>(2018) | **CNM**<br>/<br>LET | **Visibility Saliency** (Letter activity: *Eccentricity \* Crowding \* Attention in RH*)<br>unless **Lexical Processing** triggers a regression |  | **Center of Most Salient Blob/Word** | SRE<br>RND |
| **SERIF**<sup>99</sup><br>(2005) | **CPM**<br>/<br>LET | **Educated guesses \* Global Peripheral Processing**<br>*Word Length \* Word Eccentricity*<br>(+ *Word N RH Uncertainty*) | **(Descriptive)**<br>*Word N Length \* ILP* | **Center (/Beginning) of Blob N + x<sub>1...3</sub>**<br>(*Probabilistic*)<br>unless Blob N Refixation | SRE<br>RND |
| **Strategy-Tactics**<sup>7,100-102</sup><br>(1987-1998) | **T**<br>/<br>LET | **Early: Visuo-Motor Strategy \* Global Peripheral Processing** (*"Fixate longest word"*)<br>**Late: Word N+1 Processing** | **Educated Visuo-Motor Tactics**<br>*ILP* | **Early: Center of Next (Long) Blob**<br>**Late: Center of Word N + x<sub>1...2</sub>**<br>unless Blob N Refixation | CoG |
| **CoG**<sup>40,103-104</sup><br>(1991-2011) | **T**<br>/<br>LET | **Early: Fixation Activity (FA) + Spatial Integration Mechanisms in SC**<br>**Late: Word N+1 Processing** |  | **Early: Most active location in SC map**<br>**Late: Larger Saccade** | None |
| **Competition-Interaction**<sup>24,66,105</sup><br>(2001-2006) | **BM**<br>/<br>LET | **Early: Strategy-based SC-Population Activity** (Rightward oculomotor bias)<br>**Late: Letter-based SC-Population activity** (*Word Length \* Word Eccentricity*)<br>**Much Later: Linguistic-Processing related Inhibition** |  | **Early: None (CSL)**<br>**Late: Center of Word N+x**<br>**Much Later: Word N-x** | RND |
| **Multilevel Model of Reading Eye-Movement Control**<sup>106</sup> (1984) | **T**<br>/<br>LET | **Oculomotor Processes**<br>* **Global Peripheral Processing** (*Word Length*)<br>* **Lexical Processing \* Word-Buffer Content \* Comprehension Processes** |  | **None (~CSL)**<br>**Center of a selected Blob**<br>**Center of a selected Blob/Word** | None |
| **Cognitive & Peripheral Search Guidance**<sup>107-110</sup><br>(1970-1976) | **T**<br>/<br>LET | **Syntactic-Semantic Anticipations \* Global Peripheral Processing** (*Word Length*)<br>vs. **Automatic (skilled) Mode \* Attentional Verbal Buffer** |  | **Most informative Letter Location in Line**<br>vs. **None (RND Forward Saccade)** | None |
| **Internal Control**<sup>111-112</sup><br>(1974-1976) | **T**<br>/<br>LET | **Saccade Preprogramming \* Availability of Peripheral Input**<br>* **Word-Buffer Content** |  | **None (~CSL vs. RG)** | RND |
| **Preprogramming**<sup>44,113-116</sup><br>(1908-1937) | **T** | **Saccade Preprogramming \* Global Adjustments**<br>* **Local modulations by peripheral input**<sup>113-116</sup> |  | **None (~CSL vs. RG)** | RND |
| **DSA**<sup>46,117-118</sup><br>(2016) | **MM**<br>/<br>LET | **Character/Word Processing in Perceptual Span**<br>Fixation Duration (peripheral preview) * Frequency/Visibility of Words N & N+1 |  | **Character Location in Line**<br>maximizing efficiency of foveal & parafoveal processing | RND |
| **Flexible Saccade Target**<sup>45</sup><br>(2010) | **T**<br>/<br>WD | **Word segmentation Success (Yes vs. No)**<br>*Visual, morphological, semantic properties of Word N+1* |  | **Center vs. Beginning of Word N+1** | RND |
| **Extended E-Z Reader**<sup>119</sup><br>(2007) | **CPM**<br>/<br>LET | **Word N+1 Processing**<br>Letter Visibility (*Acuity*) * Lexical Processing<br>**Assuming easy word segmentation** (*undefined*) | **Default Refixation**<br>*Word Difficulty*<br>* **Word N Processing** | **Center of Word N + x<sub>1...2</sub>**<br>x = 2 if Word N+1 familiarity check before non-labile SP, unless refixation not cancelled | SRE<br>RND |

Column 1: ASM: Attention Shift Model; EMMA: Eye Movements and Movement of Attention; TDPC: Text- Dependent Probabilistic Control; SWIFT: Saccade-generation With Inhibition by Foveal Targets; SERIF: Stochastic model of Eye-movements in Reading Incorporating Foveal splitting; CoG: Center of Gravity. Column 2: T: Theory; IOM: Ideal-Observer Model; CPM: Computational Model; CNM: Connectionist Model; BM: Biological Model; MM: Mathematical Model; LET: letters; WD: Words; PIX: Pixels. Columns 3-4: Word N: the fixated word; Word N+1(x): the next word(s) in the periphery; Word N-1(x): the immediately prior word(s) in the

**Extended Data Figure 1.**
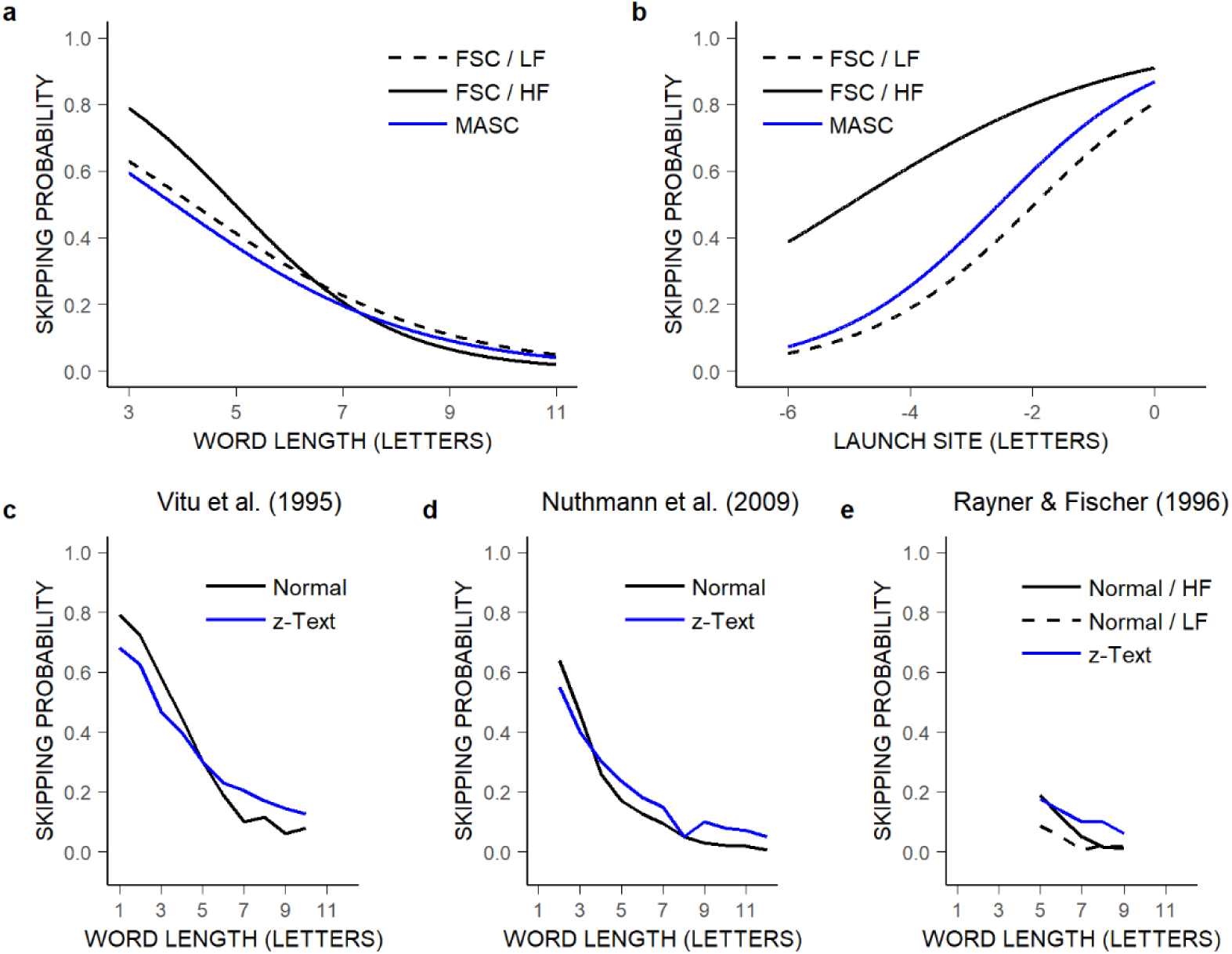
Language-related processes only mildly modulate word-skipping behavior. **a-b**, Probability of word skipping as a function of word length (**a**), and for 4-letter words as a function of saccades’ launch-site distance to the space in front of the words (in letters; **b**), for MASC (blue) across words of different frequencies and for FSC readers (black) separately for low- and high-frequency (LF and HF) words in French^124^ (dashed and solid lines). Curves for MASC and FSC readers represent partial effects computed from GLMMs reported respectively in Supplementary Table 5 (Fig. 2b) and in Albrengues et al.’s study^31^; in the latter, log word frequency was entered as an additional continuous predictor, meaning that predictions could be derived for *n* levels of log word frequency; here, LF and HF corresponded to the minimal and maximal mean log frequency across word lengths (0.01 and 9.59 log units). MASC nearly behaved as FSC readers encountering LF words. **c-e**, Previously reported relationship between word-skipping rate and word length during normal reading (black; by word frequency in the right panel) and the “reading” of meaningless z-transformed text materials (blue; all letters replaced by the letter *z*)^33,94,126,(127)^; the difference between normal- and z-reading conditions is consistent with the difference observed between FSC readers and MASC (Fig. 2b; **a**) and between HF and LF words in FSC readers (**a**), thus confirming that lexical/linguistic processes only mildly modulate default, mindless, word-skipping behavior^31–32,125^.

**Extended Data Figure 2.**
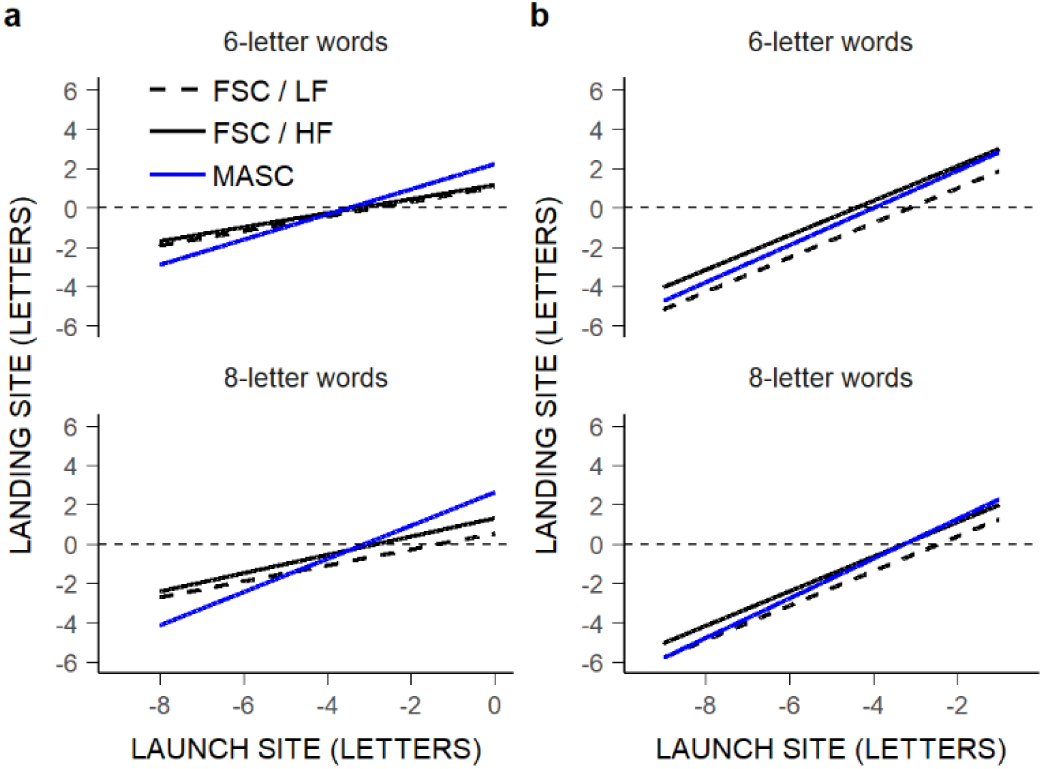
Language-related processes only mildly modulate saccades’ landing positions. **a-b**, Landing positions in 6- and 8-letter words as a function of saccades’ launch-site distance to the space in front of the words, for MASC (blue) across words of different frequencies and for FSC readers (black) separately for low- and high-frequency (LF and HF) words^124^ (dashed and solid lines); **a**: within-word landing positions; **b**: all saccades’ landing positions. The curves for MASC and FSC readers represent the partial effects computed respectively from two separate LMMs fitted to raw landing positions. In the first, the fixed structure included data set (2 levels), word length, launch-site distance, and their interaction as predictors (yielding similar estimates as LMMs fitted to the GMM-estimated mean of landing positions; Fig. 2f-g, Supplementary Tables 10,12), and in the second (fitted only to FSC data), the fixed structure included word length, launch-site distance, word frequency, and all interactions as predictors as in Albrengues et al.’s study^31^; LF and HF: the lowest and highest word frequency across words of 6 and 8 letters (-1.97 vs. 6.30 log units respectively). Differences in within-word landing positions between MASC and FSC readers were greater than differences between HF and LF words in FSC readers^46,128–134^ (**a**), and hence could not entirely be due to MASC lacking a lexicon; rather these differences resulted from comparing within-word truncated (though more standard) distributions that were not equally spread (see Supplementary Methods 1). Indeed, when all saccades’ landing positions were analyzed, differences between MASC and FSC readers were smaller, and as tiny as differences between HF and LF words in FSC readers (**b**). This confirms that lexical processing modulates, but only very mildly, the extent of default forward saccades, regardless of word boundaries^31,46^.

**Extended Data Figure 3.**
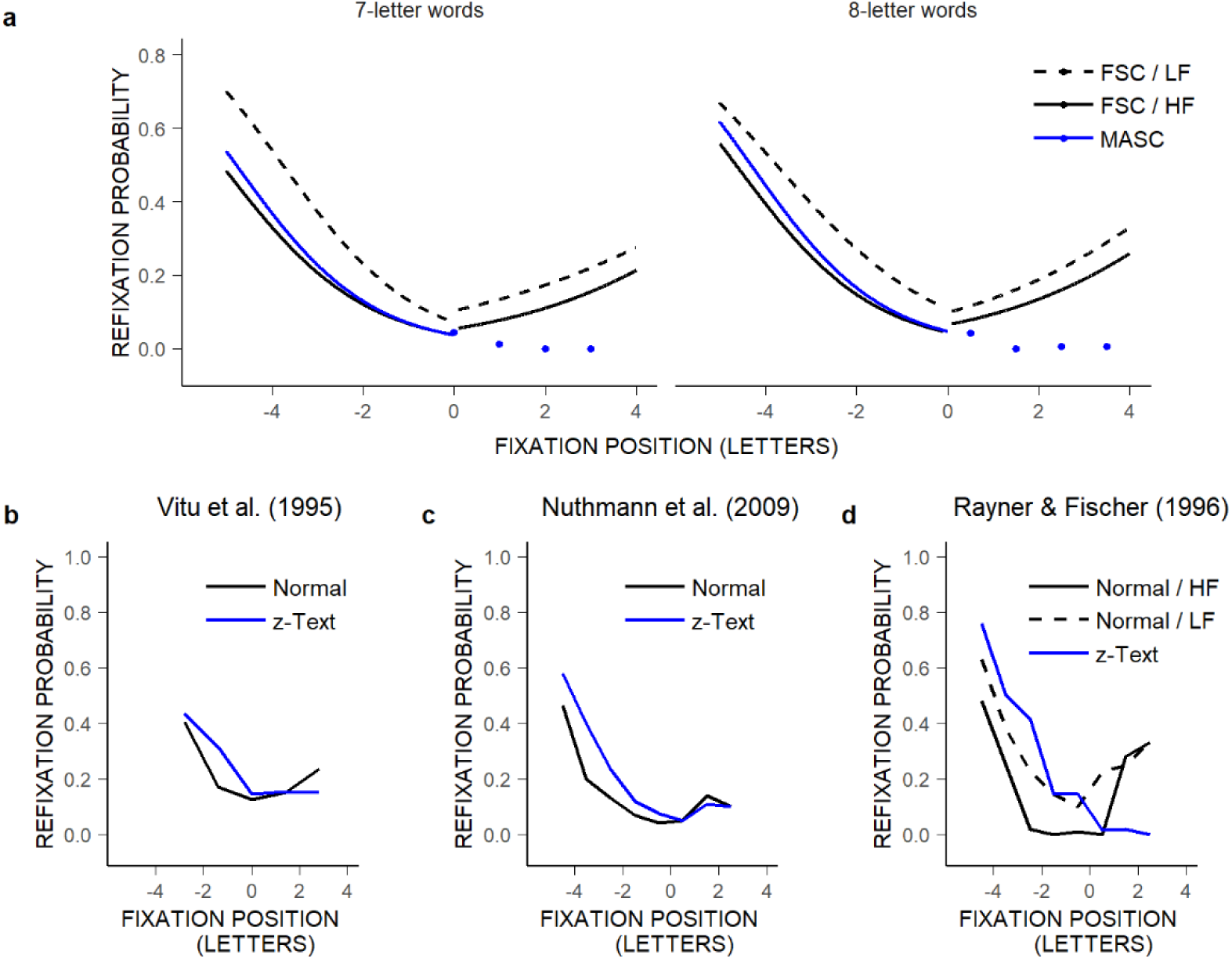
Language-related processes only partly contribute to the Refixation-OVP effect. **a-b**, Probability of within-word refixations as a function of initial fixation location in 7- and 8-letter words, for MASC (blue) across words of different frequencies and for FSC readers (black) separately for low- and high-frequency (LF and HF) words^124^ (dashed and solid lines). Dots represent means. Curves represent partial effects computed from GLMMs; the first one, reported in Supplementary Table 13, fitted MASC and FSC data associated with initial fixations in the first halves of words (Fig. 2h) and enabled representation of MASC’s left-OVP effect; the other two models fitted FSC data separately for initial fixations in the first and second halves of words using initial fixation position, word length, log word frequency, and all interactions as predictors (the random structure included a random intercept by subject, sentence pair, and word); LF and HF: -1.97 and 6.30 log units respectively. MASC, which reproduced only the left wing of U-shaped OVP curves, initiated as few refixations from the words’ centers as readers viewing HF words, but nearly as many refixations from the very-beginnings of words as readers viewing LF words, thus behaving like readers benefiting not from lexical facilitation (that suppresses unnecessary refixations from the words’ beginnings) and encountering no word-processing difficulties (which cause additional refixations from the center and likely also the end of words). **b-d**, Previously reported Refixation-OVP effect in 7-letter words during the reading of normal text (black; by word frequency -right panel) and z-transformed text (blue)^33,94,126^, revealing similarities in eye-movement behavior between z-readers and MASC (**a**), notably a Refixation-OVP effect with no right wing^137^, and a slightly greater left-OVP effect compared to normal reading. Thus, language-related processes contribute to, but do not fully explain, the Refixation-OVP effect (see Supplementary Methods 1).

**Extended Data Figure 4.**
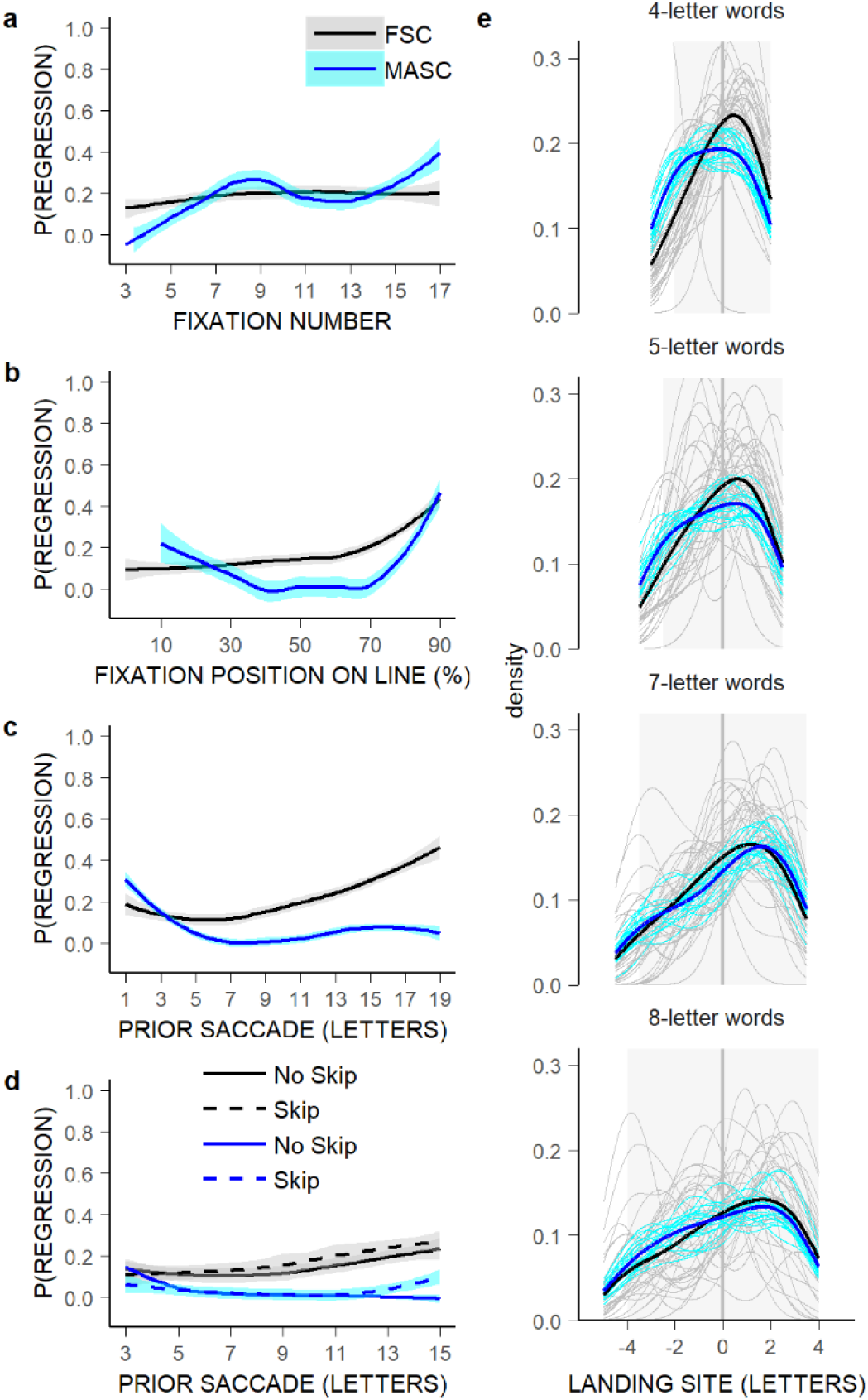
Language-related processes more greatly influence the likelihood of regressions, but not their metric. **a-d**, Mean probability of a regressive saccade (following a progressive saccade) in MASC (blue) vs. FSC readers (black) as a function of fixation number in the sentence (**a**) and fixation position on the line (in percentage of line length; **b**), and as a function of the length of the prior saccade (in letters), irrespective of how many words the saccade traversed (**c**) and separately for word-skipping and non-word-skipping saccades (dashed and solid lines; **d**). All curves were fitted with a loess smoothing function (with 0.95 confidence bands in cyan and grey). Unlike FSC readers, MASC generated regressive saccades essentially from the end parts of the sentences, regardless of prior saccade length, failing to replicate the well-established increase in regression rate with increasing prior saccade length^138–140^, thereby suggesting that regressions mostly result from language-related processes^141^. **e**, Probability density functions of the landing positions of regressive saccades, in letters relative to the centers of 4-,5-,7-, and 8-letter words, across and by subjects (thick and thin lines) for MASC and FSC readers. Both FSC readers and MASC most frequently landed to the right of the words’ centers^140^ regardless of their length^142^ (Supplementary Tables 14-15), thus suggesting that visuo-motor principles in the SC determine the landing positions of regressive saccades. Only the occurrence of regressions would primarily be under top-down control (see Supplementary Methods 1).

**Extended Data Figure 5.**
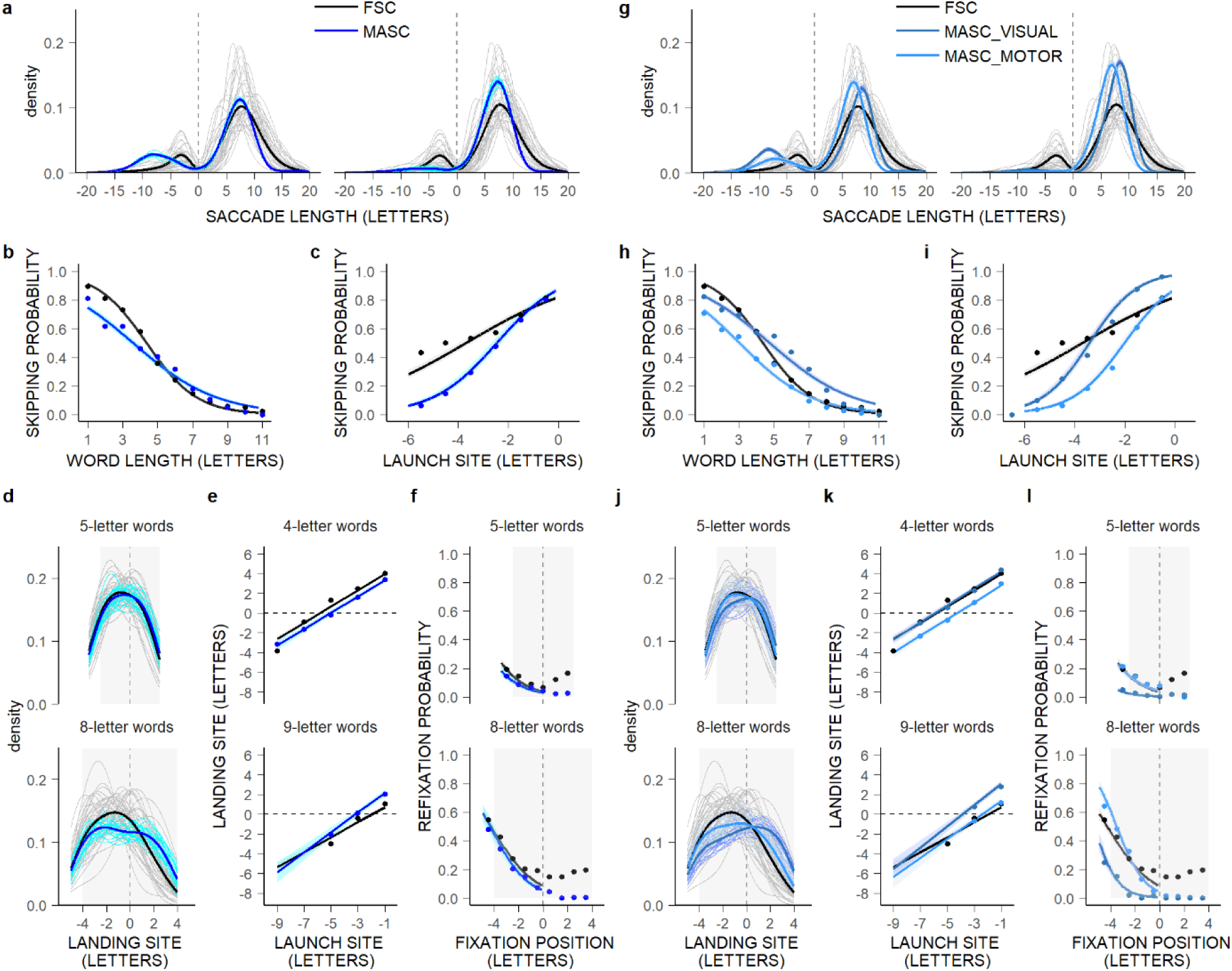
Cascaded averaging over both visual- and motor-point images in SC space accounts best for readers’ oculomotor behavior. **a-l**, First-pass oculomotor behavior for MASC (blue; **a-f**), and for MASC with averaging over visual- or motor-point images only (MASC_VISUAL and MASC_MOTOR; dark/light blue; **g-l**), compared to FSC readers (black) –see Methods, Supplementary Methods 2. **a,g**, Probability density functions of saccade lengths (in letters) across and by subjects (thick and thin lines); left panels: for comparison for data sets matched for numbers of fixations. **b-c,h-i**, Mean probability of word skipping (dots) as a function of word length (in letters; **b,h**), and for 4-letter words as a function of saccades’ launch-site distance to the space in front of the words (in letters; **c,i**), and partial effects (lines), with 0.95 confidence intervals (bands), computed from GLMMs (Supplementary Tables 20-21). **d,j**, Probability density functions of within-word landing positions (in letters relative to the centers of words; vertical grey lines) across and by subjects for 5- and 8-letter words (grey-filled rectangle areas). **e,k**, GMM-estimated means of all landing positions, in letters relative to the centers of 4- and 9-letter words, as a function of launch-site distance, and partial effects, with 0.95 confidence intervals, computed from LMMs (Supplementary Tables 26-27). **f,l**, Mean within-word refixation probability as a function of initial fixation location (in letters relative to the centers of words) for 5- and 8-letter words, and partial effects, with 0.95 confidence intervals, computed from GLMMs but only for the left wing of OVP curves (Supplementary Table 28). MASC’s first-pass behavior (**a-f**) resembled that observed when MASC and FSC were matched for numbers of fixations (Fig 2), although regressions were less likely. Averaging over visual- or motor-point images sufficed to generate word-based oculomotor behavior (**g-l**), but it did not beat averaging over both visual and motor-point images (**a-f**).

**Extended Data Figure 6.**
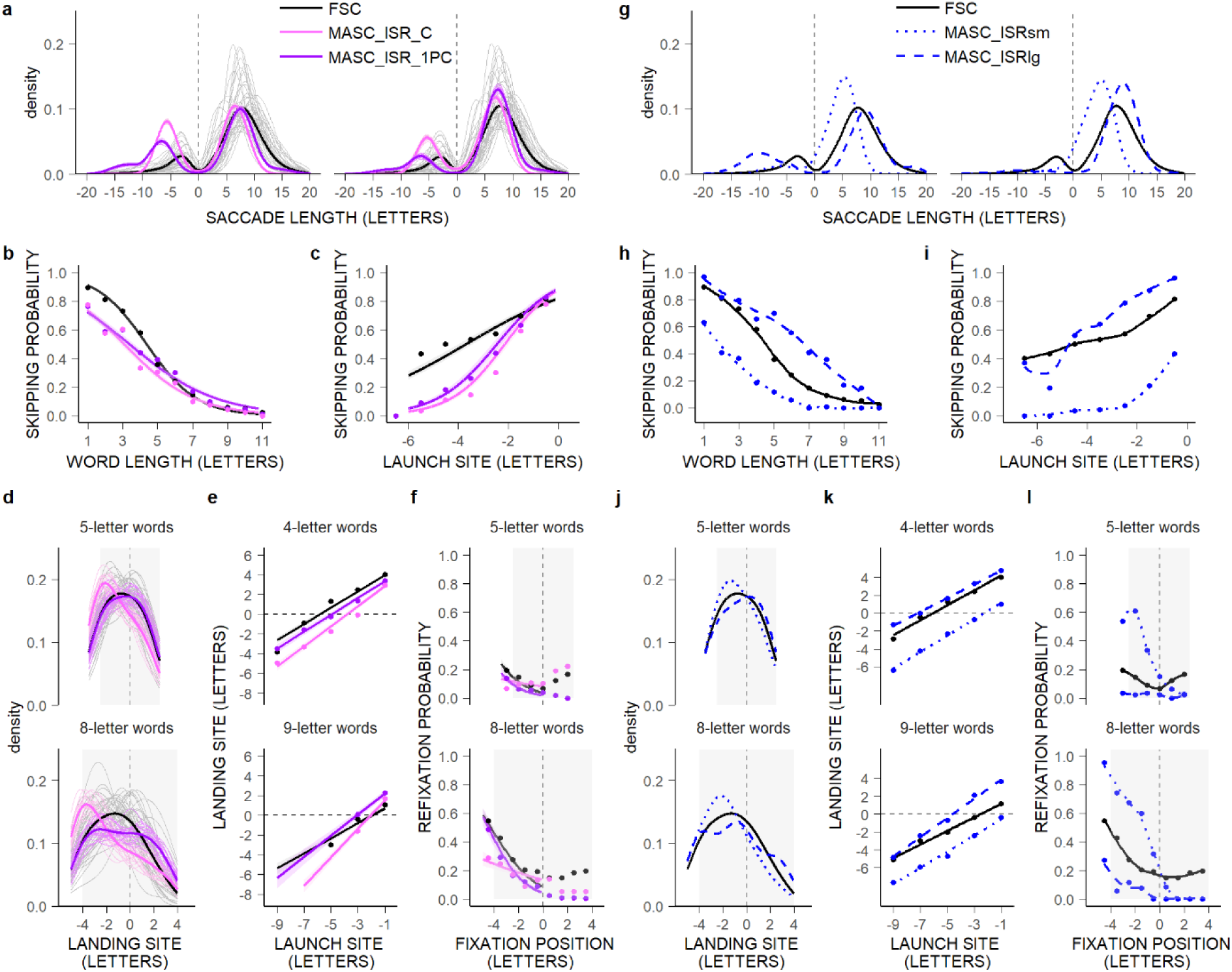
There is no need for specific ISR-parameter settings to reproduce readers’ stereotyped oculomotor behavior. **a-l**, First-pass oculomotor behavior for MASC with ISR applied to the current fixation (ISR_C; pink) or the current and immediately prior fixations (ISR_1PC; purple; **a-f**; see Supplementary Methods 2), and for MASC with the smallest and largest tested ISR-window sizes (dotted/dashed-blue lines; **g-l**) during that parameter fit (see Methods), compared to FSC readers (black). **a,g**, Probability density functions of saccade lengths across (**a,g**) and by subjects (**a**); left panels: for comparison for data sets matched for numbers of fixations. **b-c,h-i**, Mean probability of word skipping (dots) as a function of word length (in letters; **b,h**), and for 4-letter words as a function of saccades’ launch-site distance to the space in front of the words (in letters; **c,i**); in **b-c**, partial effects, with 0.95 confidence intervals, computed from GLMMs (Supplementary Tables 20-21); in h-i, Loess-smoothing curves. d,j, Probability density functions of within-word landing positions (in letters relative to the centers of words) across (**d,j**) and by subjects (**d**) for 5- and 8-letter words. **e,k**, Mean of all landing positions, in letters relative to the centers of 4- and 9-letter words, as a function of launch-site distance; in **e**, GMM-estimated means and partial effects, with 0.95 confidence intervals, computed from LMMs (Supplementary Tables 26-27); in **k**, raw means and Loess-smoothing curves. **f,l**, Mean within-word refixation probability as a function of initial fixation location, in letters relative to the centers of 5- and 8-letter words; in **f**, partial effects, with 0.95 confidence intervals, computed from GLMMs but only for the left-OVP wing (Supplementary Table 28); in l, Loess-smoothing curves. Word-based phenomena held across both MASC_ISR_C and MASC_ISR_1PC (**a-f**), and the whole range of ISR-window sizes (**g-l**), although with a slightly poorer fit than for MASC (Extended Data Fig. 5a-f).

**Extended Data Figure 7.**
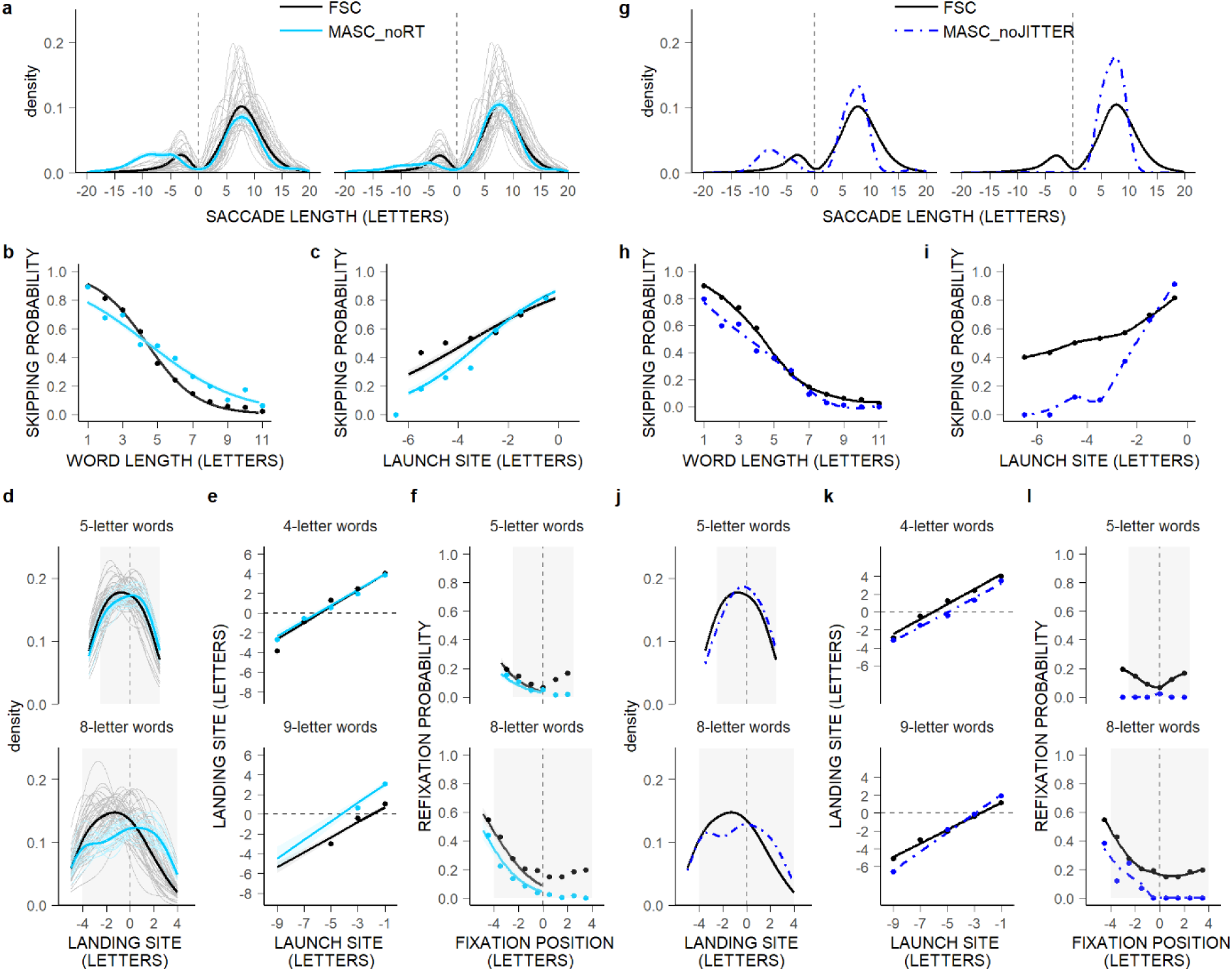
RT and population jitter contribute only mildly to readers’ stereotyped oculomotor behavior. **a-l**, First-pass oculomotor behavior for MASC with no RT (MASC_noRT; turquoise; **a-f**; see Supplementary Methods 2), and for MASC amputated from jitter over the winning population (MASC_noJITTER; dashed-blue lines; g-l) compared to FSC readers. See Extended Data Fig. 6 legend. Overall, MASC_noRT (**a-f**) and MASC_noJITTER (**g-l**) made very similar predictions to MASC (Extended Data Fig. 5a-f).

**Extended Data Figure 8.**
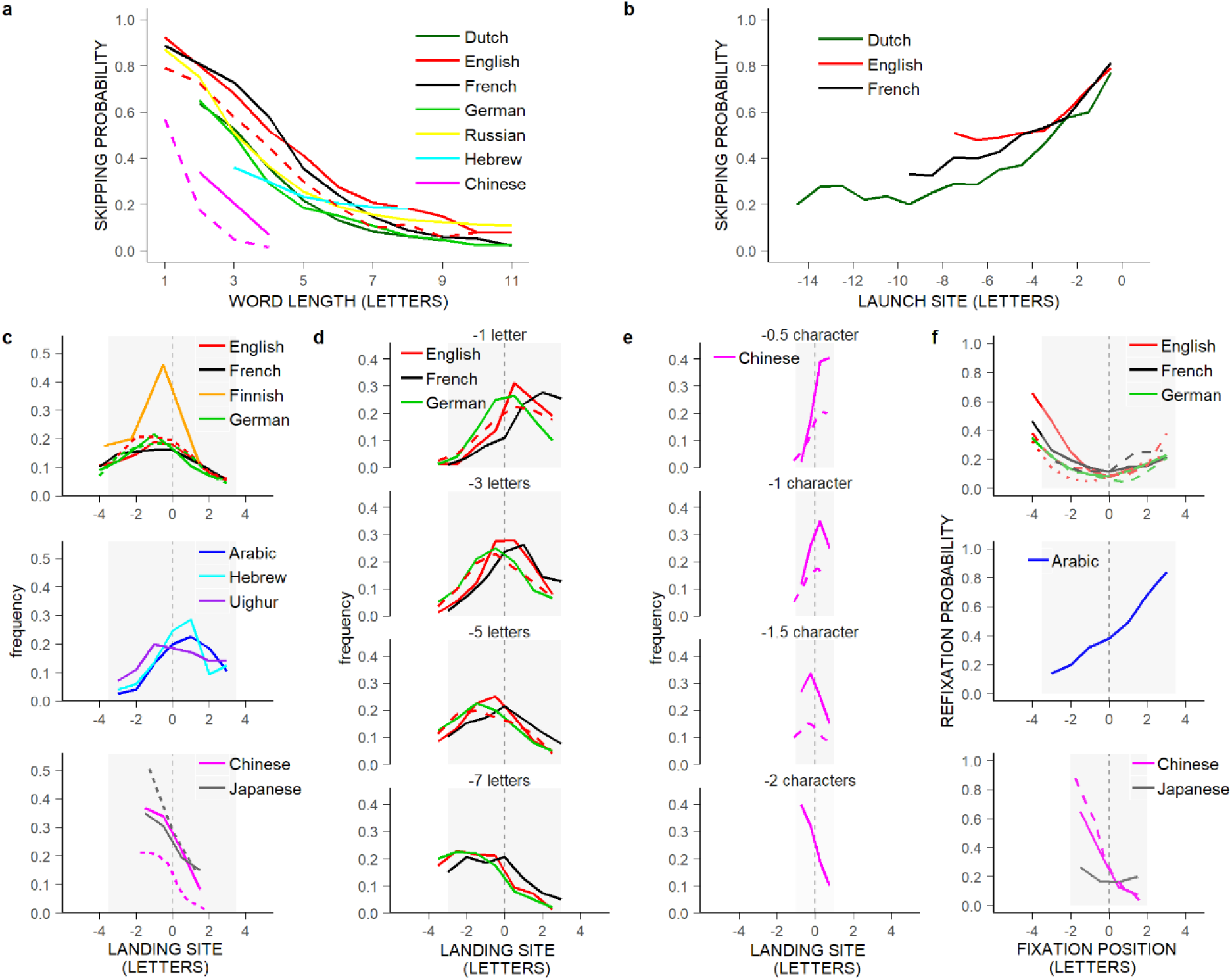
Word-based eye-movement phenomena across studies and languages. **a**, Relationship between the probability of word skipping and word length (in letters/characters) in different studies (line types) and languages (colors), including spaced-alphabetic languages read from left to right (Dutch^32^, English^33,42^, French/FSC^31^, German^4^, Russian^143^) and from right to left (Hebrew^144^), as well as left-to-right unspaced-ideographic languages (Chinese^45,118^). **b**, Relationship between the probability of word skipping and saccades’ launch-site distance to the space in front of the words for 4-letter words (in letters/characters) in alphabetic languages (Dutch^32^, English^121^, French/FSC^31^). **c**, Frequency distributions of within-word landing positions (in letters/characters relative to the centers of words) representing the PVL effect separately for 7-letter words in spaced-alphabetic languages read from left to right (English^35,43^, French/FSC^31^, Finnish^145^, German^146–147^) and from right to left (Arabic^148^, Hebrew^144^, Uighur^132^) and 4-character words in left-to-right unspaced-ideographic languages (Chinese^45,118^ and Japanese^149–150^); data for other word lengths showed similar pattern (but see^128^ for Chinese). **d-e**, Frequency distributions of within-word landing positions (in letters/characters relative to the centers of words) for different launch-site distances (-1,-3,-5,-7 letters) separately for 6-letter words in English^35,151^, French/FSC^31^, and German^147^ (**d**) and 2-character words in Chinese^45,128^ (**e**). **f**, Within-word refixation probability as a function of initial fixation location (in letters/characters relative to the centers of words) separately for 7-letter words in left-to right and right-to-left spaced-alphabetic languages (English^135,151–152^, French/FSC^31,153^, German^146–147^, and Arabic^148^) and 4-character words in left-to-right unspaced-ideographic languages (Chinese^45,118^, Japanese^149^). Color code: Arabic: blue; Chinese: pink; Dutch: dark green; English: red; Finnish: orange; French: black; German: green; Japanese: grey; Uighur: purple; Russian: yellow. All word-based eye-movement patterns are very similar across studies and languages; differences in word-skipping behavior, PVL, and OVP effects, notably between spaced and unspaced languages, are attributable to print-size differences (Extended Data Fig. 9, Supplementary Methods 3).

**Extended Data Figure 9.**
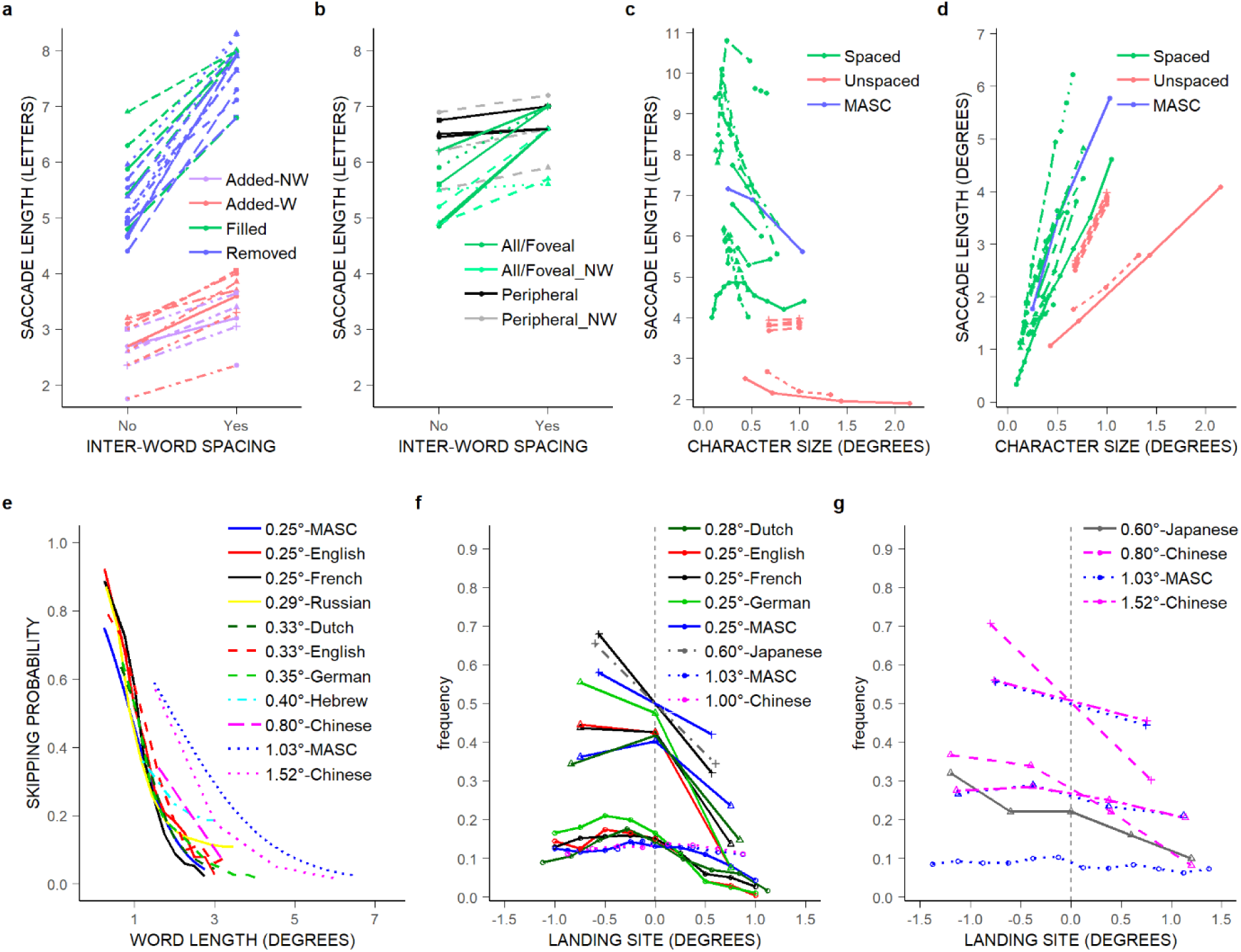
Character-print size, but not inter-word spacing, accounts for differences between spaced and unspaced languages. **a-b**, Mean forward-saccade length (in letters/characters) as a function of inter-word spacing in different studies (line types), separately for global spacing manipulations (**a:** space removal^154–160^ (blue) or space filling^155,158, 160–161^ (green) in normally spaced English/French/Spanish texts/sentences and space addition between words^150,162–163^ (“-W”; pink) or non-words^162–163^ (“-NW”; purple) in normally unspaced Chinese/Japanese texts/sentences) and gaze-contingent space-filling manipulations (**b**: in the fovea and possibly also in the (right) periphery^24,164,165^ (green), or exclusively in the (right) periphery, thus preserving the space(s) around the fixated word^164–166^ (black/grey); letters in the filled-text region were preserved or replaced by x’s/random letters (“-NW”) –for even tinier spacing effects on within-word landing-positions see^150, 156–161,163, 167– 168,(133, 169),but 150,154,167,(170)^. **c-d**, Mean forward-saccade length in letters (**c**) and in degrees (**d**) as a function of angular print size for spaced-alphabetic languages (English^171–174^, French^26,175–177^, German^178^, Slovene^179^; green) and Chinese (pink)^25,178,180^, and for MASC (blue) viewing FSC sentences in three print sizes. Paterson and Tinker^174^: we assumed a 50-cm viewing distance; Kolers et al.^172^: we divided the reported number of fixations by line length. **e**, Probability of word skipping in different studies/languages (Extended Data Fig 8a; same color code) using various print sizes (line types), and for MASC in two print-size conditions (Fig. 4h, Supplementary Table 37), re-plotted as a function of word length in degrees^181–182^. **f-g**, Frequency distributions of within-word landing positions (in degrees relative to the words’ centers) in different studies/languages (**f:** Dutch^183^, English^43^, French/FSC^31^, German^147^, Chinese^128^, Japanese^149^; **g:** Chinese^45,118^, Japanese^149^; same color code as in Extended Data Fig 8c) using different print sizes (**f**: 0.25°, 0.28°, 0.6°, 1°; **g**: 0.6°, 0.8°, 1.03°, 1.52°; solid to dotted lines), and for MASC in two print-size conditions (**f**: 1°, 0.25°; **g**: 1°), separately for two angular word sizes (**f**: 2-2.52°; **g**: 3-3.2°) and three bin sizes (**f**: 0.25°, 0.75-0.84°, 1.125-1.2°; **g**: 0.25°, 0.6-0.8°, 1.5°; circle, triangle and cross) whenever possible given reported data. All findings are described in Supplementary Methods 3.

## References

1. Yarbus, A. L. Eye Movements and Vision. (Plenum Press, New York, 1967).

2. Rayner, K. Eye movements in reading and information processing: 20 years of research. Psychol. Bull. 124(**3**), 372–422 (1998).

3. Einhauser, W., Spain, M. & Perona, P. Objects predict fixations better than early saliency. J. Vis. 8(14):18, 10.1167/8.14.18. (2008).

4. Engbert, R., Nuthmann, A., Richter, E. & Kliegl, R. Swift: A dynamical model of saccade generation during reading. Psychol. Rev. 112(**4**), 777–813 (2005).

5. Legge, G.E., Hooven, T.A., Klitz, T.S., Mansfield, J.S. & Tjan, B.S. Mr Chips 2002: new insights from an ideal-observer model of reading. Vision Res. 42, 2219–2234 (2002).

6. Najemnik, J. & Geisler, W.S. Optimal eye movement strategies in visual search. Nature 434, 387–391 (2005).

7. O’Regan, J.K. in Eye movements and their role in visual and cognitive processes (ed. Kowler, E.) 395–453 (Elsevier, Amsterdam, 1990).

8. Reichle, E.D., Rayner, K. & Pollatsek, A. The E-Z reader model of eye movement control in reading: Comparisons to other models. Behav. Brain Sci. 26, 445–526 (2003).

9. Reilly, R. & Radach, R. Some empirical tests of an interactive activation model of eye movement control in reading. Cogn. Syst. Res. 7, 34–55 (2006).

10. Zelinsky, G.J. A theory of eye movements during target acquisition. Psychol. Rev. 115, 787–835 (2008).

11. Müri, R.M. & Nyffeler, T. Neurophysiology and neuroanatomy of reflexive and volitional saccades as revealed by lesion studies with neurological patients and transcranial magnetic stimulation (TMS). Brain Cognition 68, 284–292 (2008).

12. Schall, J.D. & Cohen, J.Y. in Handbook on eye movements (eds. Liversedge, S.P., Gilchrist, I.D., & Everling, S.) 357–379 (Oxford University Press, 2011).

13. Sparks, D.L. Translation of sensory signals into commands for control of saccadic eye movements: Role of primate superior colliculus. Physiol. Rev. 66(**1**), 118–171 (1986).

14. Hall, J.A., Foster, R.E., Ebner, F.F. & Hall, W.C. Visual cortex in a reptile, the turtle (Pseydem ys scripta and Chrysem ys picta). Brain Res. 130, 197–216 (1977).

15. Stein, B.E. & Gaither, N.S. Sensory representation in reptilian optic tectum: Some comparisons with mammals. J. Comp. Neurol. 202, 69–87 (1981).

16. Adeli, H., Vitu, F. & Zelinsky, G.J. A model of the superior colliculus predicts fixation locations during scene viewing and visual search. J. Neurosci. 37(**6**), 1453–1467 (2017).

17. Schiller, P.H., Sandell, J.H., & Maunsell, J.H.R. The effect of frontal eye field and superior colliculus lesions on saccade latencies in the Rhesus monkey. J. Neurophysiol. 57(**4**), 1033–1049 (1987).

18. Braddick, O. et al. Possible blindsight in babies lacking one cerebral hemisphere. Nature 360, 461–463 (1992).

19. Wallace, M.T., McHaffie, J.G. & Stein, B.E. Visual response properties and visuotopic representation in the newborn monkey superior colliculus. J. Neurophysiol. 78(**5**), 2732–2741 (1997).

20. Land, M.F. in Handbook on eye movements (eds. Liversedge, S.P., Gilchrist, I.D., & Everling, S.) 3–15 (Oxford University Press, 2011).

21. Ottes F.P., van Gisbergen J.A. & Eggermont J.J. Visuomotor fields of the superior colliculus: a quantitative model. Vision Res. 26, 857–873 (1986).

22. Itti, I. & Koch, C. Computational modeling of visual attention. Nat. Rev. Neurosci 2, 194–203 (2001).

23. Epelboim, J., Booth, J.R. & Steinman, R.M. Much ado about nothing: the place of space in text. Vision Res. 36(**3**), 465–470 (1996).

24. Yang, S.-N. & McConkie, G.W. Saccade generation during reading: Are words necessary? Eur. J. Cogn. Psychol. 16(1/2), 226–261 (2004).

25. Shu, H., Zhou, W., Yan, M. & Kliegl, R. Font size modulates saccade-target selection in Chinese reading. Atten. Percept. Psychophys. 73, 482–490 (2011).

26. Yao-N’Dré, M., Castet, E. & Vitu, F. Inter-word eye behavior during reading is not invariant to character size: Evidence against systematic saccadic range error in reading. Vis. Cogn. 22(3-4), 415–440 (2014).

27. Kaas, J.H. The evolution of brains from early mammals to humans. WIREs Cogn Sci. 4, 33–45 (2013).

28. Hanes, DP & Wurtz, RH. Interaction of the frontal eye field and superior colliculus for saccade generation. J. Neurophysiol. 85(2), 804–815 (2000).

29. Legge, G.E., Mansfield, J.S. & Chung, S.T.L. Psychophysics of reading XX. Linking letter recognition to reading speed in central and peripheral vision. Vision Res. 41, 725–743 (2001).

30. Chanceaux, M., Vitu, F., Bendahman, L., Thorpe, S. & Grainger, J. Word Processing Speed in Peripheral Vision Measured with a Saccadic Choice Task. Vision Res. 56, 10–19 (2012).

31. Albrengues, C., Lavigne, F., Aguilar, C., Castet, E. & Vitu, F. Linguistic processes do not beat visuo-motor constraints, but they modulate where the eyes move regardless of word boundaries: Evidence against top-down word-based eye-movement control during reading. PLoS ONE 14(7), e0219666 (2019).

32. Brysbaert, M., & Vitu, F. in Eye guidance in reading and scene perception (ed. Underwood, G.) 125–147 (Elsevier Science Ltd, Oxford, 1998).

33. Vitu, F., O’Regan, J.K., Inhoff, A. & Topolski, R. Mindless reading: Eye movement characteristics are similar in scanning strings and reading texts, Percept. Psychophys. 57, 352–364 (1995).

34. Dodge, R. An experimental study of visual fixation. Psychol. Rev-Monogr. S. 4, 1–95 (1907).

35. McConkie, G.W., Kerr, P.W., Reddix, M.D. & Zola, D. Eye movement control during reading: I. The location of initial eye fixations on words. Vision Res. 28(10), 1107–1118 (1988).

36. Nuthmann, A., Vitu, F., Engbert, R. & Kliegl, R. No evidence for a saccadic range effect in visually guided and memory-guided saccades in simple saccade-targeting tasks. PLoS ONE 11(**9**), e0162449 (2016).

37. McIlwain, J.T. Distributed spatial coding in the superior colliculus: A review. Visual Neurosci. 6, 3–13 (1991).

38. Vitu, F., Casteau, S., Adeli, H., Zelinsky, G.J. & Castet, E. The magnification factor accounts for the greater hypometria and imprecision of larger saccades: Evidence from a parametric human-behavioral study. J. Vis. 17(**4**):2, 10.1167/17.4.2. (2017).

39. Findlay, J.M. & Walker, R. A model of saccade generation based on parallel processing and competitive inhibition. Behav. Brain Sci. 22(**4**), 661–720 (1999).

40. Vitu, F. About the global effect and the critical role of retinal eccentricity: Implications for eye movements in reading. J. Eye Movement Res. 2(3), 6, 10.16910/jemr.2.3.6 (2008).

41. Klein, R.M & Hilchey, M.D. in Handbook on eye movements (eds. Liversedge, S.P., Gilchrist, I.D., & Everling, S.) 471–492 (Oxford University Press, 2011).

42. Rayner, K. & McConkie, G.W. What guides a reader’s eye movements? Vision Res. 16, 829–837 (1976).

43. Rayner, K. Eye guidance in reading: Fixation location within words. Perception 8, 21–30 (1979).

44. Huey, E.B. The Psychology and Pedagogy of Reading. (Macmillan, New York, 1908).

45. Yan, M., Kliegl, R., Richter, E.M., Nuthmann, A. & Shu, H. Flexible saccade-target selection in Chinese reading. Q. J. Exp. Psychol. 63, 705–725 (2010).

46. Liu, Y., Reichle, E.D. & Li, X. The effect of word frequency and parafoveal preview on saccade length during the reading of Chinese. J. Exp. Psychol. Hum. Percept. Perform. 42, 1008–1025 (2016).

47. Luna, B., Velanova, K. & Geier, C.F. (2008). Development of eye-movement control. Brain Cogn. 68, 293–308.

48. Nuthmann, A. & Henderson, J.M. Object-based attentional selection in scene viewing. J. Vis. 10(8):20, 10.1167/10.8.20. (2010).

49. McConkie, G.W., et al. in *Vision and visual dyslexia* (ed. Stein, J.F.) 251-262 (Macmillan Press, London, 1991).

50. Sheinberg, D. & Zelinsky, G.J. in Perception and cognition: Advances in eye movement research (eds. d’Ydewalle, G. & Van Rensbergen, J.) 333–348 (North-Holland, Amsterdam, New-York, 1993).

## References

52. Geisler, W.S. & Perry, J.S. in Eye Tracking Research & Applications Symposium (ed. Duchowski, A. T.) 83–87 (New York: Association for Computing Machinery, 2002).

53. Harel, J., Koch, C. & Perona, P. in Adv. Neural Inf. Process. Syst. 19 (eds. Schölkopf, B., Platt, J. & Hofman, T.) 545–552 (NIPS, 2006).

54. Goldberg, M.E. & Wurtz, R.H. Activity of superior colliculus in behaving monkey: I. Visual receptive fields of single neurons. J. Neurophysiol. 35, 542–559 (1972).

55. Cynader, M., & Berman, N. Receptive-field organization of monkey superior colliculus. J. Neurophysiol. 35, 187–201 (1972).

56. Sparks, D.L. & Hartwich-Young, R. The deep layers of the superior colliculus. Rev. Oculomot. Res. 3, 213–255 (1989).

57. Lee, C., Rohrer, W.H. & Sparks, D.L. Population coding of saccadic eye movements by neurons in the superior colliculus. Nature 332, 357–360 (1988).

58. Anderson, R.W., Keller, E.L., Gandhi, N.J. & Das, S. Two-dimensional saccade-related population activity in superior colliculus in monkey. J. Neurophysiol. 80, 798–817 (1998).

59. McIlwain, J.T. Visual receptive fields and their images in superior colliculus of the cat. J. Neurophysiol. 38, 219–230 (1975).

60. Munoz, D.P. & Wurtz, R.H. Saccade-related activity in monkey superior colliculus: II. Spread of activity during saccades. J. Neurophysiol. 73, 2334–2348 (1995).

61. van Opstal, A.J., van Gisbergen, J.A. & Smith, A.C. Comparison of saccades evoked by visual stimulation and collicular electrical stimulation in the alert monkey. Exp. Brain Res. 79, 299–312 (1990).

62. Schiller, P.H. & Stryker, M. Single-unit recording and stimulation in superior colliculus of the alert rhesus monkey. J. Neurophysiol. 35, 915–924 (1972).

63. Robinson, D.A. Eye movements evoked by collicular stimulation in the alert monkey. Vision Res. 12, 1795–1808 (1972).

64. Sparks, D.L., Holland, R. & Guthrie, B.L. Size and distribution of movement fields in the monkey superior colliculus. Brain Res. 113, 21–34 (1976).

65. van Opstal, A.J. & van Gisbergen, J.A. Scatter in the metrics of saccades and properties of the collicular motor map. Vision Res. 29, 1183–1196 (1989).

66. Marino, R.A., Rodgers, C.K., Levy, R. & Munoz, D.P. Spatial relationships of visuomotor transformations in the superior colliculus. J Neurophysiol 100, 2564–2576 (2008).

67. Yang, S.-N. An oculomotor-based model of eye movements in reading: The competition-interaction model. Cogn. Syst. Res. 7, 56–69 (2006).

68. Fraley, C., Raftery, A.E., Murphy, T.B. & Scrucca, L. mclust Version 4 for R: Normal Mixture Modeling for Model-Based Clustering, Classification, and Density Estimation. Technical Report No. 597 (Department of Statistics, University of Washington, 2012).

69. R Development Core Team. R: A Language and Environment for Statistical Computing. (R Foundation for Statistical Computing, Vienna, Austria, 2008).

70. Fraley, C. & Raftery, A.E. Bayesian regularization for normal mixture estimation and model-based clustering. J. Classif 24, 155–181 (2007).

71. Kaas, R.E. & Raftery, A.E. Bayes Factors. J. Am. Stat. Assoc. 90(430), 773–795 (1995).

72. Bates, D., Mächler, M., Bolker, B. & Walker, S. Fitting Linear Mixed-Effects Models Using lme4. J. Stat. Softw. 67(1), 1–48 (2015).

73. Baayen, R.H. Analyzing Linguistic Data: A Practical Introduction to Statistics using R. (Cambridge University Press, Cambridge, 2008).

## References

74. McConkie, G.W. in Processing of visible language Vol. 1 (eds. Kolers, P.A., Wrolstad, M.E. & Bouma, H.) 37–48 (Plenum, New York, 1979).

75. McConkie, G.W. & Rayner, K. The span of the effective stimulus during a fixation in reading. Percept. Psychophys. 17, 578–586 (1975).

76. Legge, G.E., Klitz, T.S. & Tjan, B.S. Mr. Chips: An ideal-observer model of reading. Psychol. Rev. 104, 524–553 (1997).

77. Klitz, T.S., Legge, G.E. & Tjan, B.S. in Reading as a perceptual process (eds. Kennedy, A., Radach, R., Heller, D. & Pynte, J.) 667–682 (Elsevier, Oxford, 2000).

78. Bernard, J.-B., Moscoso Del Prado, F. & Castet, E. Predicting oculomotor strategies in reading with normal and damaged visual fields. Perception 44(S1), 70 (2015).

79. Thibadeau, R., Just, M.A. & Carpenter, P.A. A model of the time course and content of reading. Cogn. Sci. 6, 157–203 (1982).

80. Just, M.A. & Carpenter, P.A. A theory of reading: From eye fixations to Comprehension. Psychol. Rev. 87, 329–354 (1980).

81. Reilly, R. in Perception and cognition: Advances in eye movement research (ed. d’Ydewalle, G. & Van Rensbergen, J.) 191–212 (North Holland, Amsterdam, New York, 1993).

82. Pollatsek, A., Reichle, E.D. & Rayner, K. Tests of the E-Z Reader model: Exploring the interface between cognition and eye-movement control. Cogn. Psychol. 52, 1–56 (2006).

83. Reichle, E. D., Pollatsek, A. & Rayner, K. E-Z Reader: A cognitive-control, serial-attention model of eye-movement behavior during reading. Cogn. Syst. Res. 7, 4–22 (2006).

84. Reichle, E.D., Pollatsek, A. & Rayner, K. Using E-Z Reader to simulate eye movements in non-reading tasks: A unified framework for understanding the eye-mind link. Psychol. Rev. 119(**1**), 155–185 (2012).

85. Reichle, E.D., Warren, T. & McConnell, K. Using E-Z Reader to model the effect of higher-level language processing on eye movements during reading. Psychon. Bull. Rev. 16(**1**), 1–21 (2009).

86. Morrison, R.E. Manipulation of stimulus onset delay in reading: Evidence for parallel programming of saccades. J. Exp. Psychol. Hum. Percept. Perform. 10, 667–682 (1984).

87. Pollatsek, A. & Rayner, K. in Comprehension processes in reading (eds. Balota, D.A., Flores d’Arcais, G.B. & Rayner, K.) 143–163 (Erlbaum, Hillsdale, NJ, 1990).

88. Rayner, K., & Pollatsek, S. The Psychology of Reading. (Prentice-Hall, London, 1989).

89. Reichle, E.D., Pollatsek, A., Fisher, D.L. & Rayner, K. Toward a model of eye movement control in reading. Psychol. Rev. 105(**1**), 125–157 (1998).

90. Reichle, E.D., Rayner, K. & Pollatsek, A. Eye-movement control in reading: accounting for initial fixation locations and refixations within the E-Z reader model. Vision Res. 39, 4403–4411 (1999).

91. Salvucci, D.D. An integrated model of eye movements and visual encoding. Cogn. Syst. Res. 1, 201–220 (2001).

92. Heinzle, J., Hepp, K. & Martin, A.C. A biologically realistic cortical model of eye movement control in reading. Psychol. Rev. 117(**3**), 808–830 (2010).

93. Suppes, P. in Eye movements and their role in visual and cognitive processes (ed. Kowler, E.) 455–477 (Elsevier, Amsterdam, 1990).

94. Richter, E.M., Engbert, R. & Kliegl, R. Current advances in swift. Cogn. Syst. Res. 7, 25–33 (2006).

95. Nuthmann, A. & Engbert, R. Mindless reading revisited: An analysis based on the SWIFT model of eye-movement control. Vision Res. 49, 322–336 (2009).

96. Engbert, R. & Kliegl, R. Mathematical models of eye movements in reading: a possible role for autonomous saccades. Biol. Cybern. 85, 77–87 (2001).

97. Engbert, R., Longtin, A. & Kliegl, R. A dynamical model of saccade generation in reading based on spatially distributed lexical processing. Vision Res. 42, 621–636 (2002).

98. Reilly, R. & Radach, R. in Cognitive and applied aspects of eye movement research (eds. Hyönä, J., Radach, R. & Deubel, H.) 429–455 (Elsevier, Amsterdam, 2003).

99. Snell, J., van Leipsig, S., Grainger, J. & Meeter, M. OB1-Reader: A model of word recognition and eye movements in text reading. Psychol. Rev. 125(**6**), 969–984 (2005).

100. McDonald, S.A., Carpenter, R.H.S. & Shillcock, R.C. An anatomically constrained, stochastic model of eye movement control in reading. Psychol. Rev. 112, 814–840 (2005).

101. O’Regan, J.K. in Eye movements and visual cognition: Scene perception and reading (ed. Rayner, K.) 333–354 (Springer-Verlag, New York, 1992).

102. Reilly, R.G. & O’Regan, J.K. Eye movement control during reading: A simulation of some word targeting strategies. Vision Res, 38(**2**), 303–317 (1998).

103. O’Regan, J.K. & Lévy-Schoen, A. in Attention and performance XII: The psychology of reading (ed. Coltheart, M.) 363–383 (Erlbaum, Hillsdale, NJ, 1987).

104. Vitu, F. The basic assumptions of E-Z reader are not well-founded. Behav. Brain Sci. 26(**4**), 506–507 (2003).

105. Vitu, F. in Handbook on Eye Movements (eds. Liversedge, S.P., Gilchrist, I.D. & Everling, S.) 731–750 (Oxford University Press, 2011).

106. Yang, S.-N., & McConkie, G.W. Eye movements during reading: A theory of saccade initiation times. Vision Res. 41, 3567–3585 (2001).

107. Shebilske, W. in Cognition and motor processes (eds. Prinz, W. & Sanders, A. F.) 99–119 (Springer-Verlag, Berlin, Heidelberg, 1984).

108. Hochberg, J. in Basic studies on reading (eds. Levin, H. & Williams, J.P.) (Basic Books, New York, 1970).

109. Hochberg, J. On the control of saccades in reading. Vision Res. 15 (1975).

110. Hochberg, J. in Eye movements and psychological processes (eds. Monty, R.A. & Senders, J.W.) 397–417 (Erlbaum, Hillsdale, NJ, 1976).

111. Shebilske, W. in Understanding language (ed. Massaro, D.) 291–311 (Academic Press, New York, 1975).

112. Bouma, H., & De Voogd, A.H. On the control of eye saccades in reading. Vision Res. 14, 273–284 (1974).

113. Haber, R.N. in Eye movements and psychological processes (eds. Monty, R.A. & Senders, J.W.) (Erlbaum, Hillsdale, NJ, 1976).

114. Buswell, G.T. An experimental study of the eye-voice span in reading. Suppl. Educ. Monogr. 17 (1920).

115. Buswell, G.T. Fundamental Reading Habits, a Study of their Development (Chicago University Press, 1922).

116. Buswell, G.T. How adults read. Suppl. Educ. Monogr. 45 (1937).

117. Kolers, P.A. in Eye movements and psychological processes (eds. Monty, R.A. & Senders, J.W.) 371–395 (Erlbaum, Hillsdale, NJ, 1976).

118. Liu, Y., Huang, R., Gao, D. & Reichle, E.D. Further tests of a dynamic-adjustment account of saccade targeting during the reading of Chinese. Cogn. Sci. 41(**6**), 1624–1287 (2017).

119. Li, X., Liu, P. & Rayner, K. (2011). Eye movement guidance in Chinese reading: Is there a preferred viewing location? Vision Res. 51, 1146–1156.

120. Rayner, K., Li, X.S. & Pollatsek A. Extending the E-Z Reader model of eye movement control to Chinese readers. Cogn. Sci. 31: 1021–1033 (2007).

121. Engbert, R. & Krügel, A. Readers use Bayesian estimation for eye-movement control. Psychol. Sci. 21, 366–371 (2010).

122. Kerr, P.W. Eye movement control during reading: The selection of where to send the eyes. University of Illinois at Urbana-Champaign (1992).

123. McConkie, G.W., Kerr, P.W. & Dyre, B.P. in Eye movements in reading (eds. Ygge, J. & Lennerstrand, G.) 315–328 (Pergamon Press, Oxford, 1994).

124. Feng, G. Eye movements as time-series random variables: A stochastic model of eye-movement control in reading. Cogn. Syst. Res. 7, 70–95 (2006).

125. New, B., Pallier, C., Ferrand, L. & Matos, R. Une base de données lexicales du français contemporain sur interne: Lexique [A lexical data base of contemporary French on the Web: Lexicon]. L’Année Psychologique 101, 447–462 (2001).

126. Brysbaert, M., Drieghe, D. & Vitu, F. in Cognitive Processes in eye guidance (ed. Underwood, G.) 53–77 (Oxford University Press, 2005).

127. Rayner, K. & Fischer, M.H. Mindless reading revisited: Eye movements during reading and scanning are different. Percept. Psychophys. 58(**5**), 734–747 (1996).

128. Luke, S.G. & Henderson, J.M. Oculomotor and cognitive control of eye movements in reading: Evidence from mindless reading. Atten. Percept. Psychophys. 75(**6**), 1230–1242 (2013).

129. Tsai, J.L. & McConkie, G.W. in The mind’s eye: Cognitive and applied aspects of eye movement research (eds. Hyönä, J., Radach, R. & Deubel, H.) 159–176 (Elsevier, Oxford, 2003).

130. Hyönä, J., Yan, M. & Vainio, S. Morphological structure influences the initial landing position in words during reading Finnish. Q. J. Exp. Psychol. 71(**1**), 122–130 (2013).

131. Lavigne, F., Vitu, F. & d’Ydewalle, G. The influence of semantic context on initial eye landing sites in words. Acta Psychol. 104, 191–204 (2000).

132. Rayner, K, Reichle, E.D., Stroud, M.J., Williams, C.C. & Pollatsek, A. The effect of word frequency, word predictability, and font difficulty on the eye movements of young and older readers. Psychol. Aging 21(**3**), 448–465 (2006).

133. Yan, M. et al. Eye movements guided by morphological structure: Evidence from the Uighur language. Cognition 132, 181–215 (2014).

134. Zhou, W., Wang, A., Shu, H., Kliegl, R. & Yan, M. Word segmentation by alternating colors facilitate eye guidance in Chinese reading. Mem. Cogn. 46, 729–740 (2018).

135. Liu, Y., Guo, S., Yu, L., Reichle, E.D. Word predictability affects saccade length in Chinese reading: An evaluation of the dynamic-adjustment model. Psychon. Bull. Rev. 25: 1891–1899 (2018).

136. McConkie, G.W., Kerr, P. W., Reddix, M.D., Zola, D. & Jacobs, A.M. Eye movement control during reading: II. Frequency of refixating a word. Percept. Psychophys. 46, 245–253 (1989).

137. Vitu, F. The influence of parafoveal preprocessing and linguistic context on the optimal landing position effect. Percept. Psychophys. 50, 58–75 (1991).

138. Nazir, T. On the role of refixations in letter strings: The influence of oculomotor factors. Percept. Psychophys. 49(**4**), 373–389 (1991).

139. Andriessen, J.J. & De Voogd, A.H. Analysis of eye movement patterns in silent reading. IPO Annual Progress Report 8, 29–34 (1973).

140. Vitu, F. & McConkie, G.W. in Reading as a perceptual process (eds. Kennedy, A., Radach, R., Heller, D. & Pynte, J.) 301–326 (Elsevier, Oxford, 2000).

141. Vitu, F., McConkie, G.W. & Zola, D. in Eye guidance in reading and scene perception (ed. Underwood, G.) 101–124 (Elsevier Science Ltd, Oxford, 1998).

142. Vitu, F. in Cognitive processes in eye guidance (ed. Underwood, G.) 1–32 (Oxford University Press, 2005).

143. Radach, R. & McConkie, G.W. in Eye guidance in reading and scene perception (ed. Underwood, G.) 77–100 (Elsevier Science Ltd, Oxford, 1998).

144. Laurinavichyute, A.K., Sekerina, I.A., Alexeeva, S., Bagdasaryan, K. & Kliegl, R. Russian sentence corpus: Benchmark measures of eye movements in reading in Russian. Behav. Res. Methods 51, 1161–1178 (2019).

145. Deutsch, A. & Rayner, K. Initial fixation location effects in reading Hebrew words. Lang. Cognitive Proc. 14, 393–421 (1999).

146. Hyönä, J. & Bertram, R. Optimal viewing position effects in reading Finnish. Vision Res. 51, 1279–1287 (2011).

147. Nuthmann, A., Engbert, R. & Kliegl, R. Mislocated fixations during reading and the inverted optimal viewing position effect. Vision Res. 45, 2201–2217 (2005).

148. Radach, R. & Kempe, V. in Perception and cognition: Advances in eye movement research (eds. d’Ydewalle, G. & Van Rensbergen, J.) 213–225 (North-Holland, Amsterdam, New York, 1993).

149. Paterson, K.B., Almabruk, A.A.A., McGowan, V.A., White, S.J. & Jordan, T.R. Effects of word length on eye movement control: The evidence from Arabic. Psychon. Bull. Rev. 22, 1443–1450 (2015).

150. Kajii, N., Nazir, T.A. & Osaka, N. Eye movement control in reading unspaced text: The case of the Japanese script. Vision Res. 41, 2503–2510 (2001).

151. Sainio, M., Hyönä, J., Bingushi, K. & Bertram, R. The role of inter-word spacing in reading Japanese: An eye movement study. Vision Res. 47, 2575–2584 (2007).

152. Rayner, K., Sereno, S.C. & Raney, G.E. Eye movement control in reading: A comparison of two types of models. J. Exp. Psychol. Hum. Percept. Perform. 22(**5**), 1188–1200 (1996).

153. Vitu, F., McConkie, G.W., Kerr, P. & O’Regan, J.K. Fixation location effects on fixation durations during reading: An inverted optimal viewing position effect. Vision Res. 41, 3513–3533 (2001).

154. Vitu, F., O’Regan, J.K. & Mittau, M. Optimal landing position in reading isolated words and continuous text. Percept. Psychophys. 47(**6**), 583–600 (1990).

155. Epelboim, J., Booth, J.R. & Steinman, R.M. Reading unspaced text: Implications for theories of reading eye movements. Vision Res. 34, 1735–1766 (1994).

156. McGowan, V.A., White, S.J., Jordan, T.R. & Paterson, K.B. Aging and the use of interword spaces during reading: Evidence from eye movements. Psychon. Bull. Rev., 21(**3**), 740–747 (2014).

157. Mirault, J., Snell, J. & Grainger, J. Reading without spaces revisited: The role of word identification and sentence-level constraints. Acta Psychol. 195, 22–29 (2019).

158. Perea, M. & Acha, J. Space information is important for reading. Vision Res. 49, 1994– 2000 (2009).

159. Rayner, K., Fischer, M.H. & Pollatsek, A. Unspaced text interferes with both word identification and eye movement control. Vision Res. 38, 1129–1144 (1998).

160. Rayner, K., Yang, J., Schuett, S. & Slattery, T. Eye movements of older and younger readers when reading unspaced text. Exp. Psychol. 60(**5**), 354–361 (2013).

161. Veldre, A., Drieghe, D. & Andrews, S. Spelling ability selectively predicts the magnitude of disruption in unspaced text reading. J. Exp. Psychol. Hum. Percept. Perform. 43(**9**), 1612–1628 (2017).

162. Sheridan, H., Rayner, K. & Reingold, E.M. Unsegmented text delays word identification: Evidence from a survival analysis of fixation durations. Vis. Cogn. 21(**1**), 38–60 (2013).

163. Bai, X.J., Yan, G.L., Liversedge, S.P., Zang, C.L. & Rayner, K. Reading spaced and unspaced Chinese text: Evidence from eye movements. J. Exp. Psychol. Hum. Percept. Perform. 34(**5**), 1277–1287 (2008).

164. Shen, D. et al. Eye movements of second language learners when reading spaced and unspaced Chinese text. J. Exp. Psychol. Appl. 18, 192–202 (2012).

165. Morris, R.K., Rayner, K. & Pollatsek, A. Eye movement guidance in reading: The role of parafoveal letter and space information. J. Exp. Psychol. Hum. Percept. Perform. 16, 268–281 (1990).

166. Pollatsek, A. & Rayner, K. Eye movement control in reading: The role of word boundaries. J. Exp. Psychol. Hum. Percept. Perform. 8(**6**), 817–833 (1982).

167. McConkie, G.W. & Rayner, K. The span of the effective stimulus during a fixation in reading. Percept. Psychophys. 17, 578–586 (1975).

168. Winskel, H., Radach, R. & Luksaneeyanawin, S. Eye movements when reading spaced and unspaced Thai and English: A comparison of Thai-English bilinguals and English monolinguals. J. Mem Lang. 61, 339–351 (2009).

169. Zang, C., Liang, F., Bai, X., Yan, G. & Liversedge, S.P. Interword spacing and landing position effects during Chinese reading in children and adults. J. Exp. Psychol. Hum. Percept. Perform. 39(**3**), 720–734 (2013).

170. Yan, M. & Kliegl, R. CarPrice versus CarpRice: Word boundary ambiguity influences saccade target selection during the reading of Chinese sentences. J. Exp. Psychol. Learn. Mem. Cogn. 42(**11**), 1832–1838 (2016).

171. Inhoff, A.W. & Wu, C. Eye movements and the identification of spatially ambiguous words during Chinese sentence reading. Mem. Cogn. 33(**8**),1345–1356 (2005).

172. Bullimore, M.A. & Bailey, I.L. Reading and eye movements in age-related maculopathy. Optom. Vis. Sci. 72(**2**), 125–138 (1995).

173. Kolers, P.A., Duchnicky, R.L. & Ferguson, D.C. Eye movement measurement of readability of CRT displays. Hum. Factors 23(**5**), 517–527 (1981).

174. Morrison, R.E. & Rayner, K. Saccade size in reading depends upon character spaces and not visual angle. Percept. Psychophys. 30, 395–396 (1981).

175. Paterson, D.G. & Tinker, M.A. The effect of typography upon the perceptual span in reading. Am. J. Psychol. 60(**3**), 388–396 (1947).

176. Lamare, M. Des mouvements des yeux dans la lecture. Bull. Mem. Soc. Fr. Ophtalmol. 10, 354–364 (1892).

177. Landolt, E. Nouvelles recherches sur la physiologie des mouvements des yeux. Archives d’Ophtalmologie, Sept-oct., 385–395 (1891).

178. O’Regan, J.K., Lévy-Schoen, A. & Jacobs, A. The effect of visibility on eye movement parameters in reading. Percept. Psychophys. 34, 457–464 (1983).

179. Yan, M., Zhou, W., Shu, H. & Kliegl, R. Perceptual span depends on font size during the reading of Chinese sentences. J. Exp. Psychol. Learn. Mem. Cogn. 41(**1**), 209–219 (2015).

180. Franken, G., Podlesek, A. & Mozina, K. Eye-tracking study of reading speed from LCD displays: Influence of type style and type size. J. Eye Movement Res. 8(**1**):**3**, 1–8 (2015).

181. Yen, N.-S., Tsai, J.-L., Chen, P.-L., Lin, H.-Y. & Chen, A.L.P. Effects of typographic variables on eye-movement measures in reading Chinese from a screen. Behav. Inform. Technol. 30(**6**), 797–808 (2011).

182. McDonald, S.C. Effects of number-of-letters on eye movements during reading are independent from effects of spatial word length. Vis. Cogn. 13(**1**), 89–98 (2006).

183. Hautala, J., Hyönä, J. & Aro, M. Dissociating spatial and letter-based word length effects observed in readers’ eye movement patterns. Vision Res. 51, 1719–1727 (2011).

184. Vonk, W., Radach, R. & van Rijn, H. in Reading as a perceptual process (eds. Kennedy, A., Radach, R., Heller, D. & Pynte, J.) 269–299 (Elsevier, Oxford, 2004).

