## Supplementary Information for "Readers move their eyes mindlessly using midbrain visuo-motor principles"

**Supplementary Methods 1. Statistical comparison of MASC and FSC readers 6**

**Word-skipping behavior…………………………………………………………….... 9**
- The relationship between word skipping and word length…………………………... 9
- The relationship between word skipping and launch site…………………………….... 10

**Within-word landing positions………………………………………………………. 11**
- The PVL effect………………………………………………………………………… 11
- The Launch-Site effect……………………………………………………………….. 13

**All landing positions regardless of word boundaries……………………………………. 16**
- Methodological Note…………………………………………………………………. 16
- The Launch-Site effect revisited……………………………………………………... 16

- The relationship between word skipping and launch site………………………………. 34

**Within-word landing positions………………………………………………………. 38**
- The PVL effect………………………………………………………………………… 38
- The Launch-Site effect……………………………………………………………….. 43

**All landing positions: The Launch-Site effect revisited…………………………………. 49**

**Within-word refixation behavior: The OVP effect……………………………………… 57**

**Supplementary Methods 3. The respective roles of inter-word spacing and print size 62**

**MASC's predicted effects of inter-word spacing………………………………………... 62**
- Saccade length……………………………………………………………………. 63
- Word-skipping behavior………………………………………………………….. 66
- The PVL effect…………………………………………………………………… 69
- The launch-site effect…………………………………………………………….. 72
- The Refixation-OVP effect……………………………………………………… 74

**MASC' predicted effects of character size……………………………………………. 76**
- Saccade length……………………………………………………………………. 76
- Word-skipping behavior………………………………………………………….. 79
- The PVL effect…………………………………………………………………… 81
- The launch-site effect…………………………………………………………….. 85
- The Refixation-OVP effect……………………………………………………… 87

**Supplementary References 89**

**List of Supplementary Tables**

**Supplementary Table 1:** GMM-estimated shape of the distributions of saccade lengths in MASC and FSC readers………………………………………….... **7**

**Supplementary Table 2:** GLM regression coefficients for the proportion of regressions in MASC and FSC readers………………………………………….... **7**

**Supplementary Table 3:** LM regression coefficients for the mean and SD of the length of regressive saccades in MASC and FSC readers…………………… **8**

**Supplementary Table 4:** LM regression coefficients for the mean and SD of the length of progressive saccades in MASC and FSC readers………………….. **8**

**Supplementary Table 5:** Fixed effects of GLMMs for the probability of word skipping by word length in MASC and FSC readers…………………………… **9**

**Supplementary Table 6:** Fixed effects of GLMMs for the probability of word skipping by word length and launch site in MASC and FSC readers…………... **10**

**Supplementary Table 7:** GMM-estimated shape of within-word landing-site distributions by word length in MASC and FSC readers………………………... **12**

**Supplementary Table 8:** Fixed effects of LMMs for the mean and SD of within-word landing sites by word length in MASC and FSC readers………….. **13**

**Supplementary Table 9:** GMM-estimated shape of within-word landing-site distributions by word length and launch site in MASC and FSC readers……..… **14**

**Supplementary Table 10:** Fixed effects of LMMs for the mean and SD of within-word landing sites by word length and launch site in MASC and FSC readers………………………………………………………. **15**

**Supplementary Table 11:** GMM-estimated shape of overall landing-site distributions by word length and launch site in MASC and FSC readers………… **17**

**Supplementary Table 12:** Fixed effects of LMMs for the mean and SD of all landing sites by word length and launch site in MASC and FSC readers……... **19**

**Supplementary Table 13:** Fixed effects of GLMMs for the probability of within-word refixations by initial landing position and word length in MASC and FSC readers………………………………………… **21**

**Supplementary Table 14:** GMM-estimated shape of the distributions of within-word landing sites following regressions in MASC and FSC readers…. **23**

**Supplementary Table 15:** Fixed effects of LMMs for the mean and SD of within-word landing sites following regressions in MASC and FSC readers…. **24**

**Supplementary Table 16:** GMM-estimated shape of the distributions of saccade lengths in (comparison) models and FSC readers……………………... **26**

**Supplementary Table 17:** GLM regression coefficients for the proportion of regressions in (comparison) models and FSC readers……………………... **27**

**Supplementary Table 18:** LM regression coefficients for the mean and SD of the length of regressive saccades in (comparison) models and FSC readers…... **29**

**Supplementary Table 19:** LM regression coefficients for the mean and SD of the length of progressive saccades in (comparison) models and FSC readers…. **31**

**Supplementary Table 20:** Fixed effects of GLMMs for the probability of word skipping by word length in (comparison) models and FSC readers…………... **33**

**Supplementary Table 21:** Fixed effects of GLMMs for the probability of word skipping by word length and launch site in (comparison) models and FSC readers…………………………………………………………. **36**

**Supplementary Table 22:** GMM-estimated shape of within-word landing-site distributions by word length in (comparison) models and FSC readers……….. **39**

**Supplementary Table 23:** Fixed effects of LMMs for the mean and SD of within-word landing sites by word length in (comparison) models and FSC readers…………………………………………………………. **41**

**Supplementary Table 24:** GMM-estimated shape of within-word landing-site distributions by word length and launch site in (comparison) models and FSC readers………………………………………………………. **44**

**Supplementary Table 25:** Fixed effects of LMMs for the mean and SD of within-word landing sites by word length and launch site in (comparison) models and FSC readers……………………………………….. **46**

**Supplementary Table 26:** GMM-estimated shape of overall landing-site distributions by word length and launch site in (comparison) models and FSC readers……………………………………………………………. **50**

**Supplementary Table 27:** Fixed effects of LMMs for the mean and SD of all landing sites by word length and launch site in (comparison) models and FSC readers………………………………………………………. **53**

**Supplementary Table 28:** Fixed effects of GLMMs for the probability of within-word refixations by initial landing position and word length in (comparison) models and FSC readers…………………………... **59**

**Supplementary Table 29:** LM regression coefficients for the mean and SD of the length of MASC's progressive saccades by inter-word spacing………... **65**

**Supplementary Table 30:** Fixed effects of LMMs for the length of progressive saccades by line length in MASC and FSC readers……………………….. **66**

**Supplementary Table 31:** Fixed effects of GLMMs for MASC's word-skipping probability by word length and inter-word spacing………………………….. **67**

**Supplementary Table 32:** Fixed effects of GLMMs for MASC's word-skipping probability by word length, launch site, and inter-word spacing…………….. **68**

**Supplementary Table 33:** Fixed effects of LMMs for the mean and SD of MASC's within-word landing sites by word length and inter-word spacing……… **71**

**Supplementary Table 34:** Fixed effects of LMMs for the mean and SD of MASC's overall landing-site distributions by word length, launch site, and inter-word spacing……………………………………………….. **73**

**Supplementary Table 35:** Fixed effects of GLMMs for MASC's within-word refixation probability by initial landing position, word length, and inter-word spacing……………………………………………….. **75**

**Supplementary Table 36:** LM regression coefficients for the mean and SD of the length of MASC's progressive saccades by character size……………... **78**

**Supplementary Table 37:** Fixed effects of GLMMs for MASC’s word-skipping probability by word length and character size………………………………….. **79**

**Supplementary Table 38:** Fixed effects of GLMM for MASC’s word-skipping probability by word length, launch site, and character size………………….. **81**

**Supplementary Table 39:** Fixed effects of LMMs for the mean and SD of MASC’s within-word landing sites by word length and character size…………… **84**

**Supplementary Table 40:** Fixed effects of LMMs for the mean and SD of MASC’s overall landing-site distributions by word length, launch site, and character size…………………………………………………….. **86**

**Supplementary Table 41:** Fixed effects of GLMM for MASC’s within-word refixation probability by initial landing position, word length, and character size…………………………………………………….. **87**

### Supplementary Methods 1 | Statistical comparison of MASC and FSC readers

In the analyses reported in this section, the oculomotor behavior of MASC and FSC readers was compared after matching both data sets for numbers of fixations. MASC indeed made on average more fixations per sentence than FSC readers (see Methods), being deprived of a comprehension-based termination criterion. For a given model run and sentence, the number of fixations considered for analysis was randomly sampled from the distribution of the numbers of fixations per sentence for FSC readers in the corresponding sentence pair. Results were comparable to the results obtained in a second set of analyses including all comparison models and using a different data-matching procedure (i.e., data sets were matched by considering only first-pass fixations/saccades over sentences; see Supplementary Methods 2).

#### Saccade-length distributions

Gaussian-Mixture-Models (GMM) were first fitted to each individual distribution of saccade lengths in FSC and MASC data sets, yielding four GMM-parameter estimates (the number of estimated mixture components, as well as the mean, the standard deviation (SD), and the proportion of cases (*k* value) for each detected mode). These estimates, and other derived measures (see below), were compared between MASC and FSC readers using (Generalized) Linear Models ((G)LM) whenever possible.

**Typical bimodal distributions.** As shown in Supplementary Table 1, the great majority of the distributions in FSC and MASC data sets were best fitted with two (one negative and one positive) mixture components. In a very few cases in FSC readers, more than two components were detected, but the largest component (associated with the largest *k* estimate) for regressive as well as progressive saccades accounted on average for more than 96% of the data (Fig. 2a). This is consistent with the well-established bimodal trend for saccade-length distributions[1].

**Supplementary Table 1 | GMM-estimated shape of the distributions of saccade lengths in MASC and FSC readers.**

| | ALL SACCADES | | | | REGRESSIONS | | PROGRESSIONS | |
|---|---|---|---|---|---|---|---|---|
| **SL: MODES** | **1** | **2** | **3** | **4** | **2** | **Ratio** | **2** | **Ratio** |
| **FSC** | 0.000 | 0.900 | 0.075 | 0.025 | 0.025 | 0.988 | 0.100 | 0.966 |
| **MASC** | 0.000 | 1.000 | 0.000 | 0.000 | 0.000 | 1.000 | 0.000 | 1.000 |

Proportion of 1-4 mixture components in individual distributions, as estimated based on GMM (left four columns), and proportion of two mixture components, as well as mean ratios between the largest *k* estimate and the sum of *k* estimates, separately for regressive and forward saccades (right four columns), for FSC and MASC (rows). These analyses relied on a total number of 60630 and 38734 cases across subjects, in FSC and MASC respectively.

**Proportion of regressive saccades.** In Supplementary Table 2, the proportion of regressive saccades was compared between MASC and FSC data sets, by fitting GLMs to GMM-estimated regression rates (the sum of *k* estimates associated with a negative component). The intercept logit estimate, of about -1.58563 when Data Set was at its reference FSC level, indicated that regressions occurred in about 17% of the cases in FSC readers. This proportion was significantly greater in MASC (logit: 0.47632; 25%), thus indicating that MASC made slightly more regressions than FSC readers.

**Supplementary Table 2 | GLM regression coefficients for the proportion of regressions in MASC and FSC readers**

| REGRESSION RATE | Estimate | Std. Error | z value | Pr(>\|z\|) | Proportion |
|---|---|---|---|---|---|
| **FSC -(Intercept)** | -1.58563 | 0.04209 | -37.67005 | 0.00000 | 0.170 |
| **MASC** | 0.47632 | 0.06673 | 7.13809 | 0.00000 | 0.248 |

A GLM was fitted to the GMM-estimated proportion of regressions, with Data Set (with 2 levels: FSC and MASC) as a categorical predictor. Estimates and standard errors are expressed in logit unit; these can be back transformed into probabilities, using the inverse logit formula: $p = \exp^{(x)} / (1 + \exp^{(x)})$. The intercept (first line) gives the regression-rate estimate in logit units when Data Set was at its reference level: FSC; the slope estimate for MASC (second line) is the value that should be added to the intercept to get MASC's regression rate in logit unit. Corresponding regression likelihood is given in the rightmost column for each data set.

**Length of regressive saccades.** LMs were fitted respectively to the GMM-estimated mean and SD of the largest mixture component associated with regressive saccades. Estimates shown in Supplementary Table 3A-B indicate that regressions were on average of about three letters in FSC readers (intercept estimate: -3.30850). In MASC, they were much larger (estimate: -4.38332), but only slightly less variable (estimate: -1.41081).

**Supplementary Table 3 | LM regression coefficients for the mean and standard deviation (SD) of the length of regressive saccades in MASC and FSC readers.**

| A- REGRESSIONS: MEAN | Estimate | Std. Error | t value | Pr(>\|t\|) |
|---|---|---|---|---|
| **FSC -(Intercept)** | -3.30850 | 0.32175 | -10.28297 | 0.00000 |
| **MASC** | -4.38332 | 0.55728 | -7.86557 | 0.00000 |

| B- REGRESSIONS: SD | Estimate | Std. Error | t value | Pr(>\|t\|) |
|---|---|---|---|---|
| **FSC -(Intercept)** | 4.90710 | 0.29741 | 16.49971 | 0.00000 |
| **MASC** | -1.41081 | 0.51512 | -2.73878 | 0.00818 |

LMs were fitted separately to the GMM-estimated mean (A) and SD (B) of the largest negative component in individual distributions of saccade lengths (in letters), with Data Set (2 levels: FSC and MASC) as a categorical predictor. FSC was the reference level.

**Length of progressive saccades.** MASC's forward saccades even more closely matched those observed in FSC readers. As reported in Supplementary Table 4A, the mean length of progressive saccades was of about 8.43 letters in FSC readers, as classically reported[1]. This was only about 1.14 letters smaller in MASC. In addition, the variability in progressive saccade length did not differ significantly between both data sets ($p = 0.45053$; Supplementary Table 4B).

**Supplementary Table 4 | LM regression coefficients for the mean and SD of the length of progressive saccades in MASC and FSC readers.**

| A- PROGRESSIONS: MEAN | Estimate | Std. Error | t value | Pr(>\|t\|) |
|---|---|---|---|---|
| **FSC –(Intercept)** | 8.43300 | 0.18022 | 46.79243 | 0.00000 |
| **MASC** | -1.14105 | 0.31215 | -3.65543 | 0.00055 |

| B- PROGRESSIONS: SD | Estimate | Std. Error | t value | Pr(>\|t\|) |
|---|---|---|---|---|
| **FSC –(Intercept)** | 2.99550 | 0.07439 | 40.26592 | 0.00000 |
| **MASC** | 0.09788 | 0.12885 | 0.75966 | 0.45053 |

LMs were fitted separately to the GMM-estimated mean (A) and SD (B) of the largest positive component in individual distributions of saccade lengths (in letters), with Data Set (2 levels: FSC and MASC) as a categorical predictor. FSC was the reference level.

### Word-skipping behavior

**The relationship between word skipping and word length.** Previous studies reported a reduction in the likelihood of word skipping with increasing word length[2]. Here we tested this relationship by fitting a first GLMM to the data, using data set and word length as predictors of word skipping rate. The model's fixed effects are presented in Supplementary Table 5. The intercept estimate, of about 0.29826 when FSC was the reference, indicates that readers skipped 4-letter words (the reference, mean, value) in about 57% of the cases (Panel 1). The skipping rate was significantly lower in MASC (by about 10%; logit: -0.42286; Panel 1). Still, both FSC readers and MASC showed a significant decrease in word-skipping rate with increasing word length (logit: -0.75995 and -0.47735; Panels 1 and 2 respectively). This effect remained a bit smaller in MASC, as indicated by the positive slope estimate for the interaction between word length and MASC (logit: 0.28259; Panel 1; Fig. 2b). This was likely due to MASC lacking language-related processes that facilitate the identification of shorter (and usually more frequent/predictable) words and in turn mildly inflate the likelihood these words are skipped (Extended Data Fig. 1a,c-e).

**Supplementary Table 5 | Fixed effects of GLMMs for the probability of word skipping by word length in MASC and FSC readers.**

| SKIPPING BY WL | Estimate | Std. Error | z value | Pr(>\|z\|) | Proportion |
|---|---|---|---|---|---|
| **(1) FSC -(Intercept)** | 0.29826 | 0.09053 | 3.29473 | 0.00099 | 0.574 |
| **WL** | -0.75995 | 0.02634 | -28.84923 | 0.00000 | |
| **MASC** | -0.42286 | 0.15350 | -2.75476 | 0.00587 | 0.469 |
| **WL:MASC** | 0.28259 | 0.04449 | 6.35142 | 0.00000 | |
| **(2) MASC -(Intercept)** | -0.12461 | 0.12634 | -0.98630 | 0.32398 | |
| **WL** | -0.47735 | 0.03626 | -13.16460 | 0.00000 | |

GLMMs were fitted to a binary variable indicating whether a given word was skipped. The fixed structure included Data Set (2 levels: FSC and MASC), Word Length (WL; 1-11 letters; reference (mean) value: 4.01 letters), and the interaction between Data Set and WL (indicated by a colon), as predictors. The random structure included a random intercept by subject, sentence pair, and word, as well as a random effect of WL by subject. Estimates and standard errors are expressed in logit unit. Intercept estimates are converted into probabilities in the rightmost column (in Panel 1), thus giving an estimate of the likelihood of word skipping for each data set, when WL was at its reference value. Panel 1: Fixed-effects for the main GLMM, with Data-Set reference level set to FSC. Panel 2: Data-Set reference level set to MASC. In Panel 2, the fixed effects that were redundant with those reported in Panel 1 were dropped. These analyses relied on a total number of 35216 and 19973 cases across subjects in FSC and MASC respectively.

**The relationship between word skipping and launch site.** As reported in several studies, word-skipping rate is also a function of saccades' launch-site distance relative to the beginning of words, being progressively lower as saccades are initiated from closer to the words' beginning[3-4]. To test whether MASC predicted this relationship, a second GLMM was fitted to the data, with data set, word length, and saccadic launch-site distance as predictors, but considering a smaller range of word lengths than in the above GLMM given floor and ceiling effects. The resulting fixed effects are presented in Supplementary Table 6.

**Supplementary Table 6 | Fixed effects of GLMMs for the probability of word skipping by word length and launch site in MASC and FSC readers.**

| SKIPPING BY LS | Estimate | Std. Error | z value | Pr(>\|z\|) | Proportion |
|---|---|---|---|---|---|
| **(1) FSC -(Intercept)** | 1.24684 | 0.11585 | 10.76266 | 0.00000 | 0.77673 |
| **WL** | -0.98865 | 0.03168 | -31.21106 | 0.00000 | |
| **LS** | 0.52263 | 0.03315 | 15.76712 | 0.00000 | |
| **MASC** | -0.87767 | 0.19412 | -4.52116 | 0.00001 | 0.59126 |
| **WL:LS** | -0.05045 | 0.00935 | -5.39806 | 0.00000 | |
| **WL:MASC** | 0.37887 | 0.05258 | 7.20614 | 0.00000 | |
| **LS:MASC** | 0.21627 | 0.05632 | 3.84023 | 0.00012 | |
| **WL:LS:MASC** | 0.08544 | 0.01497 | 5.70588 | 0.00000 | |
| **(2) MASC -(Intercept)** | 0.36910 | 0.15998 | 2.30712 | 0.02105 | |
| **WL** | -0.60980 | 0.04231 | -14.41419 | 0.00000 | |
| **LS** | 0.73889 | 0.04565 | 16.18679 | 0.00000 | |
| **WL:LS** | 0.03498 | 0.01190 | 2.94110 | 0.00327 | |

GLMMs were fitted to a binary variable indicating whether a given word was skipped. The fixed structure included Data Set (2 levels: FSC and MASC), Word Length (WL; 2-6 letters; reference (mean) value: 3.25), saccadic Launch-Site distance (LS; 0-6 letters from the space in front of the words; reference (mean) value: -2.60 letters) and all interactions. The random structure included a random intercept by subject and by sentence pair, as well as random effects of WL and LS by subject; with a random intercept by word, the model did not converge. Estimates and standard errors are expressed in logit units; intercept estimates are converted into probabilities in the rightmost column (in Panel 1), thus giving an estimate of the likelihood of word skipping for each data set, when WL and LS were at their reference value. Panel 1: Fixed-effects of the main GLMM, with Data-Set reference level set to FSC. Panel 2: Data-Set reference level set to MASC. These analyses relied on a total of 24208 and 14561 cases across subjects in FSC and MASC respectively.

In FSC readers, the likelihood of word skipping, of about 78 % (logit: 1.24684) when both word length and launch-site distance were at their reference (mean) value (3.25 letters and -2.60 letters from the space in front of the words respectively), decreased with increasing word length (logit: -0.98865), and increasing launch-site distance (logit: 0.52263; recall the negative sign of launch site; Panel 1; Fig.2c). Both effects were present in MASC (logit: -0.60980 and 0.73889 respectively; Panel 2). Still, as suggested by the positive slope estimate for the interaction between word length and MASC (logit: 0.37887; Panel 1), the effect of word length was again slightly smaller than in FSC readers (Supplementary Table 5). Moreover, the effect of launch-site distance was larger (logit: 0.21627; Panel 1), and even more so as word length increased, as suggested by the three-way interaction between word length, launch-site distance, and MASC (logit: 0.08544). These small differences were again mostly due to mild top-down, language-related, modulations of word-skipping behavior in FSC readers (Extended Data Fig. 1b).

### Within-word landing positions

**The PVL effect.** As well known, when readers move their eyes forward along lines of text, they tend to preferentially fixate near the center of short words, and slightly to the left of the center of longer words in languages read from left to right, although sometimes landing towards the very beginning or end of words[5]. The resulting typical PVL distribution is normal in shape; it shifts towards the words' beginning, but also becomes wider, as words are longer.

Accordingly, as shown in Supplementary Table 7, most individual within-word landing-position distributions in FSC readers were best fitted with a single Gaussian mixture component (proportion of two mixture components <= 0.10; Fig. 2d). Similarly, MASC's distributions were for the most unimodal. The proportion of two mixture components was slightly higher, particularly in 6- to 8-letter words (<= 0.35), but the largest mixture component accounted on average for more than 86% of the data (>= 96% in FSC).

**Supplementary Table 7 | GMM-estimated shape of within-word landing-site distributions by word length in MASC and FSC readers.**

| **PVL EFFECT** | **4-lw** | | **5-lw** | | **6-lw** | | **7-lw** | | **8-lw** | | **9-lw** | |
|---|---|---|---|---|---|---|---|---|---|---|---|---|
| **2 Modes/ Largest *K*** | **2M** | $K_k$ | **2M** | $K_k$ | **2M** | $K_k$ | **2M** | $K_k$ | **2M** | $K_k$ | **2M** | $K_k$ |
| **FSC** | 0.00 | 1.00 | 0.02 | 0.99 | 0.10 | 0.96 | 0.07 | 0.97 | 0.10 | 0.96 | 0.02 | 0.99 |
| **MASC** | 0.00 | 1.00 | 0.05 | 0.97 | 0.25 | 0.90 | 0.35 | 0.86 | 0.20 | 0.91 | 0.00 | 1.00 |

Proportion of two mixture components (i.e., the maximal number of detected modes; "2M") and mean proportion of cases accounted for by the largest mixture component (mean of largest *k* values; "$K_k$"), as estimated after fitting GMMs to individual distributions of landing positions in 4- to 9-letter words separately for FSC and MASC. These analyses relied on a total of 22434 and 10910 cases across subjects in FSC and MASC respectively.

Further comparisons, based on GMM-estimated mean and SD of the largest mixture component, are presented in the left and right panels of Supplementary Table 8. These confirmed the previously reported relationship between mean landing position and word length in both FSC readers and MASC. As shown in the left panels, FSC readers landed on average slightly to the left of the words' center (estimate: -0.82824, when word length was at its reference (mean) value; 6.5 letters), and even more so as word length increased (estimate: -0.17202; Panel 1). In a similar manner, MASC exhibited a leftward fixation bias (estimate: -0.60055; Panel 2). Moreover, although the bias was significantly less than for FSC readers (Panel 1), it increased with increasing word length (estimate: -0.10866; Panel 2) at about the same rate as for FSC readers, as suggested by the non-significant interaction between MASC and word length ($t$ = 1.51957; Panel 1). On the other hand, as shown in the right panels of Supplementary Table 8, the variability in within-word landing positions differed between FSC readers and MASC by only about 0.09 letter (Panel 1), and it significantly increased with increasing word length in both data sets (estimates: 0.14650 and 0.20594 respectively; Panels 1-2); the spread increase with word length was only slightly greater in MASC compared to FSC readers (estimate: 0.05944; Panel 1).

**Supplementary Table 8 | Fixed effects of LMMs for the mean and SD of within-word landing sites by word length in MASC and FSC readers**.

| PVL EFFECT: MEAN / SD | Estimate | Std. Error | *t* value | Estimate | Std. Error | *t* value |
|---|---|---|---|---|---|---|
| **(1) FSC -(Intercept)** | -0.82824 | 0.05401 | -15.33633 | 1.77971 | 0.02260 | 78.75725 |
| **WL** | -0.17202 | 0.02407 | -7.14633 | 0.14650 | 0.01064 | 13.76839 |
| **MASC** | 0.22769 | 0.09354 | 2.43415 | 0.08772 | 0.03914 | 2.24108 |
| **WL:MASC** | 0.06335 | 0.04169 | 1.51957 | 0.05944 | 0.01843 | 3.22521 |
| **(2) MASC -(Intercept)** | -0.60055 | 0.07637 | -7.86321 | 1.86742 | 0.03196 | 58.43454 |
| **WL** | -0.10866 | 0.03404 | -3.19214 | 0.20594 | 0.01505 | 13.68579 |

LMMs were fitted separately to the GMM-estimated mean (left panel) and SD (right panel) of the largest mixture component in individual within-word landing position distributions split by word length. In both LMMs, the fixed structure included Data Set (2 levels: FSC and MASC), Word Length (WL; 4-9 letters; reference (mean) value: 6.5 letters), and their interaction as predictors. The random structure included both a random intercept and a random effect of WL by subject. Estimates and standard errors are expressed in letters. Panel 1: Fixed-effects for the main LMMs, with Data-Set reference level set to FSC. Panel 2: Data-Set reference level set to MASC.

**The Launch Site effect.** It is well-established that the PVL effect results from a more basic phenomenon, referred to as the launch-site effect: the typical, normal, distribution of initial landing positions in words of a given length is in fact a composite distribution that results from the summation of many normal distributions, each contingent on how far from the words' beginning the eyes come from[6]. When saccades are launched from a position near to the words' beginning, the landing-site distribution tends to peak to the right of the words' center, but as saccades are launched from further away, landing positions progressively shift towards the words' beginning, resulting in a linear relationship between launch-site distance and mean landing site. The slope of this relationship was initially reported to be in the order of 0.5 regardless of word length, but this was not later confirmed, the slope varying with word length[7], as well as the peripheral visual configuration[8-10].

In line with previous findings, the distributions of within-word landing positions in 4- to 9-letter words, contingent on saccades' launch-site distance to the space in front of the words, were most often best fitted with a single mixture component in both FSC readers and MASC (Fig. 2e).

This is summarized in Supplementary Table 9, where the mean proportion of two mixture components was most often zero, and in any case never greater than 0.02.

**Supplementary Table 9 | GMM-estimated shape of within-word landing-site distributions by word length and launch site in MASC and FSC readers.**

| LS EFFECT: | 4-lw | | 5-lw | | 6-lw | | 7-lw | | 8-lw | | 9-lw | |
|---|---|---|---|---|---|---|---|---|---|---|---|---|
| 2 MODES | Far | Near | Far | Near | Far | Near | Far | Near | Far | Near | Far | Near |
| FSC | 0.00 | 0.00 | 0.00 | 0.01 | 0.01 | 0.00 | 0.00 | 0.01 | 0.00 | 0.00 | 0.02 | 0.00 |
| MASC | 0.00 | 0.00 | 0.00 | 0.00 | 0.02 | 0.00 | 0.00 | 0.00 | 0.00 | 0.02 | 0.02 | 0.00 |

Mean proportion of two mixture components (i.e., the maximal number of detected modes), as estimated after fitting GMMs to individual landing-position distributions partitioned by word length and saccades' launch-site distance to the space in front of the words (binned in two-letter intervals) separately for FSC readers and MASC. To make the tables more digest, means were computed respectively across near and far saccadic launch-site distances separately for 4- to 9-letter words (i.e., below and above the median of launch sites, that is -5 letters for 4- to 8-letter words and -3 letters for 9-letter words). The mean of the largest *k* estimates was greater than, or equal to, 0.988 in all data sets and conditions. These analyses relied on a total of 19084 and 10050 cases across subjects in FSC and MASC respectively.

Moreover, for both MASC and FSC readers, the distributions of within-word landing positions progressively shifted from the end to the beginning of words with increasing saccadic launch-site distance (Fig. 2e-f). As shown in the left panels of Supplementary Table 10, the estimated mean landing position in 6-letter words, when saccades were launched from about 4.7 letters from the space in front of the words, was slightly to the left of the words' center (estimates: -0.67987 and -0.77251 for FSC and MASC respectively; Panels 1-2) and non-significantly different between the two data sets ($t$ = -0.73378; Panel 1). This leftward bias increased with word length (Panels 1 and 2), at the same rate for FSC readers and MASC, the interaction between word length and MASC being again non-significant ($t$ = -0.35430; Panel 1; see also Supplementary Table 8). Moreover, in both data sets, this bias increased significantly with increasing launch-site distance (estimates: 0.36196 and 0.60744 respectively; Panels 1-2), and even more so as words were longer, as suggested by the significant interaction between word length and launch site (estimates: 0.04761 and 0.10975 respectively; Panels 1-2). The slope of the linear relationship between launch-site distance and mean landing site yet remained significantly steeper in MASC compared to FSC readers (estimate: 0.24547; Panel 1), and this increased more greatly with increasing word length, as suggested by the significant three-way interaction between word length, launch-site distance, and MASC (estimate: 0.06214; Panel 1).

**Supplementary Table 10 | Fixed effects of LMMs for the mean and SD of within-word landing sites by word length and launch site in MASC and FSC readers.**

| LS EFFECT: MEAN / SD | Estimate | Std. Error | *t* value | Estimate | Std. Error | *t* value |
|---|---|---|---|---|---|---|
| **(1) FSC -(Intercept)** | -0.67987 | 0.07300 | -9.31345 | 1.31175 | 0.01880 | 69.79090 |
| **WL** | -0.23796 | 0.01689 | -14.09053 | 0.09344 | 0.00730 | 12.79650 |
| **LS** | 0.36196 | 0.01471 | 24.60874 | -0.01755 | 0.00471 | -3.72691 |
| **MASC** | -0.09264 | 0.12625 | -0.73378 | -0.19243 | 0.03240 | -5.94013 |
| **WL:LS** | 0.04761 | 0.00331 | 14.39143 | 0.00489 | 0.00207 | 2.36024 |
| **WL:MASC** | -0.01033 | 0.02916 | -0.35430 | -0.04219 | 0.01256 | -3.35947 |
| **LS:MASC** | 0.24547 | 0.02535 | 9.68150 | 0.03403 | 0.00809 | 4.20497 |
| **WL:LS:MASC** | 0.06214 | 0.00562 | 11.05461 | 0.00393 | 0.00352 | 1.11788 |
| **(2) MASC -(Intercept)** | -0.77251 | 0.10301 | -7.49930 | 1.11932 | 0.02639 | 42.42155 |
| **WL** | -0.24829 | 0.02377 | -10.44432 | 0.05125 | 0.01022 | 5.01581 |
| **LS** | 0.60744 | 0.02065 | 29.41254 | 0.01648 | 0.00658 | 2.50396 |
| **WL:LS** | 0.10975 | 0.00454 | 24.15028 | 0.00883 | 0.00284 | 3.10542 |

LMMs were fitted separately to the GMM-estimated mean (left panel) and SD (right panel) of the largest mixture component in individual within-word landing position distributions split by word length and saccades' launch-site distance to the space in front of the words (binned in two-letter intervals). In both LMMs, the fixed structure included Data Set (2 levels: FSC and MASC), Word Length (WL; 4-9 letters; reference (mean) value: 6.34 letters), Launch Site (LS; in two-letter bins and ranging between -9 and -1 letters; reference (mean) value: -4.69 letters), and all interactions as predictors. The random structure included a random intercept as well as random effects of WL and LS by subject, but without the correlation between random effects for the SD. Estimates and standard errors are expressed in letters. Panel 1: Fixed-effects for the main LMMs, with Data-Set reference level set to FSC. Panel 2: Data-Set reference level set to MASC.

As shown in the right panels of Supplementary Table 10, the variability in within-word landing positions was smaller for MASC compared to FSC readers (estimate: -0.19243; Panel 1). Moreover, it tended to mildly decrease with increasing launch-site distance (estimate: 0.01648; Panel 2), while the reverse was true in FSC readers (estimate: -0.01755; Panel 1)[6]. Still, as in the above analyses across launch sites (Supplementary Table 8, right panels), the variability in within-word landing positions significantly increased with increasing word length for both FSC readers and MASC (estimates: 0.09344 and 0.05125; Panels 1-2), but this time at a slightly slower rate for

MASC compared to FSC readers, as suggested by the significant interaction between word length and MASC (estimate: -0.04219; Panel 1).

**All landing positions regardless of word boundaries**

**Methodological Note.** Due to the long-standing assumption that eye-movement behavior during reading is determined in a word-based manner (Extended Data Table 1), saccades' landing positions are traditionally analyzed within word boundaries. This means that only the saccades that land on a word of a given length, as well as on the space in front of the word in spaced alphabetic languages, are considered for analysis, and hence that all other saccades, that either fall short or land beyond the word's end, are excluded. As a result, landing-site distributions, notably those that peak near the word boundaries (as for very-near and very-far saccades launch sites relative to the beginning of words), are truncated (Fig. 2e). The problem is that this biases estimation of the shape and mode (mean) of the distributions, but also leads to underestimate the relationship between launch-site distance and mean landing site[7,10], and even more so as the distributions are more widely spread, as in the case of long words and for FSC readers in comparison with MASC (Supplementary Table 10). The full distributions of saccades' landing positions, regardless of whether they fall within word boundaries, therefore enable fairer comparison of the launch-site effect between conditions, and here between MASC and FSC readers, compared to within-word landing-position distributions. As further developed below, when all saccades' landing positions were considered for analysis, MASC's launch-site effect differed only mildly from that observed in FSC readers, and mainly for long words.

**The Launch Site effect revisited.** In the present analyses, saccades were launched from a given word (N), and their landing positions were measured relative to the center of the next word on the line (N+1), but regardless of whether they landed on Word N, Word N+1 or one of the following words. The resulting overall distributions of landing sites were fitted with GMMs, separately for each subject, and a range of word lengths and saccadic launch-site distances. Since these analyses, in opposition with within-word landing-positions analyses, made sense regardless of word length, tested words ranged between 1 and 10 letters.

As shown in Supplementary Table 11, the proportion of two-mixture components in overall landing-position distributions remained very low; in FSC readers, it tended to be slightly greater than for within-word landing-position distributions (Supplementary Table 9) and compared to MASC, but the largest component still accounted on average for more than 93% of the data. This is already an argument against models of eye-movement control during reading that rely on the assumption that saccades are programmed towards the center of peripherally selected target words (Extended Data Table 1). Indeed, these models would predict the distributions to be bimodal, or even trimodal, with each mode aligned with the center of successive words in the sentence (Word N, N+1 and possibly also N+2)[7].

**Supplementary Table 11 | GMM-estimated shape of overall landing-site distributions by word length and launch site in MASC and FSC readers.**

| OVERALL LS EFFECT: 2 MODES | 1-lw | 2-lw | 3-lw | 4-lw | 5-lw | 6-lw | 7-lw | 8-lw | 9-lw |
|---|---|---|---|---|---|---|---|---|---|
| **Far -FSC** | 0.02 | 0.11 | 0.18 | 0.05 | 0.00 | 0.02 | 0.05 | 0.01 | 0.05 |
| **Far -MASC** | 0.00 | 0.00 | 0.00 | 0.00 | 0.00 | 0.00 | 0.00 | 0.00 | 0.00 |
| **Near -FSC** | 0.00 | 0.04 | 0.00 | 0.01 | 0.00 | 0.02 | 0.05 | 0.00 | 0.00 |
| **Near -MASC** | 0.00 | 0.00 | 0.00 | 0.00 | 0.00 | 0.00 | 0.00 | 0.00 | 0.00 |

Mean proportion of two mixture components (i.e., the maximal number of detected modes), as estimated after fitting GMMs to individual distributions partitioned by word length and saccades' launch-site distance to the space in front of the words (binned in two-letter intervals) separately for FSC readers and MASC. To make the tables more digest, means were computed respectively across near and far saccadic launch-site distances separately for 1- to 9-letter words (i.e., below and above the median of launch sites, that is -5 letters in 2- to 4-letter words, and -3 letters in 1- and 5- to 9-letter words). Since the mean of the largest *k* estimates was greater than, or equal to, 0.937 in all data sets and conditions, these are not reported in the table. These analyses relied on a total of 35654 and 19444 cases across subjects in FSC and MASC respectively.

The left panels of Supplementary Table 12 present the fixed effects of LMMs fitted to the estimated mean of the largest mixture component in overall landing-position distributions. This shows that FSC readers tended to fixate to the right of the center of short words (estimate: 1.43201, for Words N+1 of 4.55 letters on average and launch sites of -3.64 letters from the space in front of the words; Panel 1), but progressively closer to the words' beginning as word length increased and saccades were launched from further away (estimates: -0.61676 and 0.82160; Panel 1). MASC's saccades also overshot the center of short words, by about the same extent as for FSC readers ($t$ = -1.47906), and they landed closer to the words' beginning as word length increased (estimate: -0.33243; Panel 2). The effect of word length was smaller than for FSC readers, as suggested by the significant interaction between word length and MASC (estimate: 0.28433; Panel 1). However, landing positions shifted towards the words' beginning with increasing launch-site distance (estimates: 0.82160 and 0.85225 respectively; Panels 1-2) at about the same rate as for FSC readers when words were about 4-letters long on average ($t$ = 0.69398 for the interaction between MASC and launch site; Fig. 2g). The only difference was that MASC's launch-site effect slightly increased with increasing word length (estimate: 0.03537; Panel 2), while it slightly decreased with word length in FSC readers (estimate: -0.01347; Panel 1), leading to a significant interaction between word length, launch site and MASC (estimate: 0.04884; Panel 1). However, the slope difference between MASC and FSC readers for the relationship between launch-site distance and mean landing site in 6-letter words was only of about 0.10, thus more than half that observed when fitting LMMs to mean within-word landing positions (Supplementary Table 10, left panels). Thus, the above-reported disparity between MASC and FSC readers for the launch-site effect came for a great part from using truncated, within-word landing-position, distributions. The small remaining differences are attributable to top-down, language-related, processes, which mildly modulate saccade amplitude[7], exclusively in FSC readers (Extended Data Fig. 2).

**Supplementary Table 12 | Fixed effects of LMMs for the mean and SD of all landing sites by word length and launch site in MASC and FSC readers.**

| OVERALL-LS EFFECT: MEAN / SD | Estimate | Std. Error | t value | Estimate | Std. Error | t value |
|---|---|---|---|---|---|---|
| **(1) FSC -(Intercept)** | 1.43201 | 0.17118 | 8.36562 | 2.33144 | 0.05836 | 39.95132 |
| **WL** | -0.61676 | 0.01811 | -34.05536 | -0.07778 | 0.00897 | -8.67021 |
| **LS** | 0.82160 | 0.02459 | 33.41185 | -0.04187 | 0.00985 | -4.25099 |
| **MASC** | -0.44004 | 0.29752 | -1.47906 | -0.65647 | 0.10221 | -6.42268 |
| **WL:LS** | -0.01347 | 0.00578 | -2.33135 | 0.01240 | 0.00347 | 3.57042 |
| **WL:MASC** | 0.28433 | 0.03375 | 8.42330 | -0.00106 | 0.01734 | -0.06132 |
| **LS:MASC** | 0.03065 | 0.04416 | 0.69398 | 0.11762 | 0.01854 | 6.34418 |
| **WL:LS:MASC** | 0.04884 | 0.01158 | 4.21889 | -0.00887 | 0.00699 | -1.26830 |
| **(2) MASC -(Intercept)** | 0.99197 | 0.24334 | 4.07650 | 1.67497 | 0.08391 | 19.96033 |
| **WL** | -0.33243 | 0.02849 | -11.67041 | -0.07885 | 0.01484 | -5.31179 |
| **LS** | 0.85225 | 0.03668 | 23.23363 | 0.07575 | 0.01571 | 4.82269 |
| **WL:LS** | 0.03537 | 0.01003 | 3.52584 | 0.00353 | 0.00607 | 0.58172 |

LMMs were fitted separately to the GMM-estimated mean (left panel) and SD (right panel) of the largest mixture component in individual overall landing-position distributions split by word length and saccades' launch-site distance to the space in front of the words (binned in two-letter intervals). In both LMMs, the fixed structure included Data Set (2 levels: FSC and MASC), Word Length (WL; 1-10 letters; reference, mean, value: 4.55 letters), Launch Site (LS; in two-letter bins and ranging between -9 and -1 letters; reference (mean) value: -3.64 letters), and all interactions as predictors. The random structure included a random intercept as well as random effects of WL and LS by subject, but without the correlation between random effects for SD. Estimates and standard errors are expressed in letters. Panel 1: Fixed-effects for the main LMMs, with Data-Set reference level set to FSC. Panel 2: Data-Set reference level set to MASC.

The above-reported analyses revealed that within-word landing positions were less widely spread for MASC in comparison with FSC readers (Supplementary Table 10). As shown in the right panels of Supplementary Table 12, the same was true for the full distributions of landing positions, although the difference was much larger (estimate: -0.65647; Panel 1) and the distributions were about twice as wide (estimate: 2.33144 for FSC readers when word length and launch site were at their reference (mean) value; Panel 1). The variability decreased, and no longer increased, with increasing word length, at about the same rate in both data sets (estimates: -0.07778 and -0.07885 respectively; Panels 1-2), the interaction between MASC and word length being non-significant ($t$ = -0.06132). However, the SD still slightly increased with increasing launch-site distance for FSC readers, but not MASC, which showed the opposite effect (estimates: -0.04187 and 0.07575 respectively; Panels 1-2); the interaction between MASC and launch-site distance was significant (estimate: 0.11762; Panel 1).

**Within-word refixation behavior: The OVP effect.**

The Refixation-OVP effect shows that the likelihood of making two (or more) consecutive fixations on a word (or refixation) is mainly a function of where in the word the eyes initially land, as well as the word's length[11-14]. In languages read from left to right, it is minimal towards the center of short words, and slightly to the left of the center of long words, and it increases gradually as words are longer and as the initial eye fixation deviates from this optimal location, thus yielding typically U-shaped curves. FSC readers replicated this phenomenon (Fig. 2h). However, MASC only showed half of the effect, that is only the left wing associated with initial landing positions in the first, beginning, part of the words, initiating nearly no refixations from the end part of words. To statistically compare the refixation behavior of MASC and FSC readers, GLMMs were thus fitted to only a subset of the data, those associated with initial fixation positions in the beginning part of words. The resulting fixed effects are displayed in Supplementary Table 13.

FSC readers and MASC refixated words in about 13% and 11% of the cases respectively (logit: -1.85545 and -2.03858; Panels 1-2), when words were about 6 letters long and the initial fixation was about 2 letters to the left of the words' center; these two refixation rates were not significantly different ($p$ = 0.42950; Panel 1). Most importantly, for both FSC readers and MASC, the likelihood of within-word refixations increased with increasing word length (logit: 0.33450 and 0.29206 respectively; Panels 1-2), and as the initial fixation more greatly deviated from the words' center (logit: -0.64172 and -0.67665 respectively; Panels 1-2); the latter effect slightly differed depending on word length, but only for FSC readers (logit: 0.02733; Panel 1). As suggested by the non-significant interaction between MASC and initial fixation position ($p$ = 0.65025), there was no slope difference between both data sets in the left wing of the OVP effect for 6-letter words (the reference (mean) value). However, as words were longer, the slope tended to be slightly steeper for MASC in comparison with FSC readers, as suggested by the marginally significant interaction between word length, initial fixation position and MASC (logit: -0.03854, *p* = 0.08426).

**Supplementary Table 13 | Fixed effects of GLMMs for the probability of within-word refixations by initial landing position and word length in MASC and FSC readers**.

| Refixation-OVP EFFECT | Estimate | Std. Error | z value | Pr(>\|z\|) | Proportion |
|---|---|---|---|---|---|
| **(1) FSC -(Intercept)** | -1.85545 | 0.13572 | -13.67127 | 0.00000 | 0.13523 |
| **WL** | 0.33450 | 0.02166 | 15.44534 | 0.00000 | |
| **ILP** | -0.64172 | 0.04453 | -14.40999 | 0.00000 | |
| **MASC** | -0.18282 | 0.23140 | -0.79004 | 0.42950 | 0.11524 |
| **WL:ILP** | 0.02733 | 0.01279 | 2.13687 | 0.03261 | |
| **WL:MASC** | -0.04248 | 0.03483 | -1.21976 | 0.22255 | |
| **ILP:MASC** | -0.03462 | 0.07635 | -0.45342 | 0.65025 | |
| **WL:ILP:MASC** | -0.03854 | 0.02232 | -1.72650 | 0.08426 | |
| **(2) MASC -(Intercept)** | -2.03858 | 0.18955 | -10.75481 | 0.00000 | |
| **WL** | 0.29206 | 0.02829 | 10.32357 | 0.00000 | |
| **ILP** | -0.67665 | 0.06215 | -10.88667 | 0.00000 | |
| **WL:ILP** | -0.01113 | 0.01843 | -0.60379 | 0.54599 | |

GLMMs were fitted to a binary variable indicating whether a given word was refixated. The fixed structure included Data Set (2 levels: FSC and MASC), Word Length (WL; 4-9 letters; reference (mean) value: 6.41), Initial Landing Position (ILP; ranging between -5.5 and 0 letters relative to the center of words, thus excluding ILPs in the second halves of words, which yielded near-zero refixation in MASC; reference (mean) value: -2.04 letters), and all interactions. The random structure included a random intercept by subject and by sentence pair, as well as random effects of WL and ILP by subject. Estimates and standard errors are expressed in logit unit; intercept estimates are converted into probabilities in the rightmost column (in Panel 1), thus giving an estimate of the refixation likelihood in each of the data sets, when WL and ILP were at their reference value. Panel 1: Fixed-effects for the main GLMM model, with Data-Set reference level set to FSC. Panel 2: Data-Set reference level set to MASC. These analyses relied on a total of 16985 and 8579 cases across subjects in FSC and MASC respectively.

MASC's slight departure from readers' refixation behavior in the case of longer words again very likely came from the fact that it was immune to mild language-related modulations (Extended Data Fig. 3). The same was likely true for the right wing of the OVP effect, that was absent in MASC. However, note that some regressive refixations were generated from the end part of words in MASC_ISR_C, that is MASC with ISR applied only to the current fixation, despite this model, as MASC, being completely illiterate (Extended Data Fig. 6f, Supplementary Table

28). This suggests that refixations from the end part of words, as well as regressions (see below), do not entirely reflect ongoing word-identification processes.

### Regression behavior

**The variables that affect the likelihood of regressions.** In FSC readers, the likelihood of a regressive saccade mildly increased with increasing fixation number, but also as the eye progressed on the line of text, though more drastically towards the last fourth of the sentence (Extended Data Fig. 4a-b). MASC showed very roughly the same pattern, but the increase with increasing fixation number was more drastic and non-linear. Moreover, the relationship between regression probability and fixation position on the line tended to be quadratic, with more regressions towards the first and last fourth of the sentences and nearly no regression in between. Given this, it was unreasonable to fit GLMMs to the data.

Another variable that affected the probability FSC readers made regressions was the length of the prior saccade, when this moved the eye forward: As the prior saccade was longer, the likelihood of a regressive saccade became higher, and even more so when the prior saccade skipped over a word, in line with previous reports[15-17] (Extended Data Fig. 4c-d). MASC also made more regressions following very long prior forward saccades that skipped over a word, but it made nearly no regressions for a range of shorter prior saccade lengths. Given these floor effects, GLMMs could not be reasonably fitted to the data.

**The Regression-PVL effect.** Only a few studies reported the distributions of within-word landing positions of regressive saccades[17-18]. However, they consistently indicated that these distributions are normal in shape, and that they tend to invariably peak towards the centers of words. In line with these findings, the majority of the distributions in FSC readers were best fitted with a single mixture component, the proportion of two mixture components being less than or equal to 0.08, except in 6-letter words where this proportion was of about 0.14 (Supplementary Table 14, Extended Data Fig. 4e). MASC's distributions were also best fitted with a single mode, but the proportion of two mixture components remained higher in 5- to 8-letter words (0.10-0.35). The largest component in MASC's distributions still accounted on average for 87%-96% of the data (against 94%-99% in FSC readers).

**Supplementary Table 14 | GMM-estimated shape of the distributions of within-word landing sites following regressions in MASC and FSC readers.**

| RG - PVL EFFECT | 4-lw | | 5-lw | | 6-lw | | 7-lw | | 8-lw | | 9-lw | |
|---|---|---|---|---|---|---|---|---|---|---|---|---|
| **2 Modes/ Largest *K*** | **2M** | $K_k$ | **2M** | $K_k$ | **2M** | $K_k$ | **2M** | $K_k$ | **2M** | $K_k$ | **2M** | $K_k$ |
| **FSC** | 0.00 | 1,00 | 0.03 | 0,99 | 0.14 | 0,94 | 0.04 | 0,99 | 0.04 | 0,98 | 0.08 | 0,97 |
| **MASC** | 0.00 | 1,00 | 0.10 | 0,96 | 0.20 | 0,92 | 0.35 | 0,87 | 0.20 | 0,92 | 0.05 | 0,98 |

Mean proportion of two mixture components (i.e., the maximal number of detected modes), and mean proportion of cases accounted for by the largest mixture component (mean of largest *k* values; "$K_k$"), as estimated after fitting GMMs to individual distributions of regressive within-word landing positions in 4- to 9-letter words separately for FSC readers and MASC. These analyses relied on a total of 4303 and 7964 cases across subjects in FSC and MASC respectively.

As shown in the left panels of Supplementary Table 15, FSC readers fixated on average slightly to the right of the center of 6-letter words following regressions (intercept: 0.22560; Panel 1), and even more so as word length increased; note though, that the increase was only of about 1/10 letter per one-letter increment of word length (estimate: 0.09316; Panel 1). MASC did not exhibit this rightward fixation bias, making regressive saccades that landed exactly at the center of 6-letter words (intercept: -0.01272, $t$ = -0.14099; Panel 2). Still, as words were longer, regressions landed further towards the end of words (estimate: 0.13816; Panel 2), just as for FSC readers, as suggested by the non-significant interaction between word length and MASC ($t$ = 0.84667). Moreover, as shown in the right panels of Supplementary Table 15, the variability in within-word landing positions increased with word length in both FSC readers and MASC (estimates: 0.19946 and 0.20528; Panels 1-2), and at the same rate in both data sets ($t$ = 0.20929; Panel 1). The only difference was that the variability was overall greater for MASC compared to FSC readers (estimate: 0.22530; Panel 1).

**Supplementary Table 15 | Fixed effects of LMMs for the mean and SD of within-word landing sites following regressions in MASC and FSC readers.**

| RG - PVL EFFECT: MEAN / SD | Estimate | Std. Error | t value | Estimate | Std. Error | t value |
|---|---|---|---|---|---|---|
| **(1) FSC -(Intercept)** | 0.22560 | 0.07640 | 2.95285 | 1.57639 | 0.03206 | 49.16455 |
| **WL** | 0.09316 | 0.03688 | 2.52572 | 0.19946 | 0.01915 | 10.41334 |
| **MASC** | -0.23831 | 0.11820 | -2.01612 | 0.22530 | 0.04864 | 4.63228 |
| **WL:MASC** | 0.04500 | 0.05315 | 0.84667 | 0.00582 | 0.02779 | 0.20929 |
| **(2) MASC -(Intercept)** | -0.01272 | 0.09020 | -0.14099 | 1.80169 | 0.03657 | 49.26317 |
| **WL** | 0.13816 | 0.03828 | 3.60963 | 0.20528 | 0.02013 | 10.19584 |

LMMs were fitted separately to the GMM-estimated mean (left panel) and SD (right panel) of the largest mixture component in individual regressive within-word landing position distributions split by word length. In both LMMs, the fixed structure included Data Set (2 levels: FSC and MASC), Word Length (WL; 4-9 letters; reference (mean) value: 6.27 letters) and their interaction as predictors. The random structure included both a random intercept and a random effect of WL by subject. Estimates and standard errors are expressed in letters. Panel 1: Fixed-effects for the main LMM, with Data-Set reference level set to FSC. Panel 2: Data-Set reference level set to MASC.

### Supplementary Methods 2. Comparison models vs. MASC and FSC readers' behavior

In the present analyses, the oculomotor behavior predicted by alternative MASC models (MASC_noRT, MASC_VISUAL, MASC_MOTOR, MASC_IOR_C, and MASC_IOR_PC), VS models (VS_noRT, VS_RT, and VS_RT_GA), and CSL was compared to the behavior of MASC and FSC readers. Due to differences in numbers of fixations per sentence across data sets, and the fact notably that some models generated less fixations than FSC readers, data sets could hardly be matched for numbers of fixations. Thus, in the present comparisons, data sets were matched by considering only first-pass fixations/saccades over sentences (see Methods). This also enabled fairer comparison with CSL, which generated no regressive saccades.

#### Saccade length distributions

**Typical bimodal distributions.** We saw above that MASC and FSC readers exhibited bimodal distributions of saccade lengths, with one mode corresponding to regressive saccades and the other mode corresponding to forward saccades[1] (Supplementary Table 1). When only first-pass eye-movement data were considered for analysis, a similar trend was observed in these two data sets, even though the proportion of regressions was smaller, notably for MASC (Extended Data Fig. 5a, left vs. right panels), which generated regressions mainly from the end of the sentences (Extended Data Fig. 4b). As shown in Supplementary Table 16, individual distributions for FSC readers, MASC, and alternative MASC models (Extended Data Fig. 5g, 6a, 7a), were, for the most, best fitted with two (one negative and one positive) mixture components. Moreover, when three mixture components were detected, as for FSC readers, there were two positive peaks, and the largest accounted on average for more 97% of the data.

In contrast, the distributions predicted by VS models were best fitted with either three (one negative and two positive) or four (two negative and two positive) mixture components, depending on whether the model included RT (Fig. 3g). Furthermore, the largest mixture component associated with forward saccades accounted on average for only 58-67% of the data, while the largest mixture component associated with regressions in VS_noRT accounted on average for only 65% of the data. CSL generated no regressions, and its saccade-length distributions were all best fitted with a single positive mixture component (Fig. 3a). Thus, the models that best predicted the typical shape of saccade-length distributions in FSC readers were MASC models. As further developed below, MASC was also part of the models that best accounted for the proportion of regressive saccades, as well as the metrics of forward saccades (mean and SD).

**Supplementary Table 16 | GMM-estimated shape of the distributions of saccade lengths in (comparison) models and FSC readers.**

| | ALL SACCADES | | | | REGRESSIONS | | PROGRESSIONS | |
|---|---|---|---|---|---|---|---|---|
| **SL: MODES (1)** | **1** | **2** | **3** | **4** | **2** | **Ratio** | **2** | **Ratio** |
| **FSC** | 0.000 | 0.900 | 0.100 | 0.000 | 0.000 | 1.000 | 0.010 | 0.976 |
| **MASC** | 0.000 | 1.000 | 0.000 | 0.000 | 0.000 | 1.000 | 0.000 | 1.000 |
| **MASC_noRT** | 0.000 | 1.000 | 0.000 | 0.000 | 0.000 | 1.000 | 0.000 | 1.000 |
| **MASC_VISUAL** | 0.000 | 1.000 | 0.000 | 0.000 | 0.000 | 1.000 | 0.000 | 1.000 |
| **MASC_MOTOR** | 0.000 | 1.000 | 0.000 | 0.000 | 0.000 | 1.000 | 0.000 | 1.000 |
| **MASC_ISR_C** | 0.000 | 1.000 | 0.000 | 0.000 | 0.000 | 1.000 | 0.000 | 1.000 |
| **MASC_ISR_1PC** | 0.000 | 1.000 | 0.000 | 0.000 | 0.000 | 1.000 | 0.000 | 1.000 |
| **VS_noRT** | 0.000 | 0.000 | 0.000 | 1.000 | 1.000 | 0.651 | 1.000 | 0.622 |
| **VS_RT** | 0.000 | 0.000 | 1.000 | 0.000 | 0.000 | 1.000 | 1.000 | 0.669 |
| **VS_RT_GA** | 0.000 | 0.000 | 1.000 | 0.000 | 0.000 | 1.000 | 1.000 | 0.577 |
| **CSL** | 1.000 | 0.000 | 0.000 | 0.000 | 0.000 | 1.000 | 0.000 | 1.000 |

GMM-estimated proportion of 1-4 mixture components in individual saccade-length distributions (left panel), and proportion of two mixture components (vs. one single component), as well as mean ratios between the largest k estimate and the sum of k estimates, separately for regressive and progressive saccades (right panel), for FSC, MASC, and all nine comparison models. These analyses relied on a total of 56352, 31888, 33073, 28835, 32706, 32075, 36777, 13639, 28773, 33013, 23185 cases across subjects in FSC, MASC, MASC_noRT, MASC_VISUAL, MASC_MOTOR, MASC_ISR_C, MASC_ISR_1PC, VS_noRT, VS_RT, VS_RT_GA, and CSL respectively.

**Proportion of regressive saccades.** Panel 1 of Supplementary Table 17 shows the regression coefficients of GLMs fitted to the GMM-estimated regression rate, when Data Set was at its reference FSC level. The intercept logit estimate, of about -1.77235 indicated that regressions occurred in about 14% of the cases in FSC readers. The regression rate was significantly lower for

MASC (9%), as well as MASC_VISUAL and MASC_MOTOR (4% and 3% respectively), while being non-significantly different for MASC_noRT ($p$ = 0.24875). However, when ISR was applied only to the current fixation (MASC_ISR_C), or to the current and the immediately prior fixations (MASC_ISR_1PC), the proportion of regressions was inflated and significantly greater than for FSC readers (28% and 20% respectively). Still, MASC_ISR_1PC overestimated the proportion of regressions to about the same extent as MASC underestimated it.

**Supplementary Table 17 | GLM regression coefficients for the proportion of regressions in (comparison) models and FSC readers.**

| REGRESSION RATE | Estimate | Std. Error | z value | Pr(>\|z\|) | Proportion |
|---|---|---|---|---|---|
| **(1) FSC -(Intercept)** | -1.77235 | 0.04487 | -39.49698 | 0.00000 | 0.14525 |
| **MASC** | -0.58479 | 0.09133 | -6.40296 | 0.00000 | 0.08650 |
| **MASC_noRT** | 0.08785 | 0.07617 | 1.15339 | 0.24875 | 0.15650 |
| **MASC_VISUAL** | -1.31817 | 0.11845 | -11.12840 | 0.00000 | 0.04350 |
| **MASC_MOTOR** | -1.72107 | 0.13956 | -12.33182 | 0.00000 | 0.02950 |
| **MASC_ISR_C** | 0.84272 | 0.06692 | 12.59378 | 0.00000 | 0.28300 |
| **MASC_ISR_1PC** | 0.38606 | 0.07168 | 5.38556 | 0.00000 | 0.20000 |
| **VS_noRT** | 1.31873 | 0.06417 | 20.54941 | 0.00000 | 0.38850 |
| **VS_RT** | 0.46840 | 0.07065 | 6.62994 | 0.00000 | 0.21350 |
| **VS_RT_GA** | -0.89381 | 0.10120 | -8.83242 | 0.00000 | 0.06500 |
| **(2) MASC –(Intercept)** | -2.35714 | 0.07955 | -29.63211 | 0.00000 | |
| **MASC_noRT** | 0.67264 | 0.10058 | 6.68789 | 0.00000 | |
| **MASC_VISUAL** | -0.73338 | 0.13544 | -5.41470 | 0.00000 | |
| **MASC_MOTOR** | -1.13628 | 0.15425 | -7.36666 | 0.00000 | |
| **MASC_ISR_C** | 1.42751 | 0.09376 | 15.22438 | 0.00000 | |
| **MASC_ISR_1PC** | 0.97084 | 0.09722 | 9.98555 | 0.00000 | |
| **VS_noRT** | 1.90352 | 0.09183 | 20.72919 | 0.00000 | |
| **VS_RT** | 1.05318 | 0.09646 | 10.91787 | 0.00000 | |
| **VS_RT_GA** | -0.30902 | 0.12064 | -2.56144 | 0.01042 | |

GLMs were fitted to the GMM-estimated regression rate, with Data Set (10 levels: FSC, MASC, MASC_noRT, MASC_VISUAL, MASC_MOTOR, MASC_ISR_C, MASC_ISR_1PC, VS_noRT, VS_RT_GA, thus excluding CSL, that did not generate regressions) as a categorical predictor. Estimates and standard errors are expressed in logit unit. Corresponding regression rates in the different data sets are given in the rightmost column. Panel 1: Regression coefficients for the main GLM, with Data-Set reference level set to FSC. Panel 2: Data-Set reference level set to MASC. In Panel 2, the regression coefficients that were redundant with those reported in Panel 1 were dropped.

VS models more greatly misestimated the proportion of regressions than MASC: VS_RT, and even more so VS_noRT, made too many regressions (21% and 39% respectively) and VS_RT_GA made less regressions (6%) than either MASC or FSC readers. Thus, between all models, the ones that best predicted FSC readers' regression rate, were MASC_noRT, followed by MASC_ISR_1PC, MASC, and then VS_RT_GA; note that the first two models, as well as other comparison models, differed significantly from MASC (Panel 2). Thus, for this dependent variable, MASC was among the most performing models, though not the best.

**Length of regressive saccades.** As reported in Supplementary Table 18A, the estimated mean length of regressive saccades was of about -4.08 letters in FSC readers. This was significantly larger in MASC and comparison models, except for MASC_MOTOR, MASC_ISR_C, and VS_RT_GA, which did not differ significantly from FSC (*p* > = 0.06899; Panel 1). MASC still overestimated the mean length of regressions by only about 1.34 letters (Panel 1), and by about the same extent as these three models (p >= 0.08767; Panel 2). Moreover, it outperformed all other models (Panel 2), which overestimated the mean regression length by 3.03-8.60 letters (Panel 1). Additionally, as shown in Supplementary Table 18B, although MASC made regressive saccades that were more variable in length compared to FSC readers (estimate: 2.28149; Panel 1), it still did better than MASC_MOTOR (Panel 2), which overestimated the SD by about 4.67 letters (Panel 1), as well as MASC_ISR_C and VS_RT_GA (Panel 2), which underestimated it by about 2.79-3.22 letters (Panel 1). MASC however did worse than MASC_noRT and MASC_ISR_1PC (Panel 2) which did not differ significantly from FSC (*p* >= 0.12280; Panel 1). It also did worse than MASC_VISUAL (Panel 2), which overestimated the SD by only about 0.8 letter (Panel 1), as well as VS_noRT and VS_RT (Panel 2), which underestimated the SD by about 1 letter (Panel 1). Recall though that the largest negative mode in VS_noRT accounted on average for only 65% of the data (Supplementary Table 16).

**Supplementary Table 18 | LM regression coefficients for the mean and SD of the length of regressive saccades in (comparison) models and FSC readers.**

| A- REGRESSIONS: MEAN | Estimate | Std. Error | t value | Pr(>\|t\|) |
|---|---|---|---|---|
| (1) FSC -(Intercept) | -4.07981 | 0.33976 | -12.00798 | 0.00000 |
| MASC | -1.34539 | 0.58848 | -2.28622 | 0.02324 |
| MASC_noRT | -4.50975 | 0.58848 | -7.66342 | 0.00000 |
| MASC_VISUAL | -4.48351 | 0.58848 | -7.61882 | 0.00000 |
| MASC_MOTOR | -0.85659 | 0.58848 | -1.45560 | 0.14700 |
| MASC_ISR_C | -1.07566 | 0.58848 | -1.82787 | 0.06899 |
| MASC_ISR_1PC | -3.07968 | 0.58848 | -5.23330 | 0.00000 |
| VS_noRT | -8.60418 | 0.58848 | -14.62106 | 0.00000 |
| VS_RT | -3.03255 | 0.58848 | -5.15320 | 0.00000 |
| VS_RT_GA | -0.17948 | 0.58848 | -0.30499 | 0.76068 |
| (2) MASC -(Intercept) | -5.42520 | 0.48049 | -11.29096 | 0.00000 |
| MASC_noRT | -3.16436 | 0.67952 | -4.65679 | 0.00001 |
| MASC_VISUAL | -3.13812 | 0.67952 | -4.61817 | 0.00001 |
| MASC_MOTOR | 0.48880 | 0.67952 | 0.71934 | 0.47273 |
| MASC_ISR_C | 0.26973 | 0.67952 | 0.39694 | 0.69181 |
| MASC_ISR_1PC | -1.73429 | 0.67952 | -2.55224 | 0.01141 |
| VS_noRT | -7.25879 | 0.67952 | -10.68229 | 0.00000 |
| VS_RT | -1.68715 | 0.67952 | -2.48288 | 0.01382 |
| VS_RT_GA | 1.16591 | 0.67952 | 1.71580 | 0.08767 |

| B- REGRESSIONS: SD | Estimate | Std. Error | t value | Pr(>\|t\|) |
|---|---|---|---|---|
| (1) FSC -(Intercept) | 4.60834 | 0.20043 | 22.99231 | 0.00000 |
| MASC | 2.28149 | 0.34715 | 6.57197 | 0.00000 |
| MASC_noRT | 0.45640 | 0.34715 | 1.31468 | 0.19005 |
| MASC_VISUAL | 0.77964 | 0.34715 | 2.24580 | 0.02576 |
| MASC_MOTOR | 4.67321 | 0.34715 | 13.46146 | 0.00000 |
| MASC_ISR_C | -2.79213 | 0.34715 | -8.04291 | 0.00000 |
| MASC_ISR_1PC | -0.53787 | 0.34715 | -1.54937 | 0.12280 |
| VS_noRT | -1.07525 | 0.34715 | -3.09734 | 0.00222 |
| VS_RT | -1.19853 | 0.34715 | -3.45243 | 0.00067 |
| VS_RT_GA | -3.21785 | 0.34715 | -9.26923 | 0.00000 |
| (2) MASC -(Intercept) | 6.88984 | 0.28345 | 24.30701 | 0.00000 |
| MASC_noRT | -1.82509 | 0.40086 | -4.55295 | 0.00001 |
| MASC_VISUAL | -1.50185 | 0.40086 | -3.74658 | 0.00023 |
| MASC_MOTOR | 2.39172 | 0.40086 | 5.96647 | 0.00000 |
| MASC_ISR_C | -5.07362 | 0.40086 | -12.65686 | 0.00000 |
| MASC_ISR_1PC | -2.81936 | 0.40086 | -7.03329 | 0.00000 |
| VS_noRT | -3.35675 | 0.40086 | -8.37387 | 0.00000 |
| VS_RT | -3.48002 | 0.40086 | -8.68139 | 0.00000 |
| VS_RT_GA | -5.49935 | 0.40086 | -13.71888 | 0.00000 |

LMs were fitted separately to the GMM-estimated mean (A) and SD (B) of the largest negative component in individual distributions of saccade lengths (in letters), with Data Set (10 levels, excluding CSL) as a categorical predictor. Panel 1: Regression coefficients for the main LM, with Data-Set reference level set to FSC. Panel 2: Data-Set reference level set to MASC.

**Length of progressive saccades.** Supplementary Table 19A indicates that the estimated mean length of forward saccades in FSC readers was of about 8.45 letters (Panel 1). In MASC models, it was smaller, but the difference was at most of about 2 letters as for MASC_MOTOR, and less than 1 letter for MASC_noRT, MASC_VISUAL, and MASC_ISR_1PC, being only marginally significant in MASC_VISUAL (Panel 1). For both MASC and MASC_ISR_C, which did not differ significantly from one another (Panel 2), the difference was of about 1.44 and 1.81 letters (Panel 1). VS_RT_GA also underestimated the mean length of forward saccades, but more greatly than MASC (Panel 2), by as much as 4.74 letters (Panel 1), while VS_RT overestimated it by more than 2 letters (Panels 1-2). VS_noRT did slightly better than MASC (Panel 2), making forward saccades that were on average only 1.13 letters too large (Panel 1), but in that case, as for other VS models, the mean corresponded to only a portion (62%) of forward saccades (Supplementary Table 16). Across non-MASC models, only CSL beat MASC; its forward saccades were only about 0.5-letter shorter than FSC readers (Panel 1; see also Panel 2).

However, as shown in Supplementary Table 19B, neither CSL nor the other comparison models beat MASC when it came to variability. CSL and VS_RT_GA were the worst; they underestimated the SD in FSC readers by as much as 2.55 and 2.75 letters (Panel 1). For VS_noRT, just as for MASC, the SD for the largest positive mixture component was non-significantly different from that in FSC readers ($p >= 0.16750$; Panel 1), but it represented only 62% of the progressive saccades in VS_noRT against 100% in MASC (Supplementary Table 16). Moreover, for all other models, the SD differed significantly from that estimated for FSC readers (Panel 1) and MASC (Panel 2), being either smaller, as in MASC_VISUAL, MASC_MOTOR, MASC_ISR_C and MASC_ISR_1PC (estimates: -0.22578 to -0.78800; Panel 1), or larger, as in MASC_noRT and VS_RT (estimates: 0.73329, 0.47401; Panel 1), and despite the largest mixture component in VS_RT accounting for only 67% of forward saccades (Supplementary Table 16).

**Supplementary Table 19 | LM regression coefficients for the mean and SD of the length of progressive saccades in (comparison) models and FSC readers.**

| A- PROGRESSIONS: MEAN | Estimate | Std. Error | t value | Pr(>\|t\|) |
|---|---|---|---|---|
| **(1) FSC -(Intercept)** | 8.45331 | 0.15708 | 53.81661 | 0.00000 |
| **MASC** | -1.43684 | 0.27206 | -5.28125 | 0.00000 |
| **MASC_noRT** | -0.60116 | 0.27206 | -2.20964 | 0.02812 |
| **MASC_VISUAL** | -0.48720 | 0.27206 | -1.79077 | 0.07465 |
| **MASC_MOTOR** | -2.06563 | 0.27206 | -7.59246 | 0.00000 |
| **MASC_ISR_C** | -1.81427 | 0.27206 | -6.66856 | 0.00000 |
| **MASC_ISR_1PC** | -0.79696 | 0.27206 | -2.92930 | 0.00374 |
| **VS_noRT** | 1.12964 | 0.27206 | 4.15210 | 0.00005 |
| **VS_RT** | 2.37931 | 0.27206 | 8.74539 | 0.00000 |
| **VS_RT_GA** | -4.74303 | 0.27206 | -17.43349 | 0.00000 |
| **CSL** | -0.52314 | 0.27206 | -1.92285 | 0.05574 |
| **(2) MASC -(Intercept)** | 7.01648 | 0.22214 | 31.58591 | 0.00000 |
| **MASC_noRT** | 0.83567 | 0.31415 | 2.66009 | 0.00836 |
| **MASC_VISUAL** | 0.94964 | 0.31415 | 3.02285 | 0.00279 |
| **MASC_MOTOR** | -0.62880 | 0.31415 | -2.00156 | 0.04651 |
| **MASC_ISR_C** | -0.37744 | 0.31415 | -1.20144 | 0.23082 |
| **MASC_ISR_1PC** | 0.63988 | 0.31415 | 2.03685 | 0.04282 |
| **VS_noRT** | 2.56647 | 0.31415 | 8.16952 | 0.00000 |
| **VS_RT** | 3.81615 | 0.31415 | 12.14743 | 0.00000 |
| **VS_RT_GA** | -3.30619 | 0.31415 | -10.52415 | 0.00000 |
| **CSL** | 0.91370 | 0.31415 | 2.90846 | 0.00399 |

| B- PROGRESSIONS: SD | Estimate | Std. Error | t value | Pr(>\|t\|) |
|---|---|---|---|---|
| **(1) FSC -(Intercept)** | 3.02318 | 0.04896 | 61.74867 | 0.00000 |
| **MASC** | 0.07597 | 0.08480 | 0.89592 | 0.37124 |
| **MASC_noRT** | 0.73329 | 0.08480 | 8.64722 | 0.00000 |
| **MASC_VISUAL** | -0.22578 | 0.08480 | -2.66251 | 0.00831 |
| **MASC_MOTOR** | -0.27605 | 0.08480 | -3.25532 | 0.00130 |
| **MASC_ISR_C** | -0.78800 | 0.08480 | -9.29240 | 0.00000 |
| **MASC_ISR_1PC** | -0.25545 | 0.08480 | -3.01235 | 0.00288 |
| **VS_noRT** | 0.11742 | 0.08480 | 1.38467 | 0.16750 |
| **VS_RT** | 0.47401 | 0.08480 | 5.58971 | 0.00000 |
| **VS_RT_GA** | -2.75327 | 0.08480 | -32.46775 | 0.00000 |
| **CSL** | -2.55544 | 0.08480 | -30.13481 | 0.00000 |
| **(2) MASC -(Intercept)** | 3.09915 | 0.06924 | 44.76018 | 0.00000 |
| **MASC_noRT** | 0.65731 | 0.09792 | 6.71282 | 0.00000 |
| **MASC_VISUAL** | -0.30176 | 0.09792 | -3.08169 | 0.00231 |
| **MASC_MOTOR** | -0.35203 | 0.09792 | -3.59508 | 0.00040 |
| **MASC_ISR_C** | -0.86397 | 0.09792 | -8.82335 | 0.00000 |
| **MASC_ISR_1PC** | -0.33142 | 0.09792 | -3.38466 | 0.00084 |
| **VS_noRT** | 0.04145 | 0.09792 | 0.42327 | 0.67250 |
| **VS_RT** | 0.39803 | 0.09792 | 4.06494 | 0.00007 |
| **VS_RT_GA** | -2.82924 | 0.09792 | -28.89378 | 0.00000 |
| **CSL** | -2.63141 | 0.09792 | -26.87340 | 0.00000 |

LMs were fitted separately to the GMM-estimated mean (A) and SD (B) of the largest positive component in individual distributions of saccade lengths (in letters), with Data Set (11 levels) as a categorical predictor. Panel 1: Regression coefficients for the main LMs, with FSC as reference level. Panel 2: MASC as reference level.

**Word-skipping behavior**

**The relationship between word skipping and word length.** Supplementary Table 20 shows the fixed effects of GLMMs for the relationship between the likelihood of word skipping and word length[2]. In accordance with the above analyses (Supplementary Table 5), MASC's skipping rate decreased with increasing word length (logit: -0.41829; Panel 2), but at a slightly slower rate compared to FSC readers (logit: -0.69546; interaction logit: 0.27720; Panel 1; Extended Data Fig.5b). This was due to MASC skipping shorter words slightly less often than FSC readers (in about 45% against 56% of the cases for 4-letter words; logit: -0.41685; Panel 1). Most MASC models behaved similarly: all, but MASC_VISUAL and MASC_noRT, skipped short (4-letter) words slightly less than FSC readers (38-44%; logit: -0.72507 to -0.49319; Panel 1), and all with no exception showed a significant effect of word length (Panels 3-7), that was slightly smaller than in FSC readers (logit: 0.20419 to 0.31884; Panel 1; Extended Data Fig. 5h, 6b, 7b). For MASC_VISUAL and MASC_ISR_1PC, the word-length effect did not differ significantly from that observed for MASC ($p$'s >= 0.13885; Panel 2), but for other MASC models, it differed significantly, being either slightly greater or slightly smaller (logit: -0.07300 to 0.04169; Panel 2).

In VS models, the likelihood of word skipping also decreased significantly with word length (Panels 8-10), but only very mildly, the effect being much smaller than that observed in FSC readers (logit: 0.55356-0.67661; Panel 1; Fig. 3h) and MASC (logit: 0.27641-0.39948; Panel 2). Moreover, the skipping rate was overall too high or too low: 4-letter words were skipped in about 79%, 61%, and 32% of the cases in VS_noRT, VS_RT, and VS_RT_GA. In contrast, CSL only mildly underestimated FSC readers' skipping rate (by about 9%; Panel 1), and it predicted nearly the same decrease with increasing word length, as suggested by the non-significant interaction between word length and CSL ($p$ = 0.99816; Panel 1; Fig. 3b).

**Supplementary Table 20 | Fixed effects of GLMMs for the probability of word skipping by word length in (comparison) models and FSC readers.**

| SKIPPING BY WL | | Estimate | Std. Error | z value | Pr(>\|z\|) | Proportion |
|---|---|---|---|---|---|---|
| (1) | FSC -(Intercept) | 0.23646 | 0.04038 | 5.85592 | 0.00000 | 0.55884 |
| | WL | -0.69546 | 0.00801 | -86.85771 | 0.00000 | |
| | MASC | -0.41685 | 0.06946 | -6.00098 | 0.00000 | 0.45502 |
| | MASC_noRT | -0.08930 | 0.06944 | -1.28605 | 0.19843 | 0.53672 |
| | MASC_VISUAL | 0.06402 | 0.06939 | 0.92260 | 0.35622 | 0.57456 |
| | MASC_MOTOR | -0.72507 | 0.06944 | -10.44177 | 0.00000 | 0.38022 |
| | MASC_ISR_C | -0.70339 | 0.07059 | -9.96416 | 0.00000 | 0.38534 |
| | MASC_ISR_1PC | -0.49319 | 0.06925 | -7.12133 | 0.00000 | 0.43617 |
| | VS_noRT | 1.11360 | 0.07281 | 15.29409 | 0.00000 | 0.79414 |
| | VS_RT | 0.22164 | 0.06935 | 3.19621 | 0.00139 | 0.61256 |
| | VS_RT_GA | -0.99219 | 0.06918 | -14.34219 | 0.00000 | 0.31957 |
| | CSL | -0.36172 | 0.06941 | -5.21109 | 0.00000 | 0.46872 |
| | WL:MASC | 0.27720 | 0.01162 | 23.85615 | 0.00000 | |
| | WL:MASC_noRT | 0.31884 | 0.01159 | 27.50485 | 0.00000 | |
| | WL:MASC_VISUAL | 0.27774 | 0.01096 | 25.33117 | 0.00000 | |
| | WL:MASC_MOTOR | 0.20816 | 0.01208 | 17.23824 | 0.00000 | |
| | WL:MASC_ISR_C | 0.20419 | 0.01411 | 14.47260 | 0.00000 | |
| | WL:MASC_ISR_1PC | 0.29482 | 0.01164 | 25.33495 | 0.00000 | |
| | WL:VS_noRT | 0.67661 | 0.01414 | 47.85021 | 0.00000 | |
| | WL:VS_RT | 0.62705 | 0.01042 | 60.15225 | 0.00000 | |
| | WL:VS_RT_GA | 0.55356 | 0.01046 | 52.90134 | 0.00000 | |
| | WL:CSL | 0.00003 | 0.01332 | 0.00230 | 0.99816 | |
| (2) | MASC -(Intercept) | -0.18040 | 0.05524 | -3.26556 | 0.00109 | |
| | WL | -0.41829 | 0.00842 | -49.69195 | 0.00000 | |
| | MASC_noRT | 0.32760 | 0.07909 | 4.14214 | 0.00003 | |
| | MASC_VISUAL | 0.48087 | 0.07870 | 6.11036 | 0.00000 | |
| | MASC_MOTOR | -0.30830 | 0.07894 | -3.90551 | 0.00009 | |
| | MASC_ISR_C | -0.28641 | 0.08010 | -3.57559 | 0.00035 | |
| | MASC_ISR_1PC | -0.07633 | 0.07885 | -0.96815 | 0.33297 | |
| | VS_noRT | 1.53055 | 0.08205 | 18.65306 | 0.00000 | |
| | VS_RT | 0.63860 | 0.07882 | 8.10152 | 0.00000 | |
| | VS_RT_GA | -0.57515 | 0.07858 | -7.31912 | 0.00000 | |
| | CSL | 0.05525 | 0.07885 | 0.70075 | 0.48346 | |
| | WL:MASC_noRT | 0.04169 | 0.01188 | 3.50887 | 0.00045 | |
| | WL:MASC_VISUAL | 0.00056 | 0.01127 | 0.05001 | 0.96012 | |
| | WL:MASC_MOTOR | -0.06903 | 0.01235 | -5.58820 | 0.00000 | |
| | WL:MASC_ISR_C | -0.07300 | 0.01435 | -5.08773 | 0.00000 | |
| | WL:MASC_ISR_1PC | 0.01765 | 0.01192 | 1.48007 | 0.13885 | |
| | WL:VS_noRT | 0.39948 | 0.01438 | 27.78354 | 0.00000 | |
| | WL:VS_RT | 0.34989 | 0.01074 | 32.56651 | 0.00000 | |
| | WL:VS_RT_GA | 0.27641 | 0.01078 | 25.63762 | 0.00000 | |
| | WL:CSL | -0.27713 | 0.01358 | -20.41406 | 0.00000 | |
| (3) | MASC_noRT -(Intercept) | 0.14705 | 0.05561 | 2.64439 | 0.00818 | |
| | WL | -0.37664 | 0.00838 | -44.94931 | 0.00000 | |
| (4) | MASC_VISUAL -(Intercept) | 0.30032 | 0.05535 | 5.42567 | 0.00000 | |
| | WL | -0.41774 | 0.00749 | -55.77258 | 0.00000 | |
| (5) | MASC_MOTOR -(Intercept) | -0.48887 | 0.05630 | -8.68394 | 0.00000 | |

| | | | | |
|---|---|---|---|---|
| **WL** | -0.48734 | 0.00904 | -53.93726 | 0.00000 |
| **(6) MASC_ISR_C (Intercept)** | -0.46630 | 0.05721 | -8.15108 | 0.00000 |
| **WL** | -0.49113 | 0.01161 | -42.28759 | 0.00000 |
| **(7) MASC_ISR_1PC -(Intercept)** | -0.25680 | 0.05575 | -4.60610 | 0,00000 |
| **WL** | -0.40067 | 0.00844 | -47.46224 | 0,00000 |
| **(8) VS_noRT -(Intercept)** | 1.35007 | 0.05964 | 22.63810 | 0.00000 |
| **WL** | -0.01897 | 0.01165 | -1.62890 | 0.10334 |
| **(9) VS_RT -(Intercept)** | 0.45798 | 0.05584 | 8.20133 | 0.00000 |
| **WL** | -0.06842 | 0.00667 | -10.25029 | 0.00000 |
| **(10) VS_RT_GA -(Intercept)** | -0.75580 | 0.05571 | -13.56603 | 0.00000 |
| **WL** | -0.14189 | 0.00674 | -21.06652 | 0.00000 |
| **(11) CSL -(Intercept)** | -0.12489 | 0.05672 | -2.20183 | 0.02768 |
| **WL** | -0.69536 | 0.01065 | -65.27734 | 0.00000 |

GLMMs were fitted to a binary variable indicating whether a given word was skipped. The fixed structure included Data Set (11 levels), Word Length (WL; 1-11 letters; reference (mean) value: 4.03 letters), and their interaction as predictors. The random structure included a random intercept by subject; with a more complex random structure, the model did not converge. Estimates and standard errors are expressed in logit unit; the estimated probability of word skipping in each data set when WL was at its reference value is given in the rightmost column of Panel 1. Panel 1: Fixed-effects for the main GLMM, with Data-Set reference level set to FSC. Panels 2-11: Data-Set reference level set to MASC, MASC_noRT, MASC_VISUAL, MASC_MOTOR, MASC_ISR_C, MASC_ISR_1PC, VS_noRT, VS_RT, VS_RT_GA, and CSL respectively. In Panel 2, the fixed effects that were redundant with those reported in Panel 1 were dropped. In Panels 3-11, only the intercept and the effect of WL are reported. These analyses relied on a total of 34957, 20448, 18429, 21244, 22301, 13029, 19956, 8407, 17090, 21235, 20553 cases across subjects in FSC, MASC, MASC_noRT, MASC_VISUAL, MASC_MOTOR, MASC_ISR_C, MASC_ISR_1PC, VS_noRT, VS_RT, VS_RT_GA, and CSL respectively.

Thus, the relationship between word skipping rate and word length was best captured by CSL, and then MASC (models). Still, MASC (models) remained the best option, as it (they) left space for tiny language-related modulations of the word-length effect, as observed in FSC readers[7], and in line with previously reported differences between the reading of normal and z-transformed texts[19] (Extended Data Fig. 1a,c-e).

**The relationship between word skipping and launch site.** MASC (models) also beat VS models, as well as CSL, for the relationship between the likelihood of word skipping and saccades' launch-site distance to the space in front of the words[3-4], as indicated in Supplementary Table 21. In FSC readers, the word-skipping rate, of about 70% (logit: 0.83733; Panel 1) when both word length and launch-site distance were at their reference (mean) value (3.70 letters and -2.46 letters from the space in front of the words respectively), decreased as words were longer (logit: -1.11297;

Panel 1) and more eccentric (logit: 0.42632; Panel 1). All MASC models reproduced these effects (Panels 2-7; Extended Data Fig. 5c,i, 6c, 7c). All, but MASC_ISR_C and MASC_VISUAL, predicted a slightly weaker effect of word length compared to FSC readers (logit: 0.12216-0.29239; Panel 1); MASC_ISR_C slightly overestimated the effect (logit: -0.22936; Panel 1), and MASC_VISUAL perfectly predicted the effect observed in FSC readers ($p$ = 0.66122; Panel 1). Moreover, all MASC models predicted a greater effect of launch-site distance compared to FSC readers (logit: 0.13949-0.61400; Panel 1). MASC_VISUAL, MASC_MOTOR, MASC_ISR_C, and MASC_ISR_1PC however showed an even greater effect than MASC (logit: 0.06772-0.29729; Panel 2). MASC_noRT predicted to the contrary a smaller effect than MASC (logit: -0.17722; Panel 2), thus better approximating the effect observed in FSC readers. MASC was thus second best among MASC models for the relationship between word-skipping rate and launch site.

For VS models, they all predicted a significant reduction in word skipping rate with increasing word length (Panels 8-10), but this was much smaller than in FSC readers (logit: 0.42281-0.61388; Panel 1), and MASC (logit: 0.21726-0.40811; Panel 2; see also Supplementary Table 20). In addition, VS_RT_GA, predicted an increase, and not a decrease, of word-skipping rate with increasing launch-site distance (estimate: -0.14170; Panel 10), thus in contrast with the effect observed in FSC readers (logit: -0.56856; Panel 1; Fig. 3i) and MASC (logit: -0.88531; Panel 2). Furthermore, although VS_noRT and VS_RT rightly predicted a reduction in word-skipping rate with increasing launch-site distance (Panels 8-9), this effect was weaker than for FSC readers (logit: -0.11188 and -0.11473; Panel 1), rather than being stronger as would be expected in the absence of linguistic processing (Extended Data Fig. 1b), and as predicted by MASC. MASC (models) therefore remained more plausible than VS models. CSL was also not a viable alternative (Fig. 3c). It predicted much greater effects of word length (in contrast with analyses using only word length as a predictor; Supplementary Table 20) and launch-site distance than FSC readers (logit: -2.81498 and 3.52285; Panel 1) and MASC (logit: -3.02071 and 3.20604; Panel 2).

**Supplementary Table 21 | Fixed effects of GLMMs for the probability of word skipping by word length and launch site in (comparison) models and FSC readers.**

| SKIPPING BY LS | | Estimate | Std. Error | z value | Pr(>\|z\|) | Proportion |
|---|---|---|---|---|---|---|
| (1) | FSC -(Intercept) | 0.83733 | 0.05605 | 14.93834 | 0.00000 | 0.69790 |
| | WL | -1.11297 | 0.02835 | -39.25495 | 0.00000 | |
| | LS | 0.42632 | 0.01354 | 31.48728 | 0.00000 | |
| | MASC | -0.56033 | 0.09564 | -5.85852 | 0.00000 | 0.56881 |
| | MASC_noRT | -0.19699 | 0.09589 | -2.05420 | 0.03996 | 0.65483 |
| | MASC_VISUAL | 0.51117 | 0.09944 | 5.14055 | 0.00000 | 0.79388 |
| | MASC_MOTOR | -0.96548 | 0.09575 | -10.08331 | 0.00000 | 0.46801 |
| | MASC_ISR_C | -0.72308 | 0.09940 | -7.27436 | 0.00000 | 0.52853 |
| | MASC_ISR_1PC | -0.54280 | 0.09598 | -5.65553 | 0.00000 | 0.57311 |
| | VS_noRT | 1.03330 | 0.10697 | 9.66014 | 0.00000 | 0.86653 |
| | VS_RT | 0.21550 | 0.09610 | 2.24252 | 0.02493 | 0.74132 |
| | VS_RT_GA | -1.94769 | 0.09492 | -20.51995 | 0.00000 | 0.24780 |
| | CSL | 1.95360 | 0.14222 | 13.73653 | 0.00000 | 0.94218 |
| | WL:LS | -0.05969 | 0.01728 | -3.45484 | 0.00055 | |
| | WL:MASC | 0.20566 | 0.04710 | 4.36605 | 0.00001 | |
| | WL:MASC_noRT | 0.29239 | 0.04705 | 6.21399 | 0.00000 | |
| | WL:MASC_VISUAL | -0.02338 | 0.05335 | -0.43823 | 0.66122 | |
| | WL:MASC_MOTOR | 0.12216 | 0.04888 | 2.49941 | 0.01244 | |
| | WL:MASC_ISR_C | -0.22936 | 0.06088 | -3.76731 | 0.00017 | |
| | WL:MASC_ISR_1PC | 0.13013 | 0.04800 | 2.71134 | 0.00670 | |
| | WL:VS_noRT | 0.42281 | 0.06927 | 6.10380 | 0.00000 | |
| | WL:VS_RT | 0.58684 | 0.04575 | 12.82747 | 0.00000 | |
| | WL:VS_RT_GA | 0.61388 | 0.04555 | 13.47756 | 0.00000 | |
| | WL:CSL | -2.81498 | 0.13509 | -20.83766 | 0.00000 | |
| | LS:MASC | 0.31683 | 0.02421 | 13.08916 | 0.00000 | |
| | LS:MASC_noRT | 0.13949 | 0.02339 | 5.96404 | 0.00000 | |
| | LS:MASC_VISUAL | 0.61400 | 0.02862 | 21.45325 | 0.00000 | |
| | LS:MASC_MOTOR | 0.51041 | 0.02546 | 20.04604 | 0.00000 | |
| | LS:MASC_ISR_C | 0.47241 | 0.03161 | 14.94415 | 0.00000 | |
| | LS:MASC_ISR_1PC | 0.38446 | 0.02511 | 15.31207 | 0.00000 | |
| | LS:VS_noRT | -0.11188 | 0.03425 | -3.26651 | 0.00109 | |
| | LS:VS_RT | -0.11473 | 0.02220 | -5.16865 | 0.00000 | |
| | LS:VS_RT_GA | -0.56856 | 0.02290 | -24.83270 | 0.00000 | |
| | LS:CSL | 3.52285 | 0.11718 | 30.06357 | 0.00000 | |
| | WL:LS:MASC | 0.18060 | 0.03209 | 5.62873 | 0.00000 | |
| | WL:LS:MASC_noRT | 0.18583 | 0.03102 | 5.99061 | 0.00000 | |
| | WL:LS:MASC_VISUAL | 0.10222 | 0.03516 | 2.90740 | 0.00364 | |
| | WL:LS:MASC_MOTOR | 0.11931 | 0.03326 | 3.58727 | 0.00033 | |
| | WL:LS:MASC_ISR_C | 0.14178 | 0.04161 | 3.40784 | 0.00065 | |
| | WL:LS:MASC_ISR_1PC | 0.13975 | 0.03266 | 4.27944 | 0.00002 | |
| | WL:LS:VS_noRT | -0.06743 | 0.04233 | -1.59304 | 0.11115 | |
| | WL:LS:VS_RT | -0.06450 | 0.02830 | -2.27957 | 0.02263 | |
| | WL:LS:VS_RT_GA | -0.04795 | 0.03083 | -1.55529 | 0.11988 | |

| | | | | | |
|---|---|---|---|---|---|
| | WL:LS:CSL | 0.27197 | 0.09070 | 2.99866 | 0.00271 |
| **(2)** | **MASC -(Intercept)** | 0.27744 | 0.07756 | 3.57699 | 0.00035 |
| | **WL** | -0.90718 | 0.03762 | -24.11169 | 0.00000 |
| | **LS** | 0.74309 | 0.02007 | 37.02831 | 0.00000 |
| | **MASC_noRT** | 0.36300 | 0.10987 | 3.30398 | 0.00095 |
| | **MASC_VISUAL** | 1.07116 | 0.11298 | 9.48135 | 0.00000 |
| | **MASC_MOTOR** | -0.40548 | 0.10974 | -3.69503 | 0.00022 |
| | **MASC_ISR_C** | -0.16300 | 0.11293 | -1.44330 | 0.14893 |
| | **MASC_ISR_1PC** | 0.01713 | 0.10995 | 0.15579 | 0.87620 |
| | **VS_noRT** | 1.59326 | 0.11969 | 13.31172 | 0.00000 |
| | **VS_RT** | 0.77537 | 0.11005 | 7.04581 | 0.00000 |
| | **VS_RT_GA** | -1.38747 | 0.10903 | -12.72565 | 0.00000 |
| | **CSL** | 2.51326 | 0.15230 | 16.50198 | 0.00000 |
| | **WL:LS** | 0.12080 | 0.02704 | 4.46808 | 0.00001 |
| | **WL:MASC_noRT** | 0.08661 | 0.05316 | 1.62920 | 0.10327 |
| | **WL:MASC_VISUAL** | -0.22921 | 0.05881 | -3.89720 | 0.00010 |
| | **WL:MASC_MOTOR** | -0.08355 | 0.05478 | -1.52523 | 0.12720 |
| | **WL:MASC_ISR_C** | -0.43514 | 0.06573 | -6.62043 | 0.00000 |
| | **WL:MASC_ISR_1PC** | -0.07571 | 0.05400 | -1.40196 | 0.16093 |
| | **WL:VS_noRT** | 0.21726 | 0.07358 | 2.95286 | 0.00315 |
| | **WL:VS_RT** | 0.38107 | 0.05201 | 7.32722 | 0.00000 |
| | **WL:VS_RT_GA** | 0.40811 | 0.05183 | 7.87371 | 0.00000 |
| | **WL:CSL** | -3.02071 | 0.13767 | -21.94089 | 0.00000 |
| | **LS:MASC_noRT** | -0.17722 | 0.02769 | -6.40086 | 0.00000 |
| | **LS:MASC_VISUAL** | 0.29729 | 0.03223 | 9.22316 | 0.00000 |
| | **LS:MASC_MOTOR** | 0.19366 | 0.02946 | 6.57351 | 0.00000 |
| | **LS:MASC_ISR_C** | 0.15569 | 0.03492 | 4.45881 | 0.00001 |
| | **LS:MASC_ISR_1PC** | 0.06772 | 0.02916 | 2.32276 | 0.02019 |
| | **LS:VS_noRT** | -0.42868 | 0.03732 | -11.48585 | 0.00000 |
| | **LS:VS_RT** | -0.43150 | 0.02669 | -16.16841 | 0.00000 |
| | **LS:VS_RT_GA** | -0.88531 | 0.02727 | -32.46566 | 0.00000 |
| | **LS:CSL** | 3.20604 | 0.11837 | 27.08505 | 0.00000 |
| | **WL:LS:MASC_noRT** | 0.00543 | 0.03735 | 0.14536 | 0.88443 |
| | **WL:LS:MASC_VISUAL** | -0.07823 | 0.04085 | -1.91494 | 0.05550 |
| | **WL:LS:MASC_MOTOR** | -0.06122 | 0.03923 | -1.56076 | 0.11858 |
| | **WL:LS:MASC_ISR_C** | -0.03892 | 0.04651 | -0.83673 | 0.40275 |
| | **WL:LS:MASC_ISR_1PC** | -0.04067 | 0.03872 | -1.05053 | 0.29348 |
| | **WL:LS:VS_noRT** | -0.24794 | 0.04716 | -5.25716 | 0.00000 |
| | **WL:LS:VS_RT** | -0.24499 | 0.03512 | -6.97676 | 0.00000 |
| | **WL:LS:VS_RT_GA** | -0.22845 | 0.03719 | -6.14274 | 0.00000 |
| | **WL:LS:CSL** | 0.09165 | 0.09313 | 0.98416 | 0.32504 |
| **(3)** | **MASC_noRT -(Intercept)** | 0.64043 | 0.07782 | 8.23007 | 0.00000 |
| | **WL** | -0.82093 | 0.03756 | -21.85758 | 0.00000 |
| | **LS** | 0.56569 | 0.01907 | 29.65871 | 0.00000 |
| | **WL:LS** | 0.12597 | 0.02576 | 4.88956 | 0.00000 |
| **(4)** | **MASC_VISUAL -(Intercept)** | 1.34773 | 0.08218 | 16.39896 | 0.00000 |
| | **WL** | -1.13632 | 0.04521 | -25.13310 | 0.00000 |
| | **LS** | 1.03990 | 0.02522 | 41.23567 | 0.00000 |
| | **WL:LS** | 0.04211 | 0.03062 | 1.37553 | 0.16897 |
| **(5)** | **MASC_MOTOR -(Intercept)** | -0.12795 | 0.07770 | -1.64671 | 0.09962 |
| | **WL** | -0.99081 | 0.03983 | -24.87673 | 0.00000 |
| | **LS** | 0.93666 | 0.02157 | 43.42564 | 0.00000 |

| | | | | |
|---|---|---|---|---|
| WL:LS | 0.05938 | 0.02842 | 2.08906 | 0.03670 |
| (6) MASC_ISR_C (Intercept) | 0.11438 | 0.08216 | 1.39223 | 0.16385 |
| WL | -1.34115 | 0.05388 | -24.89135 | 0.00000 |
| LS | 0.89861 | 0.02857 | 31.45331 | 0.00000 |
| WL:LS | 0.08202 | 0.03785 | 2.16730 | 0.03021 |
| (7) MASC_ISR_1PC -(Intercept) | 0.29466 | 0.07789 | 3.78319 | 0.00015 |
| WL | -0.98267 | 0.03873 | -25.37062 | 0.00000 |
| LS | 0.81077 | 0.02115 | 38.33656 | 0.00000 |
| WL:LS | 0.08018 | 0.02771 | 2.89361 | 0.00381 |
| (8) VS_noRT -(Intercept) | 1.86775 | 0.09114 | 20.49333 | 0.00000 |
| WL | -0.68920 | 0.06316 | -10.91125 | 0.00000 |
| LS | 0.31348 | 0.03145 | 9.96789 | 0.00000 |
| WL:LS | -0.12690 | 0.03863 | -3.28470 | 0.00102 |
| (9) VS_RT -(Intercept) | 1.05286 | 0.07806 | 13.48760 | 0.00000 |
| WL | -0.52642 | 0.03591 | -14.65904 | 0.00000 |
| LS | 0.31164 | 0.01759 | 17.71315 | 0.00000 |
| WL:LS | -0.12418 | 0.02241 | -5.54189 | 0.00000 |
| (10) VS_RT_GA -(Intercept) | -1.10791 | 0.07660 | -14.46391 | 0.00000 |
| WL | -0.49809 | 0.03563 | -13.97942 | 0.00000 |
| LS | -0.14170 | 0.01846 | -7.67694 | 0.00000 |
| WL:LS | -0.10676 | 0.02553 | -4.18192 | 0.00003 |
| (11) CSL -(Intercept) | 2.77996 | 0.13074 | 21.26271 | 0.00000 |
| WL | -3.91010 | 0.13212 | -29.59551 | 0.00000 |
| LS | 3.93536 | 0.11634 | 33.82719 | 0.00000 |
| WL:LS | 0.21142 | 0.08896 | 2.37640 | 0.01748 |

GLMMs were fitted to a binary variable indicating whether a given word was skipped. The fixed structure included Data Set (11 levels), Word Length (WL; 3-5 letters; reference (mean) value: 3.70), Launch-Site distance (LS; 0-6 letters from the space in front of the words; reference (mean) value: -2.46 letters) and all interactions as predictors. The random structure included a random intercept by subject; with a more complex random structure, the model did not converge. Estimates and standard errors are expressed in logit units; the estimated probability of word skipping in each data set when WL and LS were at their reference (mean) value is given in the rightmost column (in Panel 1). Panel 1: Fixed-effects of the main LMM, with Data-Set reference level set to FSC. Panels 2-11: Data-Set reference level set to MASC, MASC_noRT, MASC_VISUAL, MASC_MOTOR, MASC_ISR_C, MASC_ISR_1PC, VS_noRT, VS_RT, VS_RT_GA, and CSL respectively. In Panels 3-11, only the intercept, the effects of WL and LS, and the interaction are reported. These analyses relied on a total of 13638, 7816, 6934, 7953, 8691, 4953, 7697, 3395, 6811, 8731, 8129 cases across subjects in FSC, MASC, MASC_noRT, MASC_VISUAL, MASC_MOTOR, MASC_ISR_C, MASC_ISR_1PC, VS_noRT, VS_RT, VS_RT_GA4, and CSL respectively.

### Within-word landing-positions

**The PVL effect.** As shown in Supplementary Table 22, most within-word landing-position distributions in FSC readers were best fitted with a single mixture component (proportion of two mixture components <= 0.1), thus in line with the well-known PVL effect[5] (see also Supplementary Table 7). In MASC, the proportion of two mixture components was higher for 6- to 8-letter words (0.20-0.35), but the largest detected mode still accounted for more than 86% of the data.

MASC_VISUAL and MASC_ISR_1PC behaved similarly. MASC_noRT and MASC_MOTOR did slightly better, as two peaks were detected in only 10-15% and 5-20% of the distributions, while MASC_ISR_C, as well as CSL, did worse, predicting bimodal distributions in 20-65% and 10-55% of the cases respectively, in 6- to 9-letter words. However, VS_RT models did even worse, as also shown in Fig. 3j (for comparison see Extended Data Fig. 5d,j, 6d, 7d, and Fig. 3d).

**Supplementary Table 22 | GMM-estimated shape of within-word landing-site distributions by word length in (comparison) models and FSC readers.**

| PVL EFFECT | 4-lw | | 5-lw | | 6-lw | | 7-lw | | 8-lw | | 9-lw | |
|---|---|---|---|---|---|---|---|---|---|---|---|---|
| **2 Modes/ Largest *K*** | **2M** | $K_k$ | **2M** | $K_k$ | **2M** | $K_k$ | **2M** | $K_k$ | **2M** | $K_k$ | **2M** | $K_k$ |
| **FSC** | 0.00 | 1.00 | 0.02 | 0.99 | 0.10 | 0.96 | 0.07 | 0.97 | 0.10 | 0.96 | 0.02 | 0.99 |
| **MASC** | 0.00 | 1.00 | 0.05 | 0.98 | 0.25 | 0.90 | 0.35 | 0.86 | 0.20 | 0.92 | 0.00 | 1.00 |
| **MASC_noRT** | 0.00 | 1.00 | 0.00 | 1.00 | 0.15 | 0.94 | 0.10 | 0.96 | 0.15 | 0.95 | 0.00 | 1.00 |
| **MASC_VISUAL** | 0.00 | 1.00 | 0.10 | 0.95 | 0.20 | 0.94 | 0.30 | 0.88 | 0.20 | 0.93 | 0.10 | 0.96 |
| **MASC_MOTOR** | 0.00 | 1.00 | 0.00 | 1.00 | 0.05 | 0.98 | 0.20 | 0.91 | 0.15 | 0.95 | 0.00 | 1.00 |
| **MASC_ISR_C** | 0.00 | 1.00 | 0.05 | 0.98 | 0.30 | 0.88 | 0.60 | 0.78 | 0.65 | 0.74 | 0.20 | 0.91 |
| **MASC_ISR_1PC** | 0.00 | 1.00 | 0.00 | 1.00 | 0.35 | 0.87 | 0.20 | 0.92 | 0.20 | 0.91 | 0.15 | 0.93 |
| **VS_noRT** | 0.00 | 1.00 | 0.25 | 0.90 | 0.30 | 0.87 | 0.15 | 0.94 | 0.15 | 0.95 | 0.35 | 0.86 |
| **VS_RT** | 0.00 | 1.00 | 0.90 | 0.61 | 0.80 | 0.68 | 0.45 | 0.81 | 0.80 | 0.69 | 0.40 | 0.83 |
| **VS_RT_GA** | 0.00 | 1.00 | 0.00 | 1.00 | 0.00 | 1.00 | 0.90 | 0.65 | 0.90 | 0.64 | 0.80 | 0.73 |
| **CSL** | 0.00 | 1.00 | 0.00 | 1.00 | 0.10 | 0.96 | 0.55 | 0.80 | 0.55 | 0.79 | 0.10 | 0.96 |

Proportion of two mixture components (i.e., the maximal number of detected modes; “2M”) and mean proportion of cases accounted for by the largest mixture component (mean of largest *k* values; “***K****k*”), as estimated after fitting GMMs to individual distributions of landing positions in 4- to 9-letter words separately for each data set. These analyses relied on a total of 22287, 11006, 10080, 11338, 12052, 8806, 11208, 3398, 7764, 11312, 13041 cases across subjects in FSC, MASC, MASC_noRT, MASC_VISUAL, MASC_MOTOR, MASC_ISR_C, MASC_ISR_1PC, VS_noRT, VS_RT, VS_RT_GA, and CSL respectively.

In VS_RT, the proportion of two mixture components varied between 0.40 and 0.90 for 5- to 9-letter words, with the largest mixture component accounting for only 61-83% of the data on average. In VS_RT_GA, the proportion of two detected modes was null in short, 4- to 6-letter, words, but it ranged between 0.80 and 0.90 in 7- to 9-letter words, with the largest mode accounting for as little as 64-73% of the data on average. VS_noRT did slightly better than other VS models, as the proportion of bimodal distributions did not exceed 0.35, but this proportion was relatively constant across a wider range of word lengths (5 to 9 letters; 0.25-0.35) compared to MASC. In sum, MASC models, except MASC_ISR_C, more closely reproduced the shape of within-word landing-position distributions observed in FSC readers, than either VS or CSL.

Comparison of the estimated mean and SD of the largest mixture component confirmed the supremacy of MASC (models). As shown in the left panels of Supplementary Table 23, and in line with the above analyses, when data sets were matched for numbers of fixations (Supplementary Table 8), both FSC readers and MASC showed a leftward fixation bias in 6-letter words (estimates: -0.82656 and -0.60281 respectively; Panels 1-2); MASC only landed a bit closer to the words' center than FSC readers (estimate: 0.22375; Panel 1). Moreover, this leftward fixation bias significantly increased with increasing word length in both FSC readers and MASC (estimates: -0.17200 and -0.10968; Panel 1), and at about the same rate as suggested by the non-significant interaction between word length and MASC ($t = 1.12585$). Most alternative MASC models, as well as CSL, behaved similarly. All predicted a leftward fixation bias in 6-letter words (Panels 3-7, 11), though not always smaller than in FSC readers: for MASC_MOTOR and MASC_ISR_1PC, it was not significantly different ($t$ <= 1.32617; Panel 1), and for MASC_ISR_C, it was significantly greater (estimate: -0.52206; Panel 1). In addition, all these models, but MASC_VISUAL, predicted a (marginally) significant linear relationship between mean landing position and word length (estimates: -0.08744 to -0.20827; Panels 3-7,11), and with a slope comparable to that observed in both FSC readers ($|t|$ <= 1.52230; Panel 1) and MASC ($|t|$ <= 1.52208; Panel 2).

VS models also predicted a leftward fixation bias (estimates: -0.89225 to -1.64358; Panels 8-10). However, although this bias for VS_noRT was comparable to that observed in FSC readers ($t = -0.69569$; Panel 1), the bias in VS_RT and VS_RT_GA was significantly greater than for both FSC readers (estimates: -0.51995 and -0.81702; Panels 1) and MASC (Panel 2). Moreover, VS_noRT and VS_RT predicted no significant relationship between mean landing position and word length ($t$ <= 0.30792; Panels 8-9), whereas VS_RT_GA predicted an increase in the leftward fixation bias with increasing word length (estimate: -0.53684; Panel 10) that was much stronger than for FSC readers (estimate: -0.36401; Panel 1) and MASC (estimate: -0.42717; Panel 2).

**Supplementary Table 23 | Fixed effects of LMMs for the mean and SD of within-word landing sites by word length in (comparison) models and FSC readers**.

| PVL | EFFECT: MEAN / SD | Estimate | Std. Error | *t* value | Estimate | Std. Error | *t* value |
|---|---|---|---|---|---|---|---|
| (1) | FSC -(Intercept) | -0.82656 | 0.05451 | -15.16239 | 1.77844 | 0.02623 | 67.80098 |
| | WL | -0.17283 | 0.03239 | -5.33643 | 0.14632 | 0.01560 | 9.37834 |
| | MASC | 0.22375 | 0.09442 | 2.36973 | 0.08913 | 0.04543 | 1.96176 |
| | MASC_noRT | 0.31356 | 0.09442 | 3.32092 | 0.16335 | 0.04543 | 3.59543 |
| | MASC_VISUAL | 0.57097 | 0.09442 | 6.04709 | 0.11476 | 0.04543 | 2.52595 |
| | MASC_MOTOR | 0.07214 | 0.09442 | 0.76406 | 0.07537 | 0.04543 | 1.65900 |
| | MASC_ISR_C | -0.52206 | 0.09442 | -5.52909 | -0.05851 | 0.04543 | -1.28784 |
| | MASC_ISR_1PC | 0.12522 | 0.09442 | 1.32617 | 0.07229 | 0.04543 | 1.59114 |
| | VS_noRT | -0.06569 | 0.09442 | -0.69569 | -0.05485 | 0.04543 | -1.20739 |
| | VS_RT | -0.51995 | 0.09442 | -5.50669 | -0.27072 | 0.04543 | -5.95881 |
| | VS_RT_GA | -0.81702 | 0.09442 | -8.65294 | -0.32088 | 0.04543 | -7.06294 |
| | CSL | 0.32003 | 0.09442 | 3.38938 | -0.02844 | 0.04543 | -0.62599 |
| | WL:MASC | 0.06316 | 0.05610 | 1.12585 | 0.05887 | 0.02702 | 2.17859 |
| | WL:MASC_noRT | 0.08428 | 0.05610 | 1.50236 | 0.08287 | 0.02702 | 3.06672 |
| | WL:MASC_VISUAL | 0.19178 | 0.05610 | 3.41882 | 0.05372 | 0.02702 | 1.98777 |
| | WL:MASC_MOTOR | 0.00719 | 0.05610 | 0.12821 | 0.03501 | 0.02702 | 1.29562 |
| | WL:MASC_ISR_C | -0.03544 | 0.05610 | -0.63170 | -0.01872 | 0.02702 | -0.69277 |
| | WL:MASC_ISR_1PC | 0.08540 | 0.05610 | 1.52230 | 0.03183 | 0.02702 | 1.17787 |
| | WL:VS_noRT | 0.15873 | 0.05610 | 2.82957 | 0.04024 | 0.02702 | 1.48903 |
| | WL:VS_RT | 0.16125 | 0.05610 | 2.87455 | 0.10307 | 0.02702 | 3.81421 |
| | WL:VS_RT_GA | -0.36401 | 0.05610 | -6.48900 | -0.14139 | 0.02702 | -5.23205 |
| | WL:CSL | 0.07109 | 0.05610 | 1.26730 | -0.04929 | 0.02702 | -1.82379 |
| (2) | MASC -(Intercept) | -0.60281 | 0.07709 | -7.81911 | 1.86756 | 0.03710 | 50.34519 |
| | WL | -0.10968 | 0.04580 | -2.39455 | 0.20520 | 0.02206 | 9.29971 |
| | MASC_noRT | 0.08981 | 0.10903 | 0.82375 | 0.07422 | 0.05246 | 1.41479 |
| | MASC_VISUAL | 0.34722 | 0.10903 | 3.18469 | 0.02563 | 0.05246 | 0.48861 |
| | MASC_MOTOR | -0.15161 | 0.10903 | -1.39055 | -0.01375 | 0.05246 | -0.26220 |
| | MASC_ISR_C | -0.74581 | 0.10903 | -6.84058 | -0.14764 | 0.05246 | -2.81424 |
| | MASC_ISR_1PC | -0.09853 | 0.10903 | -0.90375 | -0.01684 | 0.05246 | -0.32097 |
| | VS_noRT | -0.28944 | 0.10903 | -2.65473 | -0.14398 | 0.05246 | -2.74456 |
| | VS_RT | -0.74370 | 0.10903 | -6.82118 | -0.35985 | 0.05246 | -6.85942 |
| | VS_RT_GA | -1.04077 | 0.10903 | -9.54591 | -0.41001 | 0.05246 | -7.81562 |
| | CSL | 0.09628 | 0.10903 | 0.88305 | -0.11757 | 0.05246 | -2.24106 |
| | WL:MASC_noRT | 0.02112 | 0.06477 | 0.32607 | 0.02400 | 0.03120 | 0.76914 |
| | WL:MASC_VISUAL | 0.12863 | 0.06477 | 1.98577 | -0.00516 | 0.03120 | -0.16526 |
| | WL:MASC_MOTOR | -0.05596 | 0.06477 | -0.86398 | -0.02386 | 0.03120 | -0.76468 |
| | WL:MASC_ISR_C | -0.09859 | 0.06477 | -1.52208 | -0.07760 | 0.03120 | -2.48667 |
| | WL:MASC_ISR_1PC | 0.02224 | 0.06477 | 0.34334 | -0.02704 | 0.03120 | -0.86665 |
| | WL:VS_noRT | 0.09557 | 0.06477 | 1.47547 | -0.01863 | 0.03120 | -0.59718 |
| | WL:VS_RT | 0.09810 | 0.06477 | 1.51442 | 0.04420 | 0.03120 | 1.41648 |
| | WL:VS_RT_GA | -0.42717 | 0.06477 | -6.59465 | -0.20026 | 0.03120 | -6.41781 |
| | WL:CSL | 0.00794 | 0.06477 | 0.12251 | -0.10816 | 0.03120 | -3.46616 |
| (3) | MASC_noRT -(Intercept) | -0.51300 | 0.07709 | -6.65415 | 1.94179 | 0.03710 | 52.34601 |

| | | | | | | |
|---|---|---|---|---|---|---|
| **WL** | -0.08856 | 0.04580 | -1.93341 | 0.22920 | 0.02206 | 10.38743 |
| **(4) MASC_VISUAL -(Intercept)** | -0.25559 | 0.07709 | -3.31528 | 1.89320 | 0.03710 | 51.03618 |
| **WL** | 0.01895 | 0.04580 | 0.41376 | 0.20004 | 0.02206 | 9.06599 |
| **(5) MASC_MOTOR -(Intercept)** | -0.75442 | 0.07709 | -0.78565 | 1.85381 | 0.03710 | 49.97439 |
| **WL** | -0.16564 | 0.04580 | -3.61640 | 0.18133 | 0.02206 | 8.21829 |
| **(6) MASC_ISR_C (Intercept)** | -1.34862 | 0.07709 | -17.49315 | 1.71993 | 0.03710 | 46.36525 |
| **WL** | -0.20827 | 0.04580 | -4.54710 | 0.12760 | 0.02206 | 5.78302 |
| **(7) MASC_ISR_1PC -(Intercept)** | -0.70134 | 0.07709 | -9.09721 | 1.85073 | 0.03710 | 49.89127 |
| **WL** | -0.08744 | 0.04580 | -1.90900 | 0.17815 | 0.02206 | 8.07408 |
| **(8) VS_noRT -(Intercept)** | -0.89225 | 0.07709 | -11.57347 | 1.72358 | 0.03710 | 46.46379 |
| **WL** | -0.01410 | 0.04580 | -0.30792 | 0.18656 | 0.02206 | 8.45517 |
| **(9) VS_RT -(Intercept)** | -1.34651 | 0.07709 | -17.46572 | 1.50772 | 0.03710 | 40.64451 |
| **WL** | -0.01158 | 0.04580 | -0.25283 | 0.24940 | 0.02206 | 11.30292 |
| **(10) VS_RT_GA -(Intercept)** | -1.64358 | 0.07709 | -21.31907 | 1.45755 | 0.03710 | 39.29224 |
| **WL** | -0.53684 | 0.04580 | -11.72080 | 0.00493 | 0.02206 | 0.22356 |
| **(11) CSL -(Intercept)** | -0.50653 | 0.07709 | -6.57030 | 1.75000 | 0.03710 | 47.17586 |
| **WL** | -0.10174 | 0.04580 | -2.22130 | 0.09704 | 0.02206 | 4.39781 |

LMMs were fitted separately to the GMM-estimated mean (left panel) and SD (right panel) of the largest mixture component in individual within-word landing position distributions split by word length. In both LMMs, the fixed structure included Data Set (11 levels), Word Length (WL; 4-9 letters; reference (mean) value: 6.5 letters) and their interaction as predictors. The random structure included both a random intercept and a random effect of WL by subject. Estimates and standard errors are expressed in letters. Panel 1: Fixed-effects for the main LMM, with Data-Set reference level set to FSC. Panels 2-11: Data-Set reference level set to MASC, MASC_noRT, MASC_VISUAL, MASC_MOTOR, MASC_ISR_C, MASC_ISR_1PC, VS_noRT, VS_RT, VS_RT_GA, and CSL respectively. In Panels 3-11, only the intercept and the effect of WL are reported.

As shown in the right panels of Supplementary Table 23, the estimated SD for within-word landing positions, of about 1.78 letters for 6-letter words in FSC readers (Panel 1), increased with increasing word length (estimate: 0.14632; Panel 1). For MASC, the variability was marginally greater (estimate: 0.08913, $t$ = 1.96176; Panel 1), but it also increased with word length (estimate: 0.20520; Panel 2), though at a slightly faster rate than in FSC readers (estimate: 0.05887; Panel 1). All other models, but VS_RT_GA, also showed a spread increase with increasing word length (Panels 3-11). The slope for this relationship was either slightly greater than in FSC readers, as for MASC_noRT, MASC_VISUAL, and VS_RT (estimates: 0.05372-0.10307; Panel 1), or equivalent, as for MASC_MOTOR, MASC_ISR_C, MASC_ISR_1PC, VS_noRT, and CSL ($|t| <= 1.82379$; Panel 1), though differing from that estimated in MASC only for MASC_ISR_C and CSL (estimates: -0.07760 and -0.10816; Panel 2). The latter two, as well as MASC_MOTOR,

MASC_ISR_1PC, and VS_noRT, also best captured FSC readers' overall variability, their standard deviation being non-significantly different from that estimated in FSC readers ($|t| <= 1.28784$; Panel 1). However, the largest mixture component in MASC_ISR_C and CSL accounted on average for only 30-65% and 10-55% of the data respectively, which is much less than for (alternative) MASC (models). On the other hand, VS_RT (as well as VS_RT_GA), which also mostly predicted bimodal distributions, greatly underestimated FSC readers' variability (estimate: -0.27072; Panel 1), and more than MASC, or MASC_noRT and MASC_VISUAL, overestimated it (estimates: 0.08913-0.16335; Panel 1).

Thus, MASC_MOTOR, MASC_ISR_1PC, and VS_noRT, seemed to best replicate FSC readers' variability pattern, followed closely by MASC, MASC_noRT, and MASC_VISUAL. However, when considering the models' performance for both the mean and the SD of within-word landing-positions, it appears that MASC, as well as MASC_noRT and MASC_ISR_1PC, were the best compromise.

**The Launch-Site effect.** As summarized in Supplementary Table 24, the distributions of within-word landing positions, plotted separately for different saccadic launch-site distances to the space in front of the words, were again best fitted with a single mixture component in both MASC and FSC (Supplementary Table 9)[6]: the mean proportion of two mixture components for 4- to 9-letter words in near and far launch-site cases was null for MASC and varied between 0 and 0.02 in FSC readers. A similar pattern was observed for CSL (mean = 0). However, in VS models, the proportion of two mixture components was higher. In VS_RT, it was still no greater than 0.25, and in some conditions only, but in VS_RT_GA, it increased with word length in near launch-site cases, getting as high as 0.37 and 0.55 in 7- and 9-letter words respectively; in those instances, the largest mixture component accounted on average for 87% and 79% of the data. Thus, atypical, bimodal, distributions were more frequent in VS models, but not as much as when within-word landing-position distributions were plotted across launch-site distances (Supplementary Table 22).

**Supplementary Table 24 | GMM-estimated shape of within-word landing-site distributions by word length and launch site in (comparison) models and FSC readers.**

| LS EFFECT: 2 MODES | 4-lw Far | 4-lw Near | 5-lw Far | 5-lw Near | 6-lw Far | 6-lw Near | 7-lw Far | 7-lw Near | 8-lw Far | 8-lw Near | 9-lw Far | 9-lw Near |
|---|---|---|---|---|---|---|---|---|---|---|---|---|
| **FSC** | 0.00 | 0.00 | 0.00 | 0.01 | 0.01 | 0.00 | 0.00 | 0.01 | 0.00 | 0.00 | 0.02 | 0.00 |
| **MASC** | 0.00 | 0.00 | 0.00 | 0.00 | 0.00 | 0.00 | 0.00 | 0.00 | 0.00 | 0.00 | 0.00 | 0.00 |
| **MASC_noRT** | 0.00 | 0.00 | 0.00 | 0.00 | 0.02 | 0.00 | 0.00 | 0.00 | 0.01 | 0.00 | 0.00 | 0.00 |
| **MASC_VISUAL** | 0.00 | 0.00 | 0.00 | 0.00 | 0.02 | 0.00 | 0.00 | 0.00 | 0.00 | 0.00 | 0.00 | 0.00 |
| **MASC_MOTOR** | 0.00 | 0.00 | 0.00 | 0.00 | 0.00 | 0.00 | 0.00 | 0.00 | 0.00 | 0.00 | 0.00 | 0.00 |
| **MASC_ISR_C** | 0.00 | 0.00 | 0.00 | 0.00 | 0.00 | 0.00 | 0.00 | 0.02 | 0.00 | 0.00 | 0.00 | 0.00 |
| **MASC_ISR_1PC** | 0.00 | 0.00 | 0.00 | 0.00 | 0.02 | 0.00 | 0.00 | 0.00 | 0.00 | 0.00 | 0.00 | 0.00 |
| **VS_RT** | 0.00 | 0.04 | 0.25 | 0.03 | 0.00 | 0.11 | 0.00 | 0.03 | 0.14 | 0.10 | 0.00 | 0.00 |
| **VS_RT_GA** | 0.00 | 0.00 | 0.00 | 0.05 | 0.00 | 0.15 | 0.00 | 0.37 | 0.00 | 0.20 | 0.10 | 0.55 |
| **CSL** | 0.00 | 0.00 | 0.00 | 0.00 | 0.00 | 0.00 | 0.00 | 0.00 | 0.00 | 0.00 | 0.00 | 0.00 |

Mean proportion of two mixture components (i.e., the maximal number of detected modes), as estimated after fitting GMMs to individual landing-site distributions partitioned by word length and saccadic launch-site distance to the space in front of the words (binned in two-letter intervals), separately for each data set except VS_noRT because of a too low *n* when data were split by word length and launch site (2693 across subjects, and only 13 after removing instances where there were less than 10 observations per subject, word length, and launch-site bin). To make the tables more digest, means were computed respectively across near and far launch-site distances separately for 4- to 9-letter words (i.e., below and above the median of launch sites, that is -5 letters in 4- to 6-letter words, -4 letters in 7-letter words, and -3 letters in 8- to 9-letter words). These analyses relied on a total of 18922, 10203, 10050, 10050, 11259, 7707, 10334, 4141, 11244, 12654 cases across subjects in FSC, MASC, MASC_noRT, MASC_VISUAL, MASC_MOTOR, MASC_ISR_C, MASC_ISR_1PC, VS_RT, VS_RT_GA, and CSL respectively.

The fixed effects of LMMs for the estimated mean of the largest mixture component are presented in the left panels of Supplementary Table 25. These first indicate that both FSC readers and MASC initially fixated slightly to the left of the center of 6-letter words when saccades were launched from about 4 letters from the space in front of the words (estimates: -0.45537 and -0.36921; Panels 1-2). This leftward fixation bias increased with increasing word length (estimates: -0.20398 and -0.17270; Panels 1-2), but also as saccades were launched from further away (estimates: 0.37336 and 0.62848; Panels 1-2)[6]. Only the effect of launch-site distance differed between the two data sets; the estimated slope was steeper for MASC (estimate: 0.25512; Panel 1), and even more so as words were longer, as suggested by the significant interaction between word length, launch-site distance, and MASC (estimate: 0.05859; Panel 1; see also Supplementary Table 10).

Alternative MASC models also predicted word-length and launch-site effects (Panels 3-7), and as MASC, they all overestimated the slope of the linear relationship between launch-site distance and mean landing site (estimates: 0.19403-0.33904; Panel 1), though slightly less for MASC_noRT (estimate: -0.06109; Panel 2), and slightly more for MASC_VISUAL (estimate: 0.08392; other t's <= 0.91560; Panel 2). However, whereas MASC_MOTOR overestimated the effect of word length (estimate: -0.08645; Panel 1), MASC_noRT and MASC_ISR_1PC underestimated it (estimates: 0.16926 and 0.04625; Panel 1). Moreover, both MASC_noRT and MASC_VISUAL landed on average much further away from the beginning of (6-letter) words than FSC readers (estimates: 0.52023 and 0.66527; Panel 1) and MASC (estimates: 0.43406 and 0.57911; Panel 2), while MASC_MOTOR and MASC_ISR_C showed an even greater leftward fixation bias than FSC readers (estimates: -0.32624 and -0.62005; Panel 1) and MASC (estimates: -0.41240 and -0.70621; Panel 2). Thus, among MASC models, MASC best approximated FSC readers' distributions of within-word landing sites partitioned by word length and launch site.

MASC also beat CSL. This model predicted a smaller leftward fixation bias in 6-letter words than FSC readers (estimate: 0.16490; Panel 1) and much greater effects of word length and launch-site distance (Panel 10) than either FSC readers (estimates: -0.23539 and 0.54889; Panel 2) or MASC (estimates: -0.26667 and 0.29378; Panel 2). For VS models, they beat MASC on the launch-site effect: in VS_RT, this effect was not significantly different from that estimated in FSC readers ($t$ = -1.06281; Panel 1), while it was only mildly smaller in VS_RT_GA (estimate: -0.08395; Panel 1). However, although VS_RT showed the same effect of word length as FSC readers ($t$ = -1.00340; Panel 1) and MASC, VS_RT_GA greatly overestimated it (estimate: -0.50572; Panel 1). Moreover, both VS_RT and VS_RT_GA landed on average much closer to the beginning of (6-letter) words than FSC readers (estimates: -0.53025 and -1.66040; Panel 1) and MASC (estimates: -0.61641 and -1.74656; Panel 2). Recall, in addition, that these predictions from VS models, only applied to the subset of the data accounted for by the largest mixture component, and not all within-word landing positions as for MASC (Supplementary Table 24).

**Supplementary Table 25 | Fixed effects of LMMs for the mean and SD of within-word landing sites by word length and launch site in (comparison) models and FSC readers.**

| | LS EFFECT: MEAN / SD | Estimate | Std. Error | *t* value | Estimate | Std. Error | *t* value |
|---|---|---|---|---|---|---|---|
| **(1)** | **FSC -(Intercept)** | -0.45537 | 0.04147 | -10.98087 | 1.32117 | 0.01388 | 95.18069 |
| | **WL** | -0.20398 | 0.01276 | -15.98160 | 0.09511 | 0.00698 | 13.61963 |
| | **LS** | 0.37336 | 0.01152 | 32.41099 | -0.03409 | 0.00432 | -7.88254 |
| | **MASC** | 0.08616 | 0.07118 | 1.21040 | -0.15367 | 0.02357 | -6.52056 |
| | **MASC_noRT** | 0.52023 | 0.07164 | 7.26203 | -0.14362 | 0.02398 | -5.98783 |
| | **MASC_VISUAL** | 0.66527 | 0.07223 | 9.21050 | -0.26537 | 0.02453 | -10.82012 |
| | **MASC_MOTOR** | -0.32624 | 0.07127 | -4.57781 | -0.17734 | 0.02363 | -7.50372 |
| | **MASC_ISR_C** | -0.62005 | 0.07155 | -8.66557 | -0.07547 | 0.02390 | -3.15752 |
| | **MASC_ISR_1PC** | 0.03224 | 0.07118 | 0.45294 | -0.14940 | 0.02356 | -6.34005 |
| | **VS_RT** | -0.53025 | 0.07687 | -6.89828 | 0.13068 | 0.02825 | 4.62621 |
| | **VS_RT_GA** | -1.66040 | 0.08045 | -20.63995 | 0.21815 | 0.03133 | 6.96381 |
| | **CSL** | 0.16490 | 0.07173 | 2.29891 | -0.73906 | 0.02408 | -30.69585 |
| | **WL:LS** | 0.05421 | 0.00474 | 11.43009 | 0.00641 | 0.00262 | 2.44207 |
| | **WL:MASC** | 0.03128 | 0.02144 | 1.45879 | -0.05053 | 0.01173 | -4.30825 |
| | **WL:MASC_noRT** | 0.16926 | 0.02203 | 7.68371 | -0.04108 | 0.01205 | -3.40768 |
| | **WL:MASC_VISUAL** | 0.02308 | 0.02232 | 1.03443 | -0.03120 | 0.01222 | -2.55270 |
| | **WL:MASC_MOTOR** | -0.08645 | 0.02135 | -4.04838 | -0.04463 | 0.01167 | -3.82265 |
| | **WL:MASC_ISR_C** | -0.03658 | 0.02228 | -1.64196 | -0.06414 | 0.01220 | -5.25893 |
| | **WL:MASC_ISR_1PC** | 0.04625 | 0.02138 | 2.16310 | -0.06155 | 0.01169 | -5.26295 |
| | **WL:VS_RT** | -0.02691 | 0.02682 | -1.00340 | 0.14411 | 0.01466 | 9.82868 |
| | **WL:VS_RT_GA** | -0.50572 | 0.02990 | -16.91341 | 0.14076 | 0.01640 | 8.58096 |
| | **WL:CSL** | -0.23539 | 0.02206 | -10.67049 | -0.07723 | 0.01208 | -6.39232 |
| | **LS:MASC** | 0.25512 | 0.01935 | 13.18246 | 0.02250 | 0.00709 | 3.17334 |
| | **LS:MASC_noRT** | 0.19403 | 0.01984 | 9.77707 | -0.02939 | 0.00749 | -3.92559 |
| | **LS:MASC_VISUAL** | 0.33904 | 0.02068 | 16.39124 | -0.02043 | 0.00816 | -2.50534 |
| | **LS:MASC_MOTOR** | 0.27542 | 0.01956 | 14.08272 | 0.05831 | 0.00725 | 8.04660 |
| | **LS:MASC_ISR_C** | 0.26876 | 0.01985 | 13.53833 | 0.08780 | 0.00750 | 11.70761 |
| | **LS:MASC_ISR_1PC** | 0.25790 | 0.01942 | 13.27890 | 0.02908 | 0.00715 | 4.06678 |
| | **LS:VS_RT** | -0.02536 | 0.02386 | -1.06281 | 0.00491 | 0.01034 | 0.47438 |
| | **LS:VS_RT_GA** | -0.08395 | 0.02626 | -3.19674 | -0.10851 | 0.01208 | -8.97966 |
| | **LS:CSL** | 0.54889 | 0.02009 | 27.31674 | 0.02275 | 0.00770 | 2.95361 |
| | **WL:LS:MASC** | 0.05859 | 0.00764 | 7.67025 | 0.01491 | 0.00424 | 3.51409 |
| | **WL:LS:MASC_noRT** | 0.06249 | 0.00805 | 7.76146 | 0.00255 | 0.00446 | 0.57238 |
| | **WL:LS:MASC_VISUAL** | 0.03880 | 0.00862 | 4.50284 | 0.00474 | 0.00479 | 0.99057 |
| | **WL:LS:MASC_MOTOR** | 0.04288 | 0.00776 | 5.52683 | 0.01739 | 0.00431 | 4.03837 |
| | **WL:LS:MASC_ISR_C** | 0.08607 | 0.00855 | 10.06396 | 0.01927 | 0.00475 | 4.05626 |
| | **WL:LS:MASC_ISR_1PC** | 0.06274 | 0.00774 | 8.10350 | 0.01530 | 0.00430 | 3.55813 |
| | **WL:LS:VS_RT** | -0.00550 | 0.01123 | -0.48949 | -0.01284 | 0.00617 | -2.08216 |
| | **WL:LS:VS_RT_GA** | 0.10635 | 0.01268 | 8.38826 | -0.03059 | 0.00700 | -4.36992 |

| | | | | | | | |
|---|---|---|---|---|---|---|---|
| | **WL:LS:CSL** | -0.02624 | 0.00843 | -3.11321 | 0.00745 | 0.00468 | 1.59119 |
| **(2)** | **MASC -(Intercept)** | -0.36921 | 0.05786 | -6.38146 | 1.16750 | 0.01905 | 61.30085 |
| | **WL** | -0.17270 | 0.01723 | -10.02321 | 0.04458 | 0.00942 | 4.73070 |
| | **LS** | 0.62848 | 0.01555 | 40.41449 | -0.01159 | 0.00562 | -2.06238 |
| | **MASC_noRT** | 0.43406 | 0.08222 | 5.27957 | 0.01005 | 0.02730 | 0.36822 |
| | **MASC_VISUAL** | 0.57911 | 0.08273 | 6.99973 | -0.11170 | 0.02778 | -4.02135 |
| | **MASC_MOTOR** | -0.41240 | 0.08189 | -5.03585 | -0.02367 | 0.02699 | -0.87674 |
| | **MASC_ISR_C** | -0.70621 | 0.08214 | -8.59727 | 0.07820 | 0.02723 | 2.87228 |
| | **MASC_ISR_1PC** | -0.05392 | 0.08182 | -0.65902 | 0.00427 | 0.02693 | 0.15851 |
| | **VS_RT** | -0.61641 | 0.08681 | -7.10057 | 0.28435 | 0.03111 | 9.13929 |
| | **VS_RT_GA** | -1.74656 | 0.09000 | -19.40715 | 0.37182 | 0.03393 | 10.95778 |
| | **CSL** | 0.07874 | 0.08230 | 0.95680 | -0.58539 | 0.02738 | -21.37891 |
| | **WL:LS** | 0.11279 | 0.00599 | 18.83842 | 0.02132 | 0.00334 | 6.39195 |
| | **WL:MASC_noRT** | 0.13798 | 0.02488 | 5.54496 | 0.00945 | 0.01361 | 0.69412 |
| | **WL:MASC_VISUAL** | -0.00820 | 0.02514 | -0.32599 | 0.01933 | 0.01376 | 1.40475 |
| | **WL:MASC_MOTOR** | -0.11773 | 0.02429 | -4.84699 | 0.00590 | 0.01328 | 0.44434 |
| | **WL:MASC_ISR_C** | -0.06786 | 0.02511 | -2.70300 | -0.01362 | 0.01374 | -0.99089 |
| | **WL:MASC_ISR_1PC** | 0.01497 | 0.02431 | 0.61580 | -0.01102 | 0.01330 | -0.82871 |
| | **WL:VS_RT** | -0.05819 | 0.02921 | -1.99220 | 0.19464 | 0.01597 | 12.18848 |
| | **WL:VS_RT_GA** | -0.53700 | 0.03206 | -16.74850 | 0.19129 | 0.01758 | 10.88005 |
| | **WL:CSL** | -0.26667 | 0.02491 | -10.70449 | -0.02671 | 0.01364 | -1.95808 |
| | **LS:MASC_noRT** | -0.06109 | 0.02243 | -2.72411 | -0.05189 | 0.00830 | -6.25021 |
| | **LS:MASC_VISUAL** | 0.08392 | 0.02317 | 3.62172 | -0.04293 | 0.00891 | -4.81856 |
| | **LS:MASC_MOTOR** | 0.02030 | 0.02217 | 0.91560 | 0.03581 | 0.00809 | 4.42885 |
| | **LS:MASC_ISR_C** | 0.01364 | 0.02243 | 0.60812 | 0.06530 | 0.00831 | 7.85514 |
| | **LS:MASC_ISR_1PC** | 0.00279 | 0.02205 | 0.12640 | 0.00658 | 0.00800 | 0.82213 |
| | **LS:VS_RT** | -0.28048 | 0.02605 | -10.76716 | -0.01759 | 0.01094 | -1.60753 |
| | **LS:VS_RT_GA** | -0.33907 | 0.02826 | -11.99658 | -0.13101 | 0.01261 | -10.39325 |
| | **LS:CSL** | 0.29378 | 0.02265 | 12.97199 | 0.00026 | 0.00850 | 0.03005 |
| | **WL:LS:MASC_noRT** | 0.00391 | 0.00884 | 0.44188 | -0.01236 | 0.00491 | -2.51560 |
| | **WL:LS:MASC_VISUAL** | -0.01979 | 0.00936 | -2.11431 | -0.01017 | 0.00521 | -1.95043 |
| | **WL:LS:MASC_MOTOR** | -0.01571 | 0.00858 | -1.83117 | 0.00248 | 0.00477 | 0.51863 |
| | **WL:LS:MASC_ISR_C** | 0.02749 | 0.00930 | 2.95539 | 0.00436 | 0.00518 | 0.84146 |
| | **WL:LS:MASC_ISR_1PC** | 0.00415 | 0.00856 | 0.48459 | 0.00039 | 0.00477 | 0.08199 |
| | **WL:LS:VS_RT** | -0.06408 | 0.01181 | -5.42681 | -0.02776 | 0.00650 | -4.26840 |
| | **WL:LS:VS_RT_GA** | 0.04776 | 0.01319 | 3.61984 | -0.04550 | 0.00730 | -6.23633 |
| | **WL:LS:CSL** | -0.08483 | 0.00919 | -9.23354 | -0.00746 | 0.00512 | -1.45760 |
| **(3)** | **MASC_noRT -(Intercept)** | 0.06485 | 0.05841 | 1.11027 | 1.17756 | 0.01956 | 60.20152 |
| | **WL** | -0.03472 | 0.01795 | -1.93353 | 0.05403 | 0.00983 | 5.49823 |
| | **LS** | 0.56739 | 0.01616 | 35.11235 | -0.06348 | 0.00611 | -10.38554 |
| | **WL:LS** | 0.11670 | 0.00651 | 17.93499 | 0.00896 | 0.00361 | 2.48387 |
| **(4)** | **MASC_VISUAL -(Intercept)** | 0.20990 | 0.05914 | 3.54930 | 1.05580 | 0.02022 | 52.21636 |
| | **WL** | -0.18089 | 0.01831 | -9.88163 | 0.06391 | 0.01003 | 6.37201 |
| | **LS** | 0.71240 | 0.01718 | 41.46787 | -0.05452 | 0.00691 | -7.88463 |
| | **WL:LS** | 0.09301 | 0.00719 | 12.92878 | 0.01115 | 0.00401 | 2.78312 |
| **(5)** | **MASC_MOTOR -(Intercept)** | -0.78161 | 0.05796 | -13.48591 | 1.14384 | 0.01913 | 59.80215 |
| | **WL** | -0.29043 | 0.01712 | -16.96482 | 0.05048 | 0.00936 | 5.39515 |
| | **LS** | 0.64878 | 0.01580 | 41.05036 | 0.02422 | 0.00582 | 4.16577 |
| | **WL:LS** | 0.09709 | 0.00614 | 15.81100 | 0.02380 | 0.00341 | 6.96993 |
| **(6)** | **MASC_ISR_C (Intercept)** | -1.07542 | 0.05831 | -18.44306 | 1.24571 | 0.01946 | 64.02379 |
| | **WL** | -0.24056 | 0.01826 | -13.17455 | 0.03096 | 0.01000 | 3.09606 |
| | **LS** | 0.64212 | 0.01617 | 39.71671 | 0.05372 | 0.00613 | 8.76656 |

| | | | | | | |
|---|---|---|---|---|---|---|
| **WL:LS** | 0.14028 | 0.00712 | 19.70969 | 0.02568 | 0.00396 | 6.48350 |
| **(7) MASC_ISR_1PC -(Intercept)** | -0.42313 | 0.05785 | -7.31389 | 1.17177 | 0.01904 | 61.53444 |
| **WL** | -0.15773 | 0.01716 | -9.19388 | 0.03356 | 0.00938 | 3.57738 |
| **LS** | 0.63127 | 0.01564 | 40.36986 | -0.00501 | 0.00569 | -0.88023 |
| **WL:LS** | 0.11694 | 0.00612 | 19.11129 | 0.02171 | 0.00341 | 6.37038 |
| **(8) VS_RT -(Intercept)** | -0.98562 | 0.06472 | -15.22870 | 1.45186 | 0.02460 | 59.01103 |
| **WL** | -0.23089 | 0.02358 | -9.78984 | 0.23922 | 0.01289 | 18.55425 |
| **LS** | 0.34800 | 0.02090 | 16.65188 | -0.02918 | 0.00939 | -3.10692 |
| **WL:LS** | 0.04871 | 0.01018 | 4.78592 | -0.00644 | 0.00558 | -1.15281 |
| **(9) VS_RT_GA -(Intercept)** | -2.11577 | 0.06893 | -30.69295 | 1.53932 | 0.02808 | 54.81359 |
| **WL** | -0.70970 | 0.02704 | -26.24663 | 0.23586 | 0.01484 | 15.89055 |
| **LS** | 0.28941 | 0.02360 | 12.26224 | -0.14260 | 0.01128 | -12.63738 |
| **WL:LS** | 0.16056 | 0.01176 | 13.65525 | -0.02418 | 0.00649 | -3.72617 |
| **(10) CSL -(Intercept)** | -0.29047 | 0.05853 | -4.96277 | 0.58212 | 0.01967 | 29.59003 |
| **WL** | -0.43937 | 0.01799 | -24.41952 | 0.01787 | 0.00986 | 1.81252 |
| **LS** | 0.92225 | 0.01646 | 56.01751 | -0.01133 | 0.00638 | -1.77740 |
| **WL:LS** | 0.02797 | 0.00697 | 4.01411 | 0.01386 | 0.00388 | 3.57174 |

LMMs were fitted separately to the GMM-estimated mean (left panel) and SD (right panel) of the largest mixture component in individual within-word landing-position distributions split by word length and saccadic launch-site distance to the space in front of the words (binned in two-letter intervals). In both LMMs, the fixed structure included Data Set (10 levels, thus excluding VS_noRT), Word Length (WL; 4-9 letters; reference (mean) value: 6.40 letters), Launch Site (LS; in two-letter bins and ranging between -7 and -1 letters; reference (mean) value: -3.96 letters), and all interactions as predictors. The random structure included a random intercept as well as random effects of WL and LS by subject. Estimates and standard errors are expressed in letters. Panel 1: Fixed-effects for the main LMMs, with Data-Set reference level set to FSC. Panels 2-10: Data-Set reference level set to MASC, MASC_noRT, MASC_VISUAL, MASC_MOTOR, MASC_ISR_C, MASC_ISR_1PC, VS_RT, VS_RT_GA, and CSL respectively. In Panels 3-10, only the intercept, the effects of WL and LS and their interaction are reported.

Additionally, MASC was part of the models that best captured the variability in within-word landing positions. As shown in the right panels of Supplementary Table 25, the estimated SD in FSC readers increased with increasing word length and launch-site distance (estimates: 0.09511 and -0.03409; Panel 1). In MASC models, the SD was a bit smaller (estimates: -0.07547 to -0.26537; Panel 1), but it also significantly increased with word length (Panels 2-7), and only slightly less than for FSC readers (estimates: -0.03120 to -0.06414; Panel 1). Moreover, in MASC, MASC_noRT, and MASC_VISUAL, the variability significantly increased with increasing launch-site distance (Panels 2-4), at only slightly faster or slower rates than in FSC readers (estimates: -0.02939 to 0.02250; Panel 1). In MASC_MOTOR and MASC_ISR_C, to the contrary, the SD significantly decreased with increasing launch-site distance, thus opposite to the effect observed in FSC readers (estimates: 0.02422 and 0.05372; Panels 5-6); in MASC_ISR_1PC, the effect was non-significant ($t$ = -0.88023; Panel 7). MASC, MASC_noRT, and MASC_VISUAL thus outperformed other MASC models for this dependent variable.

These three MASC models also beat CSL and VS models. CSL overall greatly underestimated the variability observed in FSC readers (estimate: -0.73906; Panel 1), while showing neither an effect of word length nor an effect of launch site ($|t| <= -1.81252$; Panel 10). VS models overestimated the variability in within-word landing positions (estimates: 0.13068 and 0.21815; Panel 1), but roughly as little as MASC underestimated it, and they both generated word length and launch-site effects (Panels 8-9). However, they overestimated the effect of word length three times more than MASC underestimated it (estimates: 0.14411 and 0.14076; Panel 1; see also Panel 2). Furthermore, although the launch-site effect for VS_RT did not differ significantly from that observed in FSC readers ($t$ = 0.47438; Panel 1) and MASC ($t$ = -1.60753; Panel 2), this effect for VS_RT_GA was much greater than for FSC readers and MASC (estimates: -0.10851 and -0.13101; Panels 1-2). Thus, when considering the predictions for the shape, the mean, and the SD of within-word landing-position distributions, MASC appeared as the best possible model.

**All landing positions: The Launch-Site effect revisited.**

As noted in Supplementary Methods 1, standard distributions of within-word landing positions are truncated distributions that bias comparison between data sets and conditions[7,10]. We therefore re-analyzed the launch-site effect using the full distributions of saccades' landing positions regardless of word boundaries. The results of GMMs fitted to these overall landing-position distributions, measured relative to the beginning of Word N+1 and partitioned by Word N+1 length and saccadic launch-site distance to the space in front of Word N+1, are summarized in Supplementary Table 26.

**Supplementary Table 26 | GMM-estimated shape of overall landing-site distributions by word length and launch site in (comparison) models and FSC readers.**

| OVERALL LS EFFECT: 2 MODES | 1-lw | 2-lw | 3-lw | 4-lw | 5-lw | 6-lw | 7-lw | 8-lw | 9-lw |
|---|---|---|---|---|---|---|---|---|---|
| **(1) Far Launch Site** | | | | | | | | | |
| **FSC** | 0.02 | 0.12 | 0.16 | 0.06 | 0.00 | 0.02 | 0.05 | 0.01 | 0.04 |
| **MASC** | 0.00 | 0.00 | 0.00 | 0.00 | 0.00 | 0.00 | 0.00 | 0.00 | 0.00 |
| **MASC_noRT** | 0.00 | 0.00 | 0.02 | 0.00 | 0.00 | 0.00 | 0.00 | 0.00 | 0.00 |
| **MASC_VISUAL** | 0.00 | 0.00 | 0.00 | 0.00 | 0.00 | 0.00 | 0.00 | 0.00 | 0.00 |
| **MASC_MOTOR** | 0.00 | 0.00 | 0.00 | 0.00 | 0.00 | 0.00 | 0.00 | 0.00 | 0.00 |
| **MASC_ISR_C** | 0.00 | 0.03 | 0.02 | 0.00 | 0.00 | 0.00 | 0.00 | 0.00 | 0.00 |
| **MASC_ISR_1PC** | 0.00 | 0.00 | 0.00 | 0.00 | 0.00 | 0.00 | 0.00 | 0.05 | 0.00 |
| **VS_RT** | / | 0.18 | 0.41 | 0.10 | 0.05 | 0.23 | 0.08 | 0.14 | 0.33 |
| ***(mean largest k)*** | / | *0.93* | *0.80* | *0.96* | *0.98* | *0.91* | *0.95* | *0.93* | *0.86* |
| **VS_RT_GA** | / | 0.40 | 0.80 | 0.58 | 0.55 | 0.96 | 0.97 | 0.83 | 0.57 |
| ***(mean largest k)*** | / | *0.83* | *0.67* | *0.77* | *0.76* | *0.61* | *0.61* | *0.70* | *0.80* |
| **CSL** | 0.00 | 0.00 | 0.00 | 0.00 | 0.00 | 0.00 | 0.00 | 0.00 | 0.00 |
| **(2) Near Launch Site** | | | | | | | | | |
| **FSC** | 0.00 | 0.04 | 0.00 | 0.01 | 0.00 | 0.02 | 0.05 | 0.00 | 0.00 |
| **MASC** | 0.00 | 0.00 | 0.00 | 0.00 | 0.00 | 0.00 | 0.00 | 0.00 | 0.00 |
| **MASC_noRT** | 0.00 | 0.00 | 0.00 | 0.00 | 0.00 | 0.00 | 0.05 | 0.00 | 0.00 |
| **MASC_VISUAL** | 0.00 | 0.00 | 0.00 | 0.00 | 0.00 | 0.00 | 0.00 | 0.00 | 0.00 |
| **MASC_MOTOR** | 0.00 | 0.00 | 0.00 | 0.00 | 0.00 | 0.00 | 0.00 | 0.00 | 0.00 |
| **MASC_ISR_C** | 0.00 | 0.02 | 0.02 | 0.00 | 0.00 | 0.00 | 0.00 | 0.00 | 0.09 |
| **MASC_ISR_1PC** | 0.00 | 0.00 | 0.00 | 0.00 | 0.00 | 0.00 | 0.00 | 0.00 | 0.00 |
| **VS_RT** | / | 0.87 | 0.48 | 0.10 | 0.10 | 0.00 | 0.00 | 0.00 | 0.05 |
| ***(mean largest k)*** | / | *0.63* | *0.83* | *0.96* | *0.96* | *1.00* | *1.00* | *1.00* | *0.98* |
| **VS_RT_GA** | 0.50 | 0.00 | 0.82 | 0.17 | 0.00 | 0.80 | 0.75 | 0.90 | 0.80 |
| ***(mean largest k)*** | *0.83* | *1.00* | *0.70* | *0.94* | *1.00* | *0.74* | *0.72* | *0.60* | *0.65* |
| **CSL** | 0.00 | 0.00 | 0.00 | 0.00 | 0.00 | 0.00 | 0.00 | 0.00 | 0.00 |

Mean proportion of two mixture components (i.e., the maximal number of detected modes), as estimated after fitting GMMs to individual distributions partitioned by word length and saccadic launch-site distance to the space in front of the words (binned in two-letter intervals), separately for each data set, but excluding VS_noRT due to a comparatively much lower *n* (6715 across subjects, and 4503 after removing instances where there were less than 10 observations per subject, word length, and launch-site bin) and a rather poor fit. To make the tables more digest, means were computed respectively across near and far launch-site distances separately for 1- to 9-letter words (i.e., below and above the median of launch sites, that is -5 for 2- to 4-letter words and -3 for 1-, and 5- to 9-letter words; Panels 1 and 2 respectively). In most data sets and conditions, the mean proportion of cases for the largest mixture component ranged between 0.94 and 1. However, this was much lower in most conditions for VS_RT and even more so for VS_RT_GA. Only for these two data sets, are the mean *k* values for the largest mixture component reported (in italics) in the table. These analyses relied on a total of 35395, 19978, 17537, 19740, 22336, 14767, 19583, 15968, 24943, 19745 cases across subjects in FSC, MASC, MASC_noRT, MASC_VISUAL, MASC_MOTOR, MASC_ISR_C, MASC_ISR_1PC, VS_RT, VS_RT_GA, and CSL respectively.

This table shows that the proportion of two-mixture components was again relatively low in FSC readers (mean: 0-0.016) and MASC (mean: 0), as well as CSL (mean: 0), with the largest mixture component accounting for more than 94% of the data in all three data sets. This confirms the above reported pattern when MASC and FSC data sets were matched for numbers of fixations (Supplementary Table 9). For VS_RT, the proportion of two mixture components was higher, and particularly in far launch-site cases (0.05-0.41), as well as for short 2- and 3-letter words in near launch-site conditions (0.87 and 0.48). However, the data set that contrasted the most from FSC was VS_RT_GA. In that case, the predicted distributions were for the most bimodal: The proportion of two mixture components ranged between 0.40 and 0.97 in far launch-site conditions, and between 0 and 0.90 in near launch-site conditions, with the largest mode accounting on average for only 61-83% and 60-100% of the data respectively.

The fixed effects of LMMs fitted to the mean of the largest mixture component are presented in the left panels of Supplementary Table 27. This again shows great similarity between MASC and FSC readers, and even greater similarity than when only within-word landing positions were selected for analysis (Supplementary Table 25, Supplementary Methods 1). MASC overshot the center of 4-letter words (in near, 4-letter, launch-site instances), only slightly less than FSC readers (estimate: -0.48240; Panel 1), and it landed closer to the beginning of longer and more eccentric words (estimates: -0.32622 and 0.84619; Panel 2), as FSC readers (estimates: -0.61701 and 0.81771; Panel 1). The effect of word length was smaller for MASC compared to FSC readers (estimate: 0.29079; Panel 1). However, the effect of launch-site distance was not significantly different for short, 4-letter, words ($t$ = 0.86540; Panel 1) and the slope difference in longer words remained very small: this increased by only about 0.05 letters with every one-letter increment of word length, as suggested by the significant three-way interaction between word length, launch-site distance, and MASC (Panel 1; Extended Data Fig. 5e). These small differences came from the launch-site effect being slightly greater in longer words for MASC (estimate: -0.03606; Panel 2), but not for FSC readers ($t$ = -1.75384; Panel 1). They likely resulted from the fact that MASC, unlike FSC readers, did not benefit from mild top-down, language-related, modulations of saccade amplitude (Extended Data Fig. 2b).

Alternative MASC models also predicted both word-length and launch-site effects (Panels 3-7; Extended Data Fig. 5e,k, 6e, 7e). In all these models, the effect of word length was smaller than for FSC readers (estimates: 0.12541-0.35515; Panel 1), and it did not differ from the effect predicted by MASC except for MASC_ISR_C (estimate: -0.16537, $t$ = -4.57498; other |$t$s| <= 1.56502; Panel 2), which less largely underestimated the effect observed in FSC readers. Likewise, the launch-site effect did not differ from that estimated for FSC readers and MASC, except for MASC_ISR_C, which overestimated it (estimates: 0.24547 and 0.21698 respectively; Panels 1 and 2). Moreover, for all models, this effect became slightly greater with increasing word length, as suggested by the (marginally) significant interactions between word length and launch-site distance ($t$ >= 1.91165; Panels 3-7), leading to slightly greater slope differences between the models and FSC readers for longer words (estimates: 0.03511-0.09746; Panel 1), just as for MASC; only MASC_ISR_C again differed from MASC (estimate: 0.04952, $t$ = 3.23581; other |$t$s| <= -0.75924; Panel 2). In addition, MASC_ISR_C, and to some extent MASC_MOTOR, did not land far enough, initially fixating near the center of short (4-letter) words, rather than overshooting it (Panels 1, 5-6). Thus, for this dependent variable, the prediction level was comparable across all MASC models, except MASC_ISR_C and MASC_MOTOR, whose predictions were not as good.

CSL behaved pretty much like MASC, but still did not beat it for the launch-site effect. It underestimated FSC readers' mean landing positions (estimate: -0.36053; Panel 1) by about the same amount as MASC ($t$ = 0.64038; Panel 2), and, as MASC, it predicted effects of word length and launch-site distance (Panel 10; Fig. 3e). Its word-length effect more closely matched that observed for FSC readers (estimate: 0.12756; Panel 1; see also Panel 2). However, its launch-site effect (for 4-letter words) was greater than for FSC readers (estimate: 0.15718; Panel 1) and MASC (estimate: 0.12869; Panel 2), regardless of word length, as suggested by the non-significant interaction between and word length, launch-site distance, and CSL (*t* = 0.85213; Panel 1).

**Supplementary Table 27 | Fixed effects of LMMs for the mean and SD of all landing sites by word length and launch site in (comparison) models and FSC readers.**

| OVERALL LS EFFECT: MEAN / SD | Estimate | Std. Error | *t* value | Estimate | Std. Error | *t* value |
|---|---|---|---|---|---|---|
| (1) FSC -(Intercept) | 1.50520 | 0.09333 | 16.12734 | 2.31758 | 0.03466 | 66.85979 |
| WL | -0.61701 | 0.01583 | -38.97542 | -0.07715 | 0.00908 | -8.50103 |
| LS | 0.81771 | 0.01730 | 47.27196 | -0.04246 | 0.00967 | -4.39300 |
| MASC | -0.48240 | 0.16406 | -2.94048 | -0.64221 | 0.06290 | -10.21023 |
| MASC_noRT | 0.25847 | 0.16624 | 1.55479 | -0.41357 | 0.06530 | -6.33350 |
| MASC_VISUAL | 0.30427 | 0.16357 | 1.86025 | -0.83503 | 0.06234 | -13.39576 |
| MASC_MOTOR | -1.09194 | 0.16431 | -6.64557 | -0.79452 | 0.06321 | -12.57009 |
| MASC_ISR_C | -1.55095 | 0.16202 | -9.57277 | -0.41949 | 0.06056 | -6.92633 |
| MASC_ISR_1PC | -0.54047 | 0.16392 | -3.29711 | -0.62013 | 0.06276 | -9.88033 |
| VS_RT | 1.49572 | 0.16393 | 9.12415 | 1.68877 | 0.06273 | 26.91914 |
| VS_RT_GA | -3.01584 | 0.16079 | -18.75594 | -0.25877 | 0.05916 | -4.37395 |
| CSL | -0.36053 | 0.16348 | -2.20537 | -1.66319 | 0.06226 | -26.71532 |
| WL:LS | -0.01188 | 0.00677 | -1.75384 | 0.01541 | 0.00441 | 3.49730 |
| WL:MASC | 0.29079 | 0.03103 | 9.37032 | 0.00290 | 0.01850 | 0.15689 |
| WL:MASC_noRT | 0.35515 | 0.03506 | 10.12894 | -0.03169 | 0.02134 | -1.48508 |
| WL:MASC_VISUAL | 0.27442 | 0.03057 | 8.97725 | 0.04130 | 0.01815 | 2.27630 |
| WL:MASC_MOTOR | 0.26981 | 0.03135 | 8.60619 | 0.03942 | 0.01875 | 2.10272 |
| WL:MASC_ISR_C | 0.12541 | 0.02907 | 4.31466 | 0.01292 | 0.01705 | 0.75754 |
| WL:MASC_ISR_1PC | 0.27879 | 0.03156 | 8.83320 | 0.00978 | 0.01890 | 0.51765 |
| WL:VS_RT | 0.18032 | 0.03153 | 5.71960 | 0.18111 | 0.01882 | 9.62230 |
| WL:VS_RT_GA | 0.13326 | 0.02644 | 5.03971 | 0.00586 | 0.01510 | 0.38794 |
| WL:CSL | 0.12756 | 0.03104 | 4.10890 | 0.07332 | 0.01851 | 3.96000 |
| LS:MASC | 0.02849 | 0.03292 | 0.86540 | 0.11417 | 0.01909 | 5.98067 |
| LS:MASC_noRT | -0.01446 | 0.03525 | -0.41021 | 0.10491 | 0.02080 | 5.04259 |
| LS:MASC_VISUAL | 0.06334 | 0.03283 | 1.92916 | 0.09182 | 0.01903 | 4.82572 |
| LS:MASC_MOTOR | 0.04449 | 0.03362 | 1.32344 | 0.10558 | 0.01964 | 5.37522 |
| LS:MASC_ISR_C | 0.24547 | 0.02936 | 8.35941 | 0.08298 | 0.01639 | 5.06424 |
| LS:MASC_ISR_1PC | 0.05856 | 0.03287 | 1.78142 | 0.10459 | 0.01908 | 5.48135 |
| LS:VS_RT | 0.16334 | 0.03245 | 5.03400 | -0.02604 | 0.01872 | -1.39104 |
| LS:VS_RT_GA | 0.06796 | 0.02819 | 2.41041 | 0.00858 | 0.01552 | 0.55326 |
| LS:CSL | 0.15718 | 0.03270 | 4.80631 | 0.05173 | 0.01895 | 2.73007 |
| WL:LS:MASC | 0.04794 | 0.01355 | 3.53733 | -0.01231 | 0.00888 | -1.38756 |
| WL:LS:MASC_noRT | 0.04205 | 0.01509 | 2.78572 | -0.00862 | 0.00989 | -0.87186 |
| WL:LS:MASC_VISUAL | 0.05421 | 0.01327 | 4.08591 | -0.02269 | 0.00868 | -2.61433 |
| WL:LS:MASC_MOTOR | 0.03511 | 0.01391 | 2.52357 | -0.01262 | 0.00913 | -1.38308 |
| WL:LS:MASC_ISR_C | 0.09746 | 0.01193 | 8.17060 | -0.01379 | 0.00780 | -1.76827 |
| WL:LS:MASC_ISR_1PC | 0.05587 | 0.01398 | 3.99760 | -0.01337 | 0.00917 | -1.45838 |
| WL:LS:VS_RT | 0.06022 | 0.01332 | 4.52088 | -0.07622 | 0.00872 | -8.74045 |
| WL:LS:VS_RT_GA | 0.08934 | 0.01107 | 8.07199 | -0.02073 | 0.00725 | -2.85972 |
| WL:LS:CSL | 0.01146 | 0.01345 | 0.85213 | -0.01572 | 0.00881 | -1.78448 |
| (2) MASC -(Intercept) | 1.02279 | 0.13492 | 7.58073 | 1.67537 | 0.05249 | 31.92085 |
| WL | -0.32622 | 0.02669 | -12.22209 | -0.07425 | 0.01612 | -4.60678 |
| LS | 0.84619 | 0.02801 | 30.21359 | 0.07171 | 0.01646 | 4.35597 |

| | | | | | | | |
|---|---|---|---|---|---|---|---|
| | MASC_noRT | 0.74087 | 0.19269 | 3.84491 | 0.22864 | 0.07627 | 2.99779 |
| | MASC_VISUAL | 0.78668 | 0.19038 | 4.13203 | -0.19282 | 0.07375 | -2.61453 |
| | MASC_MOTOR | -0.60954 | 0.19103 | -3.19087 | -0.15231 | 0.07449 | -2.04479 |
| | MASC_ISR_C | -1.06855 | 0.18906 | -5.65203 | 0.22273 | 0.07226 | 3.08238 |
| | MASC_ISR_1PC | -0.05807 | 0.19069 | -0.30453 | 0.02208 | 0.07411 | 0.29799 |
| | VS_RT | 1.97813 | 0.19070 | 10.37309 | 2.33098 | 0.07409 | 31.46293 |
| | VS_RT_GA | -2.53344 | 0.18801 | -13.47511 | 0.38344 | 0.07109 | 5.39414 |
| | CSL | 0.12187 | 0.19031 | 0.64038 | -1.02098 | 0.07368 | -13.85668 |
| | WL:LS | 0.03606 | 0.01174 | 3.07197 | 0.00309 | 0.00770 | 0.40135 |
| | WL:MASC_noRT | 0.06436 | 0.04112 | 1.56502 | -0.03459 | 0.02515 | -1.37517 |
| | WL:MASC_VISUAL | -0.01637 | 0.03737 | -0.43803 | 0.03840 | 0.02251 | 1.70610 |
| | WL:MASC_MOTOR | -0.02098 | 0.03801 | -0.55198 | 0.03652 | 0.02300 | 1.58798 |
| | WL:MASC_ISR_C | -0.16537 | 0.03615 | -4.57498 | 0.01002 | 0.02164 | 0.46290 |
| | WL:MASC_ISR_1PC | -0.01200 | 0.03818 | -0.31416 | 0.00688 | 0.02312 | 0.29759 |
| | WL:VS_RT | -0.11046 | 0.03815 | -2.89511 | 0.17821 | 0.02306 | 7.72871 |
| | WL:VS_RT_GA | -0.15753 | 0.03407 | -4.62310 | 0.00296 | 0.02014 | 0.14683 |
| | WL:CSL | -0.16323 | 0.03776 | -4.32333 | 0.07042 | 0.02281 | 3.08738 |
| | LS:MASC_noRT | -0.04295 | 0.04157 | -1.03324 | -0.00926 | 0.02471 | -0.37469 |
| | LS:MASC_VISUAL | 0.03485 | 0.03954 | 0.88150 | -0.02235 | 0.02323 | -0.96202 |
| | LS:MASC_MOTOR | 0.01600 | 0.04019 | 0.39816 | -0.00859 | 0.02373 | -0.36188 |
| | LS:MASC_ISR_C | 0.21698 | 0.03671 | 5.91108 | -0.03119 | 0.02112 | -1.47658 |
| | LS:MASC_ISR_1PC | 0.03007 | 0.03957 | 0.75994 | -0.00958 | 0.02327 | -0.41163 |
| | LS:VS_RT | 0.13485 | 0.03922 | 3.43860 | -0.14021 | 0.02298 | -6.10148 |
| | LS:VS_RT_GA | 0.03947 | 0.03578 | 1.10321 | -0.10558 | 0.02045 | -5.16243 |
| | LS:CSL | 0.12869 | 0.03943 | 3.26386 | -0.06244 | 0.02316 | -2.69574 |
| | WL:LS:MASC_noRT | -0.00589 | 0.01788 | -0.32910 | 0.00369 | 0.01174 | 0.31439 |
| | WL:LS:MASC_VISUAL | 0.00627 | 0.01637 | 0.38327 | -0.01038 | 0.01074 | -0.96656 |
| | WL:LS:MASC_MOTOR | -0.01283 | 0.01690 | -0.75924 | -0.00031 | 0.01110 | -0.02759 |
| | WL:LS:MASC_ISR_C | 0.04952 | 0.01530 | 3.23581 | -0.00148 | 0.01004 | -0.14705 |
| | WL:LS:MASC_ISR_1PC | 0.00793 | 0.01695 | 0.46801 | -0.00105 | 0.01113 | -0.09444 |
| | WL:LS:VS_RT | 0.01228 | 0.01641 | 0.74850 | -0.06391 | 0.01077 | -5.93360 |
| | WL:LS:VS_RT_GA | 0.04140 | 0.01464 | 2.82736 | -0.00841 | 0.00962 | -0.87457 |
| | WL:LS:CSL | -0.03647 | 0.01652 | -2.20808 | -0.00341 | 0.01084 | -0.31422 |
| **(3)** | **MASC_noRT -(Intercept)** | 1.76367 | 0.13757 | 12.82001 | 1.90401 | 0.05534 | 34.40642 |
| | **WL** | -0.26186 | 0.03129 | -8.37011 | -0.10884 | 0.01931 | -5.63566 |
| | **LS** | 0.80325 | 0.03071 | 26.15372 | 0.06245 | 0.01842 | 3.38965 |
| | **WL:LS** | 0.03017 | 0.01349 | 2.23664 | 0.00678 | 0.00886 | 0.76584 |
| **(4)** | **MASC_VISUAL -(Intercept)** | 1.80947 | 0.13432 | 13.47092 | 1.48256 | 0.05181 | 28.61598 |
| | **WL** | -0.34259 | 0.02615 | -13.10107 | -0.03585 | 0.01571 | -2.28125 |
| | **LS** | 0.88104 | 0.02791 | 31.57170 | 0.04936 | 0.01639 | 3.01161 |
| | **WL:LS** | 0.04233 | 0.01141 | 3.71042 | -0.00729 | 0.00748 | -0.97421 |
| **(5)** | **MASC_MOTOR -(Intercept)** | 0.41326 | 0.13523 | 3.05595 | 1.52306 | 0.05285 | 28.81608 |
| | **WL** | -0.34720 | 0.02706 | -12.83093 | -0.03773 | 0.01640 | -2.29995 |
| | **LS** | 0.86220 | 0.02882 | 29.91181 | 0.06312 | 0.01710 | 3.69127 |
| | **WL:LS** | 0.02323 | 0.01215 | 1.91165 | 0.00279 | 0.00799 | 0.34862 |
| **(6)** | **MASC_ISR_C (Intercept)** | -0.04576 | 0.13243 | -0.34550 | 1.89810 | 0.04966 | 38.21938 |
| | **WL** | -0.49160 | 0.02438 | -20.16638 | -0.06423 | 0.01444 | -4.44920 |
| | **LS** | 1.06317 | 0.02373 | 44.80560 | 0.04052 | 0.01323 | 3.06242 |
| | **WL:LS** | 0.08558 | 0.00982 | 8.71550 | 0.00162 | 0.00644 | 0.25110 |
| **(7)** | **MASC_ISR_1PC -(Intercept)** | 0.96472 | 0.13476 | 7.15886 | 1.69746 | 0.05232 | 32.44166 |
| | **WL** | -0.33822 | 0.02730 | -12.38691 | -0.06737 | 0.01657 | -4.06457 |
| | **LS** | 0.87626 | 0.02795 | 31.34942 | 0.06213 | 0.01645 | 3.77634 |

| | | | | | | |
|---|---|---|---|---|---|---|
| **WL:LS** | 0.04399 | 0.01223 | 3.59842 | 0.00204 | 0.00804 | 0.25393 |
| **(8) VS_RT -(Intercept)** | 3.00092 | 0.13477 | 22.26736 | 4.00635 | 0.05229 | 76.61981 |
| **WL** | -0.43668 | 0.02726 | -16.01644 | 0.10396 | 0.01649 | 6.30461 |
| **LS** | 0.98105 | 0.02745 | 35.73640 | -0.06851 | 0.01603 | -4.27248 |
| **WL:LS** | 0.04834 | 0.01147 | 4.21448 | -0.06082 | 0.00753 | -8.08060 |
| **(9) VS_RT_GA -(Intercept)** | -1.51064 | 0.13093 | -11.53743 | 2.05882 | 0.04794 | 42.94388 |
| **WL** | -0.48375 | 0.02118 | -22.84007 | -0.07129 | 0.01207 | -5.90658 |
| **LS** | 0.88566 | 0.02226 | 39.78149 | -0.03388 | 0.01214 | -2.79120 |
| **WL:LS** | 0.07746 | 0.00875 | 8.84817 | -0.00532 | 0.00575 | -0.92414 |
| **(10) CSL -(Intercept)** | 1.14466 | 0.13422 | 8.52839 | 0.65439 | 0.05171 | 12.65416 |
| **WL** | -0.48945 | 0.02670 | -18.32894 | -0.00383 | 0.01614 | -0.23743 |
| **LS** | 0.97488 | 0.02775 | 35.12739 | 0.00926 | 0.01630 | 0.56847 |
| **WL:LS** | -0.00041 | 0.01162 | -0.03563 | -0.00031 | 0.00763 | -0.04128 |

LMMs were fitted separately to the GMM-estimated mean (left panel) and SD (right panel) of the largest mixture component in individual overall landing-position distributions split by word length and saccades' launch-site distance to the space in front of the words (binned in two-letter intervals). In both LMMs, the fixed structure included Data Set (10 levels, excluding VS_noRT), Word Length (WL; 1-10 letters; reference (mean) value: 4.51 letters), Launch Site (LS; in two-letter bins and ranging between -9 and -1 letters; reference (mean) value: -3.60 letters), and all interactions as predictors. The random structure included a random intercept as well as random effects of WL and LS by subject. Estimates and standard errors are expressed in letters. Panel 1: Fixed-effects for the main LMMs, with Data-Set reference level set to FSC. Panels 2-10: Data-Set reference level set to MASC, MASC_noRT, MASC_VISUAL, MASC_MOTOR, MASC_ISR_C, MASC_ISR_1PC, VS_RT, VS_RT_GA, and CSL respectively. In Panels 3-10, only the intercept, the effects of WL and LS, and the interaction were reported.

MASC (models) however clearly beat VS models. VS_RT overshot the center of 4-letter words by about 1.5 letters more than FSC readers (Panels 1,8), while VS_RT_GA undershot it by about the same amount as FSC readers overshot it (estimate: -1.51064; Panel 9). Moreover, although both models predicted an effect of word length (Panels 8-9), that less largely underestimated the effect observed in FSC readers (estimates: 0.18032 and 0.13326; Panel 1; see also Panel 2), they predicted an effect of launch-site distance that was much greater than for FSC readers (estimates: 0.16334 and 0.06796; Panel 1), and even more so as word length increased (estimates: 0.06022 and 0.08934; Panel 1; Fig. 3k).

Finally, as shown in the right panels of Supplementary Table 27, although MASC did not perfectly predict the spread in saccades' landing positions, it still outperformed CSL and VS models. MASC, and other MASC models, underestimated FSC readers' variability (estimates: -0.64221 and -0.41357 to -0.83503; Panel 1), but not as much as CSL (estimate: -1.66319; Panel 1), and much less than VS_RT overestimated it (estimate: 1.68877; Panel 1). In addition, MASC predicted a significant decrease in the spread of landing positions with increasing word length (estimate: -0.07425; Panel 2), that was not significantly different from that estimated in FSC readers ($t$ = 0.15689; Panel 1). Alternative MASC models and CSL also predicted a significant effect of word length (Panels 3-7, 10), but for MASC_VISUAL, MASC_MOTOR, and CSL the effect was significantly smaller than for FSC readers (estimates: 0.04130, 0.03942, and 0.07332; other |*ts*| <= -1.48508; Panel 1). For VS_RT, the effect of word length (Panel 8) was opposite to that observed in FSC readers (estimate: 0.18111; Panel 1). The only weakness in MASC models is that they predicted a decrease, and not an increase (as in FSC readers), in the spread of landing positions with increasing launch-site distance (0.04052-0.07171 vs. -0.04246; Panels 2-7, 1); the interaction between launch site and each of these data sets was significant (estimates: 0.08298-0.11417; Panel 1). For CSL, the effect of launch site was non-significant ($t$ = 0.56847; Panel 10) and significantly different from the effect in FSC readers (and 0.05173; Panel 1). VS_RT did predict an increase in variability with increasing launch-site distance (Panel 8), that was not significantly different from that estimated in FSC readers ($t$ = -1.39104; Panel 1) for 4-letter words, but that still differed in longer words, as suggested by the significant three-way interaction (estimate: -0.07622; Panel 1). Only VS_RT_GA seemed to do slightly better than MASC (models). It underestimated FSC readers' variability (estimate: -0.25877; Panel 1) less than MASC (estimate: 0.38344; Panel 2), and it predicted, just as MASC, a similar effect of word length as FSC readers ($t$ = 0.38794; Panel 1). Moreover, its predicted effect of launch site was comparable to that estimated in FSC readers for 4-letter words ($t$ = 0.55326; Panel 1). However, this did not hold for longer words, as suggested by the significant three-way interaction (estimate: -0.02073; Panel 1). The main problem though was that these predictions only applied to the very-few data belonging to the largest mixture component, that is as little as 74% of the data on average, and even less (61%) for some combinations of word length and launch site (Supplementary Table 26).

Thus, when considering all three indices (shape, mean and SD), it appears that MASC, as well as MASC_noRT and MASC_ISR_1PC, provided the best approximation of FSC readers' overall landing-position distributions[7] and launch-site effect.

**Within-word refixation behavior: The OVP effect.**

As reported in the above comparisons between MASC and FSC data sets, when matched for numbers of fixations, MASC's within-word refixation behavior nearly perfectly matched that observed in FSC readers when considering only the refixations that were initiated from the first halves of words (Supplementary Table 13, Fig.2h). Likewise, as shown in Supplementary Table 28, the likelihood of within-word refixations in FSC readers and MASC (of about 19% and 15% respectively when words were about 7 letters long; Panel 1) increased with increasing word length (logit: 0.25616 and 0.27659; Panels 1-2), and also as the initial fixation deviated from the words' center (logit: -0.58403 and -0.63831; Panels 1-2)[11-14]. Neither the effect of word length nor the effect of initial landing position differed significantly between the two data sets ($p$ = 0.52056 and $p$ = 0.27775; Panel 1). The only difference was that MASC's left-OVP effect did not significantly vary with word length ($p$ = 0.37716; Panel 2), while this effect in FSC readers tended to become weaker for longer words (logit: 0.02439, $p$ = 0.07656; Panel 1). This resulted in a slightly steeper slope for MASC in longer words, as suggested by the marginally significant interaction between word length, initial landing position and MASC (Extended Data Fig. 3).

Almost none of the alternative models reached that level of prediction. MASC models all predicted an increase in the likelihood of within-word refixations in longer words, and as saccades landed closer to the words' beginning (Panels 3-7; Extended Data Fig. 5l, 6f, 7f). The effect of word length did not differ from that observed in FSC readers, except for MASC_noRT which tended to mildly underestimate that effect (logit: -0.06542, $p = 0.06007$; other $p\text{'s} >= 0.13591$). However, the predicted left wing of the OVP effect was either too steep, as in MASC_VISUAL, MASC_MOTOR, and MASC_ISR_1PC (logit: -0.11780 to -0.35304; Panel 1), and steeper than for MASC at least for MASC_VISUAL and MASC_MOTOR (logit: -0.29883 and 0.11746; Panel 1), or way too weak, as in MASC_ISR_C (logit: 0.40556; Panel 1; see also Panel 2). Only the slope for MASC_noRT did not differ from that observed in FSC readers ($p = 0.93074$; Panel 2); it also did not differ from that estimated for MASC, just as the slope for MASC_ISR_1PC ($p >= 0.28846$; Panel 2). This suggests that among MASC models, both MASC and MASC_noRT, followed by MASC_ISR_1PC, best estimated the left wing of the Refixation-OVP effect. Interestingly, MASC_ISR_C generated some refixations from the right halves of words (Extended Data Fig. 6f), thus indicating that these refixations also do not exclusively reflect ongoing language processes.

VS models predicted word-length and initial-landing-position effects (Fig. 3l). However, for VS_RT and VS_RT_GA, the effect of word length was much weaker than for FSC readers (logit: -0.16921 and -0.22553; Panel 1) and MASC (logit: -0.18964 and -0.24599; Panel 2). Moreover, the effect of initial landing position was either weaker or stronger than for FSC readers (logit: 0.10108 and -0.08796; Panel 1) and MASC (logit: 0.15539, $p = 0.00881$, but -0.03385, $p = 0.52816$; Panel 1), and even more/less so as word length increased (logit: 0.20093 and 0.34541 – Panel 1; logit: 0.24417 and 0.38868 –Panel 2). For VS_noRT, the effect of word length differed only marginally from that observed in FSC readers (logit: -0.11842, $p = 0.09462$; Panel 1) and MASC (logit: -0.13859, $p = 0.06103$; Panel 2). Moreover, the effect of initial landing position was comparable to that estimated for FSC readers and MASC in the case of 7-letter words ($p = 0.53050$ and $p = 0.23450$; Panels 1-2), but it differed drastically for shorter/longer words (logit: 0.26818 and 0.31152; Panels 1-2). Thus, MASC, as well as MASC_noRT, outperformed all other models for the left wing of the Refixation-OVP effect.

**Supplementary Table 28 | Fixed effects of GLMMs for the probability of within-word refixations by initial landing position and word length in (comparison) models and FSC readers**.

| Refixation-OVP EFFECT | Estimate | Std. Error | z value | Pr(>\|z\|) | Proportion |
|---|---|---|---|---|---|
| **(1) FSC -(Intercept)** | -1.43949 | 0.06648 | -21.65372 | 0.00000 | 0.19162 |
| **WL** | 0.25616 | 0.01707 | 15.00264 | 0.00000 | |
| **ILP** | -0.58403 | 0.02839 | -20.57128 | 0.00000 | |
| **MASC** | -0.27610 | 0.11521 | -2.39645 | 0.01655 | 0.15244 |
| **MASC_noRT** | -0.52103 | 0.11690 | -4.45724 | 0.00001 | 0.12341 |
| **MASC_VISUAL** | -2.05164 | 0.13846 | -14.81731 | 0.00000 | 0.02956 |
| **MASC_MOTOR** | 0.01298 | 0.11389 | 0.11395 | 0.90928 | 0.19364 |
| **MASC_ISR_C** | -0.24832 | 0.11459 | -2.16704 | 0.03023 | 0.15606 |
| **MASC_ISR_1PC** | -0.44762 | 0.11616 | -3.85360 | 0.00012 | 0.13157 |
| **VS_noRT** | -1.13130 | 0.13934 | -8.11909 | 0.00000 | 0.07104 |
| **VS_RT** | -0.03741 | 0.11472 | -0.32612 | 0.74434 | 0.18589 |
| **VS_RT_GA** | 1.55785 | 0.11217 | 13.88850 | 0.00000 | 0.52955 |
| **WL:ILP** | 0.02439 | 0.01377 | 1.77103 | 0.07656 | |
| **WL:MASC** | 0.02069 | 0.03220 | 0.64248 | 0.52056 | |
| **WL:MASC_noRT** | -0.06542 | 0.03479 | -1.88026 | 0.06007 | |
| **WL:MASC_VISUAL** | -0.01851 | 0.06356 | -0.29121 | 0.77089 | |
| **WL:MASC_MOTOR** | -0.00173 | 0.02978 | -0.05818 | 0.95360 | |
| **WL:MASC_ISR_C** | -0.04633 | 0.03107 | -1.49119 | 0.13591 | |
| **WL:MASC_ISR_1PC** | 0.03333 | 0.03378 | 0.98673 | 0.32377 | |
| **WL:VS_noRT** | -0.11842 | 0.07085 | -1.67151 | 0.09462 | |
| **WL:VS_RT** | -0.16921 | 0.03338 | -5.06860 | 0.00000 | |
| **WL:VS_RT_GA** | -0.22553 | 0.02747 | -8.21154 | 0.00000 | |
| **ILP:MASC** | -0.05452 | 0.05023 | -1.08539 | 0.27775 | |
| **ILP:MASC_noRT** | -0.00455 | 0.05237 | -0.08691 | 0.93074 | |
| **ILP:MASC_VISUAL** | -0.35304 | 0.07513 | -4.69884 | 0.00000 | |
| **ILP:MASC_MOTOR** | -0.17177 | 0.04848 | -3.54341 | 0.00039 | |
| **ILP:MASC_ISR_C** | 0.40556 | 0.04834 | 8.38938 | 0.00000 | |
| **ILP:MASC_ISR_1PC** | -0.11780 | 0.05168 | -2.27938 | 0.02264 | |
| **ILP:VS_noRT** | 0.05378 | 0.08574 | 0.62724 | 0.53050 | |
| **ILP:VS_RT** | 0.10108 | 0.05100 | 1.98189 | 0.04749 | |
| **ILP:VS_RT_GA** | -0.08796 | 0.04430 | -1.98577 | 0.04706 | |
| **WL:ILP:MASC** | -0.04281 | 0.02541 | -1.68473 | 0.09204 | |
| **WL:ILP:MASC_noRT** | -0.04001 | 0.02691 | -1.48668 | 0.13710 | |
| **WL:ILP:MASC_VISUAL** | -0.09247 | 0.04291 | -2.15517 | 0.03115 | |
| **WL:ILP:MASC_MOTOR** | -0.13223 | 0.02470 | -5.35242 | 0.00000 | |
| **WL:ILP:MASC_ISR_C** | -0.03091 | 0.02354 | -1.31329 | 0.18909 | |
| **WL:ILP:MASC_ISR_1PC** | -0.01270 | 0.02678 | -0.47410 | 0.63543 | |
| **WL:ILP:VS_noRT** | 0.26818 | 0.05756 | 4.65941 | 0.00000 | |
| **WL:ILP:VS_RT** | 0.20093 | 0.02840 | 7.07412 | 0.00000 | |
| **WL:ILP:VS_RT_GA** | 0.34541 | 0.02039 | 16.94385 | 0.00000 | |
| **(2) MASC -(Intercept)** | -1.71607 | 0.09417 | -18.22282 | 0.00000 | |
| **WL** | 0.27659 | 0.02730 | 10.13004 | 0.00000 | |

| | | | | | |
|---|---|---|---|---|---|
| | ILP | -0.63831 | 0.04149 | -15.38361 | 0.00000 |
| | MASC_noRT | -0.24417 | 0.13462 | -1.81376 | 0.06972 |
| | MASC_VISUAL | -1.77483 | 0.15373 | -11.54546 | 0.00000 |
| | MASC_MOTOR | 0.28973 | 0.13200 | 2.19496 | 0.02817 |
| | MASC_ISR_C | 0.02813 | 0.13261 | 0.21216 | 0.83198 |
| | MASC_ISR_1PC | -0.17137 | 0.13401 | -1.27876 | 0.20098 |
| | VS_noRT | -0.85480 | 0.15458 | -5.52968 | 0.00000 |
| | VS_RT | 0.23886 | 0.13270 | 1.79990 | 0.07188 |
| | VS_RT_GA | 1.83479 | 0.13058 | 14.05080 | 0.00000 |
| | WL:ILP | -0.01886 | 0.02136 | -0.88313 | 0.37716 |
| | WL:MASC_noRT | -0.08594 | 0.04080 | -2.10644 | 0.03517 |
| | WL:MASC_VISUAL | -0.03917 | 0.06703 | -0.58430 | 0.55902 |
| | WL:MASC_MOTOR | -0.02210 | 0.03662 | -0.60341 | 0.54624 |
| | WL:MASC_ISR_C | -0.06670 | 0.03767 | -1.77040 | 0.07666 |
| | WL:MASC_ISR_1PC | 0.01295 | 0.03994 | 0.32416 | 0.74581 |
| | WL:VS_noRT | -0.13859 | 0.07399 | -1.87325 | 0.06103 |
| | WL:VS_RT | -0.18964 | 0.03960 | -4.78828 | 0.00000 |
| | WL:VS_RT_GA | -0.24599 | 0.03476 | -7.07635 | 0.00000 |
| | ILP:MASC_noRT | 0.04973 | 0.06051 | 0.82191 | 0.41113 |
| | ILP:MASC_VISUAL | -0.29883 | 0.08103 | -3.68779 | 0.00023 |
| | ILP:MASC_MOTOR | -0.11746 | 0.05717 | -2.05452 | 0.03993 |
| | ILP:MASC_ISR_C | 0.45977 | 0.05705 | 8.05933 | 0.00000 |
| | ILP:MASC_ISR_1PC | -0.06362 | 0.05993 | -1.06151 | 0.28846 |
| | ILP:VS_noRT | 0.10814 | 0.09096 | 1.18886 | 0.23450 |
| | ILP:VS_RT | 0.15539 | 0.05932 | 2.61953 | 0.00881 |
| | ILP:VS_RT_GA | -0.03385 | 0.05365 | -0.63081 | 0.52816 |
| | WL:ILP:MASC_noRT | 0.00324 | 0.03148 | 0.10280 | 0.91813 |
| | WL:ILP:MASC_VISUAL | -0.04928 | 0.04591 | -1.07352 | 0.28304 |
| | WL:ILP:MASC_MOTOR | -0.08893 | 0.02961 | -3.00349 | 0.00267 |
| | WL:ILP:MASC_ISR_C | 0.01239 | 0.02865 | 0.43230 | 0.66552 |
| | WL:ILP:MASC_ISR_1PC | 0.03063 | 0.03137 | 0.97624 | 0.32894 |
| | WL:ILP:VS_noRT | 0.31152 | 0.05984 | 5.20569 | 0.00000 |
| | WL:ILP:VS_RT | 0.24417 | 0.03277 | 7.45216 | 0.00000 |
| | WL:ILP:VS_RT_GA | 0.38868 | 0.02612 | 14.88072 | 0.00000 |
| **(3)** | **MASC_noRT -(Intercept)** | -1.95988 | 0.09628 | -20.35638 | 0.00000 |
| | WL | 0.19051 | 0.03032 | 6.28402 | 0.00000 |
| | ILP | -0.58789 | 0.04403 | -13.35101 | 0.00000 |
| | WL:ILP | -0.01627 | 0.02312 | -0.70356 | 0.48170 |
| **(4)** | **MASC_VISUAL -(Intercept)** | -3.48691 | 0.12138 | -28.72703 | 0.00000 |
| | WL | 0.23901 | 0.06112 | 3.91065 | 0.00009 |
| | ILP | -0.93352 | 0.06953 | -13.42646 | 0.00000 |
| | WL:ILP | -0.06873 | 0.04058 | -1.69366 | 0.09033 |
| **(5)** | **MASC_MOTOR -(Intercept)** | -1.42683 | 0.09253 | -15.41953 | 0.00000 |
| | WL | 0.25443 | 0.02440 | 10.42622 | 0.00000 |
| | ILP | -0.75555 | 0.03934 | -19.20654 | 0.00000 |
| | WL:ILP | -0.10805 | 0.02051 | -5.26836 | 0.00000 |
| **(6)** | **MASC_ISR_C (Intercept)** | -1.69011 | 0.09338 | -18.10009 | 0.00000 |
| | WL | 0.20922 | 0.02596 | 8.05941 | 0.00000 |
| | ILP | -0.17875 | 0.03915 | -4.56629 | 0.00000 |
| | WL:ILP | -0.00658 | 0.01910 | -0.34481 | 0.73024 |
| **(7)** | **MASC_ISR_1PC -(Intercept)** | -1.88773 | 0.09538 | -19.79187 | 0.00000 |
| | WL | 0.28952 | 0.02914 | 9.93409 | 0.00000 |

| | | | | | |
|---|---|---|---|---|---|
| | **ILP** | -0.70159 | 0.04326 | -16.21960 | 0.00000 |
| | **WL:ILP** | 0.01145 | 0.02298 | 0.49825 | 0.61831 |
| **(8)** | **VS_noRT -(Intercept)** | -2.56543 | 0.12251 | -20.94115 | 0.00000 |
| | **WL** | 0.13792 | 0.06862 | 2.00976 | 0.04446 |
| | **ILP** | -0.52588 | 0.08080 | -6.50844 | 0.00000 |
| | **WL:ILP** | 0.29192 | 0.05580 | 5.23154 | 0.00000 |
| **(9)** | **VS_RT -(Intercept)** | -1.47848 | 0.09349 | -15.81400 | 0.00000 |
| | **WL** | 0.08618 | 0.02868 | 3.00435 | 0.00266 |
| | **ILP** | -0.48280 | 0.04239 | -11.38847 | 0.00000 |
| | **WL:ILP** | 0.22484 | 0.02484 | 9.05059 | 0.00000 |
| **(10)** | **VS_RT_GA -(Intercept)** | 0.11579 | 0.09035 | 1.28156 | 0.20000 |
| | **WL** | 0.02980 | 0.02152 | 1.38501 | 0.16605 |
| | **ILP** | -0.67379 | 0.03401 | -19.80976 | 0.00000 |
| | **WL:ILP** | 0.36987 | 0.01504 | 24.59120 | 0.00000 |

GLMMs were fitted to a binary variable indicating whether a given word was refixated. The fixed structure included Data Set (10 levels, excluding CSL given the null refixation probability in 4- to 6-letter words), Word Length (WL; 5-9 letters; reference (mean) value: 6.86; 4-letter words were excluded due to floor effects in several data sets), Initial Landing Position (ILP; ranging between -5.6 and 0 letters relative to the center of words, thus excluding ILPs in the right halves of words which yielded near-zero refixation in MASC and most other models; reference (mean) value: -2.21 letters), and all interactions as predictors. The random structure included a random intercept by subject, as well as random effects of WL and ILP, but without their correlation; with a more complex random structure, GLMMs did not converge. Estimates and standard errors are expressed in logit unit; the estimated refixation probability in each data set when WL and ILP were at their reference value is given in the rightmost column (in Panel 1). Panel 1: Fixed-effects for the main GLMM model, with Data-Set reference level set to FSC. Panels 2-10: Data-Set reference level set to MASC, MASC_noRT, MASC_VISUAL, MASC_MOTOR, MASC_ISR_C, MASC_ISR_1PC, VS_noRT, VS_RT, and VS_RT_GA respectively. In Panels 3-10, only the intercept, the effects of WL and ILP, and the interaction were reported. These analyses relied on a total of 14451, 6931, 6176, 6642, 7599, 6749, 7008, 2511, 6045, 9153 cases across subjects in FSC, MASC, MASC_noRT, MASC_VISUAL, MASC_MOTOR, MASC_ISR_C, MASC_ISR_1PC, VS_noRT, VS_RT, and VS_RT_GA respectively.

**Supplementary Methods 3. The respective roles of inter-word spacing and print size.**

To determine the visual factors that are crucial for eye-movement guidance and test the general assumption in reading models (Extended Data Table 1) that only inter-word spacing, but not character-print size, matters[1,12,20-21], two additional tests were performed. First, MASC was presented with sentences from the FSC in three conditions of inter-word spacing: the normal spacing condition, and two others where inter-word spaces in the sentences were either filled or removed respectively (see Methods). Second, MASC's behavior over FSC sentences was tested for three screen-width angles resulting into three angular character sizes (0.25° as in the main analyses, and 0.51° and 1.03°). The effects of these manipulations were tested by fitting (G)LMMs to MASC's first-pass behavior, using the same procedure as in the above analyses (see Methods; Supplementary Methods 1-2), but without data set as a predictor. Since FSC readers only read the sentences in the normal-spacing and small-print-size (0.25°) conditions, MASC's predictions were compared to findings in the literature, when available, as well cross-study comparisons between spaced and unspaced languages using different font sizes. In the first series of analyses, inter-word spacing was entered as a categorical predictor with three levels, and in the second series of analyses, character size was entered as a continuous variable centered on its mean.

**MASC's predicted effects of inter-word spacing.**

Effects of inter-word spacing manipulations on eye movements during reading have been reported in a great number of studies. These effects, together with previously reported differences in oculomotor behavior between spaced and unspaced languages, have been interpreted as evidence for a central role of peripheral-word segmentation processes in eye-movement guidance. However, word-spacing manipulations are prone to several confounds and the role of word segmentation remains greatly debated[22,23]. MASC's oculomotor behavior as a function of inter-word spacing helps further disambiguating the origin of previously reported effects, while suggesting that differences in eye-movement behavior between spaced and unspaced languages cannot be attributed to differences in text segmentation.

**Saccade Length.** Numerous studies reported that the length of progressive saccades decreases drastically (by 1-3 letters on average) when spaces in normally spaced texts are globally removed[e.g.,24-28] or filled[25-26,29]. However, these effects were greatly reduced when confounding variables were controlled for, as illustrated in Extended Data Fig. 9a-b. First, whereas effects of space removal were found to be proportional to the reduction in line length that results from withdrawing spaces[24], space-filling manipulations, which allow keeping line length constant, overall yielded much smaller effects than space removal[25]. Furthermore, as revealed in several studies using gaze-contingent display changes, space-filling effects were even smaller, and often turned out to be negligible, when only peripheral spaces were filled and spaces to the right and/or left of the fixated word were preserved, regardless of peripheral linguistic content (normal vs. random or x-letters)[30-32]. Relatedly, the increase in fixation duration that generally accompanies space-filling effects was considerably reduced. These findings indicate that space-filling manipulations at best perturb online foveal word-identification, and/or fixation disengagement, rather than peripheral word segmentation and saccade targeting. In fact, when potential artifacts associated with visual-display changes during fixations[12,22], as well as the possibility of global reading-pace adjustments[22], were minimized by filling spaces (in the fovea and the periphery) during only a few random saccades, spacing effects nearly vanished[23]: Only were the proportion of small-amplitude forward saccades, associated with fixation behavior[33], and the duration of fixations, mildly inflated. Moreover, there was no change in the mean and the shape of the distributions of forward saccades when display changes also made the text meaningless (all letters replaced by random letters). Likewise, space addition between words in normally unspaced scripts was found to lengthen saccade length only modestly (effects <= 0.9 letters), nearly as little as extra spaces between nonwords[e.g.,34].

MASC lacks both (foveal) word-identification processes and fixation disengagement mechanisms, but it also has no idea of how unusual, and difficult, reading French text without spaces may be. It therefore predicted, in line with our literature review, only tiny effects of inter-word-spacing manipulations on the length of forward saccades.

In all three inter-word-spacing conditions, MASC's saccade-length distributions (Fig. 4a) were best fitted with two mixture components, associated respectively with regressive and progressive saccades. The estimated proportion of regressions remained very low, as in the above first-pass analyses (Supplementary Table 17), being numerically slightly greater in the normal and the removed-spacing conditions (0.09 and 0.07) than in the filled-spacing condition (0.02). Thus, only forward saccades were considered for analysis. Supplementary Table 29A-B presents the regression coefficients of LMs fitted respectively to the GMM-estimated mean and SD of the length of progressive saccades in the three spacing conditions. This indicates that forward saccades were on average shorter when spaces were removed than when they were preserved (estimate: -0.60630), while being slightly more variable (estimate: 0.07272). Moreover, forward saccades were slightly longer, though by only 1/6 letter, and slightly less variable (estimate: -0.19961) in the filled- compared to the normal-spacing condition.

Importantly, the reduction in saccade length with space removal was likely essentially due to lines being narrower in that condition. As shown in Supplementary Table 30, MASC, as FSC readers, made progressively shorter forward saccades as FSC sentences were shorter (estimates: 0.04780 and 0.05996; Panels 2,1). Considering that FSC sentences were on average 8.44 characters shorter when spaces were removed in comparison with the normal-spacing condition, it appears that about two third of the spacing effect was likely due to a confound with line length[24], thereby explaining why previously reported effects of space removal were overall much greater than space-filling effects[25] (Extended Data Fig. 9a-b). This, together with the negligible effect of space filling predicted by MASC, suggests that inter-word spacing has little impact on the length of forward saccades during reading beyond the effect due to line length in space-removal manipulations. Ongoing foveal processes, as well as global/offline adjustments (slowdown) of reading pace due to unusual and (seemingly) more difficult reading conditions, would come on top to modulate that effect. However, as suggested by the rather modest effects of unpredictable space-filling manipulations during the reading of English text[23] and space addition during the reading of unspaced languages[34], modulations by ongoing foveal-word identification processes would be only tiny, meaning that the larger reported effects of inter-word spacing were mainly artifactual. Consequently, the fact that Chinese/Japanese readers make much shorter forward saccades on average than Western readers (Extended Data Table 9a) cannot be attributed to the lack of inter-word spacing in ideographic scripts.

**Supplementary Table 29 | LM regression coefficients for the mean and SD of the length of MASC's progressive saccades by inter-word spacing.**

| A- PROGRESSIONS: MEAN | Estimate | Std. Error | t value | Pr(>\|t\|) |
|---|---|---|---|---|
| NORMAL -(Intercept) | 7.01648 | 0.01105 | 634.78855 | 0.00000 |
| FILLED | 0.16761 | 0.01563 | 10.72253 | 0.00000 |
| REMOVED | -0.60630 | 0.01563 | -38.78641 | 0.00000 |

| B- PROGRESSIONS: SD | Estimate | Std. Error | t value | Pr(>\|t\|) |
|---|---|---|---|---|
| NORMAL -(Intercept) | 3.09915 | 0.01625 | 190.69369 | 0.00000 |
| FILLED | -0.19961 | 0.02298 | -8.68499 | 0.00000 |
| REMOVED | 0.07272 | 0.02298 | 3.16405 | 0.00250 |

LMs were fitted separately to the GMM-estimated mean (A) and SD (B) of the positive mixture component in individual distributions of saccade lengths (in letters), with inter-word spacing as a categorical predictor (with three levels: NORMAL, FILLED, and REMOVED); Reference level set to NORMAL.

**Supplementary Table 30 | Fixed effects of LMMs for the length of progressive saccades by line length in MASC and FSC readers.**

| PROGRESSIONS: | Estimate | Std. Error | t value |
|---|---|---|---|
| **(1) FSC-(Intercept)** | 8.62949 | 0.18031 | 47.86048 |
| **MASC** | -1.41930 | 0.31044 | -4.57191 |
| **LL** | 0.05996 | 0.00316 | 18.97618 |
| **MASC: LL** | -0.01216 | 0.00312 | -3.89448 |
| **(2) MASC-(Intercept)** | 7.21019 | 0.25411 | 28.37390 |
| **LL** | 0.04780 | 0,00352 | 13.56281 |

LMMs were fitted to the length of forward saccades (in letters) in FSC readers and MASC. The fixed structure included Data Set (with two levels: FSC vs. MASC), Line Length (LL; 31-69 letters; reference (mean) value: 51.48 letters) and their interaction as predictors. The random structure included a random intercept by subject and sentence pair; with a random effect of line length by subject, the model did not converge. Panel 1: Fixed-effects of the main LMM, with Data-Set reference level set to FSC. Panel 2: Data-Set reference level set to MASC.

**Word-skipping behavior.** Very few studies tested the impact of inter-word-spacing manipulations on word-skipping behavior, and when word-skipping rate was reported this was across words of different lengths. Results overall showed that readers were less likely to skip words when spaces in normally spaced texts/sentences were globally removed[24-25,27,35-36] or filled[29,25], although these effects remained very small, being no greater than 11%, and most often ranging between 1 and 8%. Moreover, although similar trends were observed as a result of space addition in normally unspaced Chinese texts[37], other studies in unspaced (Thai and Japanese) languages showed the opposite trend[38-39].

As shown in Supplementary Table 31, MASC skipped short 4-letter words slightly more frequently when spaces were filled (53%), and even more so when they were removed (65%), than when they were preserved (48%; Panel 1; Fig.4b). This difference between spaced and unspaced conditions however reduced as words were longer, due to word-skipping rate decreasing faster with increasing word length in filled- and removed-spacing conditions compared to the normal-spacing condition (logit: -0.07251 and -0.07944; Panel 1); the effect of word length, significant in all three conditions (Panels 1-3), did not differ between filled and removed-spacing conditions ($p$ = 0.60475; Panel 2).

**Supplementary Table 31 | Fixed effects of GLMMs for MASC's word-skipping probability by word length and inter-word spacing.**

| SKIPPING BY WL | Estimate | Std. Error | z value | Pr(>\|z\|) | Proportion |
|---|---|---|---|---|---|
| **(1) NORMAL -(Intercept)** | -0.05666 | 0.03556 | -1.59340 | 0.11107 | 0.48584 |
| **WL** | -0.43407 | 0.00886 | -48.98466 | 0.00000 | |
| **FILLED** | 0.17483 | 0.02282 | 7.66026 | 0.00000 | 0.52951 |
| **REMOVED** | 0.66235 | 0.02413 | 27.45414 | 0.00000 | 0.64696 |
| **WL:FILLED** | -0.07251 | 0.01263 | -5.74082 | 0.00000 | |
| **WL:REMOVED** | -0.07944 | 0.01303 | -6.09565 | 0.00000 | |
| **(2) FILLED -(Intercept)** | 0.11816 | 0.03584 | 3.29716 | 0.00098 | |
| **WL** | -0.50658 | 0.00939 | -53.95580 | 0.00000 | |
| **REMOVED** | 0.48752 | 0.02448 | 19.91697 | 0.00000 | |
| **WL:REMOVED** | -0.00692 | 0.01338 | -0.51759 | 0.60475 | |
| **(3) REMOVED -(Intercept)** | 0.60569 | 0.03670 | 16.50544 | 0.00000 | |
| **WL** | -0.51350 | 0.00988 | -51.98508 | 0.00000 | |

GLMMs were fitted to a binary variable indicating whether a given word was skipped. The fixed structure included Inter-Word Spacing (categorical variable with 3 levels), Word Length (WL; 1-11 letters; reference (mean) value: 3.79 letters), and their interaction, as predictors. The random structure included a random intercept by subject and by sentence pair, as well as a random effect of WL; with in addition a random intercept by word, the model did not converge. Estimates and standard errors are expressed in logit unit; the estimated probability of word skipping in each spacing condition when WL was at its reference value is given in the rightmost column (in Panel 1). Panel 1: Fixed effects for the main GLMM, with the reference level for Inter-Word Spacing set to NORMAL. Panel 2-3: Inter-Word Spacing reference levels set to FILLED and REMOVED respectively.

A second GLMM was fitted to the data, using this time inter-word spacing, word length, and saccades' launch-site distance to the beginning of words as predictors of word-skipping rate. As shown in Supplementary Table 32, the probability of skipping short 3-letter words was again greater when spaces were filled (77%) than when they were preserved (69%; Panel 1), but it was no longer inflated when spaces were removed (69%; $p$ = 0.21147; Panel 1). Still, as in the above analyses without launch-site distance, the effect of word length, significant in all three spacing conditions (Panels 1-3), was stronger in unspaced conditions (logit: -0.14789 and -0.15805; Panel 1), thus implying that the likelihood of skipping longer words showed an even smaller effect of inter-word spacing. Most importantly, there was a decrease in word-skipping rate with increasing launch-site distance in all three conditions (logit: 0.76169, 0.80186 and 0.66782 in normal, filled, and removed-spacing conditions respectively; Panels 1-3), which was slightly greater and smaller in filled- and removed-spacing conditions respectively compared to the normal-spacing condition (logit: 0.04013 and -0.09384; Panel 1; Fig.4c). These mild differences attenuated with increasing word length, as suggested by the significant three-way interaction (logit: -0.04186 and 0.04063 respectively; Panel 1).

**Supplementary Table 32 | Fixed effects of GLMMs for MASC's word-skipping probability by word length, launch site, and inter-word spacing.**

| SKIPPING BY LS | | Estimate | Std. Error | z value | Pr(>\|z\|) | Proportion |
|---|---|---|---|---|---|---|
| **(1)** | **NORMAL -(Intercept)** | 0.82443 | 0.05223 | 15.78367 | 0.00000 | 0.69518 |
| | **WL** | -0.63346 | 0.01766 | -35.87358 | 0.00000 | |
| | **LS** | 0.76169 | 0.01483 | 51.34407 | 0.00000 | |
| | **FILLED** | 0.41479 | 0.03393 | 12.22568 | 0.00000 | 0.77543 |
| | **REMOVED** | -0.04176 | 0.03342 | -1.24954 | 0.21147 | 0.68625 |
| | **WL:LS** | 0.05347 | 0.01223 | 4.37377 | 0.00001 | |
| | **WL:FILLED** | -0.14789 | 0.02480 | -5.96333 | 0.00000 | |
| | **WL:REMOVED** | -0.15805 | 0.02808 | -5.62818 | 0.00000 | |
| | **LS:FILLED** | 0.04013 | 0.02109 | 1.90294 | 0.05705 | |
| | **LS:REMOVED** | -0.09384 | 0.02028 | -4.62815 | 0.00000 | |
| | **WL:LS:FILLED** | -0.04186 | 0.01704 | -2.45681 | 0.01402 | |
| | **WL:LS:REMOVED** | 0.04063 | 0.01708 | 2.37847 | 0.01738 | |
| **(2)** | **FILLED -(Intercept)** | 1.23884 | 0.05375 | 23.04797 | 0.00000 | |
| | **WL** | -0.78129 | 0.01881 | -41.52548 | 0.00000 | |
| | **LS** | 0.80186 | 0.01566 | 51.20830 | 0.00000 | |
| | **REMOVED** | -0.45638 | 0.03562 | -12.81260 | 0.00000 | |
| | **WL:LS** | 0.01166 | 0.01238 | 0.94243 | 0.34597 | |
| | **WL:REMOVED** | -0.01022 | 0.02876 | -0.35515 | 0.72248 | |
| | **LS:REMOVED** | -0.13406 | 0.02084 | -6.43147 | 0.00000 | |
| | **WL:LS:REMOVED** | 0.08244 | 0.01718 | 4.79738 | 0.00000 | |
| **(3)** | **REMOVED -(Intercept)** | 0.78242 | 0.05318 | 14.71211 | 0.00000 | |
| | **WL** | -0.79145 | 0.02277 | -34.75082 | 0.00000 | |
| | **LS** | 0.66782 | 0.01432 | 46.62033 | 0.00000 | |
| | **WL:LS** | 0.09408 | 0.01227 | 7.66908 | 0.00000 | |

GLMMs were fitted to a binary variable indicating whether a given word was skipped. The fixed structure included Inter-Word Spacing (3 levels), Word Length (WL; 2-6 letters; reference (mean) value: 3.15), Launch-Site distance (LS; 0-7 letters relative to the beginning of words; reference (mean) value: -3.13 letters) and all interactions as predictors. The random structure included a random intercept by subject and sentence pair, as well as random effects of WL and LS by subject; with, in addition, a random intercept by word, the model did not converge. Estimates and standard errors are expressed in logit units; the estimated probability of word skipping in each spacing condition, when WL and LS were at their reference (mean) value, is given in the rightmost column (in Panel 1). Panel 1: Fixed effects for the main GLMM, with the reference level for Inter-Word Spacing set to NORMAL. Panel 2-3: Inter-Word Spacing reference level set to FILLED and REMOVED, respectively.

Thus, in line with previous reports, MASC predicted only tiny differences in word-skipping rate as a function of inter-word spacing, and even more so as words were longer. The comparison however remains difficult as neither word length nor launch site were controlled in previous studies. The fact yet is that spacing effects were not reported to vary with language-related variables (i.e., word frequency), except in one single study[29]. Thus, it is quite unlikely that these effects reflected top-down, language-related, modulations of eye-movement behavior, but this will need to be further investigated at the light of MASC's predictions. Importantly, predicted and observed effects of inter-word spacing on the likelihood of word skipping were much smaller than differences in skipping rate between naturally spaced and unspaced languages (Extended Data Fig. 8a). Relatedly, MASC predicted a slightly greater effect of word length on the likelihood of word skipping when inter-word spaces were removed/filled, but this did not compare with the much stronger relationship between word length and word skipping in Chinese compared to Western languages. Therefore, it is quite unlikely that inter-word spacing is the critical factor accounting for differences in word-skipping behavior between spaced and unspaced languages.

**The PVL effect.** Studies investigating the effects of inter-word spacing manipulations on eye-movement behavior quite consistently reported that saccades tend to land slightly closer to the beginning of words in unsegmented-text conditions[26-27,35,37-39, but 24,38-39]. These tiny effects, together with the tendency for readers of unspaced languages to preferentially fixate the words' very-first characters (Extended Data Fig. 8d), have been taken as main arguments for the assumption that word segmentation is crucial for eye-movement guidance[26]. MASC suggests an alternative interpretation for effects of inter-word spacing as well as inter-language differences, by showing that visual word segmentation is unnecessary for the PVL effect observed in spaced alphabetic languages.

In all three inter-word spacing conditions, MASC's within-word landing-position distributions were best fitted with a single mixture component. The proportion of bimodal distributions was null for 4- and 5-letter words, and slightly higher for longer, 6- to 9-letter, words, but with no consistent difference between the three conditions (Normal: 0, 0.30, 0.25, and 0 respectively; Filled: 0.15, 0.05, 0.10 and 0.05; Removed: 0, 0.30, 0.20 and 0). In 6- to 9-letter words, the largest mixture component still accounted on average for 88%-100% of the data. The fixed effects of LMMs fitted to the GMM-estimated mean of the largest mixture component are presented in the left panels of Supplementary Table 33. This shows that MASC initially fixated on average a position slightly to the left of the center of 6-letter words in the normal-spacing condition (estimate: -0.11335; Panel 1), as well as in filled- and removed-spacing conditions (estimates: -0.10855 and -0.11124, respectively; Panels 2-3). This leftward fixation bias did not significantly differ between the three conditions (t <= 0.06963; Panels 1-2). Still, as words were longer, MASC's saccades landed progressively closer to the words' beginning (estimate: -0.08034; Panel 1), and even more so when spaces were removed (estimate: -0.04961; Panel 1). In these conditions, saccades thus undershot slightly more the center of 7- to 9-letter words than in the normal-spacing condition (Fig. 4d). In the filled-spacing condition, to the contrary, the effect of word length, though also significant (estimate: -0.06596; Panel 2), was no different from that predicted in the normal-spacing condition ($t$ = 0.37957; Panel 1), thus leading to a comparable leftward fixation bias in filled- and normal-spacing conditions, regardless of word length. In addition, the spread of within-word landing position distributions, as reported in the right panels of Supplementary Table 33, was comparable across all three inter-word spacing conditions ($|t| <= 0.30966$; Panel 1). Moreover, it significantly increased with word length (estimates: 0.20097, 0.22205 and 0.201316; Panels 1-3) at the same rate regardless of inter-word spacing ($t <= 1.21412$).

**Supplementary Table 33 | Fixed effects of LMMs for the mean and SD of MASC's within-word landing sites by word length and inter-word spacing**.

| PVL EFFECT: MEAN / SD | Estimate | Std. Error | *t* value | Estimate | Std. Error | *t* value |
|---|---|---|---|---|---|---|
| **(1) NORMAL -(Intercept)** | -0.11335 | 0.04873 | -2.32592 | 1.64748 | 0.02097 | 78.57416 |
| **WL** | -0.08034 | 0.02678 | -2.99964 | 0.20097 | 0.01228 | 16.36918 |
| **FILLED** | 0.00480 | 0.06892 | 0.06963 | 0.00918 | 0.02965 | 0.30966 |
| **REMOVED** | 0.00211 | 0.06892 | 0.03063 | -0.00449 | 0.02965 | -0.15153 |
| **WL:FILLED** | 0.01438 | 0.03788 | 0.37957 | 0.02108 | 0.01736 | 1.21412 |
| **WL:REMOVED** | -0.04961 | 0.03788 | -1.30989 | 0.00220 | 0.01736 | 0.12651 |
| **(2) FILLED -(Intercept)** | -0.10855 | 0.04873 | -2.22744 | 1.65666 | 0.02097 | 79.01207 |
| **WL** | -0.06596 | 0.02678 | -2.46284 | 0.22205 | 0.01228 | 18.08619 |
| **REMOVED** | -0.00269 | 0.06892 | -0.03900 | -0.01368 | 0.02965 | -0.46119 |
| **WL:REMOVED** | -0.06399 | 0.03788 | -1.68946 | -0.01888 | 0.01736 | -1.08761 |
| **(3) REMOVED -(Intercept)** | -0.11124 | 0.04873 | -2.28300 | 1.64299 | 0.02097 | 78.35986 |
| **WL** | -0.12995 | 0.02678 | -4.85201 | 0.20316 | 0.01228 | 16.54809 |

LMMs were fitted separately to the GMM-estimated mean (left panel) and SD (right panel) of the largest mixture component in individual landing position distributions. In both LMMs, the fixed structure included Inter-Word Spacing (3 levels), Word Length (WL; 4-9 letters; reference (mean) value: 6.5 letters), and their interaction as predictors. The random structure included both a random intercept and a random effect of WL by subject. Estimates and standard errors are expressed in letters. Panel 1: Fixed effects for the main LMM, with the reference level for Inter-Word Spacing set to NORMAL. Panel 2-3: Inter-Word Spacing reference level set to FILLED and REMOVED, respectively.

Thus, in line with the previously reported leftward shift in within-word landing positions during the reading of unsegmented text materials[26-27,35,37-39, but 24,38-39], MASC predicted a tiny shift in landing positions towards the beginning of long words when spaces were removed. Similar effects were reported for space-filling manipulations[26,29], but MASC did not replicate them, thus suggesting that its predicted effect of space removal was again essentially due to lines being narrower when spaces were removed. Under the assumption that saccades are guided towards the centers of peripherally selected target word(-object)s, the leftward bias in within-word landing positions in unspaced texts was attributed to difficulties in peripheral word segmentation and saccade targeting. MASC, because it relies on computation of a visual saliency map, does extract word-boundary information, but only in spaced texts. The simple fact it predicted a PVL effect regardless of inter-word spacing indicates to the contrary that visual word segmentation is unnecessary for eye-movement guidance. Previously reported effects of inter-word spacing on within-word landing positions[26-27,35,37-39] were as tiny as observed variations in within-word landing positions with linguistic variables[7] (Extended Data Fig. 2). Moreover, they were found essentially during the reading of naturally spaced languages[38]. These effects therefore very likely reflected top-down modulations of saccade amplitude, resulting from difficulties in (para)foveal word identification in unsegmented texts. On the other hand, the fact that predicted and observed effects of inter-word spacing were much smaller than previously reported differences in the PVL effect between alphabetic and ideographic scripts (Extended Data Fig. 8c) indicates that word segmentation cannot be the (sole) explanation for cross-language differences. In fact, as shown in Extended Data Fig. 9f-g, these differences were the mere artefact of character-print size being larger in unspaced- compared to spaced-language studies and landing positions being classically binned into letter-/character-based intervals regardless of print size.

**The Launch-Site effect.** To our knowledge, previous studies did not investigate whether inter-word spacing manipulations affect the launch-site effect. However, our literature review of word-based phenomena across studies and languages suggests that this effect is quite comparable between spaced[6] and unspaced languages[40-43] (Extended Data Fig. 8d-e). As summarized in the left panels of Supplementary Table 34, MASC predicted nearly no difference in the launch-site effect as a function of inter-word spacing (Fig. 4e). Saccades' overall landing-position distributions, all best fitted with a single mixture component (see Table legend), peaked to the right of the center of 4-letter words in the normal-spacing condition (estimate: 1.61521; Panel 1), and only slightly less or slightly more to the right in removed- and filled-spacing conditions respectively (estimates: -0.14954 and 0.19369; Panel 1). This rightward fixation bias decreased with increasing word length (estimates: -0.29311, -0.34899, and 0.34128, in normal-, filled-, and removed-spacing conditions; Panels 1-3) and increasing launch-site distance (estimates: 0.79472, 0.81142, and 0.80375 in normal-, filled-, and removed-spacing conditions; Panels 1-3). The effect of word length was slightly greater in both filled and removed-spacing conditions compared to the normal-spacing condition (estimates: -0.05589 and -0.04817; Panel 1). In contrast, the effect of launch site was not significantly different between the three conditions for 4-letter words ($t <= 1.27704$; Panels 1-2), and only mildly smaller in the filled- compared to the normal-spacing condition in longer words (estimate: -0.03418; Panel 1).

**Supplementary Table 34 | Fixed effects of LMMs for the mean and SD of MASC's overall landing-site distributions by word length, launch site, and inter-word spacing.**

| LS EFFECT: MEAN / SD | Estimate | Std. Error | *t* value | Estimate | Std. Error | *t* value |
|---|---|---|---|---|---|---|
| **(1) NORMAL -(Intercept)** | 1.61521 | 0.02223 | 72.64432 | 1.72217 | 0.01359 | 126.71433 |
| **WL** | -0.29311 | 0.00957 | -30.64162 | -0.07141 | 0.00585 | -12.21302 |
| **LS** | 0.79472 | 0.00908 | 87.53469 | 0.04940 | 0.00565 | 8.74131 |
| **FILLED** | 0.19369 | 0.03181 | 6.08958 | -0.00272 | 0.01944 | -0.13994 |
| **REMOVED** | -0.14954 | 0.03979 | -3.75833 | 0.01398 | 0.02432 | 0.57491 |
| **WL:LS** | 0.02761 | 0.00459 | 6.02029 | -0.01233 | 0.00280 | -4.39665 |
| **WL:FILLED** | -0.05589 | 0.01378 | -4.05454 | -0.01255 | 0.00843 | -1.48977 |
| **WL:REMOVED** | -0.04817 | 0.01950 | -2.47077 | -0.00540 | 0.01192 | -0.45333 |
| **LS:FILLED** | 0.01670 | 0.01308 | 1.27704 | -0.02342 | 0.00814 | -2.87930 |
| **LS:REMOVED** | 0.00902 | 0.01696 | 0.53193 | 0.00914 | 0.01048 | 0.87203 |
| **WL:LS:FILLED** | -0.03418 | 0.00655 | -5.22156 | -0.00342 | 0.00400 | -0.85447 |
| **WL:LS:REMOVED** | -0.01441 | 0.00865 | -1.66484 | 0.00637 | 0.00529 | 1.20450 |
| **(2) FILLED -(Intercept)** | 1.80889 | 0.02274 | 79.53512 | 1.71945 | 0.01390 | 123.68149 |
| **WL** | -0.34899 | 0.00992 | -35.16677 | -0.08396 | 0.00607 | -13.84104 |
| **LS** | 0.81142 | 0.00941 | 86.22535 | 0.02598 | 0.00585 | 4.44005 |
| **REMOVED** | -0.34322 | 0.04007 | -8.56457 | 0.01670 | 0.02450 | 0.68187 |
| **WL:LS** | -0.00657 | 0.00467 | -1.40588 | -0.01575 | 0.00286 | -5.51648 |
| **WL:REMOVED** | 0.00771 | 0.01967 | 0.39210 | 0.00715 | 0.01203 | 0.59447 |
| **LS:REMOVED** | -0.00768 | 0.01714 | -0.44779 | 0.03256 | 0.01059 | 3.07553 |
| **WL:LS:REMOVED** | 0.01977 | 0.00870 | 2.27294 | 0.00979 | 0.00532 | 1.84147 |
| **(3) REMOVED -(Intercept)** | 1.46567 | 0.03300 | 44.41990 | 1.73616 | 0.02017 | 86.08254 |
| **WL** | -0.34128 | 0.01699 | -20.08872 | -0.07681 | 0.01038 | -7.39697 |
| **LS** | 0.80375 | 0.01433 | 56.09558 | 0.05854 | 0.00882 | 6.63498 |
| **WL:LS** | 0.01321 | 0.00734 | 1.79948 | -0.00596 | 0.00449 | -1.32806 |

LMMs were fitted separately to the GMM-estimated mean (left panel) and SD (right panel) of the largest mixture component in individual landing-position distributions. The largest mixture component accounted for 100% of the data, except in three cases (1-letter words FILLED, 6-letter words NORMAL, and 7-letter words REMOVED) where this accounted on average for 99%, 98%, and 95% of the data (proportion of two mixture components: 0.04, 0.05, and 0.12 respectively). In both LMMs, the fixed structure included Inter-Word Spacing (3 levels), Word Length (WL; 1-9 letters; reference (mean) value: 4.37 letters), Launch Site (LS; in two-letter bins and ranging between -11 and -1 letters relative to the beginning of words; reference (mean) value: -3.82 letters), and all interactions as predictors. The random structure included a random intercept as well as random effects of WL and LS by subject. Estimates and standard errors are expressed in letters. Panel 1: Fixed effects for the main LMM, with the reference level for Inter-Word Spacing set to NORMAL. Panel 2-3: Inter-Word Spacing reference level set to FILLED and REMOVED, respectively.

For the estimated SD, as presented in the right panels of Supplementary Table 34, this was comparable between the three conditions of inter-word spacing ($|t|$ <= 0.68187; Panels 1-2). Moreover, in all conditions, the SD mildly decreased with increasing word length (estimates: -0.07141, -0.08396, and -0.07681, in normal-, filled-, and removed-spacing conditions respectively; Panels 1-3) and increasing launch-site distance (estimates: 0.04940, 0.02598, and 0.05854, respectively; Panels 1-3). The effect of word length was comparable across all three conditions ($|t|$ <= 1.48977; Panel 1). For the effect of launch site, it was only slightly weaker in the filled-spacing condition compared to both normal- and removed-spacing conditions (estimates: -0.02342 and -0.03256; Panels 1-2).

In summary, MASC predicted nearly no variations in the launch-site effect with inter-word spacing, thus indicating that this effect, just as the PVL effect, does not require visual word segmentation.

**The Refixation-OVP effect.** Previous studies showed that within-word refixations are overall more likely in unsegmented-text conditions[22,26,29,35,37], even though the Refixation-OVP effect remains largely unaffected by inter-word spacing manipulations[26,37]. As shown in Supplementary Table 35, MASC predicted only a slight increase in the refixation rate for 6-letter words in the removed-spacing condition compared to the normal-spacing condition (11% vs. 9%; Panel 1), and a decrease rather than an increase in refixation likelihood in the filled spacing condition (6%; Panel 1). This contrasts with previous studies showing effects of space removal/filling in the order of 6-27%[26,29,35,37]. Still, in all three conditions, the likelihood of within-word refixations increased with increasing word length (logit: 0.28904, 0.29420, and 0.23885, in normal-, filled-, and removed-spacing conditions; Panels 1-3), and as the initial landing position deviated towards the beginning of words (logit: -0.56885, -0.93478, and -0.69379, respectively; Panels 1-3). The only notable difference was the mild strengthening in the left wing of the OVP effect in the filled-spacing condition (logit: -0.36593), and to some extent also in the removed-spacing condition (logit: -0.12494, $p = 0.08393$), compared to the normal-spacing condition (Fig. 4f).

**Supplementary Table 35 | Fixed effects of GLMMs for MASC's within-word refixation probability by initial landing position, word length, and inter-word spacing**.

| Refixation-OVP EFFECT | Estimate | Std. Error | z value | Pr(>\|z\|) | Proportion |
|---|---|---|---|---|---|
| (1) NORMAL -(Intercept) | -2.25573 | 0.04720 | -47.78689 | 0.00000 | 0.09486 |
| WL | 0.28904 | 0.02846 | 10.15585 | 0.00000 | |
| ILP | -0.56885 | 0.05363 | -10.60637 | 0.00000 | |
| FILLED | -0.49841 | 0.07679 | -6.49076 | 0.00000 | 0.05985 |
| REMOVED | 0.22031 | 0.06367 | 3.46037 | 0.00054 | 0.11553 |
| WL:ILP | -0.05195 | 0.02697 | -1.92611 | 0.05409 | |
| WL:FILLED | 0.00516 | 0.04630 | 0.11136 | 0.91133 | |
| WL:REMOVED | -0.05019 | 0.03808 | -1.31819 | 0.18744 | |
| ILP:FILLED | -0.36593 | 0.08576 | -4.26700 | 0.00002 | |
| ILP:REMOVED | -0.12494 | 0.07229 | -1.72833 | 0.08393 | |
| WL:ILP:FILLED | -0.01018 | 0.04298 | -0.23679 | 0.81282 | |
| WL:ILP:REMOVED | -0.00208 | 0.03693 | -0.05620 | 0.95518 | |
| (2) FILLED -(Intercept) | -2.75414 | 0.06057 | -45.47372 | 0.00000 | |
| WL | 0.29420 | 0.03652 | 8.05596 | 0.00000 | |
| ILP | -0.93478 | 0.06692 | -13.96856 | 0.00000 | |
| REMOVED | 0.71872 | 0.07412 | 9.69704 | 0.00000 | |
| WL:ILP | -0.06212 | 0.03346 | -1.85651 | 0.06338 | |
| WL:REMOVED | -0.05535 | 0.04442 | -1.24592 | 0.21279 | |
| ILP:REMOVED | 0.24099 | 0.08263 | 2.91660 | 0.00354 | |
| WL:ILP:REMOVED | 0.00810 | 0.04191 | 0.19328 | 0.84674 | |
| (3) REMOVED -(Intercept) | -2.03542 | 0.04272 | -47.64240 | 0.00000 | |
| WL | 0.23885 | 0.02530 | 9.44231 | 0.00000 | |
| ILP | -0.69379 | 0.04847 | -14.31430 | 0.00000 | |
| WL:ILP | -0.05402 | 0.02524 | -2.14073 | 0.03230 | |

GLMMs were fitted to a binary variable indicating whether a given word was refixated. The fixed structure included Inter-Word Spacing (3 levels), Word Length (WL; 4-9 letters; reference (mean) value: 6.41), Initial Landing Position (ILP; ranging between –4.6 and 0 letters relative to the center of words, thus excluding ILP in the right half of the words which yielded near-zero refixations; reference (mean) value: -1.59 letters), and all interactions as predictors. The random structure included a random intercept, as well as random effects of WL and ILP, by subject; with a more complex random structure, GLMMs did not converge. Estimates and standard errors are expressed in logit unit; estimated refixation probability in each inter-word spacing condition when WL and ILP were at their reference value is given in the rightmost column (in Panel 1). Panel 1: Fixed effects for the main GLMM, with the reference level for Inter-Word Spacing set to NORMAL. Panel 2-3: Inter-Word Spacing reference level set to FILLED and REMOVED, respectively.

In sum, MASC predicted only mild differences in the likelihood of within-word refixations between normal-, filled-, and removed-spacing conditions. This suggests that previously reported effects of inter-word spacing, which were overall larger, mainly reflected difficulties in ongoing foveal-word processing and/or fixation disengagement. MASC however predicted only mild variations in the Refixation-OVP effect with inter-word spacing, in line with previous findings. This indicates that inter-word spacing cannot be the (sole) explanation for differences in the OVP effect between spaced and unspaced languages (Extended Data Fig. 8f).

**MASC's predicted effects of character size.**

It has long been assumed that saccades during reading are programmed in character coordinates regardless of print properties, hence the conventional use of letters/characters as a metric unit for readers' oculomotor behavior[1,12,20-21,44]. However, as revealed by our literature review, many studies showed differences in eye-movement behavior as a function of angular print size (Extended Data Fig. 9c-d). Existing models of oculomotor control during reading cannot account for these effects (Extended Data Table 1). They also cannot (reasonably) account for differences in eye-movement behavior between spaced and unspaced languages, which, as we have seen above, cannot be (solely) attributed to text segmentation. The fact yet is that there were great differences in character print size across studies and languages. These differences were never considered as a possible cause for differences in eye-movement behavior between spaced and unspaced languages. MASC, which does predict previously reported changes in eye-movement behavior with character-print size, explains data across languages using only print size as a predictor.

**Saccade Length.** The general assumption that character-print size does not matter for eye-movement guidance was established only based on a couple influential studies suggesting non-significant variations in the numbers of letters traversed with viewing distance, and hence angular print size[20-21,44]. However, many other studies were conducted which did reveal significant effects of font size on the character count per saccade. As illustrated in Extended Data Fig. 9c, the great majority of studies suggested an overall decrease in the length of progressive saccades expressed in numbers of letters with increasing angular character size, but other studies reported inverted U-shaped curves. These inconsistent patterns were however simply the effect of saccades being expressed in letters. When the same data were replotted after converting saccade amplitudes to degrees of visual angle, effects of print size were far more consistent: In all studies, there was a positive linear relationship between forward saccade length and character size (Extended Data Fig. 9d). Importantly, slopes only mildly differed between studies, and they were not systematically different between spaced and unspaced languages. MASC predicted these relationships.

As we have seen above, MASC mostly moved forward during its first pass over FSC sentences in the normal 0.25° print-size condition, making regressions essentially from the end of the sentences. This was even more true for larger print sizes: in the 0.51° condition, 50% of the distributions were estimated with a single mixture component, while 100% of the distributions were unimodal in the 1.03° condition (Fig. 4g). Thus, only forward saccades were analyzed. The fixed effects of LMs fitted to the GMM-estimated mean and SD of the length of forward saccades (in letters) are presented in Supplementary Table 36A-B. This shows that MASC predicted a decrease in the length of forward saccades with increasing print size. This effect did not compare with the above-reported tiny effect of inter-word spacing (Supplementary Table 29, Fig. 4a). Indeed, for every one-degree increment in character size, the length of saccades decreased by about 1.9 letters, while becoming much less variable (estimate: -1.83237), in line with findings showing a reduction in saccade amplitude with larger characters[10,45-47,but 20-21,44]. Yet, MASC's predicted slopes could hardly be compared to previously reported effects of print size on the character count per saccade due to great variability across studies (Extended Data Fig. 9c).

However, saccade length, when expressed in degrees of visual angle, was found to increase with increasing angular print size in all studies and languages (Extended Data Fig. 9d). MASC predicted a similar relationship, its curve falling in between the curves from different studies and languages. As shown in Supplementary Table 36C-D, its saccades increased by as much as about 5.02° with every one-degree increment in character size, while becoming slightly more variable (estimate: 0.34211). This suggests that saccades are programmed in degrees of visual angle, and not in letters/characters as commonly assumed. As revealed by cross-language comparisons, Chinese/Japanese readers traverse on average much less characters than readers of Western alphabetic languages (Extended Data Fig. 9a,c). The fact yet is that these populations were tested with much larger print sizes (0.60-1.52° compared to 0.25-0.35° in spaced-language studies). Thus, it is quite likely that differences in forward saccade length between spaced and unspaced languages were for a great part due to differences in angular print size, rather than differences in text segmentation.

**Supplementary Table 36 | LM regression coefficients for the mean and SD of the length of MASC's progressive saccades by character size.**

| A- PROGRESSIONS: MEAN (letters) | Estimate | Std. Error | t value | Pr(>\|t\|) |
|---|---|---|---|---|
| Intercept | 6.45910 | 0.01737 | 371.82689 | 0.00000 |
| CS | -1.94524 | 0.05357 | -36.31266 | 0.00000 |

| B- PROGRESSIONS: SD (letters) | Estimate | Std. Error | t value | Pr(>\|t\|) |
|---|---|---|---|---|
| (Intercept) | 2.27121 | 0.03119 | 72.82592 | 0.00000 |
| CS | -1.83237 | 0.09617 | -19.05281 | 0.00000 |

| C- PROGRESSIONS: MEAN (degrees) | Estimate | Std. Error | t value | Pr(>\|t\|) |
|---|---|---|---|---|
| Intercept | 3.67041 | 0.02700 | 135.93077 | 0.00000 |
| CS | 5.02037 | 0.08327 | 60.29123 | 0.00000 |

| D- PROGRESSIONS: SD (degrees) | Estimate | Std. Error | t value | Pr(>\|t\|) |
|---|---|---|---|---|
| (Intercept) | 0.83776 | 0.00197 | 425.65566 | 0.00000 |
| CS | 0.34211 | 0.00607 | 56.36649 | 0.00000 |

LMs were fitted separately to the GMM-estimated mean (A,C) and SD (B,D) of the positive peak in individual distributions of saccade lengths, expressed in letters (A-B) or in degrees of visual angle (C-D). Character size (CS) was entered as a continuous predictor centered on its mean (0.60°).

**Word-skipping behavior.** It was previously found that the probability of word skipping tends to decrease with increasing print size[10,45-46]. To test whether MASC predicted that effect, a first GLMM was fitted to simulated data using word length (in letters) and character size as predictors. As shown in Supplementary Table 37A, MASC did predict a reduction in the likelihood of word skipping with increasing angular print size (logit: -2.32753; Fig 4h). Its word-skipping rate additionally varied with word length, being lower for longer words (logit: -0.67567), and even more so as characters were larger (logit: -0.59660).

**Supplementary Table 37 | Fixed effects of GLMMs for MASC's word-skipping probability by word length and character size.**

| A- SKIPPING BY WL (letters) | Estimate | Std. Error | z value | Pr(>\|z\|) | Proportion |
|---|---|---|---|---|---|
| **Intercept** | -1.00834 | 0.02925 | -34.47723 | 0.00000 | 0.26730 |
| **WL** | -0.67567 | 0.00770 | -87.76303 | 0.00000 | |
| **CS** | -2.32753 | 0.04841 | -48.07749 | 0.00000 | |
| **WL:CS** | -0.59660 | 0.02342 | -25.46935 | 0.00000 | |

| B- SKIPPING BY WL (degrees) | Estimate | Std. Error | z value | Pr(>\|z\|) | Proportion |
|---|---|---|---|---|---|
| **Intercept** | -2.10680 | 0.04563 | -46.16899 | 0.00000 | 0.10844 |
| **WL** | -1.67463 | 0.03345 | -50.06929 | 0.00000 | |
| **CS2(0.51°)** | 1.40165 | 0.04900 | 28.60223 | 0.00000 | |
| **CS4(1.03°)** | 1.92499 | 0.04887 | 39.39266 | 0.00000 | |
| **WL:CS2(0.51°)** | 0.47672 | 0.03864 | 12.33661 | 0.00000 | |
| **WL:CS4(1.03°)** | 0.85727 | 0.03686 | 23.25409 | 0.00000 | |

GLMMs were fitted to a binary variable indicating whether a given word was skipped. A: The fixed structure included Character Size (CS; 0.25-1.03°; reference (mean) value: 0.65°), Word Length (WL; 1-11 letters; reference (mean) value: 4.02 letters), and the interaction between CS and WL as predictors. The random structure included a random intercept by subject and by sentence pair, as well as a random effect of WL by subject; with also a random intercept by word, the model did not converge. B: Given the correlation between character size and the words' spatial extent, the fixed structure included Character Size as a categorical variable (CS1: 0.25 (the reference); CS2: 0.51°; CS3: 1.03°), Word Length (WL; 0.25-6.63°; reference (mean) value: 2.16°), and the interaction between CS and WL as predictors. The random structure included both a random intercept and a random effect of WL by subject. Estimates and standard errors are expressed in logit unit; the estimated probability of word skipping when WL and CS were at their reference value is given in the rightmost column.

To our knowledge, there has been no report in the literature of the relationship between the likelihood of word skipping and word length as a function of print size. However, this relationship was reported in many studies using respectively different print sizes. MASC's predicted differences in the effect of word length between character sizes of 0.25° and 1.03° nearly perfectly matched the differences observed between spaced and unspaced languages using font sizes of about 0.25-0.35° and 1.52° respectively (Extended Data Fig. 8a). This indicates that character size indeed strongly affects the relationship between word skipping and word length, while already arguing for an interpretation of cross-language differences in terms of print size rather than inter-word spacing –recall that inter-word-spacing manipulations only very mildly influenced word-skipping behavior by word length, and much less than print size (Fig 4b,h, Supplementary Table 31). The additional fact that the effect of print size reversed (i.e., word-skipping rate was greater and varied less largely with word length for larger characters) when word length was expressed in degrees of visual angle (Supplementary Table 37B), just as for Chinese-reading data in comparison with spaced-alphabetic languages (Extended Data Fig. 9e), confirms the predominant role of character print size at the expense of inter-word spacing.

MASC's predicted effects of character size were further investigated by fitting a second GLMM to simulated data, with word length, saccades' launch-site distance to the space in front of the words, and character size as predictors. As reported in Supplementary Table 38, word-skipping rate again decreased with increasing word length and character size (logit: -1.66844 and -5.53199), with the effect of word length being stronger for larger characters (logit: -2.46884). In addition, word skipping became less likely as saccades were launched from further away (logit: 1.83959), and even more so as characters were larger (logit: 2.74064) and words longer (logit: 0.12913); the three-way interaction was also significant (logit: 0.17156). Future studies should test these additional predictions. Moreover, if, as suggested above, character print size is responsible for differences in word-skipping behavior between spaced and unspaced languages, then the influence of launch site on word-skipping rate should be stronger in ideographic scripts, when tested using larger characters.

**Supplementary Table 38 | Fixed effects of GLMM for MASC's word-skipping probability by word length, launch site, and character size.**

| SKIPPING BY LS | Estimate | Std. Error | z value | Pr(>\|z\|) | Proportion |
|---|---|---|---|---|---|
| Intercept | -1.36660 | 0.07988 | -17.10790 | 0.00000 | 0.20317 |
| WL | -1.66844 | 0.03026 | -55.12984 | 0.00000 | |
| LS | 1.83959 | 0.02180 | 84.39485 | 0.00000 | |
| CS | -5.53199 | 0.21446 | -25.79448 | 0.00000 | |
| WL:LS | 0.12913 | 0.01509 | 8.55598 | 0.00000 | |
| WL:CS | -2.46884 | 0.09161 | -26.94866 | 0.00000 | |
| LS:CS | 2.74064 | 0.06750 | 40.60157 | 0.00000 | |
| WL:LS:CS | 0.17156 | 0.04373 | 3.92327 | 0.00009 | |

A GLMM was fitted to a binary variable indicating whether a given word was skipped. The fixed structure included Character Size (CS; 0.25-1.03°; reference (mean) value: 0.65°), Word Length (WL; 2-6 letters; reference (mean) value: 3.21 letters), saccades' Launch-Site distance to the space in front of the words (LS; 0-7 letters; reference (mean) value: -2.62 letters), and all interactions as predictors. The random structure included a random intercept by subject and sentence pair, as well as random effects of WL and LS by subject; with also a random intercept by word, the model did not converge. Estimates and standard errors are expressed in logit units; the estimated probability of word skipping when WL, LS, and CS were at their reference value are given in the rightmost column.

**The PVL effect.** In the framework of predominant word-based models of eye-movement control during reading, the PVL effect is a central phenomenon, which results from saccades being invariably aimed at the center of target words combined with SRE, a bias to traverse a constant, optimal, number of letters. This effect should therefore hold regardless of print properties/size and be affected only in conditions where peripheral word segmentation and saccade-targeting mechanisms are more difficult, such as in the absence of inter-word spacing. However, as we have seen above, the PVL effect shows only tiny variations with inter-word spacing[26-27,35,37-39, but 24,38-39] (Supplementary Table 33), and the much-greater differences in the PVL effect between spaced and unspaced languages (Extended Data Fig. 8c) call for an alternative interpretation, especially as print size was much greater in Chinese/Japanese studies compared to studies in spaced-alphabetic languages. Only a few studies investigated the PVL effect as a function of print size but results overall showed, in contradiction with word-based models, that saccades tend to land on average closer to the words' beginning when characters are larger[10,45-46]. MASC's predictions were consistent with these findings, while suggesting that cross-language differences in the PVL effect were attributable to print size.

MASC's predicted distributions of within-word landing positions, expressed in letters relative to the center of words, were best fitted with a single mixture component, regardless of print size. The proportion of bimodal distributions was overall greater in longer, 6- to 9-letter words (0-0.45), than in 4- and 5-letter words (between 0 and 0.05), but this did not vary consistently with character size (6-letter words: 0.25, 0.35 and 0.45; 7-letter words: 0.35, 0.15, and 0; 8-letter words: 0.20, 0, and 0.15; 9-letter words: 0, 0, and 0.45; respectively for 0.25°, 0.51° and 1.03° character sizes). Moreover, across all conditions, the largest mixture component accounted on average for 81% to 100% of the data. The fixed effects of LMMs fitted to the estimated mean and SD of the largest mixture component are presented respectively in the left and right panels of Supplementary Table 39A. These indicate that MASC's initial landing positions, again biased to the left of the words' center (estimate: -1.01082 for 6-letter words and an average print size of 0.6°), deviated further towards the beginning of words as word length increased (estimate: -0.24465) and, even more radically, as character size increased. For every one-degree increment in print size, landing positions shifted by about 1.2 letters towards the beginning of 6-letter words, and even more so as words were longer (e.g., by about 2.56 letters in 9-letter words), as suggested by the significant interaction between character size and word length (estimate: -0.45208), while being also much less widely scattered (estimates: -0.43923 and -0.26414 respectively). Thus, whereas MASC predicted very little variations in within-word landing positions with inter-word spacing (Supplementary Table 33, Fig. 4d), it predicted drastic changes with character size, as further illustrated in Fig. 4j.

Studies in unspaced languages revealed for the great majority that Chinese/Japanese readers tend to fixate quite systematically towards the very beginning of words regardless of their length[e.g.,43], thus in contrast with the typical PVL effect reported for Western alphabetic languages (Extended Data Fig. 8c). MASC's substantial predicted effect of print size on within-word landing positions suggests that these differences were largely due to characters in Chinese and Japanese studies being on average two to four times greater in size than in spaced-language studies (0.6°-1.5° and 0.25-0.40 respectively). It follows that cross-language differences should reduce when landing positions are expressed in degrees of visual angle and words are matched in angular extent. Tsai and McConkie[42] directly compared the distributions of initial landing positions in English and Chinese words of comparable angular size, using angular bin sizes of 0.25°. They found, in line with our prediction, great overlap between the distributions, and in particular no marked preference for the very beginning of Chinese words, unlike previous studies using letters as a metric; the only difference was that the distributions were overall flatter for Chinese compared to English. Further evidence for our assumption is provided in Extended Data Fig. 9f-g, where within-word landing-position distributions were replotted for words of comparable angular extents across studies in different languages and print sizes; landing positions originally reported in letters were converted to degrees of visual angle and binned into either small, medium, or large angular intervals. Distributions were narrower and peaked closer to the words' beginning as bin size increased and the number of bins decreased, regardless of language. This is a trivial observation, but this yet explains why most Chinese/Japanese studies using (half-)character bins (or 0.6-1° bins) suggested a fixation preference for the very-beginning of words in contrast with space-languages studies using one-letter bins and hence 0.25-0.40° bins. Moreover, for a given bin size and word extent, all distributions were very-much alike regardless of language and print size; again, the only notable difference was that the distributions tended to be flatter for larger print sizes (notably above 0.6°)[42]. As illustrated in Extended Data Fig. 9f-g and further confirmed by statistical analyses (see below),

MASC's predicted distributions showed the same trends, being nearly indistinguishable from observed distributions.

**Supplementary Table 39 | Fixed effects of LMMs for the mean and SD of MASC's within-word landing sites by word length and character size**.

| A- PVL EFFECT: MEAN / SD (letters) | Estimate | Std. Error | *t* value | Estimate | Std. Error | *t* value |
|---|---|---|---|---|---|---|
| **Intercept** | -1.01082 | 0.03107 | -32.53193 | 1.74959 | 0.01763 | 99.21760 |
| **WL** | -0.24465 | 0.01819 | -13.44681 | 0.13529 | 0.00930 | 14.54597 |
| **CS** | -1.20859 | 0.09582 | -12.61330 | -0.43923 | 0.05438 | -8.07727 |
| **WL:CS** | -0.45208 | 0.05610 | -8.05773 | -0.26414 | 0.02868 | -9.20906 |

| B- PVL EFFECT: MEAN / SD (degrees) | Estimate | Std. Error | *t* value | Estimate | Std. Error | *t* value |
|---|---|---|---|---|---|---|
| **(1) Intercept (CS: 0.25°)** | -0.15082 | 0.01076 | -14.0216 | 0.43340 | 0.00721 | 60.13269 |
| **WL** | -0.07114 | 0.04067 | -1.74904 | 0.25700 | 0.01750 | 14.68699 |
| **CS2(0.51°)** | -0.06642 | 0.01617 | -4.10784 | 0.07420 | 0.01057 | 7.01702 |
| **CS4(1.03°)** | -0.22674 | 0.01929 | -11.7569 | 0.15554 | 0.01187 | 13.10438 |
| **WL:CS2(0.51°)** | 0.10585 | 0.05785 | 1.82987 | -0.00542 | 0.02495 | -0.21751 |
| **WL:CS4(1.03°)** | 0.15044 | 0.06186 | 2.43216 | 0.01181 | 0.02736 | 0.43160 |
| **(2) Intercept (CS2: 0.51°)** | -0.21724 | 0.01207 | -17.9931 | 0.50760 | 0.00774 | 65.60647 |
| **WL** | 0.03471 | 0.04113 | 0.84392 | 0.25157 | 0.01778 | 14.14974 |
| **CS4(1.03°)** | -0.16032 | 0.02005 | -7.99575 | -0.08134 | 0.01220 | 6.66844 |
| **WL:CS4(1.03°)** | 0.04459 | 0.06216 | 0.71734 | 0.01723 | 0.02754 | 0.62579 |
| **(3) Intercept (CS4: 1.03°)** | -0.37756 | 0.01601 | -23.5857 | 0.58895 | 0.00943 | 62.45047 |
| **WL** | 0.07930 | 0.04660 | 1.70170 | 0.26881 | 0.02103 | 12.78062 |

LMMs were fitted separately to the GMM-estimated mean (left panel) and SD (right panel) of the largest mixture component in individual within-word landing position distributions (A: in letters, B: in degrees) for different word lengths and character sizes. A: In both LMMs, the fixed structure included Word Length (WL; 4-9 letters; reference (mean) value: 6.5 letters), character print size (CS: 0.25-1.03°; reference (mean) value: 0.60°), and their interaction as predictors. The random structure included both a random intercept and a random effect of WL by subject. Estimates and standard errors are expressed in letters. B: In both LMMs, the fixed structure included Word Length (WL; 0.25-2.55°; reference (mean) value: 1.4°), character print size as a categorical predictor given the correlation between character size and the words' spatial extent (CS: 0.25° (the reference); CS2: 0.51°; CS4: 1.03°), and their interaction as predictors. The random structure included both a random intercept and a random effect of WL by subject. Estimates and standard errors are expressed in degrees. Words larger than 2.55° were excluded from analysis due to the very small proportion of single detected peaks in landing-position distributions notably for the largest character size (CS2 (0.51°): 1, 1, 1, 1, 0.85, 0.5, 0.85, 1, and 1 for 0.51-, 1.02-, 1.53-, 2.04-, 2.55, 3.06-, 3.57-, 4.08-, and 4.59-degree words respectively; CS4(1.03°): 1, 0.75, 0.15, 0.05, 0.1, 0.3, 0.6, 0.35, and 0.5 for 1.02-, 2.06-, 3.09-, 4.12-, 5.15-, 6.18-, 7.21-, 8.24-, and 9.27-degree words respectively).

MASC's predicted distributions of within-word landing positions, expressed in degrees of visual angle, were best fitted with a single mixture component, at least for words subtending no more than 2.55 degrees of visual angle (Supplementary Table 39B legend). LMMs were fitted to the estimated mean and SD of the largest mixture component only for this subset of words using both word length (in degrees of visual angle) and character size as predictors. As shown in the left panels of Supplementary Table 39B, mean within-word landing positions did not vary with word length ($|t|$ <= -1.74904; Panels 1-3). Moreover, they only very mildly shifted towards the words' beginning with increasing character size (estimates: -0.06642 and -0.22674 for 0.51° and 1.03° characters; Panel 1), and even less so as words were longer (estimates: 0.10585 and 0.15044 for 0.51° and 1.03° characters; Panel 1). Most importantly, as indicated in the right panels of Supplementary Table 39B, the SD increased not only with increasing word length as in the above letter-based analyses (estimates: 0.25700, 0.25157, and 0.26881 for 0.25°, 0.51°, and 1.03° characters respectively; Panels 1-3), but also with increasing character size (estimates: 0.07420 and 0.15554 for 0.51° and 1.03° characters; Panel 1) regardless of word length ($t$ <= 0.43160; Panel 1). This resulted in very large distributions for large-printed words, thus in line with the rather flat distributions reported by Tsai and McConkie[42] during Chinese reading (see also Extended Data Fig. 9f-g).

**The Launch Site effect.** The launch-site effect, just like the PVL effect, is a central phenomenon for word-based models of eye-movement control during reading. Because it is assumed to reflect a compromise between a word-center saccade-targeting strategy and SRE[6], it should remain unaffected by print size. It was yet found, in contradiction with this assumption that the relationship between saccades' launch site and mean landing sites during the reading of French words is weaker for large-printed words[10]. MASC replicated this pattern. Its overall landing-site distributions were all best fitted with a single mixture component. As suggested by the fixed effects of LMMs fitted to the GMM-estimated mean, presented in the left panel of Supplementary Table 40, MASC tended to fixate on average closer to the beginning of large-printed words (estimate: -2.68262), and even more so as words were longer (estimate: -0.20403), in line with the above-reported within-word landing-position analyses (Supplementary Table 39A). Its saccades also landed progressively closer to the words' beginning as their launch-site distance to the space in front of the words increased (estimate: 0.87331). Importantly, this relationship did not vary with character size in the case of 4-letter words ($t$ = 0.72670), but it did for longer words, becoming slightly weaker with increasing print size (estimate: -0.01972), thus in agreement with previous findings[10] (Fig. 4k). Moreover, as shown in the right panel of Supplementary Table 40, MASC's landing positions were less widely spread as character size increased (estimate: -1.07157), and even more so as words were shorter (estimate: 0.08143) and less eccentric (estimate: -0.04418), also as previously found[10].

**Supplementary Table 40 | Fixed effects of LMMs for the mean and SD of MASC's overall landing-site distributions by word length, launch site, and character size.**

| LS EFFECT: MEAN / SD | Estimate | Std. Error | *t* value | Estimate | Std. Error | *t* value |
|---|---|---|---|---|---|---|
| Intercept | -0.01228 | 0.01017 | -1.20815 | 1.12065 | 0.02027 | 55.29046 |
| WL | -0.42189 | 0.00399 | -105.76005 | -0.03260 | 0.00275 | -11.85356 |
| LS | 0.87331 | 0.00447 | 195.21139 | 0.04999 | 0.00258 | 19.37081 |
| CS | -2.68262 | 0.03049 | -87.97737 | -1.07157 | 0.06206 | -17.26580 |
| WL:LS | 0.03377 | 0.00186 | 18.15822 | 0.01172 | 0.00117 | 10.04903 |
| WL:CS | -0.20403 | 0.01154 | -17.68551 | 0.08143 | 0.00805 | 10.11444 |
| LS:CS | 0.00961 | 0.01322 | 0.72670 | -0.04418 | 0.00759 | -5.82226 |
| WL:LS:CS | -0.01972 | 0.00541 | -3.64403 | 0.01760 | 0.00340 | 5.17482 |

LMMs were fitted separately to the GMM-estimated mean (left panel) and SD (right panel) of the largest, unique, mixture component in individual overall landing-position distributions. All distributions were indeed best fitted with a single mixture component; this accounted therefore for 100% of the data in all conditions. In both LMMs, the fixed structure included Word Length (WL; 1-10 letters; reference (mean) value: 4.53 letters), Launch Site (LS; in two-letter bins and ranging between -9 and -1 letters relative to the space in front of the words; reference (mean) value: -3.66 letters), Character Size (CS; 0.25-1.03°; reference (mean) value: 0.63°), and all interactions as predictors. The random structure included a random intercept as well as random effects of WL and LS by subject, but without the correlation between random effects for the SD. Estimates and standard errors are expressed in letters.

Only a few studies investigated the launch-site effect in unspaced languages[42-43], and even less so for saccades' overall landing positions[40-41]. Moreover, estimated slopes for the linear relationship between saccades' launch site and mean landing site, when they were reported, were quite variable across studies. It thus remains unclear whether there are inter-language differences in the launch-site effect. MASC suggests that there should be none when character size is controlled.

**The Refixation-OVP effect.** Previous studies did not investigate whether the Refixation-OVP effect varies with print size. However, our literature review of word-based eye-movement behavior across studies and languages revealed that this effect, or at least the left wing of this effect, tends to be stronger in unspaced languages (Extended Data Fig.8f). As we have seen above, this cannot be due to differences in text segmentation (Supplementary Table 35), but this could again be the effect of character size, that was much greater in unspaced-language studies. In line with this assumption, MASC predicted a strengthening of the OVP effect with increasing print size. GLMM estimates of MASC's within-word refixation probability are reported in Supplementary Table 41. These first indicate that refixations were more likely in longer words as well as large-printed words (logit: 0.71534 and 0.18587 respectively). Most importantly, the increase in the refixation rate as the initial fixation location deviated from the center of words (logit: -1.62435) was stronger as characters were larger (logit: -2.15317).

**Supplementary Table 41 | Fixed effects of GLMM for MASC's within-word refixation probability by initial landing position, word length, and character size**.

| Refixation-OVP effect | Estimate | Std. Error | z value | Pr(>\|z\|) | Proportion |
|---|---|---|---|---|---|
| Intercept | -2.49017 | 0.09435 | -26.39133 | 0.00000 | 0.07655 |
| WL | 0.71534 | 0.01982 | 36.08928 | 0.00000 | |
| ILP | -1.62435 | 0.03482 | -46.65461 | 0.00000 | |
| CS | 0.18587 | 0.27851 | 0.66738 | 0.50453 | |
| WL:ILP | 0.09061 | 0.01792 | 5.05731 | 0.00000 | |
| WL:CS | 0.93197 | 0.05216 | 17.86613 | 0.00000 | |
| ILP:CS | -2.15317 | 0.09608 | -22.40969 | 0.00000 | |
| WL:ILP:CS | 0.20274 | 0.04834 | 4.19353 | 0.00003 | |

A GLMM was fitted to a binary variable indicating whether a given word was refixated. The fixed structure included Word Length (WL; 4-9 letters; reference (mean) value: 6.34), Initial Landing Position (ILP; ranging between –5.6 and 0 letters from the center of words, thus excluding ILP in the right half of words which yielded near-zero refixations; reference (mean) value: -2.06 letters), Character Size (CS; 0.25-1.03°; reference (mean) value: 0.66), and all interactions as predictors. The random structure included a random intercept as well as random effects of WL and ILP by subject; with a more complex random structure, the GLMM did not converge. Estimates and standard errors are expressed in logit unit; the estimated refixation probability when WL, ILP, and CS were at their reference value is given in the rightmost column.

Thus, the reason the (left wing of the) Refixation-OVP effect tends to be greater in unspaced ideographic languages compared to spaced alphabetic languages very likely came from differences in character print sizes, just as inter-language differences in saccade length, word-skipping behavior, and (within-word) landing positions.

### Supplementary References

1. Rayner, K. Eye movements in reading and information processing: 20 years of research. *Psychol. Bull.* **124(3)**, 372-422 (1998).
2. Rayner, K. & McConkie, G.W. What guides a reader's eye movements? *Vision Res.* **16**, 829-837 (1976).
3. Brysbaert, M., & Vitu, F. in *Eye guidance in reading and scene perception* (ed. Underwood, G.) 125-147 (Elsevier Science Ltd, Oxford, 1998).
4. Kerr, P.W. *Eye movement control during reading: The selection of where to send the eyes.* University of Illinois at Urbana-Champaign (1992).
5. Rayner, K. Eye guidance in reading: Fixation location within words. *Perception* **8**, 21-30 (1979).
6. McConkie, G.W., Kerr, P.W., Reddix, M.D. & Zola, D. Eye movement control during reading: I. The location of initial eye fixations on words. *Vision Res.* **28(10)**, 1107-1118 (1988).
7. Albrengues, C., Lavigne, F., Aguilar, C., Castet, E. & Vitu, F. Linguistic processes do not beat visuo-motor constraints, but they modulate where the eyes move regardless of word boundaries: Evidence against top-down word-based eye-movement control during reading. *PLoS ONE* **14(7)**, e0219666 (2019).
8. Krügel A, & Engbert R. On the launch-site effect for skipped words during reading. *Vision Res.* **50**, 1532-1539 (2010).
9. Krügel A, Vitu F, & Engbert R. Fixation positions after skipping saccades: A single space makes a large difference. *Attention Percept. & Psychophys.* **74(8)**,1556-1561 (2012).

10. Yao-N'Dré, M., Castet, E. & Vitu, F. Inter-word eye behavior during reading is not invariant to character size: Evidence against systematic saccadic range error in reading. *Vis. Cogn.* **22(3-4)**, 415-440 (2014).
11. McConkie, G.W., Kerr, P. W., Reddix, M.D., Zola, D. & Jacobs, A.M. Eye movement control during reading: II. Frequency of refixating a word. *Percept. Psychophys.* **46**, 245-253 (1989).
12. O'Regan, J.K. in *Eye movements and their role in visual and cognitive processes* (ed. Kowler, E.) 395-453 (Elsevier, Amsterdam, 1990).
13. O'Regan, J. K., Lévy-Schoen, A., Pynte, J., & Brugaillère, B. Convenient fixation location within isolated words of different lengths and structures. *J. Exp. Psychol. Human* **10**, 250-257 (1984).
14. Vitu, F., O'Regan, J.K. & Mittau, M. Optimal landing position in reading isolated words and continuous text. *Percept. Psychophys.* **47(6)**, 583-600 (1990).
15. Andriessen, J.J. & De Voogd, A.H. Analysis of eye movement patterns in silent reading. *IPO Annual Progress Report* **8**, 29-34 (1973).
16. Vitu, F. & McConkie, G.W. in *Reading as a perceptual process* (eds. Kennedy, A., Radach, R., Heller, D. & Pynte, J.) 301-326 (Elsevier, Oxford, 2000).
17. Vitu, F., McConkie, G.W. & Zola, D. in *Eye guidance in reading and scene perception* (ed. Underwood, G.) 101-124 (Elsevier Science Ltd, Oxford, 1998).
18. Radach, R. & McConkie, G.W. in *Eye guidance in reading and scene perception* (ed. Underwood, G.) 77-100 (Elsevier Science Ltd, Oxford, 1998).
19. Vitu, F., O'Regan, J.K., Inhoff, A. & Topolski, R. Mindless reading: Eye movement characteristics are similar in scanning strings and reading texts, *Percept. Psychophys.* **57**, 352-364 (1995).
20. Huey, E.B. *The Psychology and Pedagogy of Reading*. (Macmillan, New York, 1908).

21. Morrison, R.E. & Rayner, K. Saccade size in reading depends upon character spaces and not visual angle. *Percept. Psychophys.* **30**, 395–396 (1981).

22. Epelboim, J., Booth, J.R. & Steinman, R.M. Much ado about nothing: the place of space in text. *Vision Res.* **36(3)**, 465-470 (1996).

23. Yang, S.-N. & McConkie, G.W. Saccade generation during reading: Are words necessary? *Eur. J. Cogn. Psychol.* **16(1/2)**, 226-261 (2004).

24. Epelboim, J., Booth, J.R. & Steinman, R.M. Reading unspaced text: Implications for theories of reading eye movements. *Vision Res.* **34**, 1735-1766 (1994).

25. McGowan, V.A., White, S.J., Jordan, T.R. & Paterson, K.B. Aging and the use of interword spaces during reading: Evidence from eye movements. *Psychon. Bull. Rev.* **21(3)**, 740-747 (2014).

26. Rayner, K., Fischer, M.H. & Pollatsek, A. Unspaced text interferes with both word identification and eye movement control. *Vision Res.* **38**, 1129-1144 (1998).

27. Rayner, K., Yang, J., Schuett, S., & Slattery, T. Eye movements of older and younger readers when reading unspaced text. *Exp. Psychol.* **60(5)**, 354-361 (2013).

28. Shen, D. et al. Eye movements of second language learners when reading spaced and unspaced Chinese text. *J. Exp. Psychol. Appl.* **18**, 192–202 (2017).

29. Sheridan, H., Rayner, K. & Reingold, E.M. Unsegmented text delays word identification: Evidence from a survival analysis of fixation durations. *Vis. Cogn.* **21(1)**, 38-60 (2013).

30. McConkie, G.W. & Rayner, K. The span of the effective stimulus during a fixation in reading. *Percept. Psychophys.* **17**, 578-586 (1975).

31. Morris, R.K., Rayner, K. & Pollatsek, A. Eye movement guidance in reading: The role of parafoveal letter and space information. *J. Exp. Psychol. Hum. Percept. Perform.* **16**, 268-281 (1990).

32. Pollatsek, A. & Rayner, K. Eye movement control in reading: The role of word boundaries. *J. Exp. Psychol. Hum. Percept. Perform.* **8(6)**, 817-833 (1982).

33. Vitu, F., Lancelin, D., Jean, A., & Farioli, F. Influence of foveal distractors on saccadic eye movements: A dead zone for the global effect. *Vision Res*. **46**, 4684-4708 (2006).

34. Bai, X.J., Yan, G.L., Liversedge, S.P., Zang, C.L. & Rayner, K. Reading spaced and unspaced Chinese text: Evidence from eye movements. *J. Exp. Psychol. Hum. Percept. Perform.* **34(5)**, 1277–1287 (2008).

35. Mirault, J., Snell, J. & Grainger, J. Reading without spaces revisited: The role of word identification and sentence-level constraints. *Acta Psychol.* **195**, 22-29 (2019).

36. Perea, M. & Acha, J. Space information is important for reading. *Vision Res.* **49**, 1994–2000 (2009).

37. Zang, C., Liang, F., Bai, X., Yan, G. & Liversedge, S.P. Interword spacing and landing position effects during Chinese reading in children and adults. *J. Exp. Psychol. Hum. Percept. Perform.* **39(3)**, 720-734 (2013).

38. Winskel, H., Radach, R. & Luksaneeyanawin, S. Eye movements when reading spaced and unspaced Thai and English: A comparison of Thai-English bilinguals and English monolinguals. *J. Mem Lang.* **61**, 339-351 (2009).

39. Sainio, M., Hyönä, J., Bingushi, K. & Bertram, R. The role of interword spacing in reading Japanese: An eye movement study. *Vision Res.* **47**, 2575–2584 (2007).

40. Liu, Y., Reichle, E.D. & Li, X. The effect of word frequency and parafoveal preview on saccade length during the reading of Chinese. *J. Exp. Psychol. Hum. Percept. Perform.* **42**, 1008–1025 (2016).

41. Liu, Y., Huang, R., Gao, D. & Reichle, E.D. Further tests of a dynamic-adjustment account of saccade targeting during the reading of Chinese. *Cogn. Sci.* **41(6)**, 1624-1287 (2017).

42. Tsai, J.L. & McConkie, G.W. in *The mind's eye: Cognitive and applied aspects of eye movement research* (eds. Hyönä, J., Radach, R. & Deubel, H.) 159-176 (Elsevier, Oxford, 2003).

43. Yan, M., Kliegl, R., Richter, E.M., Nuthmann, A. & Shu, H. Flexible saccade-target selection in Chinese reading. *Q. J. Exp. Psychol.* **63**, 705–725 (2010).

44. O'Regan, J.K., Lévy-Schoen, A. & Jacobs, A. The effect of visibility on eye movement parameters in reading. *Percept. Psychophys.* **34**, 457-464 (1983).

45. Shu, H., Zhou, W., Yan, M. & Kliegl, R. Font size modulates saccade-target selection in Chinese reading. *Atten. Percept. Psychophys.* **73**, 482–490 (2011).

46. Yan, M., Zhou, W., Shu, H. & Kliegl, R. Perceptual span depends on font size during the reading of Chinese sentences. *J. Exp. Psychol. Learn. Mem. Cogn.* ***41*****(1)**, 209–219 (2015).

47. Yen, N.-S., Tsai, J.-L., Chen, P.-L., Lin, H.-Y. & Chen, A.L.P. Effects of typographic variables on eye-movement measures in reading Chinese from a screen. *Behav. Inform. Technol.* **30(6)**, 797-808 (2011).
